# Substrate recognition and selectivity in SARS-CoV-2 main protease: Unveiling the role of subsite interactions through dynamical nonequilibrium molecular dynamics simulations

**DOI:** 10.1101/2023.12.01.569046

**Authors:** H. T. Henry Chan, A. Sofia F. Oliveira, Adrian J. Mulholland, Christopher J. Schofield, Fernanda Duarte

## Abstract

The main protease (M^pro^) of the SARS-CoV-2 coronavirus employs a cysteine-histidine dyad in its active site to catalyse hydrolysis of the viral polyproteins. It is well established that binding of the substrate P1-Gln in the S1 subsite of M^pro^ active site is crucial for catalysis and this interaction has been employed to inform inhibitor design; however, how M^pro^ dynamically recognises and responds to substrate binding remains difficult to probe by experimental methods. We thus employed the dynamical nonequilibrium molecular dynamics (D-NEMD) approach to probe the response of M^pro^ to systematic substrate variations. The results emphasise the importance of P1-Gln for initiating a productive enzymatic reaction. Specifically, substituting P1-Gln with alanine disrupts the conformations of the Cys145 and His41 dyad, causing Cys145 to transition from the productive *gauche* conformation to the non-productive *trans* conformation. Importantly, our findings indicate that M^pro^ exhibits dynamic responses to substrate binding and likely to substrate-mimicking inhibitors within each of the S4-S2′ subsites. The results inform on the substrate selectivity requirements and shed light on the observed variations in hydrolytic efficiencies of M^pro^ towards different substrates. Some interactions between substrate residues and enzyme subsites involve more induced fit than others, implying that differences in functional group flexibility may optimise the binding of a substrate or inhibitor in a particular subsite.

## Introduction

Since the onset of the global pandemic, the SARS-CoV-2 coronavirus main protease (M^pro^) has attracted considerable attention as a target for COVID-19 treatment. M^pro^, which is encoded as the non-structural protein 5 gene (nsp5) in the viral polyprotein, is a cysteine protease that cleaves the viral polyproteins at specific sites to release the individual nsps.^1-4^ As these nsps are essential for viral maturation and replication, M^pro^ inhibition is an effective antiviral strategy.^2-4^ The main protease is highly conserved among coronaviruses, with 96% sequence identity between the SARS-CoV and SARS-CoV-2 M^pro^s. By contrast, M^pro^ has low sequence similarity with human proteases, reducing the risk of side effects caused by its inhibition.^5, 6^ Various M^pro^ inhibitors such as nirmatrelvir (PF-07321332 in Paxlovid)^7^ and ensitrelvir (S-217622 in Xocova)^8^ have been developed for COVID-19 treatment.

The predominantly homodimeric M^pro^ employs an active site catalytic dyad, composed of the nucleophilic Cys145 and general base His41,^9-13^ to enable selective hydrolysis at eleven sites in the viral polyproteins. M^pro^ selectively cleaves its substrates between the P1 and P1′ residues in the consensus sequence: [P4=Ala/Val/Pro/Thr]-[P3=X]-[P2=Leu/Phe/Val]-[P1=Gln]|[P1′=Ser/Ala/Asn], where “|” denotes the cleavage site and X any proteinogenic amino acid.^12, 14-19^ The P4-P1′ residues bind in the corresponding S4-S1′ subsites on M^pro^. The presence of Gln at P1 is a conserved feature of M^pro^ substrates of SARS-CoV-2 and other coronaviruses.^18, 20^ We have used a mass spectrometry-based assay to monitor SARS-CoV-2 M^pro^ catalysed hydrolysis of the eleven polyprotein-derived peptide substrates, all of which possess a P1-Gln residue but display different turnover efficiencies by M^pro^, and which demonstrated M^pro^ inhibition by multiple P1-Gln-containing peptides.^21^ Indeed, the S1 pocket is a binding hotspot for potent M^pro^ inhibitors,^21-23^ and has been targeted using various groups, including unmodified Gln,^17, 21, 24^ Gln sidechain-mimicking cycloglutamines (γ-lactams)^5-7, 25, 26^ and δ-lactams,^27^ His,^28^ and heterocycles such as triazoles.^8^ Some inhibitors position hydrophobic groups at the P1 equivalent position, including a Met residue,^29^ a cyclobutylmethyl group (boceprevir), a propyl group (telaprevir), and a butyl group (narlaprevir); the latter three compounds display inhibition in the µM concentration range.^30-33^ Note, however, that in some cases, the relatively high reactivity of a covalently reacting warhead (*e.g.* aldehyde or α-ketoamide), or the irreversibility of the covalent addition, likely compensates for non-optimal binding in the S1 pocket (and elsewhere).^29^

Despite this apparent promiscuity of S1 binding groups, M^pro^ selectively hydrolyses substrates with a P1 Gln during polyprotein processing. Investigating how M^pro^ dynamically recognises the P1-Gln residue and how this relates to binding in other subsites on either side of the hydrolysed amide is important in understanding how M^pro^ achieves substrate selectivity.^34^ Previous efforts to elucidate protease-substrate interactions by biophysical methods, principally by low temperature crystallography, have been insufficient to investigate the dynamics of these interactions.

We report dynamical-nonequilibrium molecular dynamics (D-NEMD)^35-38^ simulations to investigate the responses of M^pro^ to an instantaneous substitution of active site cleft binding substrate residues with Ala residues. D-NEMD simulations has been successfully employed to investigate a variety of dynamic processes in biomolecular systems, including signal propagation and allosteric effects arising from ligand and substrate binding, and pH-induced changes, in nicotinic acetylcholine receptors,^39, 40^ β-lactamases,^41^ and the SARS-CoV-2 spike protein.^42-45^ Recently, we used D-NEMD simulations to investigate changes in SARS-CoV-2 M^pro^ following removal of a peptide substrate from a noncovalent complex, which identified allosteric effects.^46^ In the D-NEMD approach, the system of interest is firstly simulated using equilibrium molecular dynamics (MD); the resultant configurations are used to initiate multiple short nonequilibrium MD simulations where a perturbation is applied. The response of the system to the perturbation is quantified by comparing the equilibrium and nonequilibrium propagations. Using the Kubo-Onsager relation,^37, 38^ the response to the perturbation is evaluated by averaging responses obtained from multiple (tens to hundreds) pairs of equilibrium and nonequilibrium simulations. Such averaging enables one to assess the significance of the response by statistical measures,^37^ such as the standard error of the mean (SEM).^46^

Out of the eleven native M^pro^ substrate sequences, we selected two representatives for investigation (**Figure 1**). One was the nsp4/5 P6-P5′ 11-mer substrate (TSAVLQ|SGFRK, “s01”), which exhibits the most efficient hydrolysis of studied peptides by M^pro^.^21^ The second was the nsp8/9 substrate (SAVKLQ|NNELS, “s05”), which is conserved in coronaviruses^18^ and whose P4-P1 sequence manifests optimal activity amongst screened proteinogenic amino acid derived sequences.^17^ Our combined D-NEMD/alanine-substitution study, akin to traditional alanine scanning in experimental^47-49^ and computational studies,^50-53^ provides insights into the role of the original residue in these substrates while minimising disruption to the peptide fold and backbone conformation. Alanine substitutions were performed at each residue in the P4-P2′ region, including positions for which M^pro^ demonstrates apparent selectivity (P4, P2, P1, and P1′) and those displaying little apparent selectivity (P3 and P2′). We also performed Ala substitutions at P2 for the nsp5/6 substrate “s02” (SGVTFQ|SAVKR), and our designed inhibitory peptide “p12” (KYTFWQ|YSQFY) (**Figure 1**).^21^ Analysis of s02 and p12 enables us to probe differences in the structural responses with an alternative residue (Phe or Trp) in the S2 subsite. The dynamical response of M^pro^ towards changes in each subsite informs on structural adaptations associated with the binding of individual residues. The results not only highlight the importance of the P1-Gln residue in substrate recognition by M^pro^, but also those of the P4, P2, and P1′ positions.

**Figure 1:**
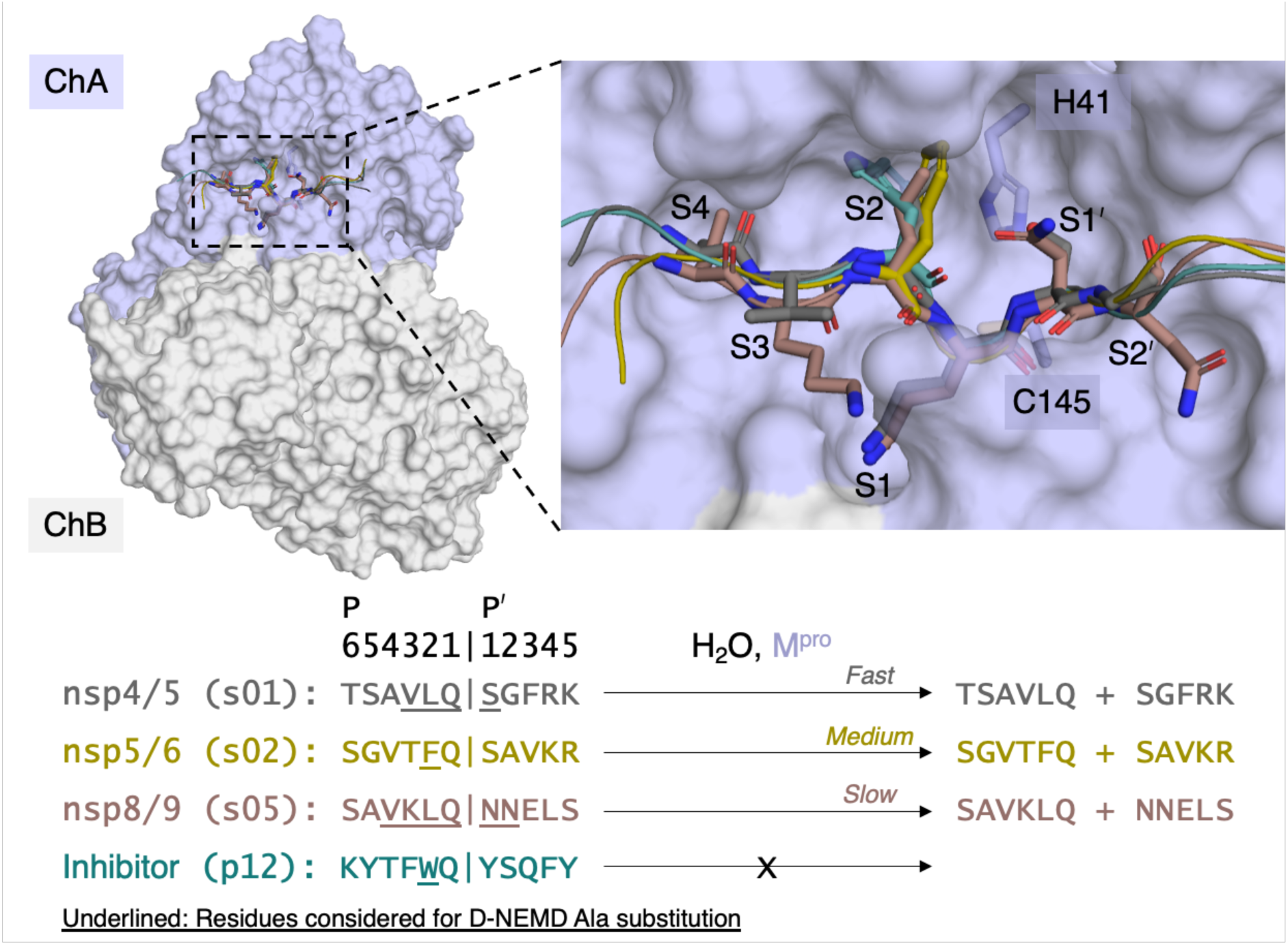
Overview of the M^pro^-peptide complexes in this study. A view of dimeric M^pro^ complexed with the peptide substrates (s01, s02, s05) and inhibitor (p12) prior to MD simulations is presented, with the P4-P2′ residues of s01 and s05, and the P2 residues of s02 and p12, shown as sticks.

## Results and Discussion

### (1) Equilibrium MD simulations of M^pro^-peptide complexes

For MD simulations, we employed models of dimeric M^pro^ noncovalently complexed with the three selected substrate peptides (s01, s02, s05) and inhibitor peptide (p12) employed for MD simulations were prepared following the protocol described previously (**Figure 1**).^21^ The neutral state of the His41-Cys145 pair and the protonation states of histidine residues near the active site were modelled as previously validated (**Supplementary Information Section SM**).^21^ In each case, the peptide was present at the substrate binding site formed predominantly by residues of the M^pro^ chain denoted as chain A (ChA). The substrate peptide adopted an extended conformation, with the P1-P1′ scissile amide bond positioned close to the ChA His41-Cys145 catalytic dyad, and the conserved P1-Gln sidechain extending through the S1 pocket away from His41 (**Figure 2a,b**). The substrate binding site in chain B (ChB) was left unoccupied; unless otherwise specified, we refer to ChA residues in subsequent descriptions. Each M^pro^-peptide complex was subjected to five equilibrium MD simulations of 200 ns, totalling 1 µs per complex. Note that for the M^pro^-s05 complex, this D-NEMD study utilised the same equilibrium simulations as previously reported, the analysis of which is not repeated here.^46^

**Figure 2:**
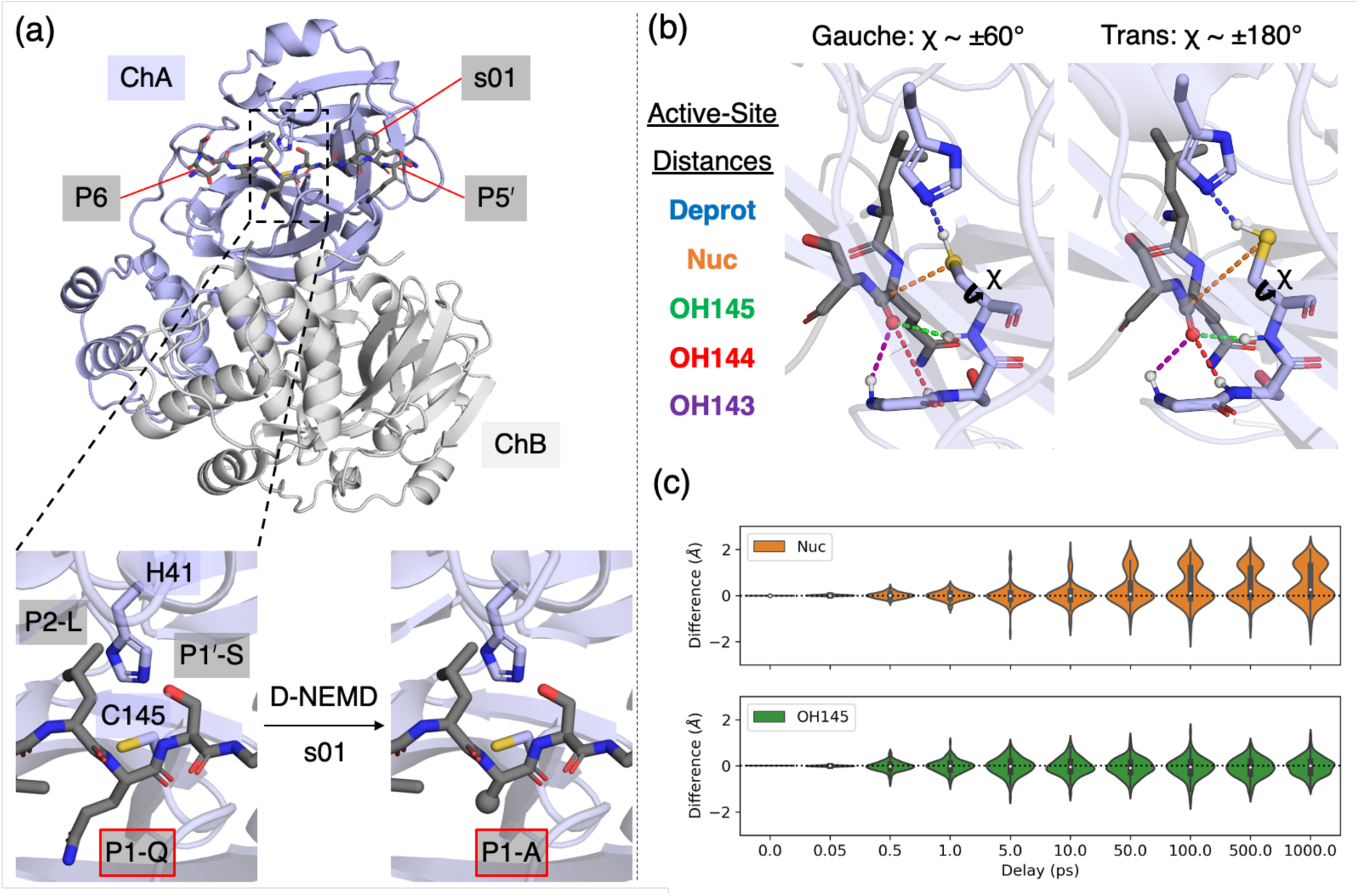
P1 Gln-to-Ala substitution. (a) View of the dimeric M^pro^-s01 complex prior to MD simulations, showing the P1 Gln-to-Ala substitution studied by D-NEMD simulations. (b) The five active-site distances are illustrated using two equilibrium MD-derived snapshots, corresponding to the *gauche* and *trans* Cys145 conformations, which show distinct χ dihedrals. (c) Distributions of the differences in *d*^Nuc^ and *d*^OH145^ following the P1 substitution in s01.

Our analyses confirm the stable binding of the s01, s02, and p12 peptides in the simulations, with all systems observed to be equilibrated from *t* = 50 ns onwards, based on analyses of root mean square deviation (RMSD) and fluctuation (RMSF), protein secondary structure content, and M^pro^-peptide interactions, including 12 conserved hydrogen bonds (HBs 1-12) across the P4-P4′ positions^21^ (**Supplementary Information Section S1**). Similar M^pro^-peptide interactions were observed for both the substrate (s01, s02) and inhibitor (p12) peptides. To investigate if the complexes represent pre-reaction complexes, we assessed binding of the scissile P1-P1′ amide in the active site by monitoring five relevant distances (**Figures 2b and S1.8**). These include distances relevant to deprotonation of Cys145 by His41 *d*(His41_Nɛ2 – Cys145_Hγ) (“*d*^Deprot^”), nucleophilic Cys145 to the scissile amide carbonyl carbon *d*(Cys145_Sγ – P1_C) (“*d*^Nuc^”), and binding of the carbonyl oxygen in the oxyanion hole by the backbone NHs of Cys145, Ser144, and Gly143 *d*(P1_O – Cys145/Ser144/Gly143_H) (“*d*^OH145^/*d*^OH144^/*d*^OH143^”, respectively). In all cases, His41 was apparently poised to deprotonate Cys145 and the scissile amide carbonyl was bound stably in the oxyanion hole. *d*^Nuc^ showed a bimodal distribution, consistent with previous studies,^10, 54^ which resulted from the different *gauche* and *trans* conformations of the Cys145 sidechain (**Figures 2b and S1.8-11**). Across all the modelled M^pro^-peptide complexes, we observed that the N-terminal (P side) of the peptides formed more, and more stable, interactions with M^pro^ than the P′ side, in accord with previous studies (**Figure S1.17**).^21^ Throughout all simulations, the other nitrogen atom (Nδ1) of the His41 sidechain consistently formed a HB to a water molecule, establishing a HB network that connects the sidechains of His41, His164, and Asp187 (**Figures S1.18-19**); the presence of this bridging water molecule (BW) is a consistent feature in crystallographic observations and is probably important in stabilising the positioning and positive charge of His41 during the M^pro^ catalytic cycle.^21, 55-57^

Equilibrium MD simulations were also performed for the s01 and s05 variants with the P1-Gln replaced by Ala, named s01-QP1A and s05-QP1A. s01-QP1A and s05-QP1A remained stably bound throughout the MD simulations, and maintained most of the conserved HBs with the protein, except for HBs 6-7, which require a P1-Gln sidechain (**Figure S1.12**). In both cases, the P1-P1′ amide was stably bound in the oxyanion hole (**Figure S1.8**). However, analysis of *d*^Nuc^ distances indicated the predominance of the non-productive *trans* Cys145 conformation relative to the productive *gauche* conformation, when compared to the corresponding wild type (wt) peptides (**Figures S1.8-11**).

This observation raised the questions of whether and how residue identity at each substrate position affects the M^pro^ active site configuration. These questions were addressed using D-NEMD, by “substituting” each residue of interest individually with Ala. Initial configurations of M^pro^ complexed with wt peptides were taken from the five equilibrium MD simulations from *t* = 50 ns up to 195 ns in 5 ns intervals, yielding 150 pairs of short equilibrium and nonequilibrium propagations in each case (*N* = 150). The subtraction technique in vector evaluation helps to distinguish significant coherent displacements caused by the perturbation while reducing random fluctuations of mobile atoms.^39, 41, 46^

Given the lack of conservation, ordered structure, and well-defined M^pro^ subsites outside of P4-P2′ of substrate peptides,^12, 14, 18, 20, 21^ substitution was only considered in the peptide P4-P2′ positions, excluding cases where the original residue was Ala or Gly (*e.g.*, P4 and P2′ of s01) (**Figure 1**). D-NEMD responses were measured up to 1 ns as in previous studies^46^ and were quantified by the following approaches. First, to characterise any statistically significant changes in the conformation of the protein and peptide binding in the active site, differences in the five active-site distances between nonequilibrium (*d*^neq^) and equilibrium (*d*^eqm^) propagations at equivalent timepoints, Δ*d* = *d*^neq^ – *d*^eqm^, were evaluated. Second, to consider the overall effect on M^pro^-peptide binding, similarly, the differences in the donor-acceptor distances (Δ*d*^D-A^) of the conserved HBs 1-5 and 8-12, all of which involve the peptide backbone, were calculated. HBs 6-7, which are dependent on the P1-Gln sidechain, were excluded. Third, the displacement vector, **v** = **r**^neq^ – **r**^eqm^ where **r** is atomic position, of each of the 612 M^pro^ and 11 peptide Cα atoms, was measured and averaged across the 150 D-NEMD replicas, with Cα atoms chosen to represent the most substantial structural changes in the protein.^38, 46^ In regions of interest according to the above analyses, average displacement vectors were evaluated additionally for non-hydrogen atoms.

### (2) Effect of the P1 Gln-to-Ala substitution

The role of the P1-Gln, a residue conserved across all natural M^pro^ substrates,^5, 12^ was investigated by substituting P1-Gln with Ala in s01 and s05, with the response analysed using D-NEMD simulations (**Figure 2a**; **Section S2**). Analysis of the changes in HBs between the P1 scissile amide carbonyl and the oxyanion hole (Δ*d*^OH145^/Δ*d*^OH144^/Δ*d*^OH143^, **Figures 2b,c and S2.1**; **Table S2.1**) showed narrow distributions, indicating these interactions remained stable upon substitution. The distance relevant to deprotonation of Cys145 by His41 (“*d*^Deprot^”) showed a wider spread, likely due to the weaker HB involving a thiol SH than those with OH or NH, but was overall centred around zero. The largest differences were observed in the Cys145_Sγ – P1_C (*d*^Nuc^) distance, with predominantly positive differences for both s01 and s05, especially after 10 ps post perturbation. For example, the average Δ*d*^Nuc^ at *t* = 100 ps was (0.37 ± 0.06) Å for s01 and (0.19 ± 0.05) Å for s05 (error bar = SEM), indicating a statistically significant increase. This increase in distance was primarily caused by movement of Cys145 Sγ away from the scissile amide carbonyl, which was itself not significantly affected according to the unaffected *d*^OH^ distances; the Cys145 sidechain switched from the productive *gauche* to the non-productive *trans* conformation.

Analysis of HBs 1-3 and 8-12 (**Tables S2.2-3**; **Figure S2.2**; note HBs 6-7 require a Gln sidechain and hence do not exist here) showed little change with Δ*d*^D-A^ values around zero. However, a wider distribution was observed for HB4, probably due to its lower stability and involvement of the flexible Gln189 sidechain.^21^ Notably, there was a consistent and statistically significant decrease in the HB5 *d*^D-A^ distance between the peptide P1 backbone N and His164 backbone O (**Table S2.2**), with Δ*d*^D-A^(HB5) = (-0.23 ± 0.03) Å for s01 and (-0.16 ± 0.02) Å for s05 at *t* = 100 ps. Surprisingly, this observation suggests tighter binding of the P1 backbone following removal of the P1-Gln sidechain.

To better understand the effects of the P1 Gln-to-Ala substitution, we visualized the response of the M^pro^ protein by analysing the non-hydrogen atomic displacement vectors averaged across 150 D-NEMD replicas, with a focus on the M^pro^ S1 pocket (**Figure 3**). Initially (at *t* = 0.5 ps) following the substitution, the largest displacements were observed in the M^pro^ His163 sidechain, which moved outwards into the space originally occupied by the P1-Gln sidechain, and in the Phe140 backbone, which moved likely due to the loss of HB7. At *t* = 5 ps, the signal propagated outwards from the S1 pocket, with displacements observed in neighbouring residues such as the Asn142 sidechain. Interestingly, the P1-Ala backbone and Cβ atoms moved inwards (**Figure S2.7**), resulting in the observed decrease in *d*^D-A^(HB5). At *t* = 50 ps, the Cys145 Sγ atom manifested displacement away from the P1 scissile amide carbonyl, accompanied by movement of the Glu166 sidechain closer into S1 and shift of the Leu141 sidechain. Residues in the Ser46-Leu50 helix moved away from the active site. The movement of the Cys145 Sγ away from the P1 peptide carbonyl, caused by a change in the Cys145 sidechain conformation from *gauche* to *trans*, continued to later timepoints, including *t* = 500 ps, at which point His41 also moved away from the active site. Similar effects on Cys145 were observed in the case of s05 from *t* = 50 ps onwards (**Figure S2.6**).

**Figure 3:**
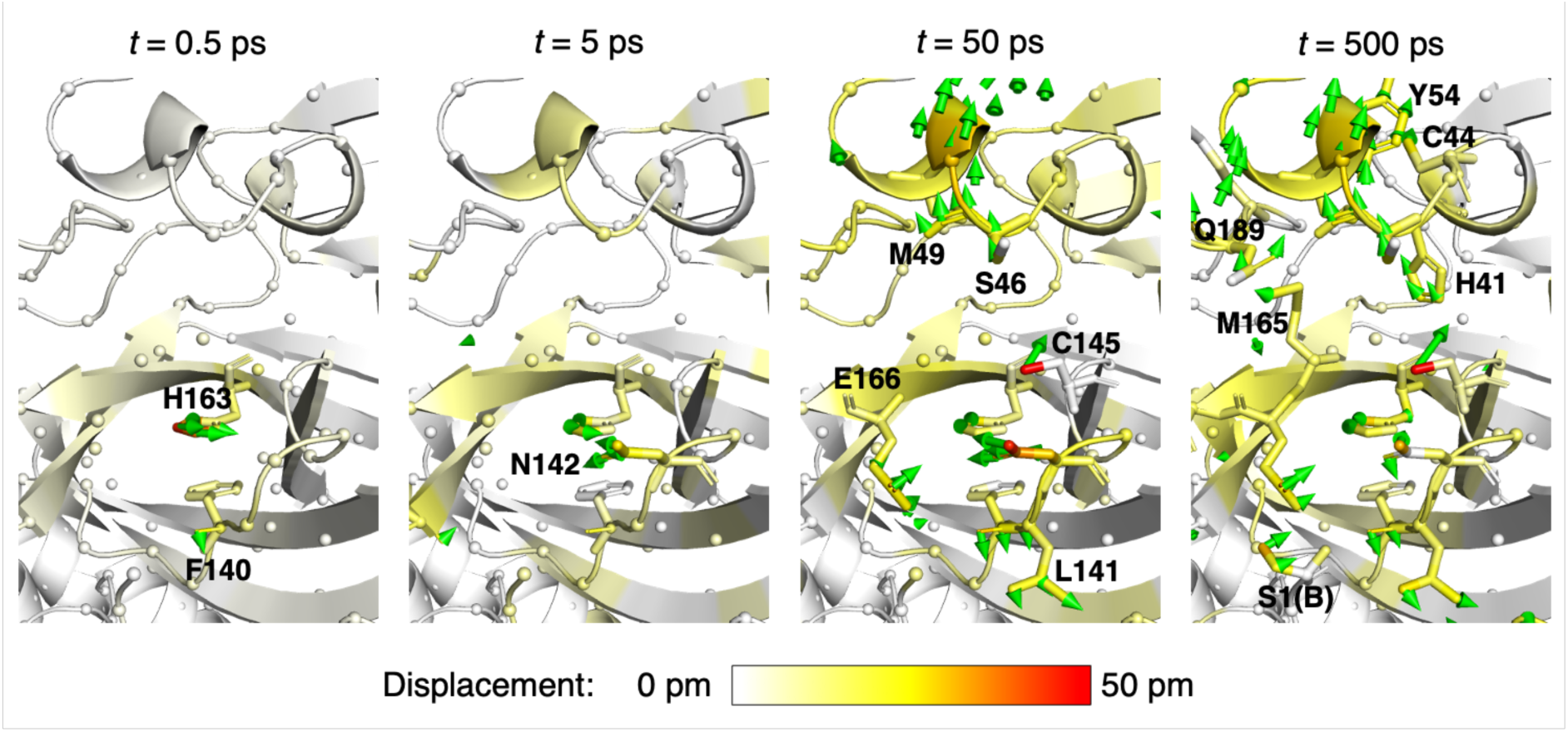
The response of the M^pro^ S1 subsite residues to P1 Gln-to-Ala substitution in s01. The response was measured by averaging the non-hydrogen displacement vectors between the equilibrium and nonequilibrium propagations at selected timepoints following the substitution. Displacement magnitudes are displayed on a white-yellow-red scale, using the structure prior to MD. Significant vectors with length ≥20 pm are shown as green arrows with a scale-up factor of 5.^58, 59^ Residues that show such significant displacements located within 10 Å of the substrate P1 residue are visualised as sticks.

In summary, the results demonstrate that recognition of the conserved P1-Gln is communicated to M^pro^ residues beyond those directly binding to its sidechain. This substitution induces structural adaptations in residues that constitute the S1 pocket and the helices above the active site (**Figure S2.6**). Notably, the substitution causes a change in Cys145 from the productive *gauche* to the non-productive *trans* conformation, along with a potential movement of the His41 sidechain away from Cys145 Sγ, thus hindering initiation of covalent reaction. Conversely, the binding of the P1-Gln sidechain, and its associated interactions with His163 and Phe140 in the S1 pocket, helps position the P1 backbone in the active site, and activates the catalytic dyad residues His41 and Cys145 into their productive conformations. This recognition and communication process ensures that M^pro^ selectively targets the backbone carbonyl of a Gln residue.

### (3) Effect of P2 substitution

To investigate if there is coordinated dynamic behaviour of M^pro^ regions upon binding of the P2 residue, we performed D-NEMD simulations on s01 and s05 by substituting P2-Leu with Ala (**Section S3**). P2 is a Leu in nine of the eleven native substrates, with the exceptions being s02 and the nsp6/7 substrate (“s03”), which have Phe and Val at P2, respectively.

Our results revealed that substitution of P2-Leu with Ala had minimal impact on the active-site distances and the HB *d*^D-A^ distances (**Tables S3.1-3**; **Figures S3.1-2**), implying no significant impact on the Cys145 sidechain conformation. Displacement vector analysis showed that M^pro^ remained mostly unperturbed relative to equilibrium following the substitution at the P2 site (**Figures 4 and S3.5-6**). The immediate response of M^pro^ was mostly localised to S2 residues, such as the Met49 and Met165 sidechains. These changes were observed after tens of picoseconds following the perturbation. These findings suggest that only a small structural rearrangement of M^pro^ residues is required to accommodate the P2-Leu.

**Figure 4:**
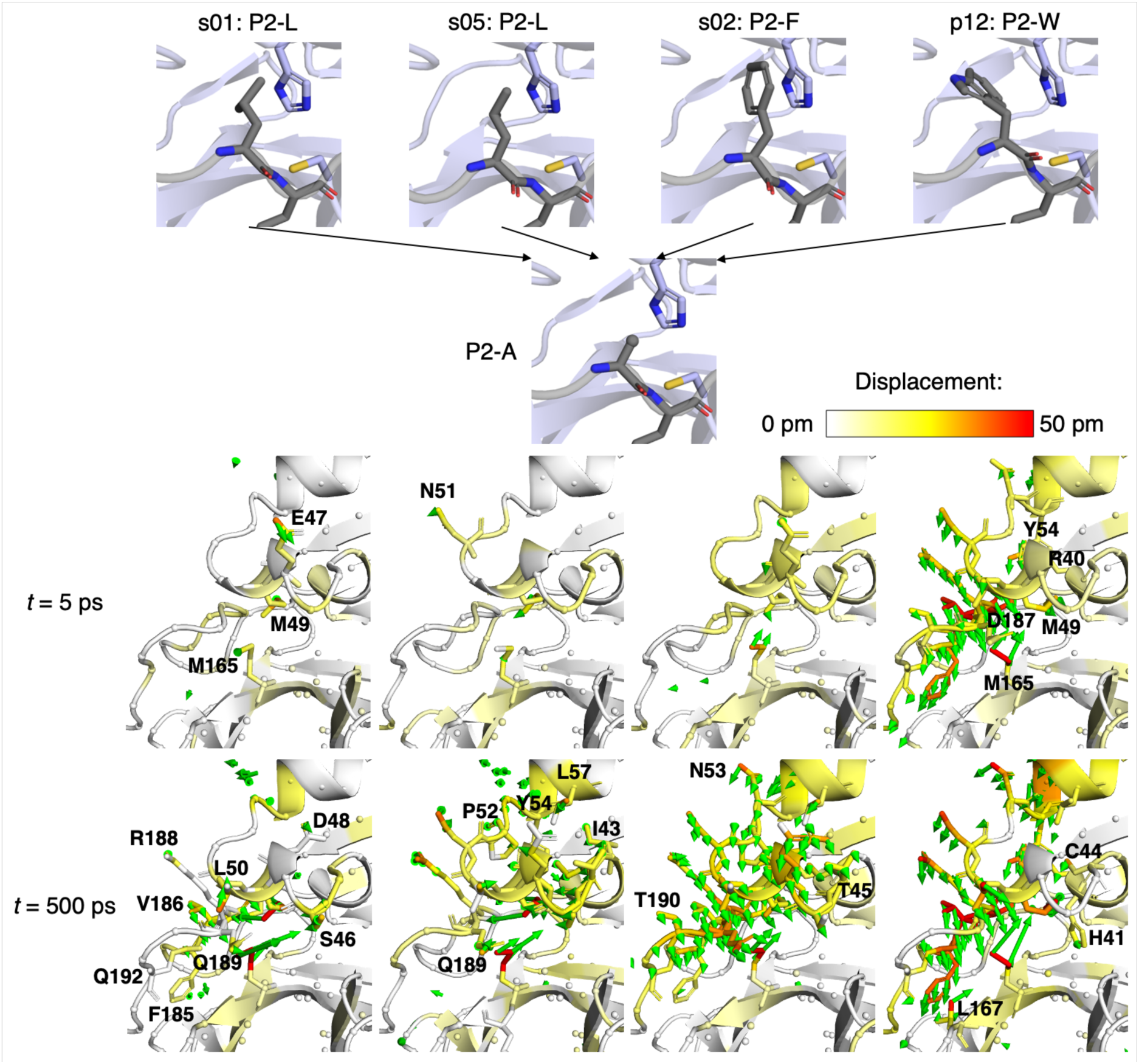
M^pro^ response to P2 substitution in substrate and inhibitor peptides. The structural response was measured by averaging the non-hydrogen displacement vectors between the equilibrium and nonequilibrium propagations at selected timepoints following the substitution. Displacement magnitudes are displayed on a white-yellow-red scale, using the structure prior to MD. Significant vectors with length ≥20 pm are shown as green arrows with a scale-up factor of 5.^58, 59^ Residues that show such significant displacements located within 10 Å of the peptide P2 residue are visualised as sticks.

We have previously reported the competitive inhibition of M^pro^ by peptides that resist hydrolysis, which we attributed to binding of a P2-Trp in the S2 pocket.^21^ To investigate if there are differences in M^pro^ dynamics when binding an aromatic ring in S2, we substituted the P2-Phe residue in s02 and the P2-Trp residue in p12 with Ala, and analysed the response by D-NEMD simulations. While these substitutions did not substantially affect the active-site distances, the flexible HB4 *d*^D-A^ decreased in both systems, with average Δ*d*^D-A^ = (-0.37 ± 0.12) Å and (-0.23 ± 0.11) Å at *t* = 100 ps for s02 and p12, respectively. For s02, *d*^D-A^(HB1) was also reduced. As the *d*^D-A^ of HBs 2-3 that bind the peptide P3 backbone remained unaffected, the decreases in *d*^D-A^ of HB4, and in the s02 case also HB1, suggested an impact of the P2-Phe/Trp on the conformations of Thr190 and/or Gln189, as visualised by displacement vectors (**Figure 4**).

Compared to s01 and s05, where responses were mostly localised on Met49 and Met165, the s02 and p12 substitutions resulted in substantial motions. These included a collective movement towards the substrate binding pocket of the residues in the Ser46-Leu50 helix adjacent to the substrate binding pocket, in the Asn53-Arg60 helix, which is located further away from the substrate binding pocket, and in the connecting loop in between (Asn51-Pro52). There were also outward movements of the Phe185-Gln192 loop, whose residues line the S4 and S2 pockets. Notably, significant displacements were observed in both the backbone and sidechain atoms of Asp187. Given the BW-mediated link between the Asp187 carboxylate, His164, and His41 of the catalytic dyad (**Figures S1.18-19**), displacement in Asp187 may have a cascading effect on His41 and influence the catalytic activity of M^pro^. Indeed, in the hundreds of picoseconds timescale (*e.g.*, *t* = 500 ps) for p12, we observed small but statistically significant displacements in the His41 sidechain, which moved towards the P1′-Tyr of p12. This observation demonstrates the operation of communication between the P2-Trp and His41, as previously proposed,^21^ possibly mediated by BW (**Figures S1.18-19**).

In conclusion, the rearrangement involved in M^pro^ residues in the Phe185-Gln192 loop, including Asp187, to accommodate the larger P2-Phe/Trp in s02/p12, may explain the poor or lack of hydrolytic activity towards these peptides.^21^

### (4) Substitution of P3 and P4 residues

We then investigated the effects of substituting P3 and P4 residues on M^pro^ dynamics. The P3 position has been assumed to confer little selectivity, with a range of residues including Val, Thr, Lys, Arg, and Met found across the eleven native polyprotein derived substrates.^5, 12^ While the P3 backbone NH and O atoms in all substrates are positioned to hydrogen bond with the Glu166 backbone (HBs 2-3), the P3 sidechain is solvent exposed and demonstrates flexibility relative to the neighbouring P4 and P2 residues.^21^

Upon substitution of the respective P3-Val and P3-Lys in s01 and s05 with Ala and analysis of the system response using the D-NEMD approach (**Section S4A**), no significant deviations in the active-site distances were observed, implying no significant impact on Cys145. The most notable change was observed in HB4, resulting from the P3 Lys-to-Ala substitution in s05 (**Table S4A.2**). In this case, the *d*^D-A^ distance increased on average from *t* = 50 ps onwards, with Δ*d* = (0.74 ± 0.13) Å at *t* = 500 ps. During the equilibrium MD simulations, the P3-Lys sidechain occasionally adopted conformations which folded back towards the peptide P2 backbone NH, with the simultaneous coordination of Gln189 sidechain carbonyl oxygen to the P2 backbone NH (HB4) and the P3-Lys amino group. This coordination and shielding from solvent by the P3-Lys sidechain likely contributes to positioning of Gln189,^18^ but these effects were lost upon the substitution of the P3-Lys sidechain with Ala. In terms of displacement, the effect of substituting P3 was minimal and less consistent over the observed timescales, compared to substitutions of P1 or P2, which was expected given the lack of a well-defined pocket on M^pro^ that encloses the P3 sidechain and the associated lack of substrate selectivity (**Figures S4A.5-6**).

To investigate substitution of the P4 residue, we focused on s05 since P4 was already Ala in s01. Substituting P4-Val with Ala in s05 did not significantly affect the active-site distances (**Section S4B**), implying little or no impact on Cys145 conformation. However, there was an average decrease in *d*^D-A^ of HBs 1 and 4 (**Table S4B.2**). Further analysis of the displacement vectors revealed that in the picosecond timescale following the substitution (*e.g.*, *t* = 5 ps), the M^pro^ loop residues Val186-Ala193 responded by closing around the S4 pocket, with movements observed in both the backbone and sidechain atoms (**Figures 5 and S4B.5-6**). The signal propagated downstream along the loop to residues including Thr196 at *t* = 500 ps, as the loop adapted to the substitution of the P4-Val sidechain. This dynamic response is consistent with previous crystallographic observations that the Val186-Gly195 loop is structurally variable depending on the bound substrate sequence,^20^ and echoes the coherent D-NEMD response observed when the full s05 substrate peptide is removed.^46^ In comparison to the M^pro^ response when the P2-Leu was substituted with Ala in the same s05 substrate, the S4 pocket needed to undergo more significant structural adaptation to accommodate the substrate P4-Val. This was in contrast to the relatively rigid S2 pocket that accommodates the predominantly conserved P2-Leu residue.

**Figure 5:**
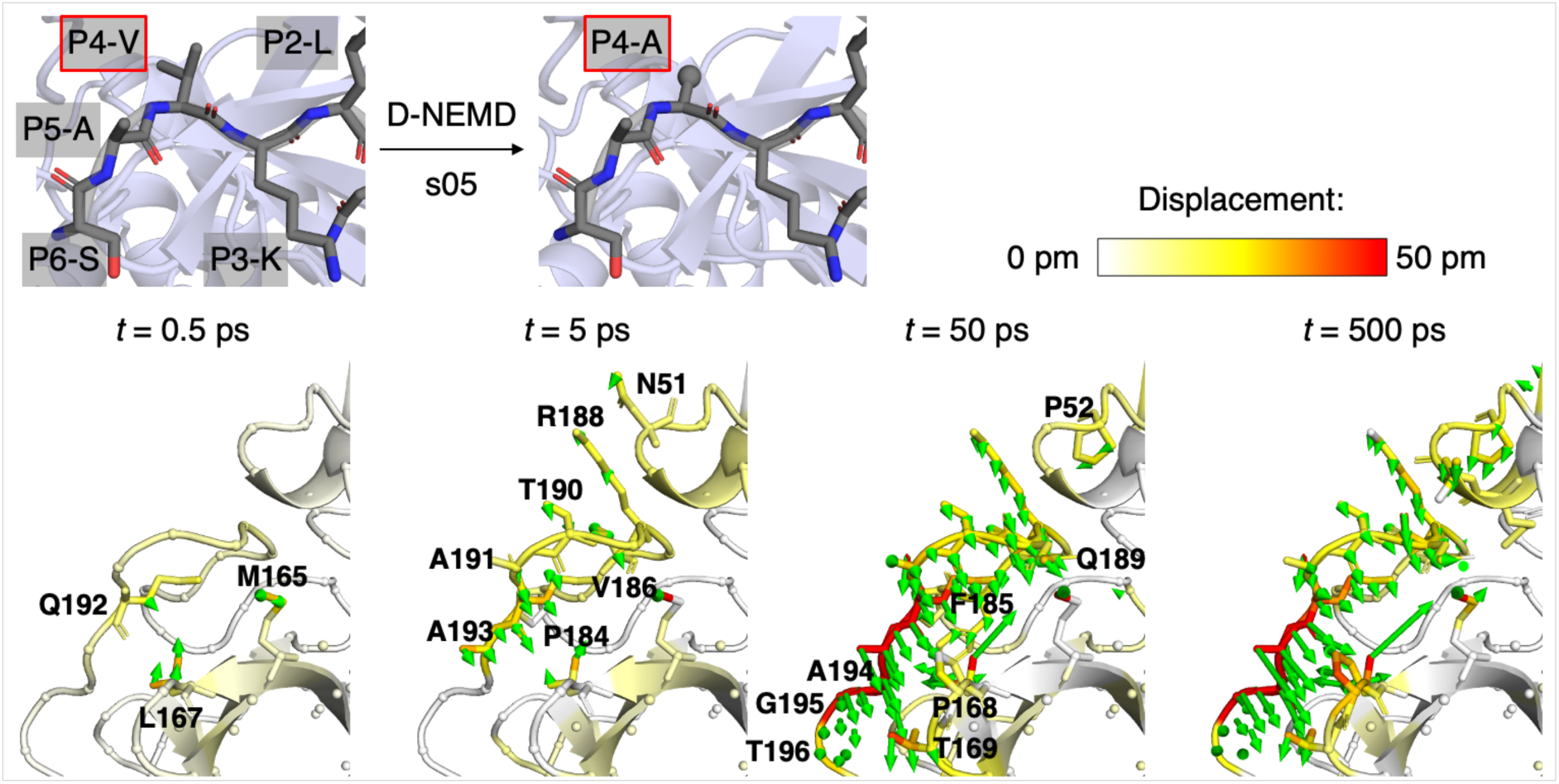
M^pro^ response to P4 substitution in s05. The response was measured by averaging the non-hydrogen displacement vectors between the equilibrium and nonequilibrium propagations at selected timepoints following substitution. Displacement magnitudes are displayed on a white-yellow-red scale, using the structure prior to MD. Significant vectors with length ≥20 pm are shown as green arrows with a scale-up factor of 5.^58, 59^ Residues that show such significant displacements located within 10 Å of the substrate P4 residue are visualised as sticks.

### (5) Substitution of P1′ and P2′ residues

The available biophysical and kinetic evidence is that the S subsites on M^pro^, responsible for binding the substrate P residues, are more well-defined and form more HB interactions with the substrate than the S′ subsites, which bind residues C-terminal of the scissile amide (**Figures S1.12, S1.17**).^21^ On the P′ side, substrate selectivity has only been reported at the P1′ position, which requires a small, neutral residue, with Ser, Ala, and Asn observed among SARS-CoV-2 M^pro^ native substrates,^18^ and with Gly present at the P1′ position in the nsp5/6 sequence of SARS-CoV.^60^ Here, we analysed the response of M^pro^ to Ala substitution at the P1′-Ser/Asn site in s01/s05, respectively, using D-NEMD simulations (**Section S5A**). Among the active-site distances analysed, we observed an average decrease in the *d*^Deprot^ distance, with s05 exhibiting a consistently negative Δ*d*^Deprot^ from *t* = 0.25 ps onwards (**Table S5A.1**). Analysis of displacement vectors for s05 revealed movement of the His41 sidechain to form a closer association with Cys145 and the scissile amide carbonyl, *e.g.*, at *t* = 5 ps and 500 ps (**Figure 6**). In the equilibrium MD simulations, His41 was observed to engage in a HB with the P1′-Asn sidechain in s05 (25% of the frames) and to a lesser extent with the P1′-Ser sidechain in s01 (18%). These HBs may compete with the coordination of His41 to Cys145, which is necessary for initiating deprotonation. Hence, these HBs potentially contribute to the experimentally observed slow hydrolysis of s05, compared to other substrates with P1′-Ser (*e.g.*, s01 and s02) or P1′-Ala (*e.g.*, the nsp9/10-derived substrate “s06”, ATVRLQ|AGNAT, which also has P4-Val and P2-Leu like s05),^21^ despite the optimal P4-P1 sequence in s05.^17^ Indeed, a P1′ Asn-to-Ala substitution in a similar coronavirus nsp8/9-derived substrate is experimentally observed to result in approximately three times more efficient hydrolysis than the wt peptide with P1′-Asn.^18^ However, more experimental validation is required to confirm our hypothesis, since s05 is the only native substrate possessing a P1′-Asn.^21^ Importantly, no significant changes were observed in the other active-site distances, implying the absence of any impact on Cys145 conformation or on P1 carbonyl binding in the oxyanion hole. Among the M^pro^-peptide HBs (**Tables S5A.2-3**), an average distance decrease was observed in HB4 in s01 after *t* = 300 ps, accompanied by a displacement of the Gln189 sidechain towards the P2 backbone (**Figure S5A.6**).

**Figure 6:**
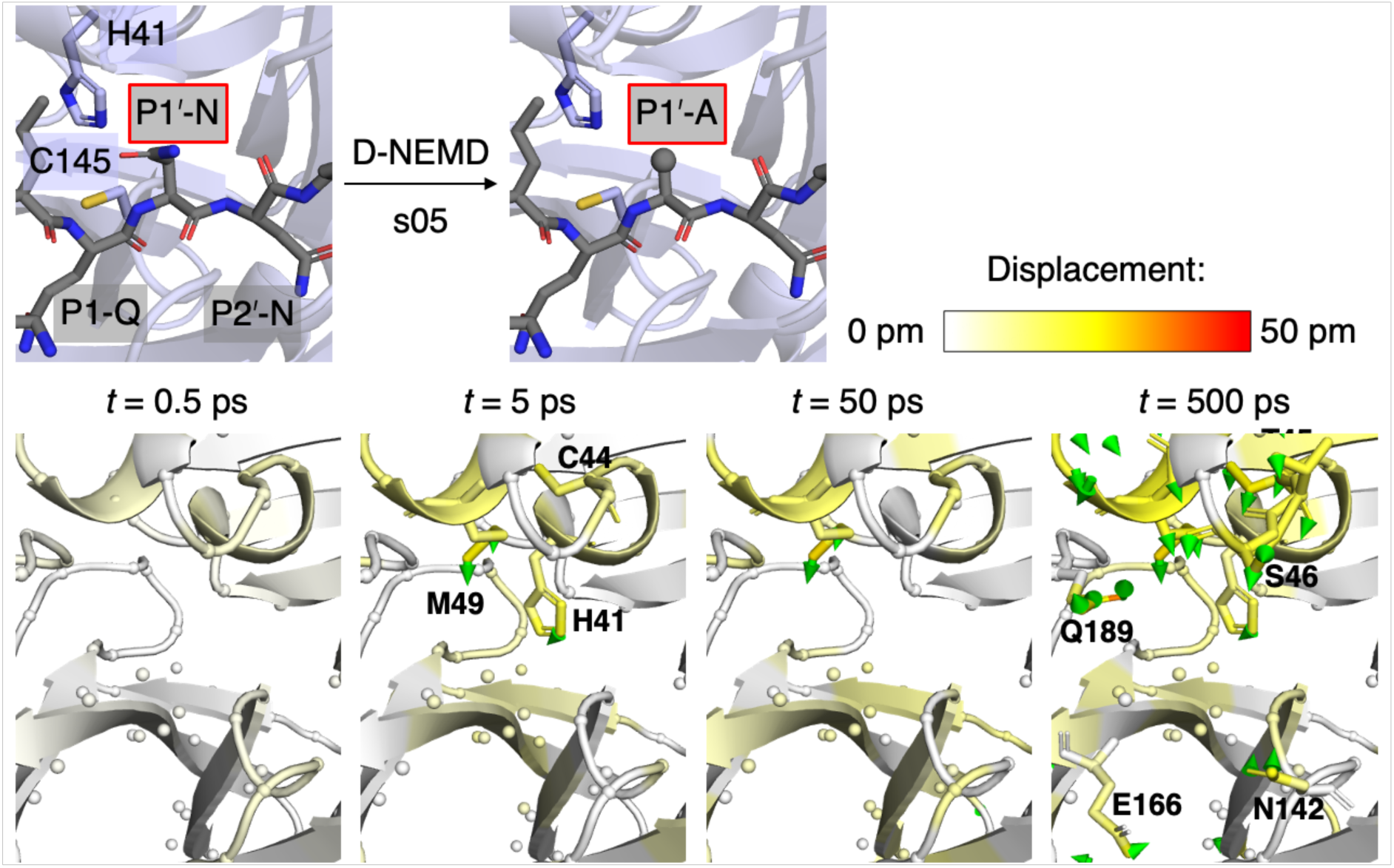
M^pro^ response to P1′ substitution in s05. The response was measured by averaging the non-hydrogen displacement vectors between the equilibrium and nonequilibrium propagations at selected timepoints following the substitution. Displacement magnitudes are displayed on a white-yellow-red scale, using the structure prior to MD. Significant vectors with length ≥20 pm are shown as green arrows with a scale-up factor of 5.^58, 59^ Residues that show such significant displacements located within 10 Å of the substrate P1′ residue are visualised as sticks.

At the P2′ position, Asn in s05 was substituted with Ala (**Section S5B**); no substitution was performed for s01, which has a P2′-Gly. Overall, the substitution did not cause significant deviations in the active-site or HB distances, including HBs 10-11 involving the P2′ backbone. There were no significant displacements in M^pro^ residues. These observations are consistent with the lack of substrate selectivity at P2′, and the solvent exposed and flexible nature of its sidechain.

## Conclusions

The D-NEMD method has emerged as a powerful tool for probing signal propagation and allosteric communication networks in proteins. By combining equilibrium and nonequilibrium simulations, the D-NEMD approach enables the identification of statistically significant changes by averaging responses across hundreds of replicas.

In a recent study, we utilised D-NEMD simulations to identify communication pathways from the SARS-CoV-2 M^pro^ active site to an experimentally identified allosteric site,^61^ and to sites relevant to drug resistant substitutions. This was achieved by entirely annihilating the native substrate s05 in the M^pro^-s05 complex.^46^ The work reported here employed an Ala substitution strategy to investigate the structural response of M^pro^ to binding of substrate residues in each of its active site cleft subsites. We sought to elucidate how the response relates to the overall processes of substrate recognition and selectivity (**Figure 7**).

**Figure 7:**
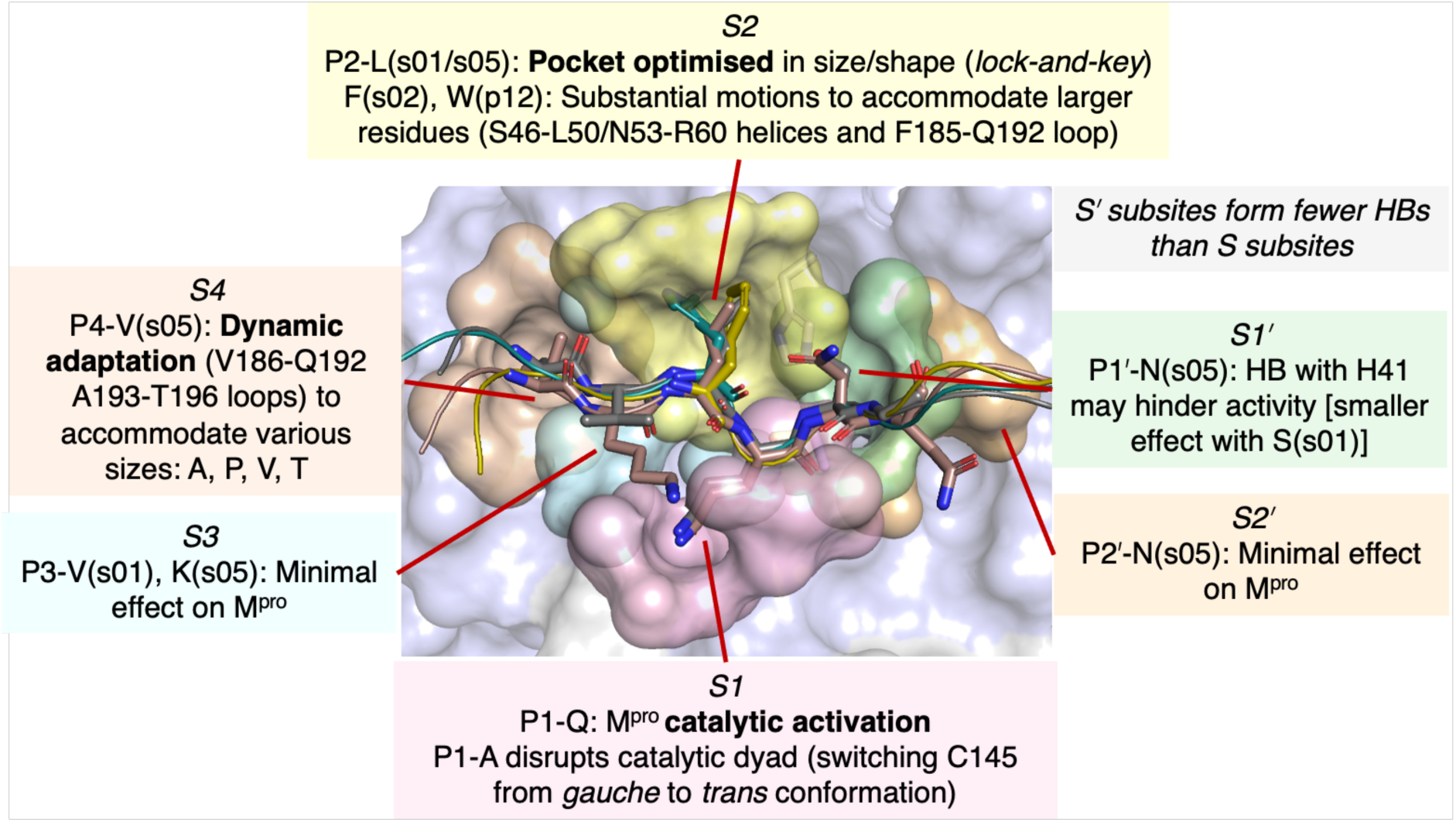
Summary of the D-NEMD derived M^pro^ responses to peptide Ala substitutions. Carbon atoms of the peptides s01, s02, s05, and p12 are coloured grey, olive, brown, and teal, respectively. Each of the S4-S2**′** M^pro^ subsite is represented as a surface defined by 3 Å contact with the corresponding peptide residue.

Our results reveal substantial differences in the nature and magnitude of the responses of M^pro^ residues when sidechain moieties at each subsite were removed through Ala substitution. Substitutions at positions with minimal experimentally observed selectivity requirement, such as the P3 and P2′ positions, had negligible effect on the conformations of M^pro^ residues. However, substitutions at the P1 and P1′ positions induced significant conformational changes, affecting the catalytic dyad conformation and consequently catalysis. At P1, Ala substitution led to significant conformational changes in the catalytic dyad, switching Cys145 from an active *gauche* to an inactive *trans* conformation, thereby inhibiting reaction. Notably, the binding of P1-Gln, a conserved residue among all native substrates of M^pro^, would thus activate the catalytic dyad, ensuring M^pro^ selectively cleaves at the designated positions in the viral polyproteins. Interactions with the P1′ sidechain were found to influence the conformation and activity of His41.

Additionally, we explored the dynamic behaviour of the S4 and S2 subsites in accommodating the corresponding P4 and P2 substrate residues. The S4 pocket is known to accommodate residues of various sizes, including Ala, Pro, Val, and Thr. This is likely facilitated by the dynamic adaptation of the S4 pocket lined by the Val186-Gln192 loop and possibly collective movements of downstream residues (Ala193-Thr196). The dynamic perturbation of this region upon substrate removal and its relevance to drug-resistant M^pro^ substitutions were highlighted in our previous D-NEMD study.^46^ Thus, accommodating the P4 residue requires M^pro^ to undergo induced fit. By contrast, the S2 pocket is minimally affected by the truncation of P2-Leu to alanine, perhaps reflecting a binding interaction resembling a *lock-and-key* model, where the pocket is optimised in size and shape for accommodating the P2-Leu, which is present in nine of the eleven polyprotein cleavage site sequences. Interestingly, the rigid, Leu-mimicking, 6,6-dimethyl-3-azabicyclo[3.1.0]hexane group in nirmatrelvir was observed to fully occupy this pocket (**Figure S1.20**).^7, 62^ However, accommodation of alternative residues, such as P2-Phe of s02 and P2-Trp of the inhibitor peptide p12, which is not efficiently hydrolysed by M^pro^, led to more significant adaptations in M^pro^ residues, including those related to the catalytic mechanism, potentially hindering peptide hydrolysis.

The plotting of statistically significant displacement vectors of each Cα or non-hydrogen atom in response to specific peptide substitutions (**Figures S2.3-4, S3.3-4, S4A.3-4, S4B.3-4, S5A.3-4, S5B.3-4**) provides a comprehensive analysis of protein conformational changes. We propose that each of these plots acts as a dynamic fingerprint representing the response to a specific perturbation, with peaks corresponding to residues or atoms that display the largest structural adaptations. Such dynamic fingerprint mapping could be a valuable tool for characterising cases involving modifications to bound ligands, in a similar manner as free energy perturbation methods inform on binding free energy differences. Such analysis could be utilised to characterise whether and how a ligand modification is tolerated and/or affects the structure and dynamics of the target receptor. This is particularly relevant in cases where drug resistance might arise due to distal substitutions altering protein dynamic behaviour, as observed in certain drug targets, such as the HIV-1 protease.^63^ Therefore, the dynamic mapping offered by the D-NEMD approach has the potential to provide useful insights into residues susceptible to drug resistant substitutions due to changes in receptor dynamics.

Future possible application of the D-NEMD approach could provide a dynamic view of how M^pro^ responds when substrates are converted into products, similar to D-NEMD studies on protein responses upon ATP or GTP hydrolysis.^64-66^ This is particularly intriguing given that product fragments have identical residue sidechains to the original substrate and the observation of stable M^pro^-product complexes both experimentally, using C145A M^pro^,^11^ and computationally.^9, 10^ An interesting question is how the release of products from M^pro^, and more generally from any protease, occurs. Equilibrium MD-based free energy calculations have suggested weakening of P1-Gln binding upon product formation in M^pro^,^67^ but further analysis would be required to evaluate how this process is facilitated.

Overall, the D-NEMD results inform on how M^pro^ achieves substrate selectivity and hydrolytic efficiencies towards different substrates (**Figure 7**). The findings and approach presented here may help in optimising M^pro^ inhibitors, especially in terms of optimising binding and, where appropriate, covalent reaction with the nucleophilic cysteine. The observed differences in induced fit among substrate residue-enzyme subsite interactions are particularly intriguing. For example, the S2-P2 interaction appears relatively unaffected, as supported by the binding mode of nirmatrelvir compared to that of the P2-Leu residue predominantly observed among substrates (**Figure S1.20**).^62^ By contrast, the S4-P4 and S1-P1 interactions involve substantial induced fit, with the latter linking to the promotion of a catalytically productive state of the His41-Cys145 dyad. D-NEMD simulations may help guide the design of functional group, with consideration of their flexibility for optimal binding in a particular subsite, in future combinatorial computational-experimental drug discovery campaigns.

## Methods

### Equilibrium molecular dynamics (MD) setup

The SARS-CoV-2 M^pro^ dimer (protein structure taken from PDB code 6YB7;^68^ 1.25 Å resolution) noncovalently complexed with a polyprotein-derived^19^ native substrate peptide (s01, s02, s05) or an inhibitor peptide (p12)^21^ in the chain A (ChA) active site was prepared as described previously.^21, 46^ The substituted s01-QP1A and s05-QP1A peptides were modelled by truncating the P1-Gln sidechains to Ala in s01 and s05 respectively. Cys145 and His41 were both neutral, and the setup of titratable and histidine residues (**Table SM.1**) was identical to previous setups.^21, 46^ Using the GROMACS (v 2019.2/2020.3/2020.4)^69^ MD package and the AMBERFF99SB-ILDN force field,^70^ and following previously reported procedures,^21, 46^ each M^pro^-peptide complex was solvated with TIP3P water^71^ in a rhombic dodecahedral box with edges at least 1 nm away from the complex and neutralised by sodium ions, with the total number of atoms being 86,305, 86,326, 86,304, 86,241, 86,301, and 86,297 for the s01, s02, s05, p12, s01-QP1A, and s05-QP1A complexes respectively. The system was subjected to energy minimisation until convergence with maximum force <1000 kJ mol^-1^ nm^-1^. Starting from random velocities from the Maxwell-Boltzmann distribution at 298.15 K, five replicas were initiated by a 200 ps (1 fs step) NVT and 200 ps (1 fs step) NPT equilibration, followed by a 200 ns (2 fs step) production MD run. A temperature of 298.15 K was maintained by velocity rescaling with a stochastic term, with separate coupling to the protein and non-protein (time constant = 0.1 ps).^72^ A pressure of 1.0 bar was maintained by a Parrinello-Rahman barostat (time constant = 2 ps).^73, 74^ Van der Waals interactions were cut off at 1 nm. Long-range electrostatic interactions beyond 1 nm were calculated using smooth particle mesh Ewald.^75, 76^ Frames were saved every 100 ps and velocities were saved every 1 ns. Trajectories were analysed using GROMACS (v 2019.2)^69^ and DSSP (v 2.0.4).^77, 78^ Hydrogen bonds were defined according to the default criteria in gmx hbond: donor-acceptor distance *d*^D-A^ ≤ 3.5 Å and hydrogen-donor-acceptor angle ∠^H-D-A^ ≤ 30°.

### Dynamical nonequilibrium MD (D-NEMD) setup

D-NEMD simulations were set up and analysed similarly to previously reported procedures.^46^ For every M^pro^-peptide complex system, frames were extracted from the five equilibrium MD simulations in 5 ns intervals from *t* = 50 ns to 195 ns, resulting in 5 × 30 = 150 configurations. To initiate the nonequilibrium simulations, in these configurations, the residue of interest was instantaneously substituted with Ala by truncation up to the Cβ atom and its bonded protons (Hβ1-3), with the Cγ atom modified to Hβ3 at the same position, and the simulations were run using the Ala topology. No new atoms were added, except in the s05 P3 Lys-to-Ala substitution, for which a water molecule was replaced by a sodium ion to preserve overall system charge neutrality. The corresponding equilibrium simulations were run using the original unmodified topology. To obtain the responses over different timescales, the pairs of equilibrium and nonequilibrium simulations were analysed over two lengths: (i) over 10 ps saving frames every 50 fs; (ii) over 1 ns saving frames every 10 ps. For the s05 P1 Gln-to-Ala substitution, the lengths and analysis frequencies of the equilibrium-nonequilibrium simulation pairs were as previously reported.^46^

### D-NEMD analysis

The M^pro^-peptide complex structure at a certain post-substitution timepoint was extracted from the nonequilibrium trajectory and superimposed onto the structure from the equilibrium trajectory at the equivalent timepoint. The two structures were compared by evaluating the equilibrium-to-nonequilibrium displacement vector (*x*, *y*, *z*) of every non-hydrogen atom, and the displacement vectors were averaged across the 150 replicas to yield the average vector, which, according to the Kubo-Onsager relation,^37, 38^ estimates the macroscopic time-dependent response of the M^pro^-peptide complex to the substitution. The magnitude of the average vector *v* was evaluated by:

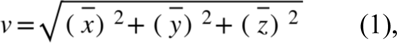

where 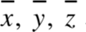 are the displacements in the x, y, and z directions, respectively, of the average vector **v**. Based on error propagation, the standard deviation in *v*, *s*^v^, can be evaluated from the standard deviations in *x*, *y*, and *z*, denoted respectively by *s*^x^, *s*^y^, and *s*^z^:

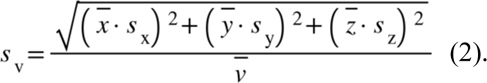

The standard error of the mean (SEM; *N* = 150) in the magnitude of the average vector was evaluated to assess statistical significance:

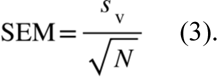

A response was only considered significant if the magnitude was greater than zero by at least 2 × SEM. Insignificant responses were treated as zero. The deviation vectors were visualised on the pre-MD M^pro^-peptide complex with PyMOL (v 2.3.0)^58^ and the modevectors.py script.^59^ The five active-site distances and ten HB *d*^D-A^ distances were output from both the equilibrium (*d*^eqm^) and nonequilibrium (*d*^neq^) propagations. The distance differences Δ*d* = *d*^neq^ – *d*^eqm^ at equivalent timepoints were calculated, plotted as a violin plot to visualise the distributions, and averaged across the 150 replicas with statistical significance assessed by SEM.

## Data Availability

Detailed setup and analyses of the equilibrium and nonequilibrium MD simulations are included in the **Supplementary Information**. Simulation data, including input files, M^pro^-peptide complex structures, and trajectories, are available on GitHub (https://github.com/duartegroup/Mpro_per_Subsite_D-NEMD).

## Author Contributions

All authors conceptualised the study. HTHC carried out the MD and D-NEMD simulations and analyses with assistance from ASFO on analyses. HTHC wrote the original draft. All authors contributed to discussions and review of the manuscript. CJS and FD supervised the study.

## Declaration of Interest

There are no conflicts to declare.

## Acknowledgements

HTHC thanks the Clarendon Fund, New College Oxford, and the EPSRC Centre for Doctoral Training in Synthesis for Biology and Medicine (EP/L015838/1) for a studentship, generously supported by AstraZeneca, Diamond Light Source, Defence Science and Technology Laboratory, Evotec, GlaxoSmithKline, Janssen, Novartis, Pfizer, Syngenta, Takeda, UCB and Vertex. AJM and ASFO thank EPSRC (grant number EP/M022609/1), BBSRC (grant numbers BB/R016445/1, BB/X009831/1 and BB/W003449/1) and ERC (Advanced Grant PREDACTED https://cordis.europa.eu/project/id/101021207) for support (funding from the European Research Council (ERC) under the European Union’s Horizon 2020 research and innovation programme, Grant agreement No. 101021207). ASFO also thanks Oracle for Research for the Research Fellowship. This project made use of time on HPC granted via the UK High-End Computing Consortium for Biomolecular Simulation, HECBioSim (http://hecbiosim.ac.uk), supported by EPSRC (grant no. EP/R029407/1).

## Notes

https://github.com/duartegroup/Mpro_per_Subsite_D-NEMD

### Competing Interest Statement

The authors have declared no competing interest.

