## Supplementary material for "Substrate recognition and selectivity in SARS-CoV-2 main protease: Unveiling the role of subsite interactions through dynamical nonequilibrium molecular dynamics simulations": SI

### Section S1: Equilibrium MD simulations of M^pro-peptide complexes

The peptides considered here include the substrate peptides s01 (nsp4/5) and s02 (nsp5/6), the inhibitor peptide p12,[1] and the P1 Gln-to-Ala substituted peptides s01-QP1A and s05-QP1A. The equilibrium molecular dynamics (MD) simulations for s05 (nsp8/9) have been reported.[2] The analyses in **Figures S1.1-15** are presented similar to those previously reported.[2]

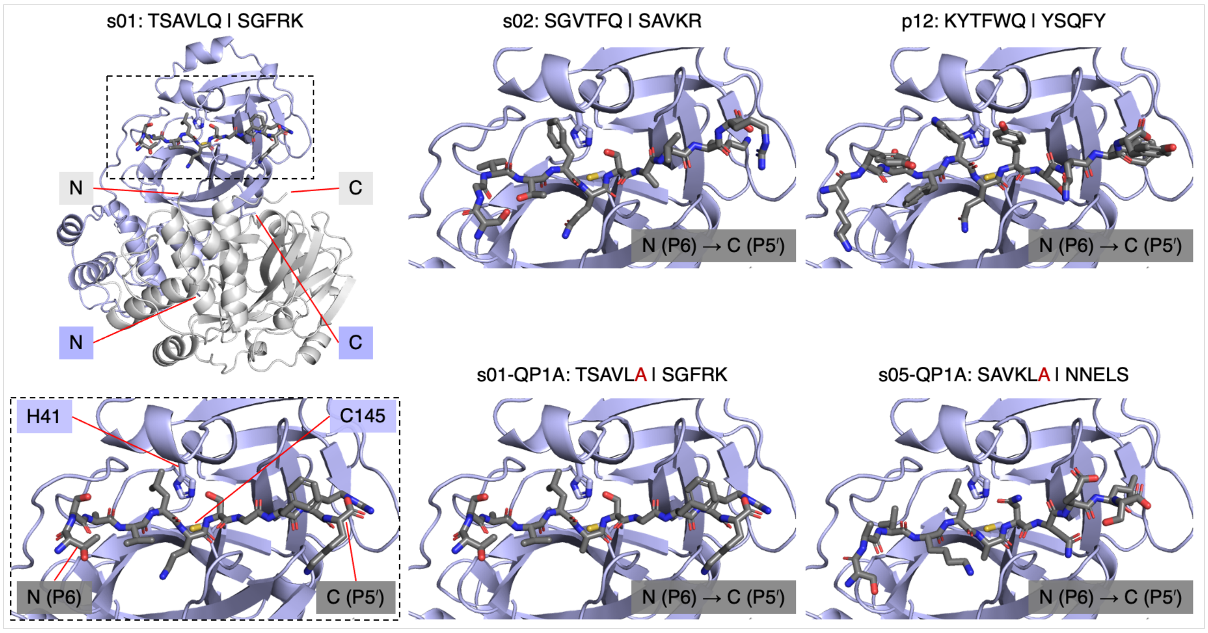

**Figure S1.1**: View of the starting non-covalent complexes of dimeric M^pro (chains A and B shown as blue and grey cartoons, respectively, with the active site His41 and Cys145 shown as sticks and labelled in the s01-bound complex; PDB 6YB7)[3] and the comparatively modelled s01, s02, p12, s01-QP1A, and s05-QP1A peptides (all shown as dark grey sticks).[1] Hydrogens are omitted for clarity. Chains A and B are subsequently referred to as ChA and ChB, respectively. The N- and C-terminal residues of each chain are labelled in the s01-bound complex.

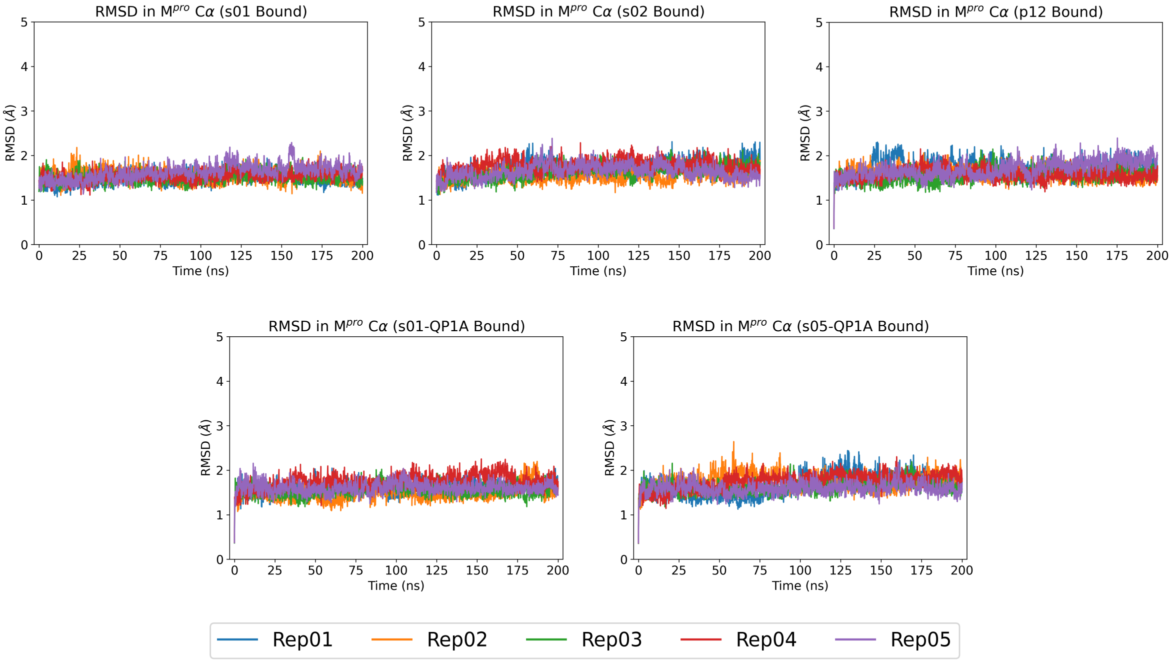

**Figure S1.2**: Time evolution of M^pro Cα root mean square deviation (RMSD) in each of the five 200 ns production molecular dynamics (MD) simulations relative to the crystallographically observed structure (PDB 6YB7)[3] for each M^pro-peptide complex. Each simulation was subjected to 200 ps NVT equilibration, during which time the protein and peptide non-hydrogen atoms were restrained. This was followed by 200 ps NPT equilibration, during which time the restraints were released for the native substrate (s01, s02) bound M^pro systems, and backbone restraints were retained in the other cases (p12, s01-QP1A, s05-QP1A bound M^pro). Note that due to protein dynamic movements during the equilibration phases, the RMSD measured at $t$ = 0 relative to crystal structure was non-zero.

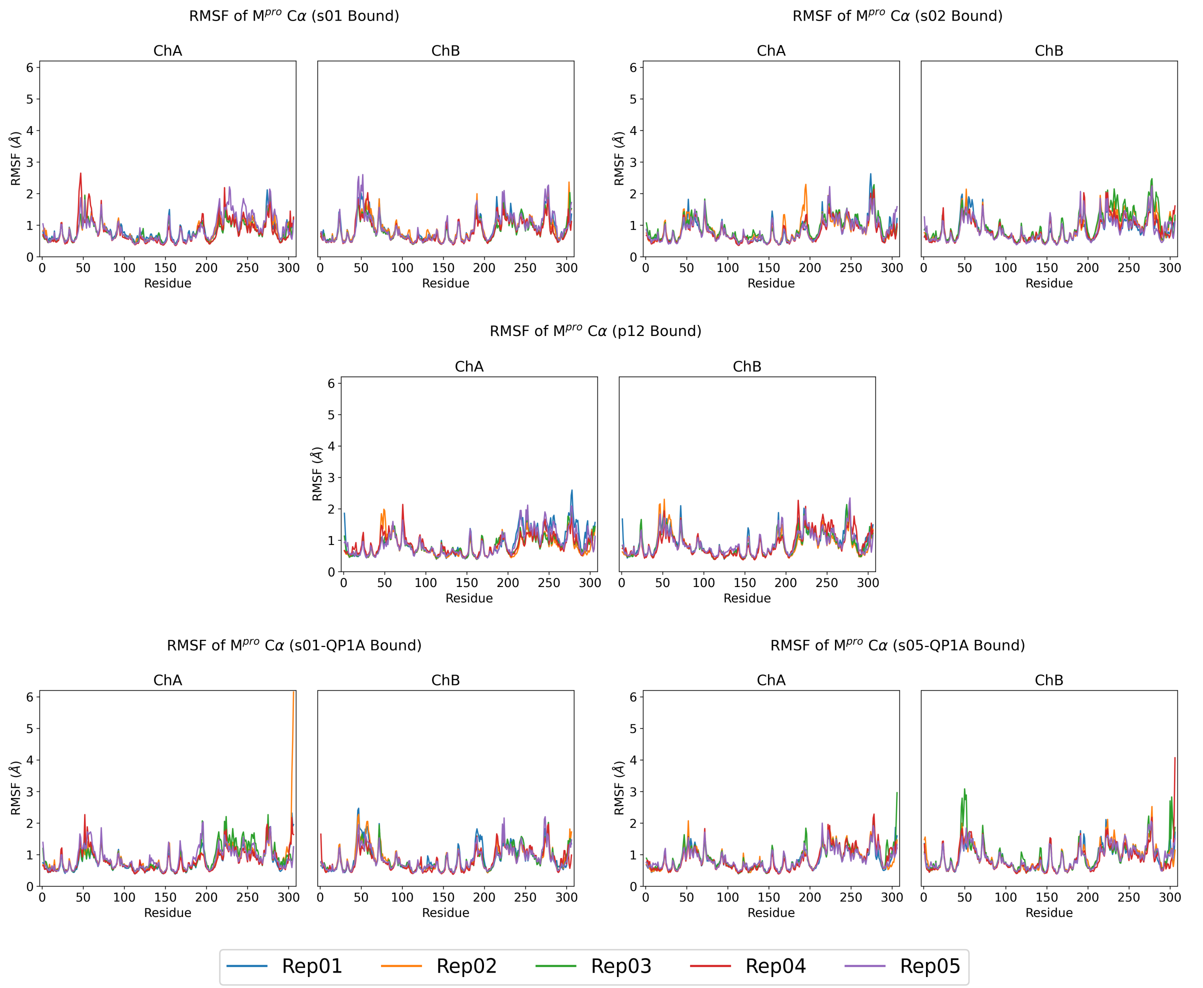

**Figure S1.3**: M$^{pro}$ Cα root mean square fluctuation (RMSF) in each of the five 150 ns equilibrium MD simulations, for each M$^{pro}$-peptide complex. The first 50 ns of each 200 ns production MD simulation was discarded as equilibration.

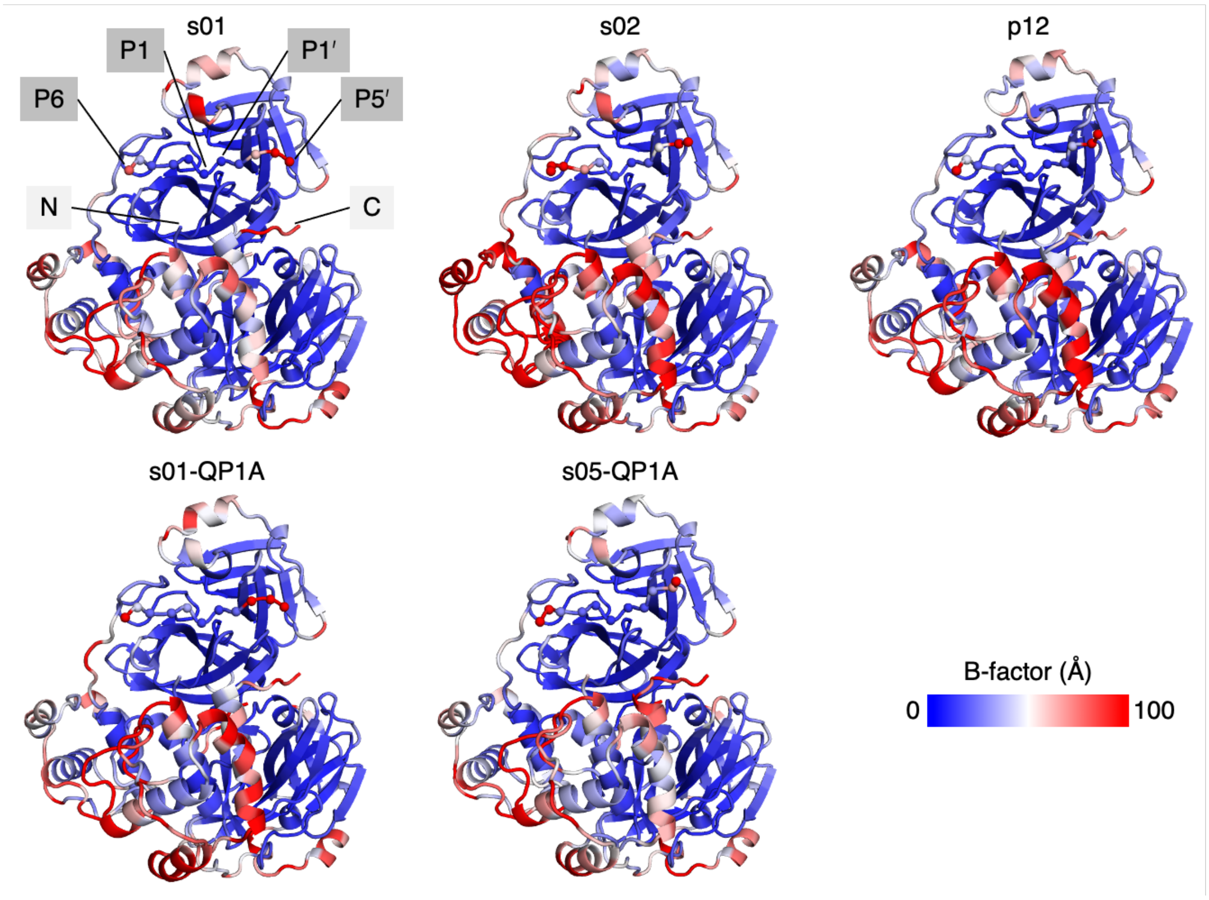

**Figure S1.4**: Views of the global average structure of each M$^{pro}$-peptide complex over the combined 5 × 150 ns equilibrium MD simulations. The colouring is based on a 0-100 Å$^2$ blue-white-red scale of the MD-derived B-factors, with rigid regions in blue and flexible regions in red. The N- and C-terminal residues of ChB are labelled in the s01-bound complex. The peptide Cα atoms are shown as spheres, with the terminal (P6 and P5′) and the scissile amide (between P1 and P1′) positions labelled for s01.

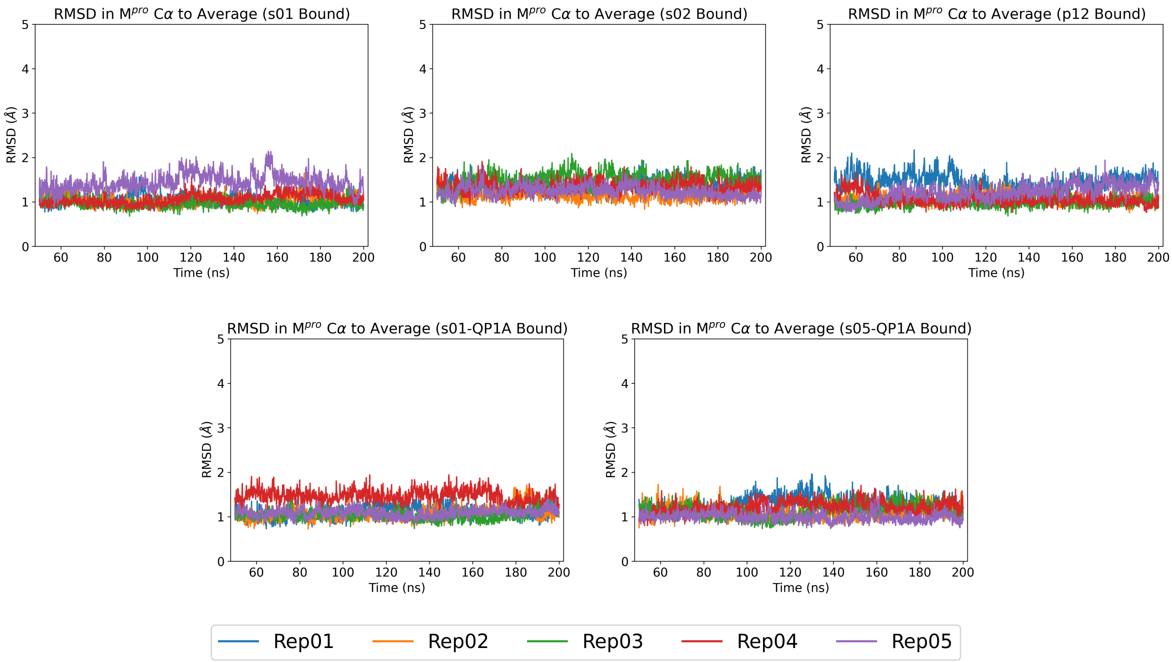

**Figure S1.5**: Time evolution of M$^{pro}$ Cα RMSD in each of the five 150 ns equilibrium MD simulations (starting from $t$ = 50 ns), relative to the equilibrium MD-derived global average structure of the complex, for each M$^{pro}$-peptide complex.

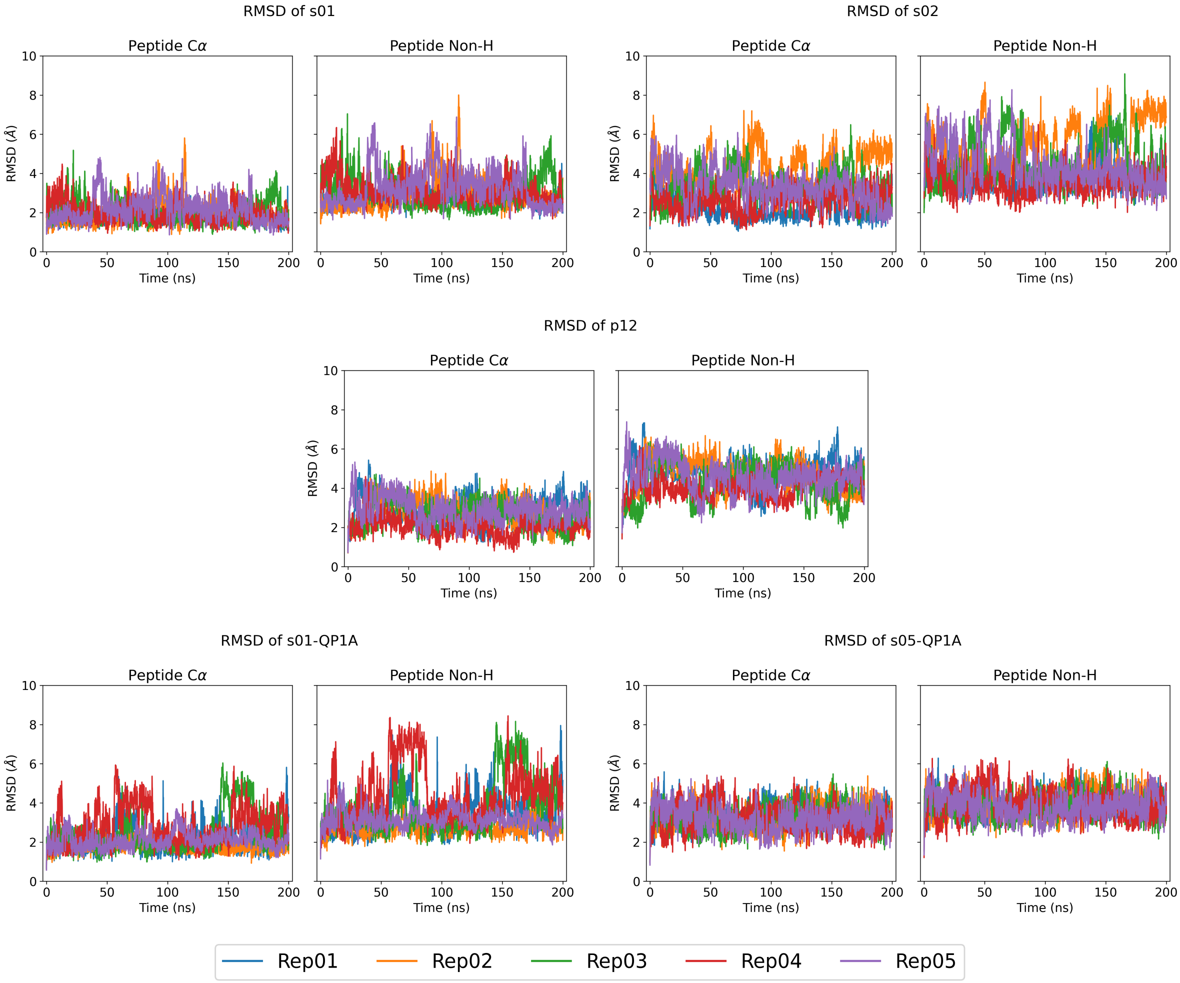

**Figure S1.6**: Time evolution of the peptide Cα and non-hydrogen atom RMSDs over the 5 × 200 ns production MD simulations relative to the starting conformation, with trajectories fitted based on $M^{pro}$ Cα atoms, for each $M^{pro}$-peptide complex.

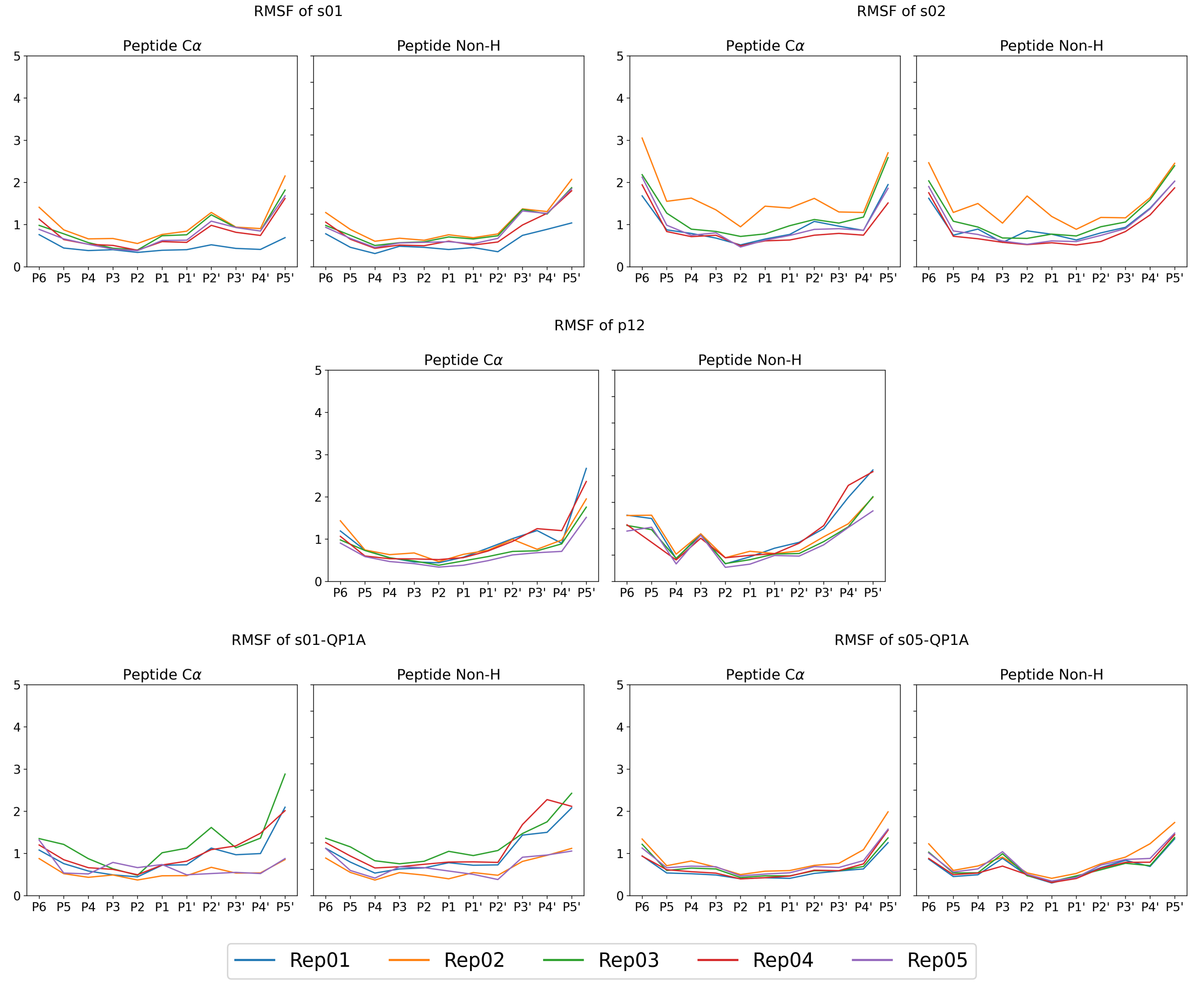

**Figure S1.7**: Per-residue RMSF of the peptide (left) Cα and (right) non-hydrogen atoms in the 5 × 150 ns equilibrium MD simulations, for each M$^{pro}$-peptide complex.

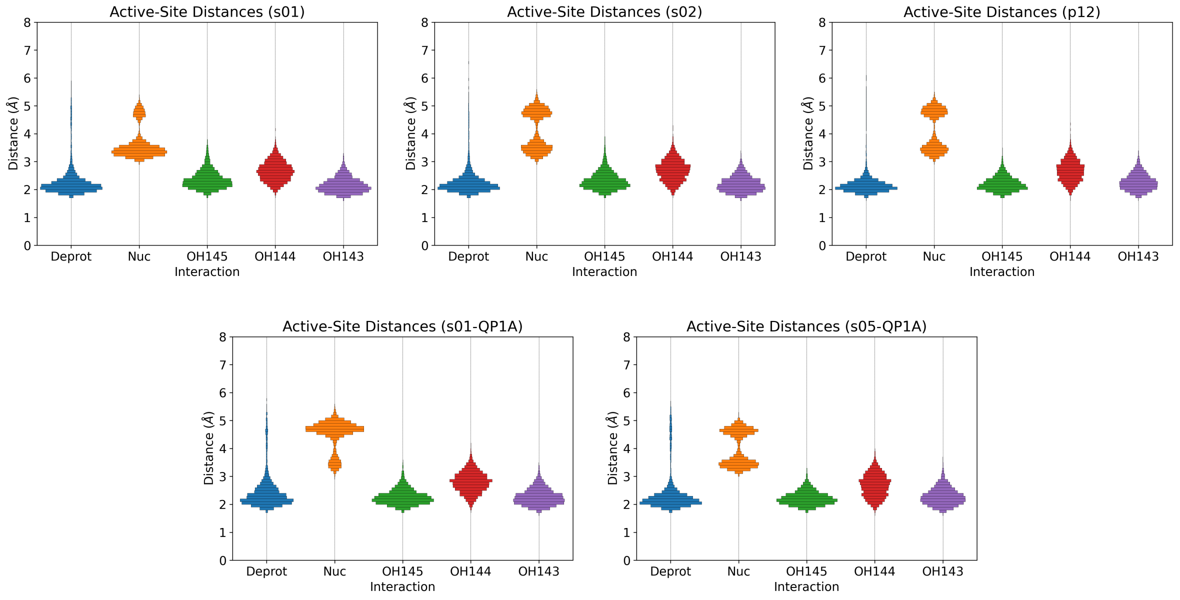

**Figure S1.8**: Distributions of distances in the active site that relate to the initiation of reaction between M$^{pro}$ Cys145 and the scissile amide bond, observed over the combined 5 × 150 ns equilibrium MD simulations, for each M$^{pro}$-peptide complex. These include the His41-Nϵ2 – Cys145-Hγ distance ($d_{Deprot}$, blue); the Cys145-Sγ – peptide P1-C distance ($d_{Nuc}$, orange); and the distances from the peptide scissile amide carbonyl P1-O to the backbone amide H atoms of Cys145 ($d_{OH145}$, green), Ser144 ($d_{OH144}$, red), and Gly143 ($d_{OH143}$, purple), which together form the oxyanion hole (see **main text Figure 2**).

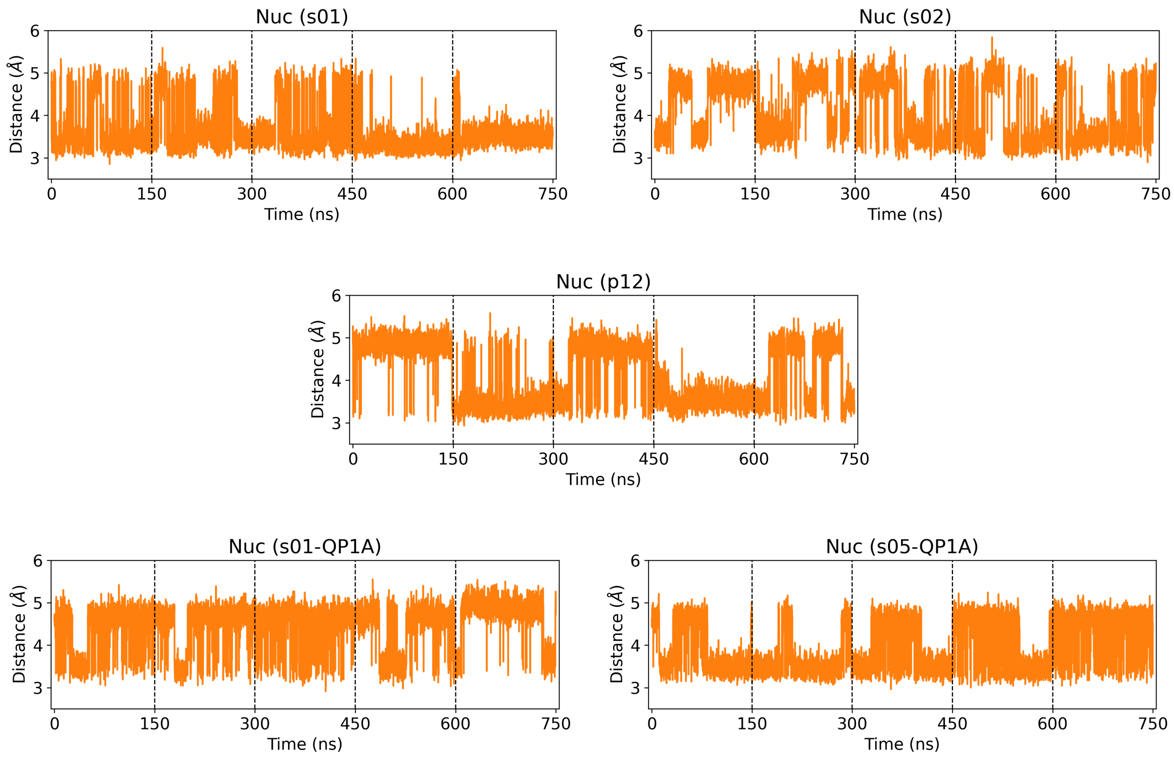

**Figure S1.9**: Time evolution of $d_{\mathrm{Nuc}}$ over the combined 5 × 150 ns equilibrium MD simulations.

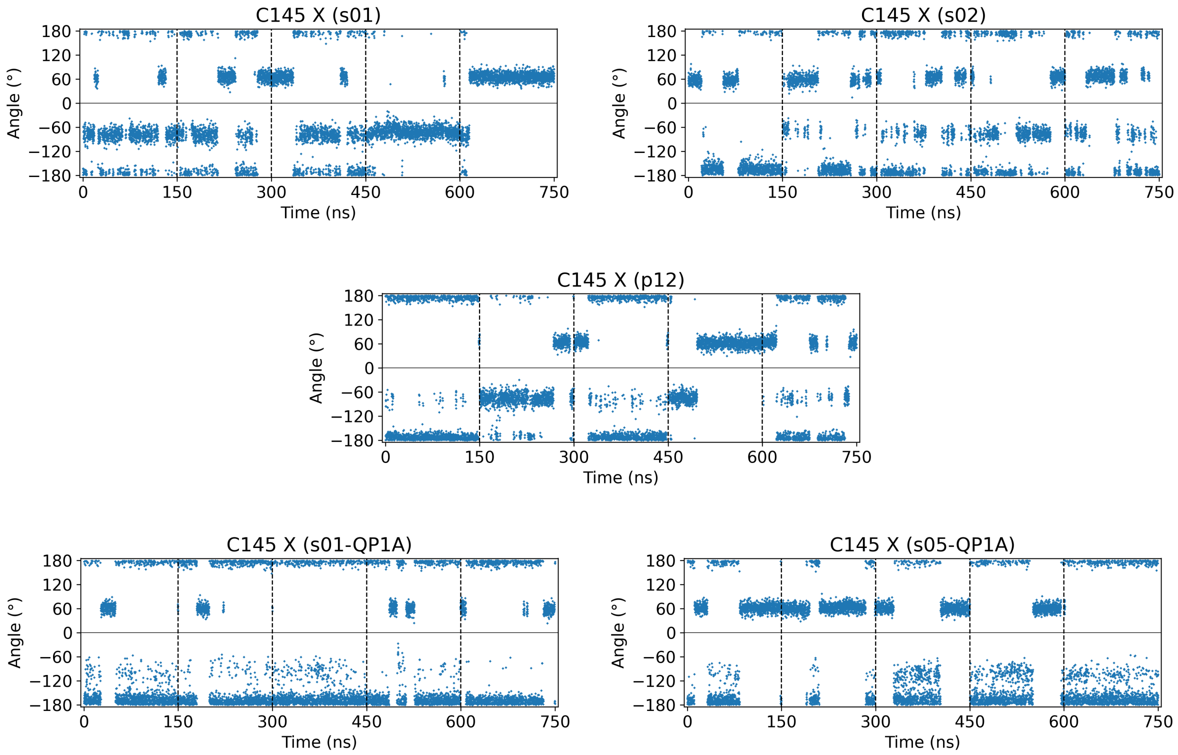

**Figure S1.10**: Time evolution of the Cys145 $\chi$ dihedral (N-Cα-Cβ-Sγ) over the combined 5 × 150 ns equilibrium MD simulations. By comparison to the time evolution of $d_{\mathrm{Nuc}}$ (**Figure S1.9**), $d_{\mathrm{Nuc}}$ is short (~3.5 Å) when Cys145 adopts a *gauche* conformation ($\chi$ centred around ±60°), and long (~5.0 Å) when Cys145 adopts a *trans* geometry ($\chi$ centred around ±180°).

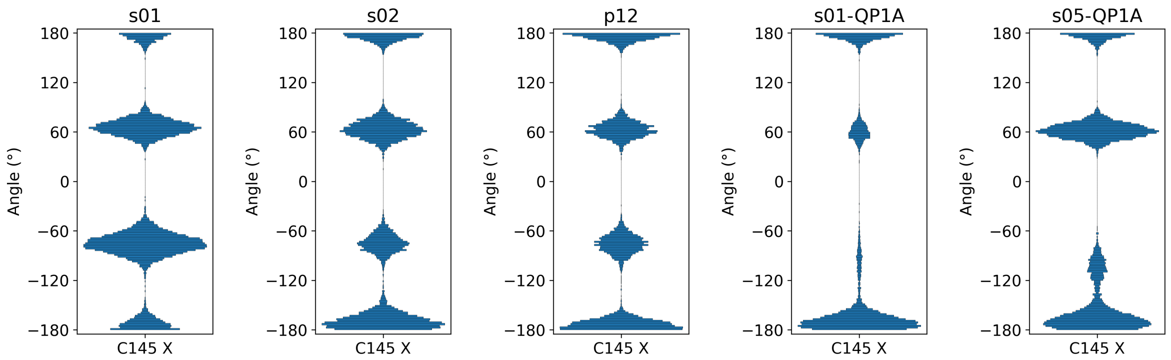

**Figure S1.11**: Distributions of the Cys145 χ dihedral (N-Cα-Cβ-Sγ) over the combined 5 × 150 ns equilibrium MD simulations. By defining the *gauche* conformer to be when -120° < χ < 120° and the *trans* conformer otherwise, the *gauche*:*trans* percentage population ratio observed when M^pro is complexed with s01, s02, p12, s01-QP1A, and s05-QP1A are 82:18, 53:47, 52:48, 19:81, 52:48, respectively. For comparison, the *gauche*:*trans* ratio in the reported equilibrium MD simulations of the M^pro-s05 complex[2] is 68:32.

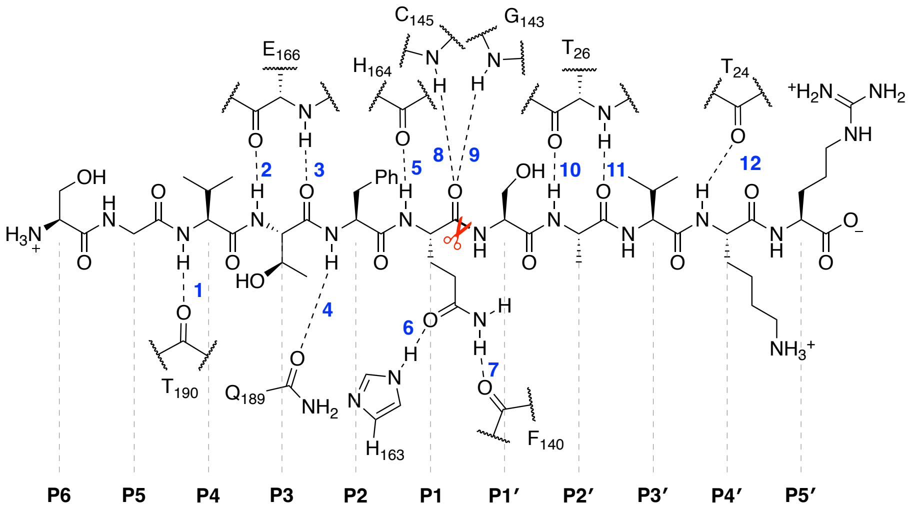

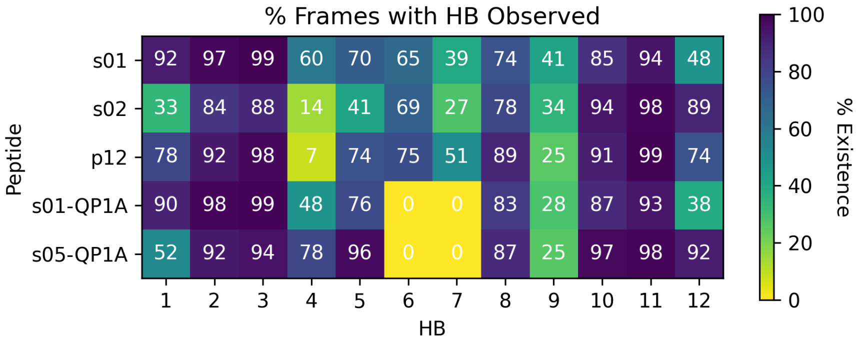

**Figure S1.12**: (top) Illustration of the 12 conserved M^pro-substrate hydrogen bonds (HBs)[1] exemplified by s02; (bottom) a heatmap showing the percentage of frames (analysed every ns) where the HB is observed, over the combined 5 × 150 ns equilibrium MD simulations, for each M^pro-peptide complex. Note that HBs 6 and 7 are dependent on the P1-Gln sidechain and, thus, are not formed in the cases of s01-QP1A and s05-QP1A.

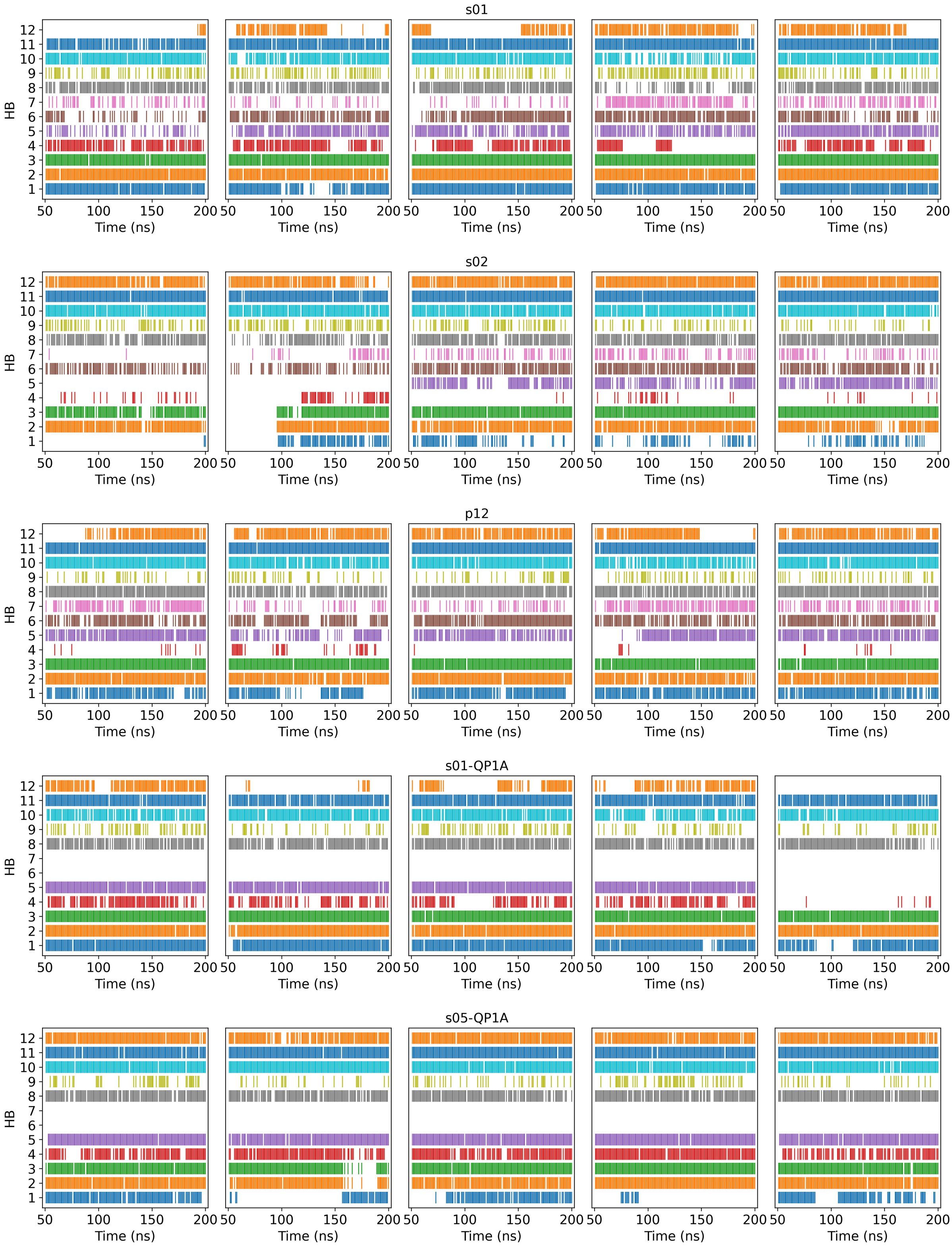

**Figure S1.13**: Event plots showing the presence of the 12 conserved HBs (frames analysed every ns) in each of the five 150 ns equilibrium MD simulations for each M$^{pro}$-peptide complex.

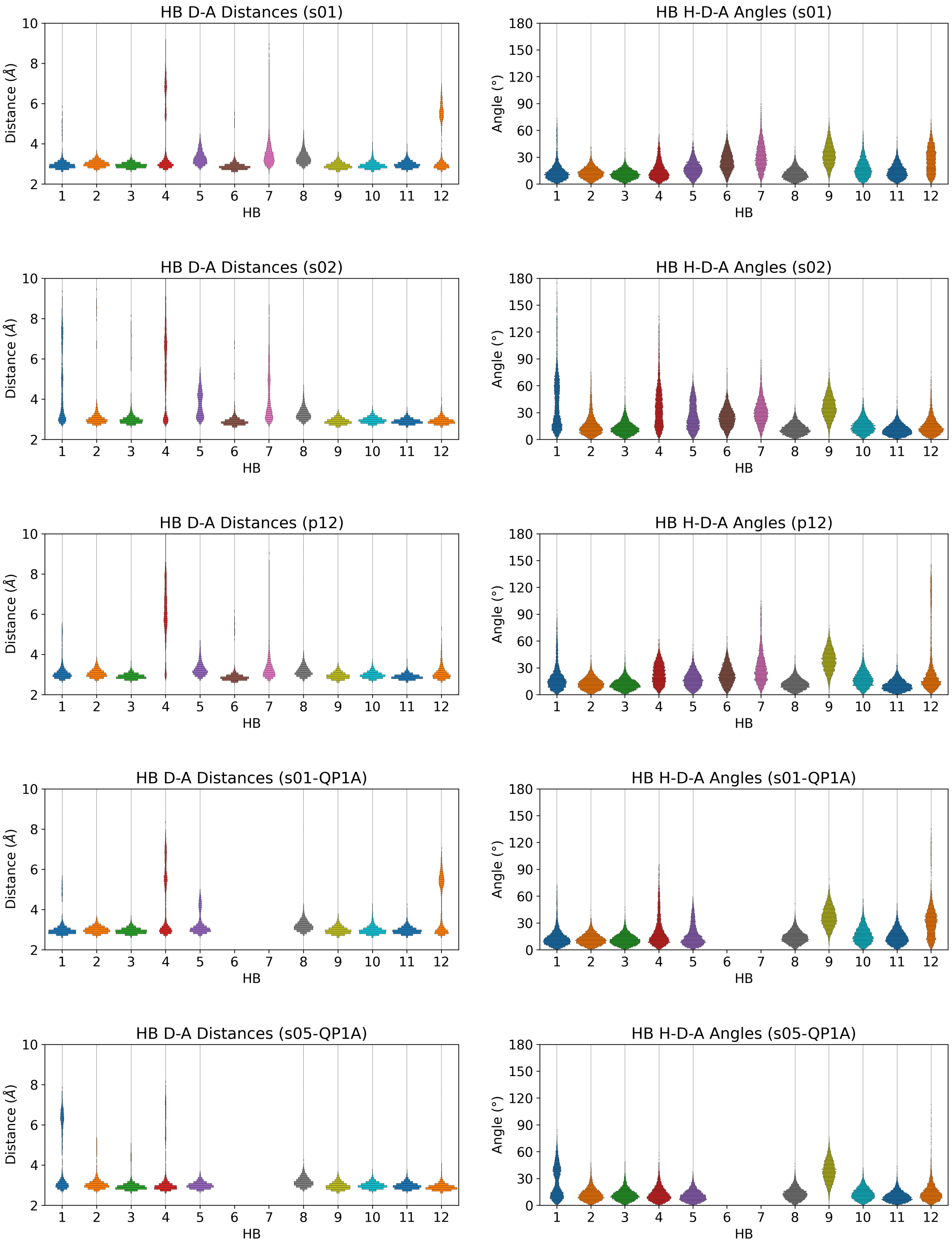

**Figure S1.14**: Distributions of (left) the donor-acceptor distances ($d_{\text{D-A}}$) and (right) the hydrogen-donor-acceptor angles ($\angle_{\text{H-D-A}}$) for each of the 12 conserved HBs, observed over the combined 5 × 150 ns equilibrium MD simulations, for each $M^{pro}$-peptide complex. HBs are defined based on a combined distance ($d_{\text{D-A}} \leq 3.5$ Å) and angle ($\angle_{\text{H-D-A}} \leq 30°$) criteria.

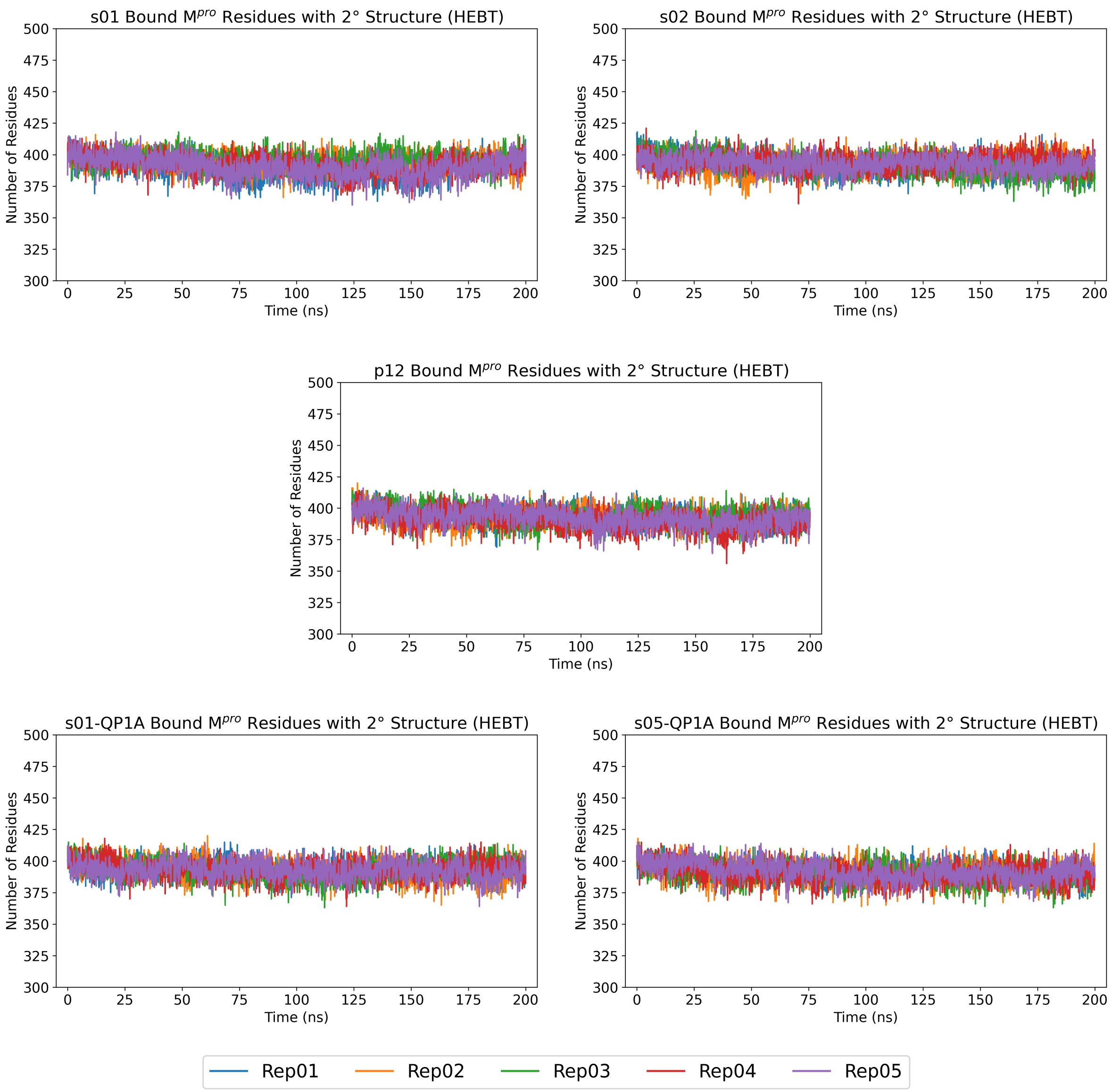

**Figure S1.15**: Time evolution of the number of $M^{pro}$ residues possessing secondary structure (H = α-helix; E = extended strand; B = β-bridge; T = hydrogen bonded turn) in each of the five 200 ns production MD simulations for each $M^{pro}$-peptide complex, as defined by DSSP (v 2.0.4).[4,5]

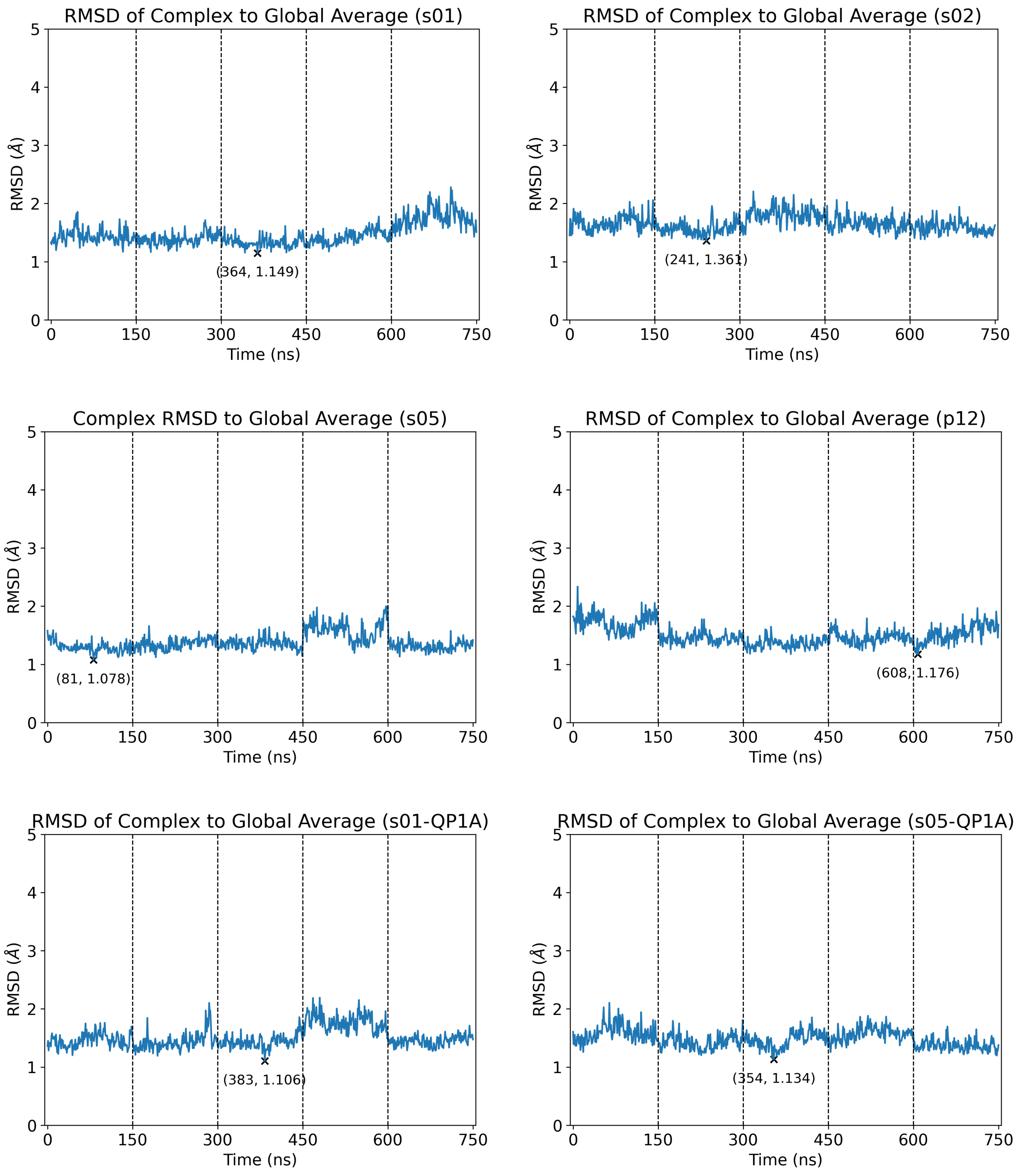

**Figure S1.16**: Time evolution of the M$^{pro}$-peptide (s01, s02, s05, p12, s01-QP1A, s05-QP1A) complex RMSD in the combined 5 × 150 ns equilibrium MD simulations (starting from $t$ = 50 ns; frames analysed every ns), relative to the equilibrium MD-derived global average structure of the complex, for each M$^{pro}$-peptide complex. The frame with the lowest RMSD is labelled and considered as a representative structure of the complex.

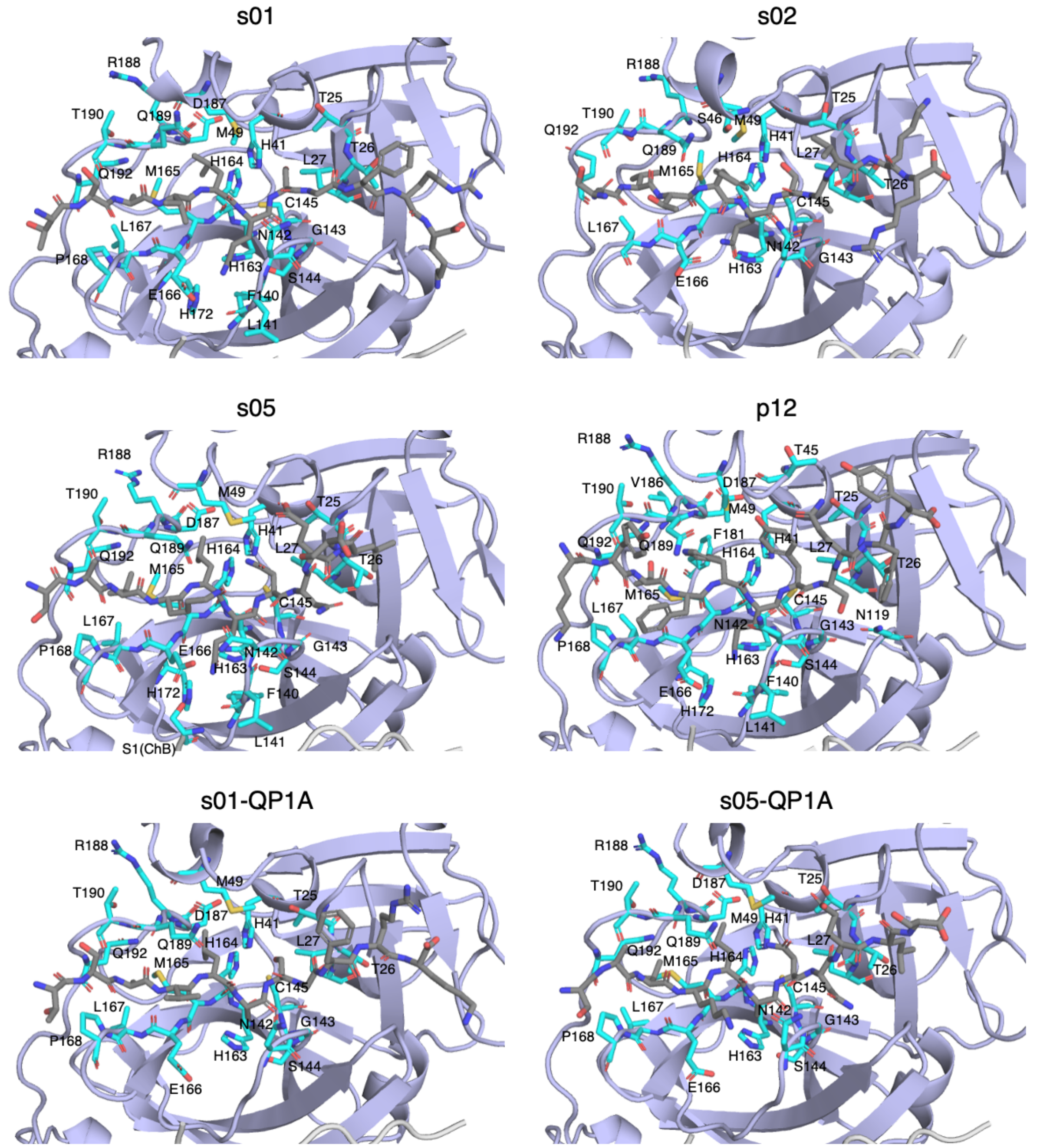

**Figure S1.17**: View of the representative structures of each M$^{pro}$-peptide complex. M$^{pro}$ residues located within 3 Å of the peptide P4-P2′ residues in the structure are shown as cyan sticks and labelled. Hydrogen atoms and water molecules are omitted for clarity.

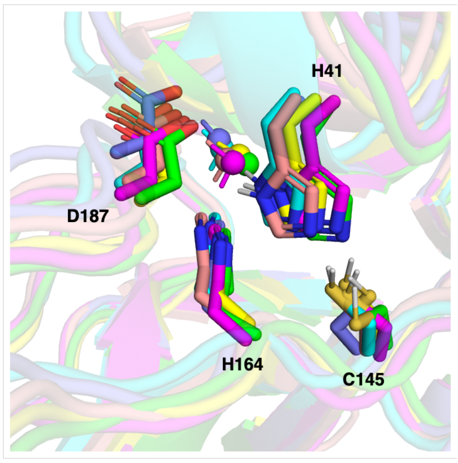

**Figure S1.18**: Superimposed views of the conserved bridging water molecule (BW) which bridges His41, His164, and His187, in the representative structures of M$^{pro}$ complexed with s01 (green), s02 (cyan), s05 (magenta), p12 (slate), s01-QP1A (yellow), and s05-QP1A (salmon), aligned using the M$^{pro}$ Cα atoms. BW is in the same colour as the protein carbon atoms, with its oxygen atom shown as a sphere. Carbon-bonded hydrogen atoms are omitted for clarity.

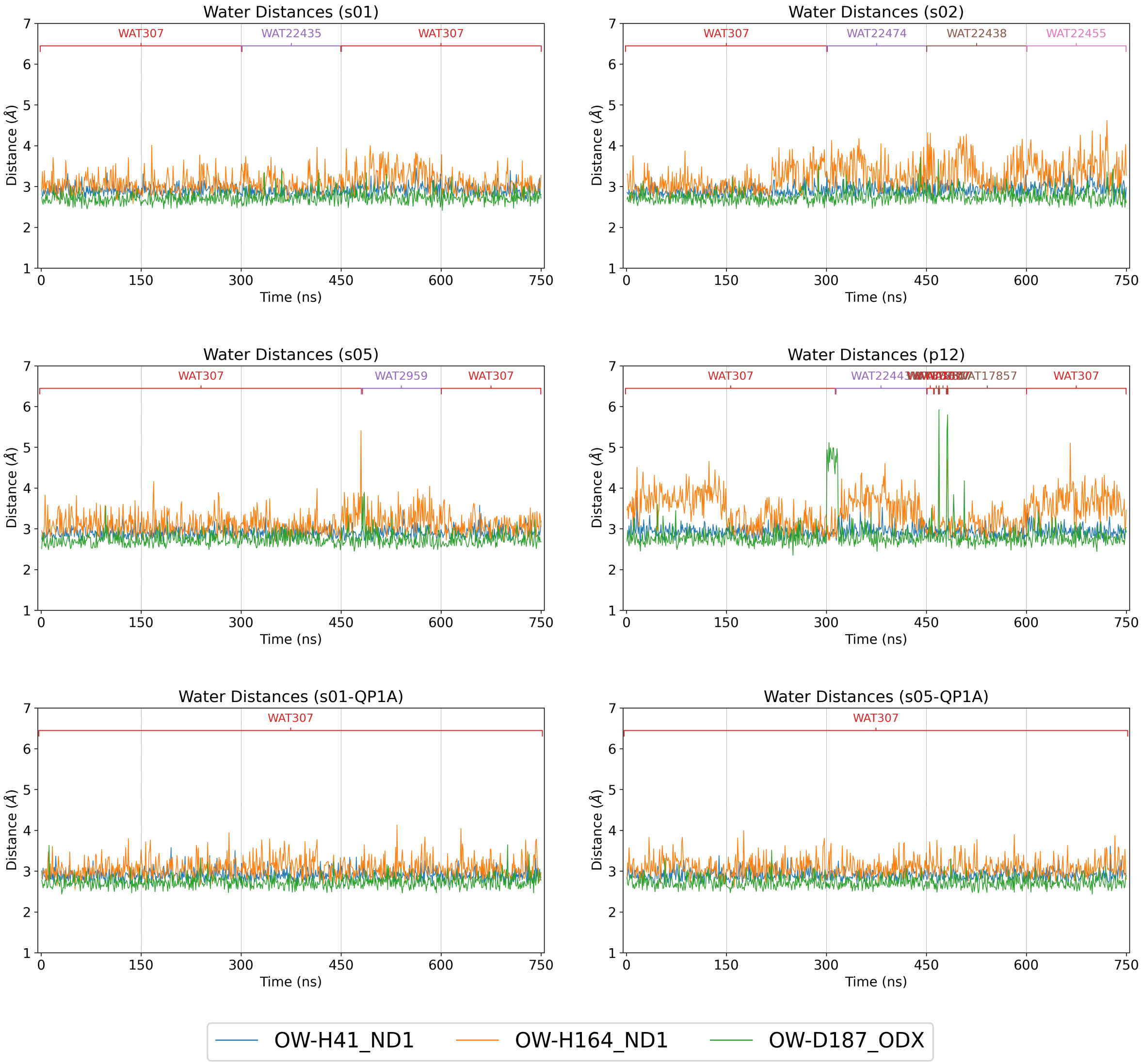

**Figure S1.19**: Time evolution of the distances between the oxygen atom of the conserved water molecule BW (the water closest to His41_Nδ1 is considered in each frame, analysed every ns), and His41_Nδ1, His164_Nδ1, and Asp187_Oδ (taking the carboxylate oxygen atom closest to the water) over the combined 5 × 150 ns equilibrium MD simulations.

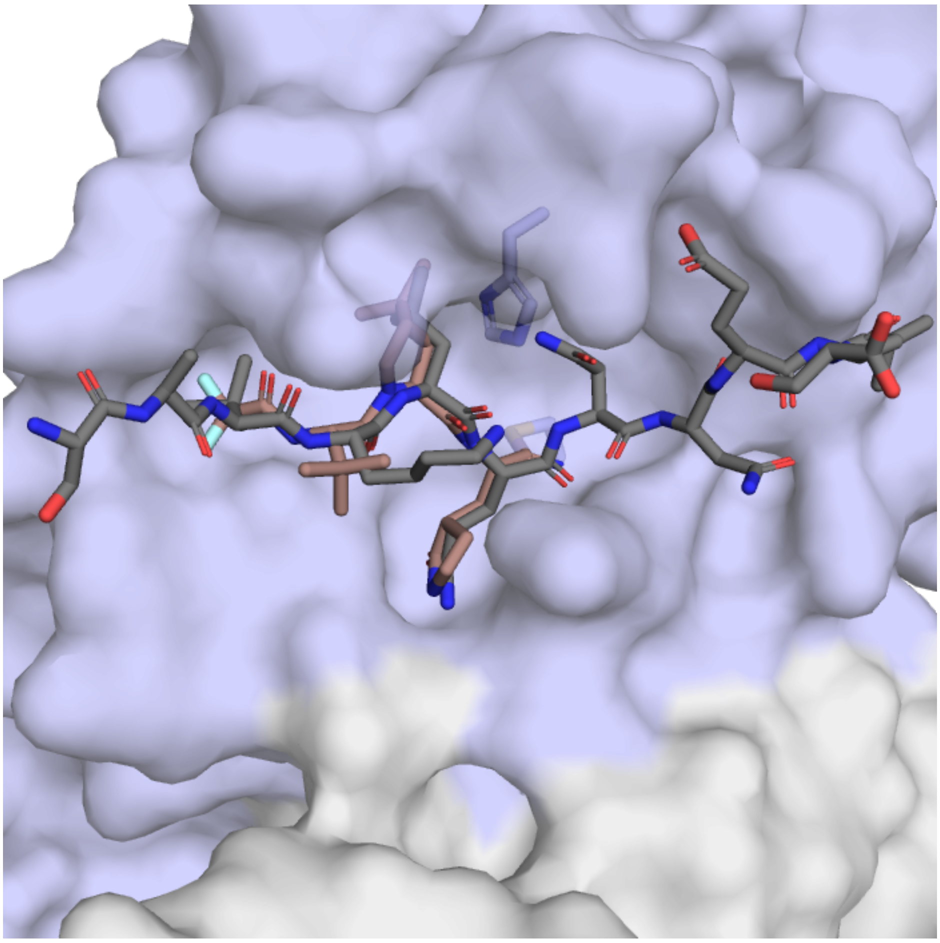

**Figure S1.20**: View of crystallographically observed dimeric $M^{pro}$ (chains A and B respectively shown as blue and light grey surfaces; His41-Cys145 dyad as sticks) with Cys145 covalently modified by nirmatrelvir (brown sticks) (PDB 7VH8; 1.59 Å resolution),[6] in comparison with the binding mode of s05 (dark grey sticks) in the equilibrium MD-derived representative structure, aligned using $M^{pro}$ Cα atoms (RMSD = 1.095 Å). Hydrogens are omitted for clarity.

### Section S2: Effect of the P1 Gln-to-Ala substitution

**Table S2.1**: The average differences in the five active-site distances ($\Delta d = d_{neq} - d_{eqm}$) following the P1 Gln to Ala substitution in s01 and s05. Positive and negative differences with magnitudes > 0.1 Å are in green and red, respectively.

| **Active Site** | s01: P1 Q->A | | | | s05: P1 Q->A | | |
|---|---|---|---|---|---|---|---|
| **Deprot** | t (ps) | Av Diff (Å) | SEM (Å) | | t (ps) | Av Diff (Å) | SEM (Å) |
| | 0.05 | 0.00 | 0.00 | | 0.05 | 0.00 | 0.00 |
| | 0.1 | -0.01 | 0.00 | | 0.1 | 0.00 | 0.00 |
| | 0.25 | -0.04 | 0.01 | | 0.25 | -0.03 | 0.01 |
| | 0.5 | -0.01 | 0.01 | | 0.5 | -0.01 | 0.01 |
| | 1 | -0.06 | 0.04 | | 1 | -0.05 | 0.04 |
| | 2 | -0.10 | 0.04 | | 2 | -0.06 | 0.05 |
| | 3 | -0.02 | 0.06 | | 3 | -0.14 | 0.06 |
| | 5 | -0.08 | 0.05 | | 5 | -0.07 | 0.04 |
| | 10 | -0.01 | 0.06 | | 10 | -0.06 | 0.06 |
| | 20 | -0.05 | 0.04 | | 20 | -0.04 | 0.06 |
| | 30 | -0.07 | 0.06 | | 30 | -0.02 | 0.05 |
| | 50 | -0.08 | 0.05 | | 50 | -0.10 | 0.06 |
| | 100 | -0.06 | 0.06 | | 100 | -0.17 | 0.07 |
| | 300 | -0.10 | 0.08 | | 300 | -0.05 | 0.05 |
| | 500 | -0.04 | 0.06 | | 500 | -0.05 | 0.06 |
| | 1000 | -0.03 | 0.06 | | 1000 | -0.04 | 0.07 |
| **Nuc** | t (ps) | Av Diff (Å) | SEM (Å) | | t (ps) | Av Diff (Å) | SEM (Å) |
| | 0.05 | 0.01 | 0.00 | | 0.05 | 0.02 | 0.00 |
| | 0.1 | 0.03 | 0.00 | | 0.1 | 0.03 | 0.00 |
| | 0.25 | -0.01 | 0.01 | | 0.25 | -0.04 | 0.01 |
| | 0.5 | 0.01 | 0.01 | | 0.5 | 0.01 | 0.01 |
| | 1 | 0.00 | 0.02 | | 1 | -0.01 | 0.01 |
| | 2 | 0.04 | 0.03 | | 2 | -0.03 | 0.02 |
| | 3 | 0.00 | 0.03 | | 3 | -0.02 | 0.02 |
| | 5 | 0.02 | 0.03 | | 5 | -0.02 | 0.03 |
| | 10 | 0.04 | 0.04 | | 10 | 0.07 | 0.04 |
| | 20 | 0.23 | 0.05 | | 20 | 0.10 | 0.05 |
| | 30 | 0.22 | 0.05 | | 30 | 0.05 | 0.04 |
| | 50 | 0.30 | 0.05 | | 50 | 0.12 | 0.05 |
| | 100 | 0.37 | 0.06 | | 100 | 0.19 | 0.05 |
| | 300 | 0.32 | 0.06 | | 300 | 0.13 | 0.04 |
| | 500 | 0.44 | 0.06 | | 500 | 0.12 | 0.05 |
| | 1000 | 0.52 | 0.06 | | 1000 | 0.10 | 0.05 |
| **OH145** | t (ps) | Av Diff (Å) | SEM (Å) | | t (ps) | Av Diff (Å) | SEM (Å) |
| | 0.05 | 0.00 | 0.00 | | 0.05 | 0.00 | 0.00 |
| | 0.1 | 0.02 | 0.00 | | 0.1 | 0.03 | 0.00 |
| | 0.25 | -0.01 | 0.01 | | 0.25 | -0.02 | 0.01 |
| | 0.5 | -0.02 | 0.02 | | 0.5 | 0.01 | 0.01 |
| | 1 | -0.02 | 0.02 | | 1 | -0.04 | 0.02 |
| | 2 | -0.02 | 0.03 | | 2 | -0.02 | 0.02 |
| | 3 | 0.00 | 0.03 | | 3 | -0.03 | 0.02 |
| | 5 | -0.03 | 0.03 | | 5 | -0.01 | 0.02 |
| | 10 | -0.03 | 0.03 | | 10 | -0.02 | 0.03 |
| | 20 | -0.04 | 0.03 | | 20 | -0.03 | 0.03 |
| | 30 | -0.03 | 0.03 | | 30 | 0.02 | 0.03 |
| | 50 | -0.07 | 0.03 | | 50 | -0.01 | 0.02 |
| | 100 | -0.08 | 0.04 | | 100 | 0.00 | 0.03 |
| | 300 | -0.03 | 0.03 | | 300 | -0.03 | 0.03 |
| | 500 | -0.08 | 0.03 | | 500 | 0.01 | 0.03 |
| | 1000 | -0.03 | 0.03 | | 1000 | 0.01 | 0.03 |
| **OH144** | t (ps) | Av Diff (Å) | SEM (Å) | | t (ps) | Av Diff (Å) | SEM (Å) |
| | 0.05 | -0.01 | 0.00 | | 0.05 | 0.00 | 0.00 |
| | 0.1 | -0.03 | 0.00 | | 0.1 | -0.02 | 0.00 |
| | 0.25 | -0.05 | 0.01 | | 0.25 | -0.05 | 0.01 |
| | 0.5 | -0.04 | 0.02 | | 0.5 | -0.01 | 0.01 |
| | 1 | -0.04 | 0.03 | | 1 | -0.08 | 0.03 |
| | 2 | -0.08 | 0.03 | | 2 | -0.07 | 0.03 |
| | 3 | -0.06 | 0.03 | | 3 | -0.07 | 0.03 |
| | 5 | -0.02 | 0.04 | | 5 | -0.04 | 0.03 |
| | 10 | 0.05 | 0.04 | | 10 | -0.01 | 0.03 |
| | 20 | -0.11 | 0.04 | | 20 | 0.00 | 0.04 |
| | 30 | 0.02 | 0.04 | | 30 | 0.00 | 0.03 |
| | 50 | 0.00 | 0.04 | | 50 | 0.00 | 0.04 |
| | 100 | 0.04 | 0.04 | | 100 | -0.05 | 0.03 |
| | 300 | 0.08 | 0.04 | | 300 | -0.04 | 0.03 |
| | 500 | 0.06 | 0.04 | | 500 | -0.01 | 0.04 |
| | 1000 | 0.06 | 0.04 | | 1000 | -0.01 | 0.04 |
| **OH143** | t (ps) | Av Diff (Å) | SEM (Å) | | t (ps) | Av Diff (Å) | SEM (Å) |
| | 0.05 | 0.01 | 0.00 | | 0.05 | 0.00 | 0.00 |
| | 0.1 | -0.01 | 0.00 | | 0.1 | -0.03 | 0.00 |
| | 0.25 | 0.04 | 0.01 | | 0.25 | 0.03 | 0.01 |
| | 0.5 | 0.05 | 0.02 | | 0.5 | 0.03 | 0.01 |
| | 1 | 0.04 | 0.02 | | 1 | 0.04 | 0.03 |
| | 2 | 0.06 | 0.03 | | 2 | 0.04 | 0.03 |
| | 3 | 0.03 | 0.03 | | 3 | 0.06 | 0.03 |
| | 5 | 0.00 | 0.03 | | 5 | -0.02 | 0.03 |
| | 10 | 0.05 | 0.03 | | 10 | 0.04 | 0.03 |
| | 20 | 0.05 | 0.03 | | 20 | 0.03 | 0.03 |
| | 30 | 0.01 | 0.03 | | 30 | -0.01 | 0.03 |
| | 50 | 0.04 | 0.03 | | 50 | 0.04 | 0.03 |
| | 100 | 0.10 | 0.03 | | 100 | -0.04 | 0.03 |
| | 300 | 0.02 | 0.03 | | 300 | -0.02 | 0.03 |
| | 500 | 0.06 | 0.03 | | 500 | 0.02 | 0.03 |
| | 1000 | 0.07 | 0.03 | | 1000 | -0.01 | 0.03 |

**Table S2.2**: The average differences in the donor-acceptor distances of HBs 1-5 following the P1 Gln to Ala substitution in s01 and s05. Positive and negative differences with magnitudes $> 0.1$ Å are in green and red, respectively.

| HB1-5 | s01: P1 Q->A | | | | s05: P1 Q->A | | |
|---|---|---|---|---|---|---|---|
| **HB1** | t (ps) | Av Diff (Å) | SEM (Å) | | t (ps) | Av Diff (Å) | SEM (Å) |
| | 0.05 | 0.00 | 0.00 | | 0.05 | 0.00 | 0.00 |
| | 0.1 | 0.00 | 0.00 | | 0.1 | 0.00 | 0.00 |
| | 0.25 | 0.00 | 0.00 | | 0.25 | -0.01 | 0.00 |
| | 0.5 | 0.00 | 0.00 | | 0.5 | -0.01 | 0.00 |
| | 1 | -0.02 | 0.01 | | 1 | 0.00 | 0.02 |
| | 2 | -0.01 | 0.02 | | 2 | -0.02 | 0.02 |
| | 3 | 0.02 | 0.02 | | 3 | -0.05 | 0.03 |
| | 5 | 0.01 | 0.02 | | 5 | -0.01 | 0.03 |
| | 10 | -0.01 | 0.02 | | 10 | 0.00 | 0.03 |
| | 20 | 0.06 | 0.03 | | 20 | 0.04 | 0.03 |
| | 30 | 0.01 | 0.04 | | 30 | 0.03 | 0.03 |
| | 50 | 0.01 | 0.03 | | 50 | -0.03 | 0.03 |
| | 100 | -0.01 | 0.03 | | 100 | -0.05 | 0.04 |
| | 300 | 0.03 | 0.04 | | 300 | 0.03 | 0.04 |
| | 500 | -0.06 | 0.03 | | 500 | -0.07 | 0.04 |
| | 1000 | -0.02 | 0.04 | | 1000 | -0.06 | 0.04 |
| **HB2** | t (ps) | Av Diff (Å) | SEM (Å) | | t (ps) | Av Diff (Å) | SEM (Å) |
| | 0.05 | 0.00 | 0.00 | | 0.05 | 0.00 | 0.00 |
| | 0.1 | -0.01 | 0.00 | | 0.1 | -0.01 | 0.00 |
| | 0.25 | 0.01 | 0.00 | | 0.25 | 0.00 | 0.00 |
| | 0.5 | 0.01 | 0.01 | | 0.5 | -0.01 | 0.01 |
| | 1 | 0.00 | 0.01 | | 1 | 0.00 | 0.02 |
| | 2 | -0.01 | 0.02 | | 2 | 0.02 | 0.02 |
| | 3 | 0.00 | 0.01 | | 3 | -0.07 | 0.02 |
| | 5 | -0.02 | 0.02 | | 5 | -0.03 | 0.02 |
| | 10 | 0.01 | 0.02 | | 10 | -0.03 | 0.02 |
| | 20 | 0.01 | 0.02 | | 20 | -0.01 | 0.02 |
| | 30 | 0.00 | 0.02 | | 30 | -0.03 | 0.02 |
| | 50 | 0.00 | 0.02 | | 50 | 0.01 | 0.02 |
| | 100 | 0.02 | 0.02 | | 100 | 0.01 | 0.02 |
| | 300 | 0.02 | 0.02 | | 300 | -0.01 | 0.03 |
| | 500 | 0.02 | 0.02 | | 500 | 0.04 | 0.03 |
| | 1000 | 0.00 | 0.02 | | 1000 | 0.05 | 0.03 |
| **HB3** | t (ps) | Av Diff (Å) | SEM (Å) | | t (ps) | Av Diff (Å) | SEM (Å) |
| | 0.05 | -0.01 | 0.00 | | 0.05 | -0.01 | 0.00 |
| | 0.1 | -0.01 | 0.00 | | 0.1 | -0.01 | 0.00 |
| | 0.25 | 0.00 | 0.00 | | 0.25 | 0.00 | 0.00 |
| | 0.5 | -0.01 | 0.01 | | 0.5 | -0.01 | 0.01 |
| | 1 | -0.01 | 0.01 | | 1 | -0.01 | 0.01 |
| | 2 | -0.04 | 0.01 | | 2 | -0.01 | 0.02 |
| | 3 | -0.01 | 0.01 | | 3 | -0.03 | 0.02 |
| | 5 | -0.04 | 0.02 | | 5 | 0.01 | 0.02 |
| | 10 | -0.02 | 0.01 | | 10 | -0.01 | 0.02 |
| | 20 | -0.02 | 0.02 | | 20 | -0.02 | 0.02 |
| | 30 | -0.02 | 0.01 | | 30 | -0.02 | 0.01 |
| | 50 | 0.00 | 0.01 | | 50 | -0.01 | 0.01 |
| | 100 | 0.01 | 0.01 | | 100 | 0.01 | 0.01 |
| | 300 | -0.01 | 0.01 | | 300 | -0.02 | 0.02 |
| | 500 | 0.01 | 0.01 | | 500 | 0.03 | 0.02 |
| | 1000 | -0.02 | 0.02 | | 1000 | 0.00 | 0.02 |
| **HB4** | t (ps) | Av Diff (Å) | SEM (Å) | | t (ps) | Av Diff (Å) | SEM (Å) |
| | 0.05 | 0.00 | 0.00 | | 0.05 | 0.00 | 0.00 |
| | 0.1 | -0.02 | 0.00 | | 0.1 | -0.02 | 0.00 |
| | 0.25 | -0.01 | 0.00 | | 0.25 | 0.00 | 0.00 |
| | 0.5 | -0.02 | 0.01 | | 0.5 | 0.00 | 0.01 |
| | 1 | 0.02 | 0.03 | | 1 | 0.02 | 0.02 |
| | 2 | 0.01 | 0.03 | | 2 | 0.00 | 0.03 |
| | 3 | 0.03 | 0.03 | | 3 | 0.01 | 0.03 |
| | 5 | -0.01 | 0.03 | | 5 | -0.06 | 0.04 |
| | 10 | 0.01 | 0.03 | | 10 | -0.04 | 0.04 |
| | 20 | 0.06 | 0.04 | | 20 | 0.01 | 0.05 |
| | 30 | 0.00 | 0.06 | | 30 | -0.03 | 0.06 |
| | 50 | -0.02 | 0.07 | | 50 | -0.03 | 0.08 |
| | 100 | 0.11 | 0.08 | | 100 | 0.04 | 0.07 |
| | 300 | -0.03 | 0.11 | | 300 | 0.01 | 0.10 |
| | 500 | -0.04 | 0.13 | | 500 | 0.13 | 0.12 |
| | 1000 | 0.09 | 0.14 | | 1000 | 0.26 | 0.16 |
| **HB5** | t (ps) | Av Diff (Å) | SEM (Å) | | t (ps) | Av Diff (Å) | SEM (Å) |
| | 0.05 | 0.01 | 0.00 | | 0.05 | 0.01 | 0.00 |
| | 0.1 | -0.02 | 0.00 | | 0.1 | -0.03 | 0.00 |
| | 0.25 | -0.09 | 0.01 | | 0.25 | -0.11 | 0.01 |
| | 0.5 | -0.08 | 0.01 | | 0.5 | -0.06 | 0.01 |
| | 1 | -0.10 | 0.02 | | 1 | -0.08 | 0.02 |
| | 2 | -0.11 | 0.03 | | 2 | -0.09 | 0.03 |
| | 3 | -0.15 | 0.03 | | 3 | -0.04 | 0.03 |
| | 5 | -0.12 | 0.03 | | 5 | -0.09 | 0.03 |
| | 10 | -0.13 | 0.03 | | 10 | -0.13 | 0.02 |
| | 20 | -0.18 | 0.04 | | 20 | -0.14 | 0.02 |
| | 30 | -0.17 | 0.03 | | 30 | -0.15 | 0.03 |
| | 50 | -0.18 | 0.03 | | 50 | -0.20 | 0.02 |
| | 100 | -0.23 | 0.03 | | 100 | -0.16 | 0.02 |
| | 300 | -0.25 | 0.03 | | 300 | -0.18 | 0.03 |
| | 500 | -0.26 | 0.04 | | 500 | -0.13 | 0.03 |
| | 1000 | -0.24 | 0.03 | | 1000 | -0.19 | 0.02 |

**Table S2.3**: The average differences in the donor-acceptor distances of HBs 8-12 following the P1 Gln to Ala substitution in s01 and s05. Positive and negative differences with magnitudes > 0.1 Å are in green and red, respectively.

| **HB8-12** | s01: P1 Q->A | | | | s05: P1 Q->A | | |
|---|---|---|---|---|---|---|---|
| **HB8** | t (ps) | Av Diff (Å) | SEM (Å) | | t (ps) | Av Diff (Å) | SEM (Å) |
| | 0.05 | 0.00 | 0.00 | | 0.05 | 0.00 | 0.00 |
| | 0.1 | 0.02 | 0.00 | | 0.1 | 0.03 | 0.00 |
| | 0.25 | -0.01 | 0.01 | | 0.25 | -0.02 | 0.01 |
| | 0.5 | -0.02 | 0.02 | | 0.5 | 0.01 | 0.01 |
| | 1 | -0.02 | 0.02 | | 1 | -0.04 | 0.02 |
| | 2 | -0.02 | 0.03 | | 2 | -0.03 | 0.02 |
| | 3 | -0.01 | 0.03 | | 3 | -0.03 | 0.02 |
| | 5 | -0.04 | 0.03 | | 5 | -0.01 | 0.02 |
| | 10 | -0.04 | 0.03 | | 10 | -0.03 | 0.02 |
| | 20 | -0.05 | 0.03 | | 20 | -0.03 | 0.03 |
| | 30 | -0.04 | 0.03 | | 30 | 0.01 | 0.03 |
| | 50 | -0.08 | 0.03 | | 50 | -0.02 | 0.02 |
| | 100 | -0.08 | 0.04 | | 100 | 0.00 | 0.03 |
| | 300 | -0.04 | 0.03 | | 300 | -0.03 | 0.03 |
| | 500 | -0.10 | 0.03 | | 500 | 0.00 | 0.03 |
| | 1000 | -0.04 | 0.03 | | 1000 | 0.00 | 0.03 |
| **HB9** | t (ps) | Av Diff (Å) | SEM (Å) | | t (ps) | Av Diff (Å) | SEM (Å) |
| | 0.05 | 0.01 | 0.00 | | 0.05 | 0.00 | 0.00 |
| | 0.1 | -0.02 | 0.00 | | 0.1 | -0.03 | 0.00 |
| | 0.25 | 0.02 | 0.01 | | 0.25 | 0.01 | 0.01 |
| | 0.5 | 0.02 | 0.01 | | 0.5 | 0.02 | 0.01 |
| | 1 | 0.02 | 0.02 | | 1 | 0.01 | 0.02 |
| | 2 | 0.03 | 0.02 | | 2 | 0.00 | 0.02 |
| | 3 | 0.01 | 0.02 | | 3 | 0.03 | 0.02 |
| | 5 | 0.00 | 0.02 | | 5 | 0.00 | 0.02 |
| | 10 | 0.04 | 0.02 | | 10 | 0.03 | 0.02 |
| | 20 | 0.03 | 0.02 | | 20 | 0.01 | 0.02 |
| | 30 | 0.00 | 0.02 | | 30 | -0.01 | 0.02 |
| | 50 | 0.02 | 0.02 | | 50 | 0.03 | 0.02 |
| | 100 | 0.05 | 0.02 | | 100 | -0.01 | 0.02 |
| | 300 | 0.02 | 0.02 | | 300 | 0.00 | 0.02 |
| | 500 | 0.03 | 0.02 | | 500 | 0.03 | 0.02 |
| | 1000 | 0.03 | 0.02 | | 1000 | 0.00 | 0.02 |
| **HB10** | t (ps) | Av Diff (Å) | SEM (Å) | | t (ps) | Av Diff (Å) | SEM (Å) |
| | 0.05 | 0.00 | 0.00 | | 0.05 | 0.00 | 0.00 |
| | 0.1 | 0.00 | 0.00 | | 0.1 | 0.00 | 0.00 |
| | 0.25 | 0.04 | 0.00 | | 0.25 | 0.03 | 0.00 |
| | 0.5 | 0.03 | 0.01 | | 0.5 | 0.01 | 0.01 |
| | 1 | 0.03 | 0.01 | | 1 | 0.02 | 0.01 |
| | 2 | 0.02 | 0.02 | | 2 | 0.01 | 0.02 |
| | 3 | 0.01 | 0.02 | | 3 | 0.00 | 0.02 |
| | 5 | -0.01 | 0.02 | | 5 | 0.02 | 0.02 |
| | 10 | 0.00 | 0.02 | | 10 | 0.02 | 0.02 |
| | 20 | 0.01 | 0.02 | | 20 | -0.01 | 0.02 |
| | 30 | 0.03 | 0.02 | | 30 | 0.01 | 0.02 |
| | 50 | 0.08 | 0.02 | | 50 | 0.02 | 0.02 |
| | 100 | 0.06 | 0.02 | | 100 | 0.04 | 0.02 |
| | 300 | 0.07 | 0.02 | | 300 | 0.03 | 0.02 |
| | 500 | -0.01 | 0.02 | | 500 | 0.05 | 0.02 |
| | 1000 | 0.04 | 0.02 | | 1000 | 0.04 | 0.02 |
| **HB11** | t (ps) | Av Diff (Å) | SEM (Å) | | t (ps) | Av Diff (Å) | SEM (Å) |
| | 0.05 | 0.00 | 0.00 | | 0.05 | 0.00 | 0.00 |
| | 0.1 | 0.00 | 0.00 | | 0.1 | 0.00 | 0.00 |
| | 0.25 | 0.01 | 0.00 | | 0.25 | 0.02 | 0.00 |
| | 0.5 | 0.00 | 0.01 | | 0.5 | 0.00 | 0.00 |
| | 1 | -0.02 | 0.01 | | 1 | 0.03 | 0.01 |
| | 2 | -0.01 | 0.02 | | 2 | 0.00 | 0.02 |
| | 3 | 0.02 | 0.02 | | 3 | -0.02 | 0.02 |
| | 5 | -0.01 | 0.02 | | 5 | 0.01 | 0.02 |
| | 10 | 0.01 | 0.02 | | 10 | 0.01 | 0.02 |
| | 20 | 0.01 | 0.02 | | 20 | -0.01 | 0.02 |
| | 30 | 0.01 | 0.02 | | 30 | 0.00 | 0.02 |
| | 50 | 0.02 | 0.02 | | 50 | 0.00 | 0.02 |
| | 100 | 0.01 | 0.02 | | 100 | 0.04 | 0.02 |
| | 300 | -0.03 | 0.02 | | 300 | 0.00 | 0.02 |
| | 500 | -0.02 | 0.02 | | 500 | 0.03 | 0.02 |
| | 1000 | 0.02 | 0.02 | | 1000 | 0.02 | 0.02 |
| **HB12** | t (ps) | Av Diff (Å) | SEM (Å) | | t (ps) | Av Diff (Å) | SEM (Å) |
| | 0.05 | 0.00 | 0.00 | | 0.05 | 0.00 | 0.00 |
| | 0.1 | 0.00 | 0.00 | | 0.1 | 0.00 | 0.00 |
| | 0.25 | 0.00 | 0.00 | | 0.25 | 0.00 | 0.00 |
| | 0.5 | 0.00 | 0.00 | | 0.5 | 0.00 | 0.00 |
| | 1 | -0.01 | 0.02 | | 1 | 0.00 | 0.02 |
| | 2 | -0.01 | 0.03 | | 2 | -0.02 | 0.02 |
| | 3 | 0.03 | 0.03 | | 3 | 0.03 | 0.02 |
| | 5 | -0.01 | 0.03 | | 5 | -0.01 | 0.02 |
| | 10 | -0.04 | 0.03 | | 10 | -0.01 | 0.03 |
| | 20 | -0.01 | 0.03 | | 20 | -0.02 | 0.02 |
| | 30 | -0.03 | 0.03 | | 30 | -0.02 | 0.03 |
| | 50 | 0.01 | 0.03 | | 50 | -0.04 | 0.03 |
| | 100 | -0.02 | 0.03 | | 100 | 0.07 | 0.03 |
| | 300 | 0.01 | 0.05 | | 300 | 0.01 | 0.03 |
| | 500 | 0.01 | 0.04 | | 500 | -0.01 | 0.03 |
| | 1000 | -0.09 | 0.05 | | 1000 | 0.01 | 0.04 |

**Table S2.4**: The ten most perturbed Cα atoms in the $M^{pro}$-peptide complex in terms of magnitude of the average displacement vector *v*, at selected time points following the P1 Gln to Ala substitution in s01 and s05. Only displacements ≥ 0.01 Å are shown.

| s01 t(ps) | CA Rank | Atom | Chain | v (Å) | SEM (Å) |
|---|---|---|---|---|---|
| **0.05** | 1 | 6ALA_CA | C | 0.03 | 0.00 |
| | 2 | 140PHE_CA | A | 0.01 | 0.00 |
| | 3 | 144SER_CA | A | 0.01 | 0.00 |
| | 4 | 163HIE_CA | A | 0.01 | 0.00 |
| | 5 | 165MET_CA | A | 0.01 | 0.00 |
| | 6 | 166GLU_CA | A | 0.01 | 0.00 |
| | 7 | 1SER_CA | B | 0.01 | 0.00 |
| | 8 | 2GLY_CA | B | 0.01 | 0.00 |
| | 9 | 5LEU_CA | C | 0.01 | 0.00 |
| | 10 | 7SER_CA | C | 0.01 | 0.00 |
| **0.5** | 1 | 6ALA_CA | C | 0.17 | 0.02 |
| | 2 | 140PHE_CA | A | 0.12 | 0.01 |
| | 3 | 163HIE_CA | A | 0.12 | 0.01 |
| | 4 | 141LEU_CA | A | 0.11 | 0.01 |
| | 5 | 162MET_CA | A | 0.11 | 0.01 |
| | 6 | 7SER_CA | C | 0.08 | 0.01 |
| | 7 | 142ASN_CA | A | 0.06 | 0.01 |
| | 8 | 8GLY_CA | C | 0.06 | 0.01 |
| | 9 | 139SER_CA | A | 0.05 | 0.01 |
| | 10 | 161TYR_CA | A | 0.05 | 0.00 |
| **5** | 1 | 6ALA_CA | C | 0.31 | 0.03 |
| | 2 | 73VAL_CA | A | 0.20 | 0.05 |
| | 3 | 72ASN_CA | A | 0.18 | 0.06 |
| | 4 | 170GLY_CA | A | 0.15 | 0.04 |
| | 5 | 196THR_CA | B | 0.15 | 0.05 |
| | 6 | 24THR_CA | A | 0.13 | 0.04 |
| | 7 | 74GLN_CA | A | 0.13 | 0.04 |
| | 8 | 244GLN_CA | A | 0.13 | 0.04 |
| | 9 | 245ASP_CA | A | 0.13 | 0.04 |
| | 10 | 24THR_CA | B | 0.13 | 0.04 |
| **50** | 1 | 6ALA_CA | C | 0.43 | 0.03 |
| | 2 | 72ASN_CA | A | 0.33 | 0.08 |
| | 3 | 47GLU_CA | A | 0.30 | 0.07 |
| | 4 | 11LYS_CA | C | 0.30 | 0.09 |
| | 5 | 8GLY_CA | C | 0.29 | 0.04 |
| | 6 | 23GLY_CA | A | 0.28 | 0.06 |
| | 7 | 72ASN_CA | B | 0.25 | 0.08 |
| | 8 | 278GLY_CA | B | 0.24 | 0.07 |
| | 9 | 24THR_CA | A | 0.23 | 0.06 |
| | 10 | 48ASP_CA | A | 0.23 | 0.05 |
| **500** | 1 | 6ALA_CA | C | 0.38 | 0.04 |
| | 2 | 11LYS_CA | C | 0.30 | 0.11 |
| | 3 | 47GLU_CA | A | 0.28 | 0.09 |
| | 4 | 215GLY_CA | A | 0.27 | 0.10 |
| | 5 | 1THR_CA | C | 0.27 | 0.13 |
| | 6 | 53ASN_CA | A | 0.25 | 0.08 |
| | 7 | 277ASN_CA | A | 0.25 | 0.10 |
| | 8 | 54TYR_CA | A | 0.24 | 0.06 |
| | 9 | 2SER_CA | C | 0.23 | 0.08 |
| | 10 | 9PHE_CA | C | 0.22 | 0.06 |
| **1000** | 1 | 6ALA_CA | C | 0.47 | 0.04 |
| | 2 | 9PHE_CA | C | 0.37 | 0.06 |
| | 3 | 10ARG_CA | C | 0.37 | 0.08 |
| | 4 | 11LYS_CA | C | 0.37 | 0.13 |
| | 5 | 71GLY_CA | A | 0.35 | 0.08 |
| | 6 | 73VAL_CA | A | 0.34 | 0.09 |
| | 7 | 24THR_CA | A | 0.32 | 0.07 |
| | 8 | 56ASP_CA | A | 0.32 | 0.09 |
| | 9 | 59ILE_CA | A | 0.32 | 0.08 |
| | 10 | 277ASN_CA | B | 0.31 | 0.10 |

| s05 t(ps) | CA Rank | Atom | Chain | v (Å) | SEM (Å) |
|---|---|---|---|---|---|
| **0.05** | 1 | 6ALA_CA | C | 0.02 | 0.00 |
| | 2 | 144SER_CA | A | 0.01 | 0.00 |
| | 3 | 164HIE_CA | A | 0.01 | 0.00 |
| | 4 | 166GLU_CA | A | 0.01 | 0.00 |
| | 5 | 1SER_CA | B | 0.01 | 0.00 |
| | 6 | 5LEU_CA | C | 0.01 | 0.00 |
| | 7 | | | | |
| | 8 | | | | |
| | 9 | | | | |
| | 10 | | | | |
| **0.5** | 1 | 6ALA_CA | C | 0.20 | 0.01 |
| | 2 | 163HIE_CA | A | 0.16 | 0.01 |
| | 3 | 162MET_CA | A | 0.15 | 0.01 |
| | 4 | 140PHE_CA | A | 0.08 | 0.01 |
| | 5 | 141LEU_CA | A | 0.08 | 0.01 |
| | 6 | 172HIE_CA | A | 0.08 | 0.00 |
| | 7 | 161TYR_CA | A | 0.07 | 0.00 |
| | 8 | 165MET_CA | A | 0.07 | 0.01 |
| | 9 | 166GLU_CA | A | 0.07 | 0.01 |
| | 10 | 164HIE_CA | A | 0.06 | 0.01 |
| **5** | 1 | 6ALA_CA | C | 0.30 | 0.03 |
| | 2 | 222ARG_CA | B | 0.21 | 0.05 |
| | 3 | 223PHE_CA | B | 0.20 | 0.04 |
| | 4 | 274ASN_CA | B | 0.19 | 0.05 |
| | 5 | 72ASN_CA | A | 0.18 | 0.06 |
| | 6 | 275GLY_CA | B | 0.16 | 0.04 |
| | 7 | 11SER_CA | C | 0.16 | 0.08 |
| | 8 | 224THR_CA | B | 0.15 | 0.04 |
| | 9 | 5LEU_CA | C | 0.15 | 0.02 |
| | 10 | 163HIE_CA | A | 0.14 | 0.02 |
| **50** | 1 | 6ALA_CA | C | 0.40 | 0.03 |
| | 2 | 72ASN_CA | B | 0.24 | 0.08 |
| | 3 | 47GLU_CA | A | 0.21 | 0.06 |
| | 4 | 226THR_CA | A | 0.19 | 0.06 |
| | 5 | 73VAL_CA | B | 0.19 | 0.06 |
| | 6 | 46SER_CA | A | 0.18 | 0.06 |
| | 7 | 225THR_CA | A | 0.17 | 0.06 |
| | 8 | 278GLY_CA | A | 0.17 | 0.09 |
| | 9 | 23GLY_CA | B | 0.17 | 0.07 |
| | 10 | 93THR_CA | A | 0.16 | 0.05 |
| **500** | 1 | 6ALA_CA | C | 0.42 | 0.03 |
| | 2 | 1SER_CA | C | 0.38 | 0.09 |
| | 3 | 191ALA_CA | B | 0.33 | 0.09 |
| | 4 | 190THR_CA | B | 0.29 | 0.08 |
| | 5 | 274ASN_CA | B | 0.23 | 0.09 |
| | 6 | 275GLY_CA | B | 0.22 | 0.08 |
| | 7 | 52PRO_CA | A | 0.21 | 0.08 |
| | 8 | 141LEU_CA | A | 0.19 | 0.04 |
| | 9 | 215GLY_CA | A | 0.19 | 0.10 |
| | 10 | 51ASN_CA | A | 0.18 | 0.07 |
| **1000** | 1 | 6ALA_CA | C | 0.48 | 0.03 |
| | 2 | 72ASN_CA | A | 0.41 | 0.11 |
| | 3 | 1SER_CA | C | 0.39 | 0.10 |
| | 4 | 47GLU_CA | A | 0.36 | 0.09 |
| | 5 | 71GLY_CA | A | 0.34 | 0.09 |
| | 6 | 48ASP_CA | A | 0.32 | 0.06 |
| | 7 | 2ALA_CA | C | 0.31 | 0.06 |
| | 8 | 45THR_CA | A | 0.30 | 0.06 |
| | 9 | 46SER_CA | A | 0.30 | 0.08 |
| | 10 | 52PRO_CA | A | 0.29 | 0.09 |

**Table S2.5**: The ten most perturbed non-hydrogen atoms in the M$^{pro}$-peptide complex in terms of magnitude of the average displacement vector *v*, at selected time points following the P1 Gln to Ala substitution in s01 and s05.

| *s01 t(ps)* | *Non-H Rank* | *Atom* | *Chain* | *v (Å)* | *SEM (Å)* |
|---|---|---|---|---|---|
| **0.05** | 1 | 6ALA_CB | C | 0.09 | 0.01 |
| | 2 | 140PHE_O | A | 0.05 | 0.00 |
| | 3 | 163HIE_CE1 | A | 0.04 | 0.00 |
| | 4 | 166GLU_OE1 | A | 0.04 | 0.00 |
| | 5 | 163HIE_NE2 | A | 0.04 | 0.00 |
| | 6 | 144SER_OG | A | 0.03 | 0.00 |
| | 7 | 163HIE_ND1 | A | 0.03 | 0.00 |
| | 8 | 166GLU_OE2 | A | 0.03 | 0.00 |
| | 9 | 166GLU_CB | A | 0.03 | 0.00 |
| | 10 | 6ALA_CA | C | 0.03 | 0.00 |
| **0.5** | 1 | 163HIE_CE1 | A | 0.46 | 0.02 |
| | 2 | 163HIE_NE2 | A | 0.39 | 0.02 |
| | 3 | 6ALA_CB | C | 0.36 | 0.03 |
| | 4 | 163HIE_ND1 | A | 0.35 | 0.02 |
| | 5 | 140PHE_O | A | 0.23 | 0.02 |
| | 6 | 163HIE_CD2 | A | 0.20 | 0.01 |
| | 7 | 6ALA_CA | C | 0.17 | 0.02 |
| | 8 | 140PHE_C | A | 0.17 | 0.01 |
| | 9 | 163HIE_N | A | 0.15 | 0.01 |
| | 10 | 162MET_SD | A | 0.15 | 0.01 |
| **5** | 1 | 6ALA_CB | C | 0.59 | 0.04 |
| | 2 | 6ALA_CA | C | 0.32 | 0.03 |
| | 3 | 73VAL_CG1 | A | 0.29 | 0.07 |
| | 4 | 72ASN_OD1 | A | 0.28 | 0.10 |
| | 5 | 142ASN_OD1 | A | 0.28 | 0.06 |
| | 6 | 196THR_CG2 | B | 0.27 | 0.08 |
| | 7 | 142ASN_ND2 | A | 0.27 | 0.06 |
| | 8 | 47GLU_OE2 | B | 0.26 | 0.09 |
| | 9 | 244GLN_OE1 | A | 0.26 | 0.08 |
| | 10 | 163HIE_CE1 | A | 0.26 | 0.03 |
| **50** | 1 | 6ALA_CB | C | 0.80 | 0.04 |
| | 2 | 72ASN_OD1 | A | 0.65 | 0.15 |
| | 3 | 47GLU_OE2 | A | 0.65 | 0.16 |
| | 4 | 72ASN_ND2 | A | 0.55 | 0.14 |
| | 5 | 145CYS_SG | A | 0.54 | 0.06 |
| | 6 | 72ASN_CG | A | 0.53 | 0.12 |
| | 7 | 47GLU_CD | A | 0.48 | 0.13 |
| | 8 | 72ASN_ND2 | B | 0.48 | 0.13 |
| | 9 | 142ASN_ND2 | A | 0.46 | 0.08 |
| | 10 | 47GLU_OE1 | A | 0.45 | 0.16 |
| **500** | 1 | 6ALA_CB | C | 0.75 | 0.05 |
| | 2 | 145CYS_SG | A | 0.70 | 0.07 |
| | 3 | 11LYS_NZ | C | 0.66 | 0.29 |
| | 4 | 142ASN_OD1 | B | 0.53 | 0.15 |
| | 5 | 11LYS_CE | C | 0.52 | 0.24 |
| | 6 | 306GLN_NE2 | A | 0.52 | 0.20 |
| | 7 | 11LYS_CD | C | 0.49 | 0.21 |
| | 8 | 236LYS_NZ | B | 0.49 | 0.25 |
| | 9 | 47GLU_OE2 | A | 0.48 | 0.23 |
| | 10 | 277ASN_OD1 | A | 0.47 | 0.20 |
| **1000** | 1 | 6ALA_CB | C | 0.87 | 0.04 |
| | 2 | 145CYS_SG | A | 0.76 | 0.07 |
| | 3 | 10ARG_NH2 | C | 0.55 | 0.23 |
| | 4 | 244GLN_OE1 | A | 0.54 | 0.23 |
| | 5 | 244GLN_NE2 | A | 0.54 | 0.25 |
| | 6 | 11LYS_OC2 | C | 0.49 | 0.16 |
| | 7 | 279ARG_NH1 | A | 0.48 | 0.21 |
| | 8 | 235MET_SD | B | 0.48 | 0.18 |
| | 9 | 6ALA_CA | C | 0.47 | 0.04 |
| | 10 | 10ARG_CZ | C | 0.47 | 0.19 |

| *s05 t(ps)* | *Non-H Rank* | *Atom* | *Chain* | *v (Å)* | *SEM (Å)* |
|---|---|---|---|---|---|
| **0.05** | 1 | 6ALA_CB | C | 0.12 | 0.01 |
| | 2 | 163HIE_CE1 | A | 0.04 | 0.00 |
| | 3 | 163HIE_NE2 | A | 0.04 | 0.00 |
| | 4 | 140PHE_O | A | 0.04 | 0.00 |
| | 5 | 163HIE_ND1 | A | 0.04 | 0.00 |
| | 6 | 166GLU_OE1 | A | 0.03 | 0.00 |
| | 7 | 144SER_OG | A | 0.03 | 0.00 |
| | 8 | 166GLU_CB | A | 0.03 | 0.00 |
| | 9 | 165MET_O | A | 0.03 | 0.00 |
| | 10 | 172HIE_ND1 | A | 0.02 | 0.00 |
| **0.5** | 1 | 163HIE_CE1 | A | 0.50 | 0.02 |
| | 2 | 6ALA_CB | C | 0.45 | 0.02 |
| | 3 | 163HIE_NE2 | A | 0.41 | 0.02 |
| | 4 | 163HIE_ND1 | A | 0.40 | 0.02 |
| | 5 | 163HIE_N | A | 0.21 | 0.01 |
| | 6 | 162MET_SD | A | 0.20 | 0.01 |
| | 7 | 6ALA_CA | C | 0.20 | 0.01 |
| | 8 | 163HIE_CD2 | A | 0.19 | 0.01 |
| | 9 | 140PHE_O | A | 0.19 | 0.02 |
| | 10 | 163HIE_CB | A | 0.19 | 0.01 |
| **5** | 1 | 6ALA_CB | C | 0.60 | 0.03 |
| | 2 | 163HIE_CE1 | A | 0.39 | 0.03 |
| | 3 | 163HIE_ND1 | A | 0.33 | 0.03 |
| | 4 | 163HIE_NE2 | A | 0.31 | 0.03 |
| | 5 | 6ALA_CA | C | 0.31 | 0.03 |
| | 6 | 142ASN_ND2 | A | 0.29 | 0.06 |
| | 7 | 72ASN_OD1 | A | 0.29 | 0.10 |
| | 8 | 72ASN_ND2 | A | 0.28 | 0.10 |
| | 9 | 72ASN_CG | A | 0.28 | 0.09 |
| | 10 | 72ASN_CB | A | 0.26 | 0.08 |
| **50** | 1 | 6ALA_CB | C | 0.79 | 0.04 |
| | 2 | 236LYS_NZ | B | 0.48 | 0.18 |
| | 3 | 72ASN_OD1 | B | 0.44 | 0.14 |
| | 4 | 74GLN_NE2 | B | 0.43 | 0.12 |
| | 5 | 236LYS_NZ | A | 0.42 | 0.18 |
| | 6 | 236LYS_CE | B | 0.41 | 0.15 |
| | 7 | 76ARG_NH1 | A | 0.40 | 0.15 |
| | 8 | 6ALA_CA | C | 0.40 | 0.03 |
| | 9 | 47GLU_OE2 | A | 0.39 | 0.17 |
| | 10 | 76ARG_NH2 | A | 0.38 | 0.15 |
| **500** | 1 | 6ALA_CB | C | 0.80 | 0.03 |
| | 2 | 236LYS_NZ | B | 0.75 | 0.26 |
| | 3 | 236LYS_CE | B | 0.67 | 0.22 |
| | 4 | 222ARG_NH1 | B | 0.67 | 0.22 |
| | 5 | 222ARG_NH2 | B | 0.59 | 0.28 |
| | 6 | 222ARG_CZ | B | 0.54 | 0.22 |
| | 7 | 60ARG_NH1 | B | 0.51 | 0.18 |
| | 8 | 47GLU_OE1 | A | 0.50 | 0.22 |
| | 9 | 1SER_OG | C | 0.49 | 0.14 |
| | 10 | 107GLN_OE1 | B | 0.46 | 0.15 |
| **1000** | 1 | 67LEU_CD1 | A | 0.86 | 0.13 |
| | 2 | 6ALA_CB | C | 0.86 | 0.04 |
| | 3 | 222ARG_NH1 | A | 0.82 | 0.29 |
| | 4 | 222ARG_NH2 | A | 0.78 | 0.31 |
| | 5 | 72ASN_OD1 | A | 0.73 | 0.21 |
| | 6 | 107GLN_OE1 | B | 0.72 | 0.15 |
| | 7 | 47GLU_OE2 | A | 0.70 | 0.19 |
| | 8 | 222ARG_CZ | A | 0.69 | 0.26 |
| | 9 | 47GLU_CG | A | 0.65 | 0.14 |
| | 10 | 142ASN_ND2 | A | 0.65 | 0.10 |

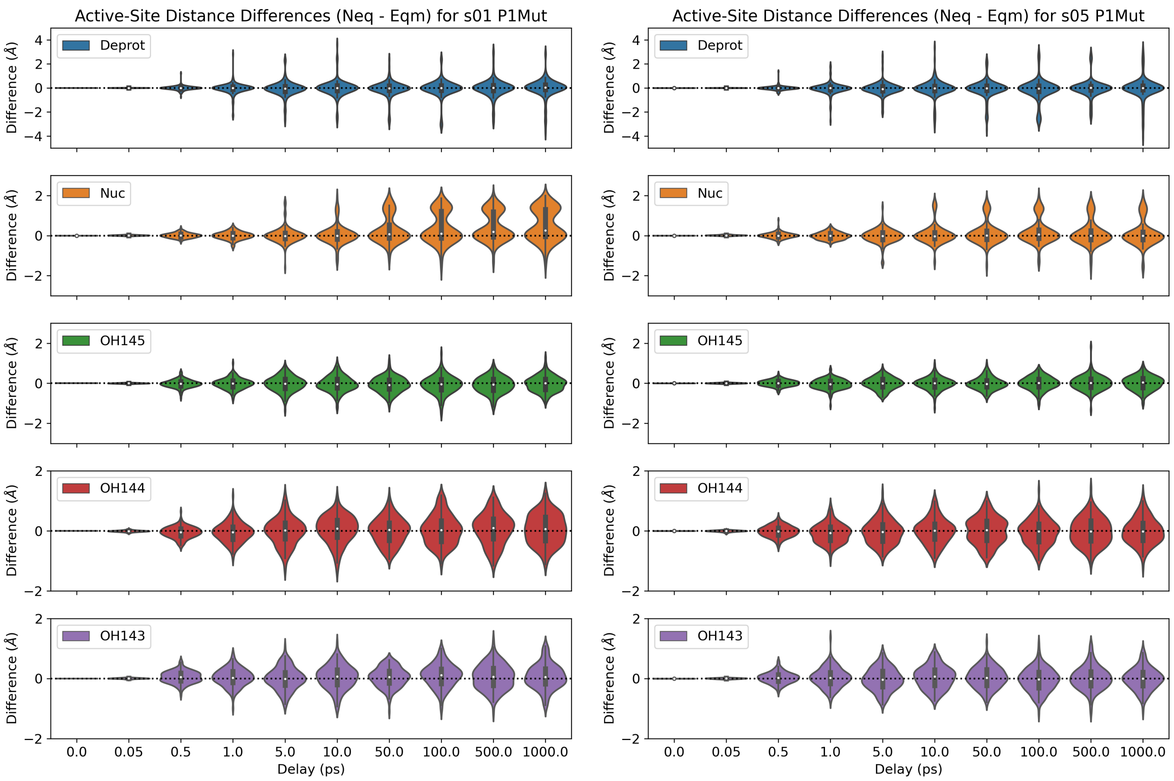

**Figure S2.1**: Distributions of the distance differences in the five active-site distances ($\Delta d = d_{\text{neq}} - d_{\text{eqm}}$) following the P1 Gln to Ala substitution in s01 and s05.

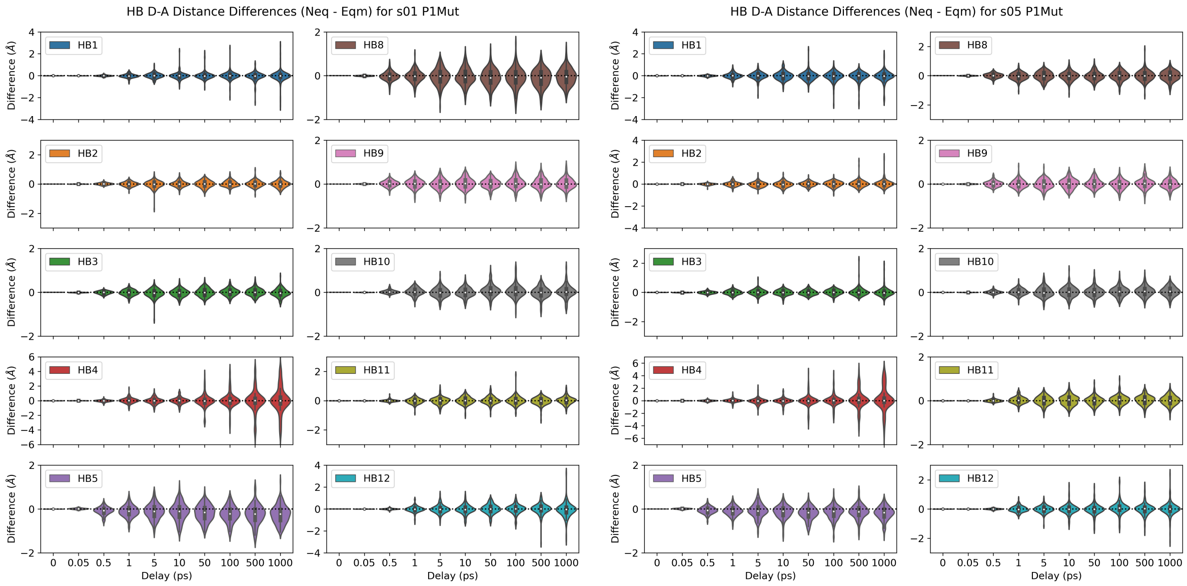

**Figure S2.2**: Distributions of the distance differences in the ten donor-acceptor distances of HBs 1-5 and 8-12 following the P1 Gln to Ala substitution in s01 and s05.

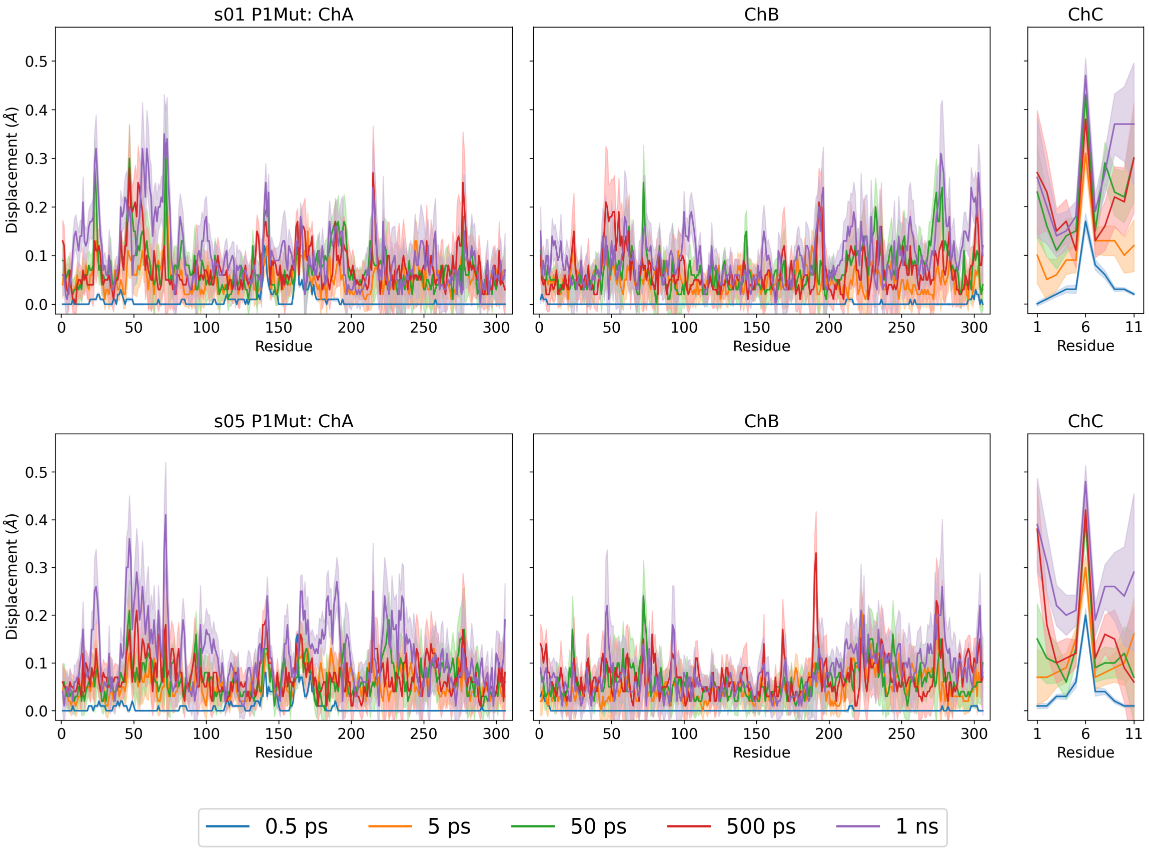

**Figure S2.3**: Magnitudes of the average Cα displacement vectors, with ± standard error of the mean (SEM) shown as shaded regions, following the P1 Gln to Ala substitution in s01 and s05.

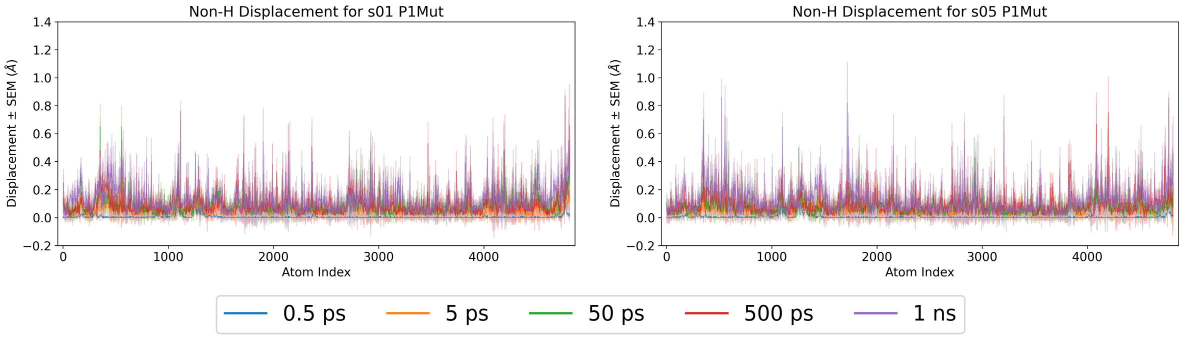

**Figure S2.4**: Magnitudes of the average non-hydrogen atom displacement vectors in the $M^{pro}$-peptide complex, with ± SEM shown as shaded regions, following the P1 Gln to Ala substitution in s01 and s05.

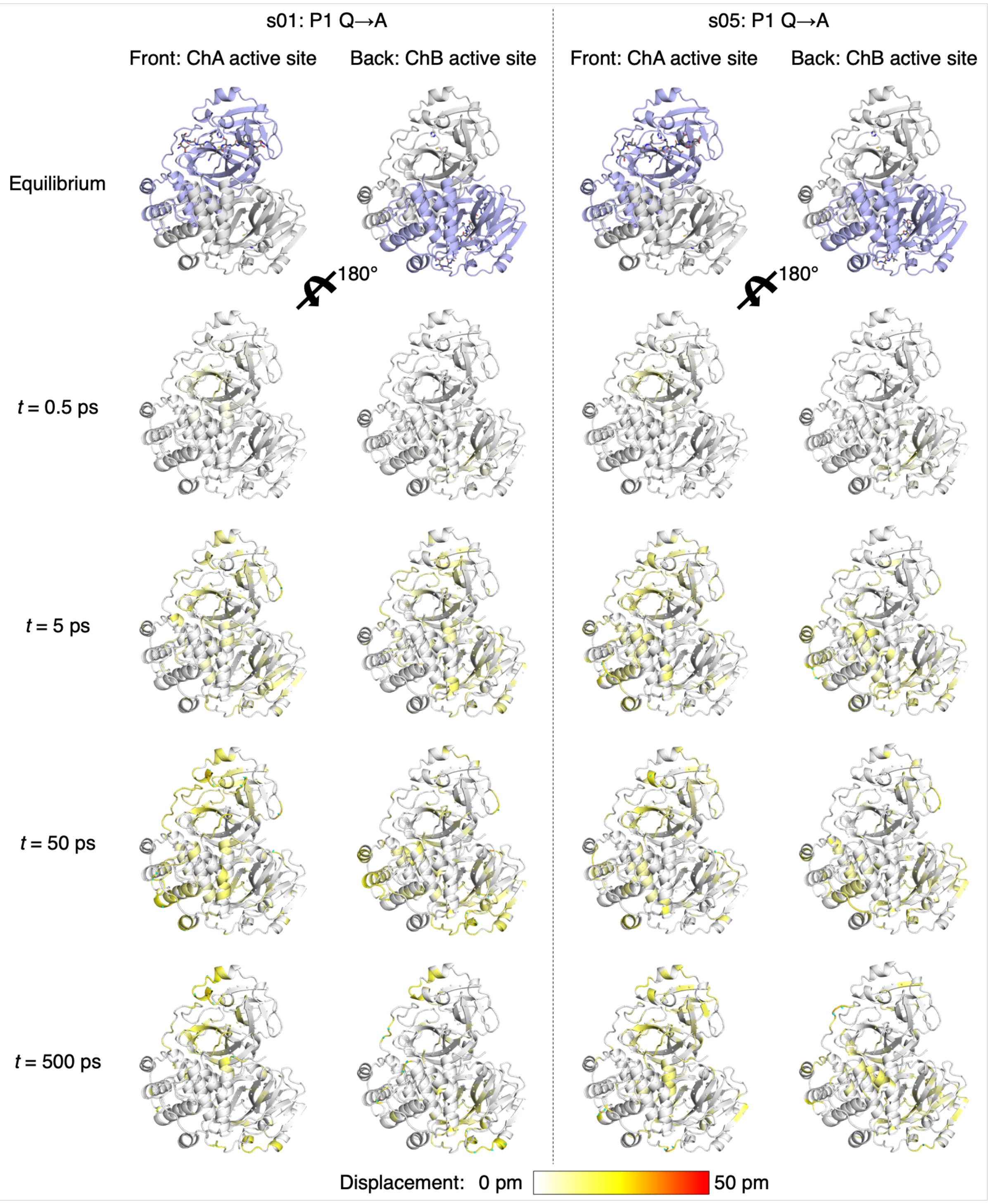

**Figure S2.5**: Views of the M$^{pro}$ response to the P1 Gln to Ala substitution in s01 and s05 from averaging Cα displacement vectors, shown using the structure prior to MD simulations. Displacement magnitudes are represented on a white-yellow-red scale.[7] Significant vectors with length ≥20 pm are displayed as cyan arrows with a scale-up factor of 5.[8]

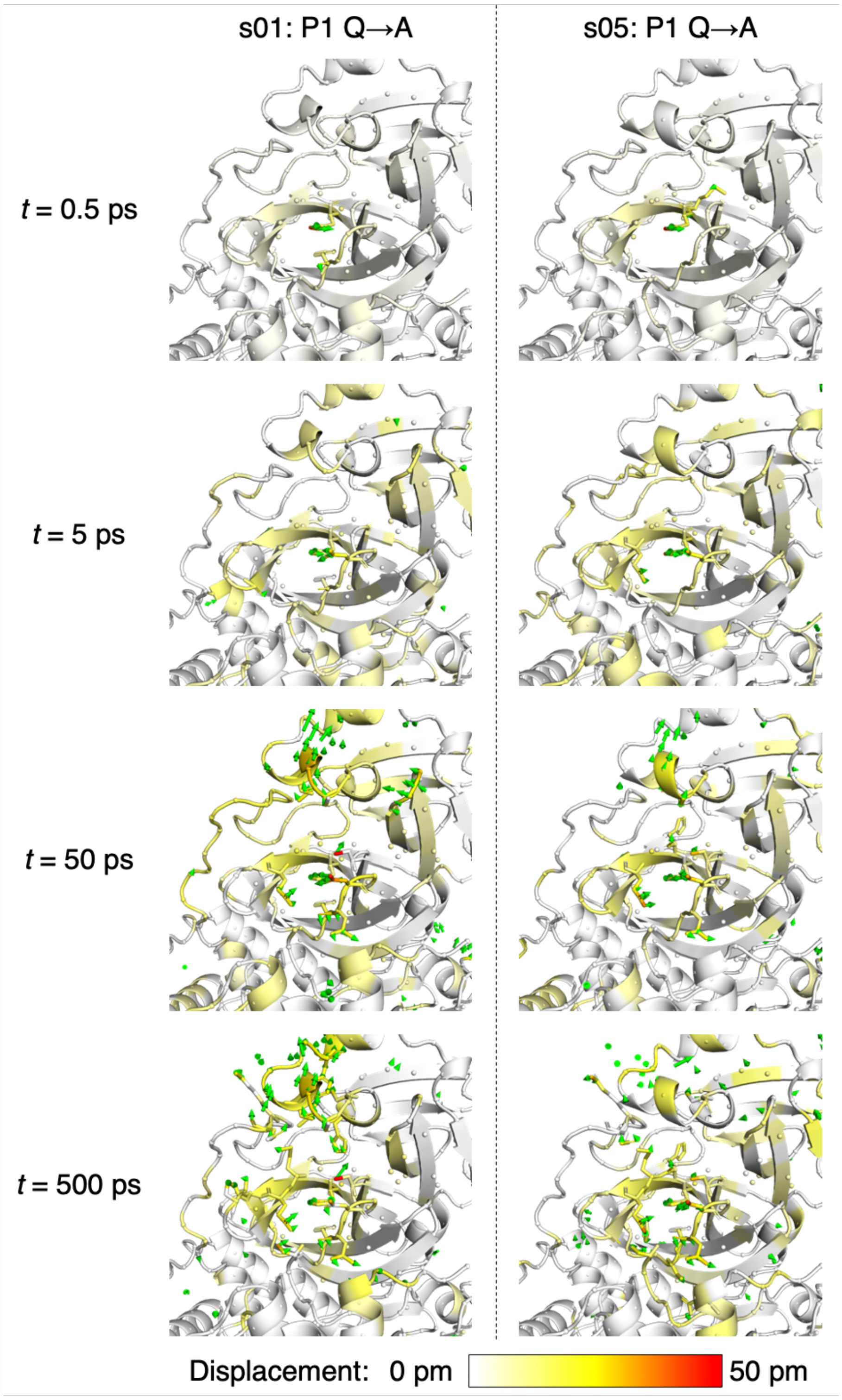

**Figure S2.6**: Views of the M$^{pro}$ response to the P1 Gln to Ala substitution in s01 and s05 from averaging non-hydrogen atom displacement vectors, shown using the structure prior to MD simulations, with a focus on the S1 subsite. Displacement magnitudes are represented on a white-yellow-red scale.[7] Significant vectors with length ≥20 pm are displayed as green arrows with a scale-up factor of 5.[8] Residues that show such significant displacements and that are within 10 Å of the substrate P1 residue are shown as sticks.

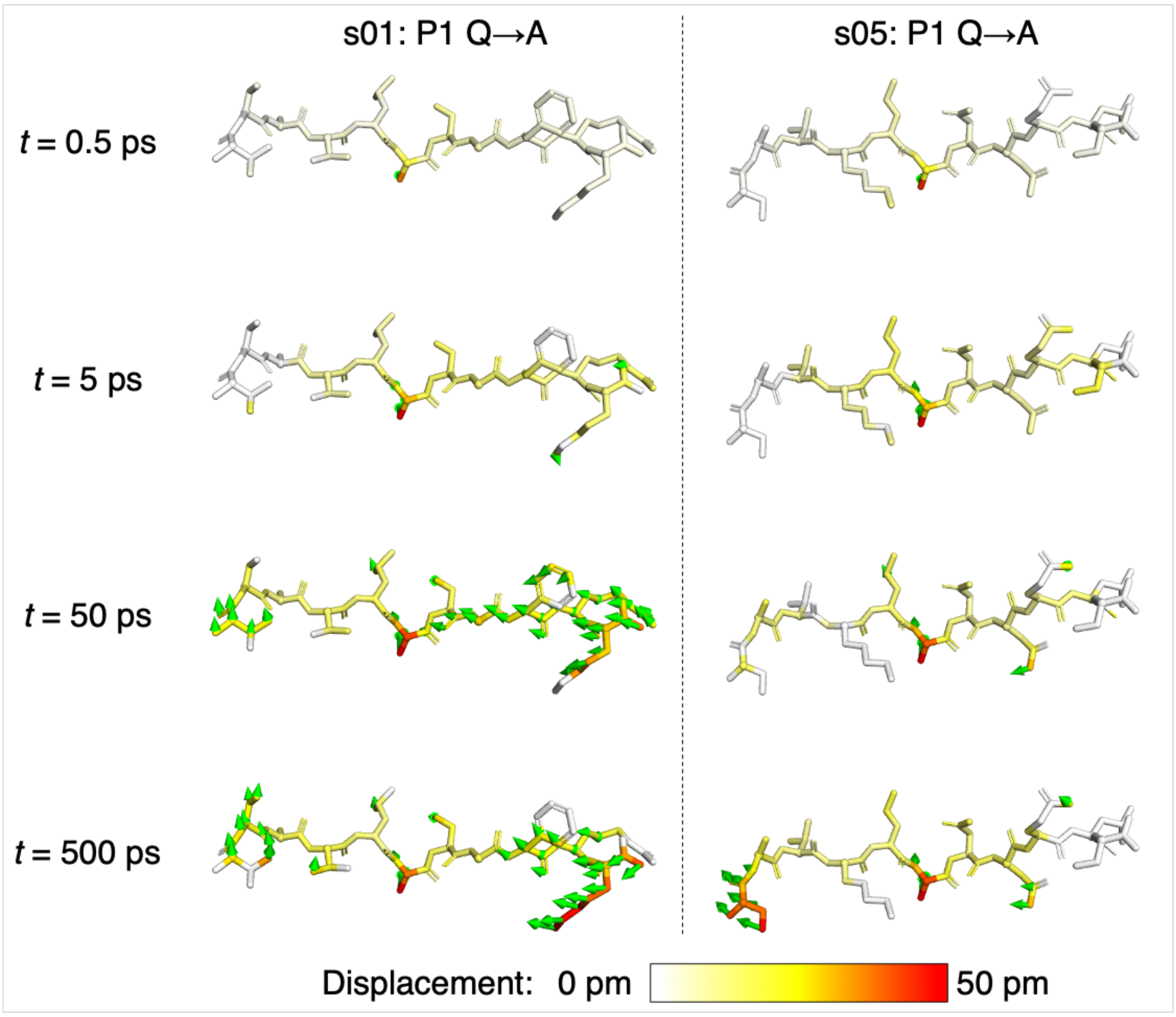

**Figure S2.7**: Views of the peptide response to the P1 Gln to Ala substitution in s01 and s05 from averaging non-hydrogen atom displacement vectors, shown using the structure prior to MD simulations. Displacement magnitudes are represented on a white-yellow-red scale.[7] Significant vectors with length ≥20 pm are displayed as green arrows with a scale-up factor of 5.[8]

### Section S3: Effect of P2 substitution

**Table S3.1**: The average differences in the five active-site distances ($\Delta d = d_{neq} - d_{eqm}$) following the shown P2 substitutions in s01, s05, s02, and p12. Positive and negative differences with magnitudes > 0.1 Å are in green and red, respectively.

| Active Site | s01: P2 L->A | | | s05: P2 L->A | | | s02: P2 F->A | | | p12: P2 W->A | | |
|---|---|---|---|---|---|---|---|---|---|---|---|---|
| **Deprot** | t (ps) | Av Diff (Å) | SEM (Å) | t (ps) | Av Diff (Å) | SEM (Å) | t (ps) | Av Diff (Å) | SEM (Å) | t (ps) | Av Diff (Å) | SEM (Å) |
| | 0.05 | 0.00 | 0.00 | 0.05 | 0.00 | 0.00 | 0.05 | 0.00 | 0.00 | 0.05 | 0.01 | 0.00 |
| | 0.1 | 0.00 | 0.00 | 0.1 | 0.01 | 0.00 | 0.1 | 0.00 | 0.00 | 0.1 | 0.01 | 0.00 |
| | 0.25 | 0.02 | 0.01 | 0.25 | 0.01 | 0.01 | 0.25 | 0.01 | 0.01 | 0.25 | 0.03 | 0.01 |
| | 0.5 | 0.02 | 0.01 | 0.5 | 0.00 | 0.01 | 0.5 | -0.01 | 0.01 | 0.5 | -0.01 | 0.02 |
| | 1 | -0.04 | 0.03 | 1 | -0.02 | 0.03 | 1 | -0.02 | 0.04 | 1 | -0.01 | 0.02 |
| | 2 | -0.03 | 0.04 | 2 | -0.06 | 0.05 | 2 | 0.00 | 0.04 | 2 | -0.06 | 0.03 |
| | 3 | 0.06 | 0.04 | 3 | -0.07 | 0.06 | 3 | 0.01 | 0.05 | 3 | -0.04 | 0.03 |
| | 5 | -0.03 | 0.06 | 5 | -0.11 | 0.05 | 5 | 0.00 | 0.05 | 5 | -0.03 | 0.03 |
| | 10 | 0.06 | 0.06 | 10 | -0.03 | 0.06 | 10 | -0.04 | 0.04 | 10 | -0.07 | 0.04 |
| | 20 | 0.06 | 0.05 | 20 | -0.04 | 0.06 | 20 | -0.04 | 0.04 | 20 | -0.05 | 0.03 |
| | 30 | 0.02 | 0.07 | 30 | 0.00 | 0.07 | 30 | 0.01 | 0.05 | 30 | -0.03 | 0.04 |
| | 50 | 0.06 | 0.07 | 50 | -0.06 | 0.07 | 50 | 0.07 | 0.05 | 50 | -0.02 | 0.04 |
| | 100 | 0.07 | 0.07 | 100 | 0.04 | 0.08 | 100 | 0.02 | 0.05 | 100 | -0.05 | 0.03 |
| | 300 | -0.04 | 0.08 | 300 | 0.03 | 0.07 | 300 | -0.08 | 0.04 | 300 | 0.00 | 0.04 |
| | 500 | 0.03 | 0.06 | 500 | 0.10 | 0.06 | 500 | 0.12 | 0.06 | 500 | -0.02 | 0.05 |
| | 1000 | 0.01 | 0.08 | 1000 | 0.05 | 0.06 | 1000 | 0.15 | 0.06 | 1000 | -0.02 | 0.04 |
| **Nuc** | t (ps) | Av Diff (Å) | SEM (Å) | t (ps) | Av Diff (Å) | SEM (Å) | t (ps) | Av Diff (Å) | SEM (Å) | t (ps) | Av Diff (Å) | SEM (Å) |
| | 0.05 | 0.00 | 0.00 | 0.05 | 0.00 | 0.00 | 0.05 | 0.00 | 0.00 | 0.05 | 0.00 | 0.00 |
| | 0.1 | 0.00 | 0.00 | 0.1 | 0.00 | 0.00 | 0.1 | 0.00 | 0.00 | 0.1 | 0.00 | 0.00 |
| | 0.25 | 0.00 | 0.00 | 0.25 | 0.01 | 0.00 | 0.25 | 0.00 | 0.00 | 0.25 | 0.01 | 0.00 |
| | 0.5 | 0.00 | 0.01 | 0.5 | 0.01 | 0.01 | 0.5 | -0.02 | 0.01 | 0.5 | -0.01 | 0.01 |
| | 1 | 0.01 | 0.01 | 1 | 0.00 | 0.01 | 1 | -0.04 | 0.01 | 1 | -0.02 | 0.01 |
| | 2 | 0.00 | 0.02 | 2 | -0.02 | 0.02 | 2 | -0.03 | 0.02 | 2 | 0.00 | 0.02 |
| | 3 | -0.05 | 0.02 | 3 | 0.02 | 0.02 | 3 | -0.04 | 0.02 | 3 | -0.02 | 0.02 |
| | 5 | 0.00 | 0.03 | 5 | -0.04 | 0.02 | 5 | 0.00 | 0.03 | 5 | -0.05 | 0.02 |
| | 10 | -0.02 | 0.03 | 10 | 0.00 | 0.03 | 10 | -0.03 | 0.02 | 10 | -0.01 | 0.02 |
| | 20 | 0.03 | 0.03 | 20 | 0.00 | 0.03 | 20 | -0.04 | 0.02 | 20 | -0.02 | 0.03 |
| | 30 | -0.02 | 0.03 | 30 | 0.03 | 0.03 | 30 | -0.04 | 0.03 | 30 | -0.01 | 0.03 |
| | 50 | -0.03 | 0.04 | 50 | 0.02 | 0.04 | 50 | -0.05 | 0.03 | 50 | -0.03 | 0.03 |
| | 100 | -0.04 | 0.04 | 100 | 0.07 | 0.04 | 100 | -0.04 | 0.04 | 100 | -0.05 | 0.04 |
| | 300 | 0.03 | 0.05 | 300 | 0.09 | 0.05 | 300 | -0.02 | 0.05 | 300 | -0.03 | 0.05 |
| | 500 | 0.12 | 0.05 | 500 | 0.05 | 0.05 | 500 | -0.04 | 0.06 | 500 | -0.05 | 0.06 |
| | 1000 | 0.15 | 0.05 | 1000 | 0.00 | 0.05 | 1000 | -0.12 | 0.06 | 1000 | -0.04 | 0.06 |
| **OH145** | t (ps) | Av Diff (Å) | SEM (Å) | t (ps) | Av Diff (Å) | SEM (Å) | t (ps) | Av Diff (Å) | SEM (Å) | t (ps) | Av Diff (Å) | SEM (Å) |
| | 0.05 | 0.00 | 0.00 | 0.05 | 0.00 | 0.00 | 0.05 | 0.00 | 0.00 | 0.05 | 0.00 | 0.00 |
| | 0.1 | 0.01 | 0.00 | 0.1 | 0.00 | 0.00 | 0.1 | 0.01 | 0.00 | 0.1 | 0.01 | 0.00 |
| | 0.25 | 0.01 | 0.01 | 0.25 | -0.01 | 0.00 | 0.25 | 0.04 | 0.01 | 0.25 | 0.02 | 0.01 |
| | 0.5 | 0.02 | 0.01 | 0.5 | -0.01 | 0.01 | 0.5 | 0.04 | 0.01 | 0.5 | 0.00 | 0.01 |
| | 1 | 0.06 | 0.02 | 1 | -0.02 | 0.02 | 1 | 0.05 | 0.02 | 1 | 0.03 | 0.02 |
| | 2 | 0.04 | 0.03 | 2 | -0.02 | 0.02 | 2 | 0.09 | 0.03 | 2 | 0.05 | 0.02 |
| | 3 | 0.09 | 0.03 | 3 | -0.04 | 0.02 | 3 | -0.01 | 0.03 | 3 | 0.07 | 0.02 |
| | 5 | 0.10 | 0.03 | 5 | -0.04 | 0.02 | 5 | 0.03 | 0.03 | 5 | 0.06 | 0.02 |
| | 10 | 0.06 | 0.03 | 10 | 0.00 | 0.02 | 10 | 0.05 | 0.03 | 10 | 0.05 | 0.02 |
| | 20 | 0.03 | 0.03 | 20 | 0.01 | 0.03 | 20 | 0.02 | 0.04 | 20 | 0.03 | 0.02 |
| | 30 | 0.05 | 0.03 | 30 | -0.02 | 0.03 | 30 | 0.02 | 0.03 | 30 | 0.03 | 0.03 |
| | 50 | 0.00 | 0.03 | 50 | 0.01 | 0.03 | 50 | 0.01 | 0.03 | 50 | 0.04 | 0.03 |
| | 100 | 0.05 | 0.04 | 100 | -0.06 | 0.03 | 100 | -0.02 | 0.03 | 100 | 0.05 | 0.03 |
| | 300 | 0.06 | 0.04 | 300 | 0.00 | 0.03 | 300 | 0.00 | 0.04 | 300 | 0.01 | 0.03 |
| | 500 | 0.00 | 0.04 | 500 | 0.01 | 0.03 | 500 | 0.08 | 0.04 | 500 | 0.08 | 0.03 |
| | 1000 | 0.04 | 0.03 | 1000 | 0.02 | 0.03 | 1000 | 0.01 | 0.04 | 1000 | 0.07 | 0.03 |
| **OH144** | t (ps) | Av Diff (Å) | SEM (Å) | t (ps) | Av Diff (Å) | SEM (Å) | t (ps) | Av Diff (Å) | SEM (Å) | t (ps) | Av Diff (Å) | SEM (Å) |
| | 0.05 | 0.00 | 0.00 | 0.05 | 0.00 | 0.00 | 0.05 | 0.00 | 0.00 | 0.05 | 0.00 | 0.00 |
| | 0.1 | 0.01 | 0.00 | 0.1 | 0.00 | 0.00 | 0.1 | 0.01 | 0.00 | 0.1 | 0.02 | 0.00 |
| | 0.25 | 0.01 | 0.00 | 0.25 | -0.01 | 0.00 | 0.25 | 0.02 | 0.01 | 0.25 | -0.01 | 0.00 |
| | 0.5 | 0.02 | 0.01 | 0.5 | 0.00 | 0.01 | 0.5 | 0.06 | 0.01 | 0.5 | 0.02 | 0.01 |
| | 1 | 0.04 | 0.02 | 1 | -0.02 | 0.02 | 1 | 0.06 | 0.02 | 1 | 0.03 | 0.02 |
| | 2 | 0.03 | 0.03 | 2 | -0.03 | 0.03 | 2 | 0.10 | 0.04 | 2 | 0.04 | 0.03 |
| | 3 | 0.02 | 0.04 | 3 | -0.03 | 0.03 | 3 | -0.01 | 0.03 | 3 | 0.05 | 0.04 |
| | 5 | 0.08 | 0.03 | 5 | -0.02 | 0.03 | 5 | 0.03 | 0.03 | 5 | 0.04 | 0.03 |
| | 10 | 0.13 | 0.03 | 10 | 0.03 | 0.03 | 10 | 0.03 | 0.04 | 10 | 0.03 | 0.03 |
| | 20 | -0.03 | 0.04 | 20 | -0.01 | 0.03 | 20 | 0.00 | 0.04 | 20 | 0.03 | 0.03 |
| | 30 | 0.06 | 0.04 | 30 | 0.02 | 0.03 | 30 | 0.03 | 0.04 | 30 | 0.01 | 0.04 |
| | 50 | 0.02 | 0.04 | 50 | 0.01 | 0.04 | 50 | -0.03 | 0.04 | 50 | 0.01 | 0.04 |
| | 100 | 0.03 | 0.04 | 100 | -0.05 | 0.03 | 100 | -0.02 | 0.04 | 100 | 0.03 | 0.03 |
| | 300 | 0.08 | 0.04 | 300 | -0.01 | 0.04 | 300 | 0.06 | 0.04 | 300 | 0.04 | 0.04 |
| | 500 | -0.03 | 0.04 | 500 | -0.03 | 0.04 | 500 | 0.04 | 0.04 | 500 | 0.08 | 0.04 |
| | 1000 | 0.03 | 0.04 | 1000 | -0.07 | 0.04 | 1000 | 0.00 | 0.04 | 1000 | -0.03 | 0.04 |
| **OH143** | t (ps) | Av Diff (Å) | SEM (Å) | t (ps) | Av Diff (Å) | SEM (Å) | t (ps) | Av Diff (Å) | SEM (Å) | t (ps) | Av Diff (Å) | SEM (Å) |
| | 0.05 | 0.00 | 0.00 | 0.05 | 0.00 | 0.00 | 0.05 | 0.00 | 0.00 | 0.05 | 0.00 | 0.00 |
| | 0.1 | 0.01 | 0.00 | 0.1 | 0.00 | 0.00 | 0.1 | 0.01 | 0.00 | 0.1 | 0.01 | 0.00 |
| | 0.25 | 0.00 | 0.00 | 0.25 | -0.01 | 0.00 | 0.25 | 0.00 | 0.00 | 0.25 | -0.02 | 0.00 |
| | 0.5 | 0.02 | 0.01 | 0.5 | 0.00 | 0.01 | 0.5 | 0.00 | 0.01 | 0.5 | 0.01 | 0.01 |
| | 1 | -0.01 | 0.02 | 1 | 0.00 | 0.02 | 1 | 0.02 | 0.02 | 1 | -0.03 | 0.02 |
| | 2 | 0.01 | 0.03 | 2 | 0.00 | 0.03 | 2 | -0.05 | 0.03 | 2 | -0.05 | 0.03 |
| | 3 | -0.01 | 0.03 | 3 | 0.01 | 0.03 | 3 | -0.02 | 0.03 | 3 | -0.05 | 0.03 |
| | 5 | -0.02 | 0.03 | 5 | 0.00 | 0.03 | 5 | 0.00 | 0.03 | 5 | -0.05 | 0.03 |
| | 10 | -0.01 | 0.03 | 10 | -0.01 | 0.03 | 10 | 0.01 | 0.03 | 10 | -0.08 | 0.03 |
| | 20 | 0.02 | 0.03 | 20 | -0.05 | 0.03 | 20 | 0.00 | 0.03 | 20 | 0.02 | 0.03 |
| | 30 | 0.01 | 0.03 | 30 | 0.02 | 0.03 | 30 | 0.05 | 0.03 | 30 | 0.00 | 0.03 |
| | 50 | 0.02 | 0.03 | 50 | -0.02 | 0.03 | 50 | 0.02 | 0.03 | 50 | -0.04 | 0.03 |
| | 100 | 0.03 | 0.03 | 100 | 0.03 | 0.03 | 100 | 0.04 | 0.03 | 100 | -0.06 | 0.03 |
| | 300 | -0.03 | 0.03 | 300 | -0.02 | 0.03 | 300 | 0.04 | 0.03 | 300 | 0.02 | 0.03 |
| | 500 | -0.02 | 0.03 | 500 | -0.01 | 0.03 | 500 | -0.02 | 0.03 | 500 | -0.04 | 0.03 |
| | 1000 | 0.03 | 0.03 | 1000 | -0.01 | 0.03 | 1000 | 0.02 | 0.03 | 1000 | -0.05 | 0.03 |

**Table S3.2**: The average differences in the donor-acceptor distances of HBs 1-5 following the shown P2 substitutions in s01, s05, s02, and p12. Positive and negative differences with magnitudes > 0.1 Å are in green and red, respectively.

| HB1-5 | s01: P2 L->A | | | s05: P2 L->A | | | s02: P2 F->A | | | p12: P2 W->A | | |
|---|---|---|---|---|---|---|---|---|---|---|---|---|
| **HB1** | t (ps) | Av Diff (Å) | SEM (Å) | t (ps) | Av Diff (Å) | SEM (Å) | t (ps) | Av Diff (Å) | SEM (Å) | t (ps) | Av Diff (Å) | SEM (Å) |
| | 0.05 | 0.00 | 0.00 | 0.05 | 0.00 | 0.00 | 0.05 | 0.00 | 0.00 | 0.05 | 0.00 | 0.00 |
| | 0.1 | 0.00 | 0.00 | 0.1 | 0.00 | 0.00 | 0.1 | -0.01 | 0.00 | 0.1 | 0.00 | 0.00 |
| | 0.25 | 0.00 | 0.00 | 0.25 | -0.01 | 0.00 | 0.25 | -0.03 | 0.00 | 0.25 | -0.01 | 0.00 |
| | 0.5 | -0.01 | 0.00 | 0.5 | 0.00 | 0.01 | 0.5 | -0.06 | 0.01 | 0.5 | -0.02 | 0.01 |
| | 1 | -0.04 | 0.01 | 1 | 0.00 | 0.02 | 1 | -0.04 | 0.02 | 1 | 0.00 | 0.02 |
| | 2 | 0.00 | 0.02 | 2 | -0.03 | 0.02 | 2 | -0.06 | 0.04 | 2 | -0.04 | 0.02 |
| | 3 | 0.03 | 0.02 | 3 | 0.00 | 0.03 | 3 | -0.07 | 0.04 | 3 | -0.05 | 0.02 |
| | 5 | 0.01 | 0.02 | 5 | 0.01 | 0.03 | 5 | -0.02 | 0.04 | 5 | -0.09 | 0.02 |
| | 10 | 0.00 | 0.02 | 10 | 0.03 | 0.03 | 10 | -0.08 | 0.04 | 10 | -0.05 | 0.03 |
| | 20 | 0.02 | 0.02 | 20 | 0.00 | 0.03 | 20 | -0.02 | 0.04 | 20 | -0.05 | 0.03 |
| | 30 | -0.04 | 0.03 | 30 | 0.02 | 0.03 | 30 | -0.06 | 0.05 | 30 | -0.07 | 0.03 |
| | 50 | -0.01 | 0.03 | 50 | -0.01 | 0.03 | 50 | -0.04 | 0.05 | 50 | -0.05 | 0.03 |
| | 100 | -0.01 | 0.03 | 100 | 0.01 | 0.03 | 100 | -0.16 | 0.07 | 100 | -0.04 | 0.04 |
| | 300 | -0.01 | 0.04 | 300 | 0.01 | 0.04 | 300 | -0.16 | 0.09 | 300 | 0.00 | 0.04 |
| | 500 | -0.04 | 0.03 | 500 | 0.02 | 0.04 | 500 | -0.29 | 0.09 | 500 | -0.02 | 0.04 |
| | 1000 | -0.08 | 0.04 | 1000 | 0.04 | 0.05 | 1000 | -0.40 | 0.10 | 1000 | 0.01 | 0.04 |
| **HB2** | t (ps) | Av Diff (Å) | SEM (Å) | t (ps) | Av Diff (Å) | SEM (Å) | t (ps) | Av Diff (Å) | SEM (Å) | t (ps) | Av Diff (Å) | SEM (Å) |
| | 0.05 | 0.00 | 0.00 | 0.05 | 0.00 | 0.00 | 0.05 | 0.00 | 0.00 | 0.05 | 0.00 | 0.00 |
| | 0.1 | 0.00 | 0.00 | 0.1 | 0.00 | 0.00 | 0.1 | 0.01 | 0.00 | 0.1 | 0.00 | 0.00 |
| | 0.25 | 0.00 | 0.00 | 0.25 | 0.00 | 0.00 | 0.25 | 0.01 | 0.00 | 0.25 | 0.00 | 0.01 |
| | 0.5 | 0.01 | 0.01 | 0.5 | -0.01 | 0.01 | 0.5 | 0.00 | 0.01 | 0.5 | -0.01 | 0.01 |
| | 1 | 0.00 | 0.01 | 1 | 0.00 | 0.01 | 1 | 0.01 | 0.01 | 1 | 0.00 | 0.02 |
| | 2 | -0.01 | 0.02 | 2 | -0.01 | 0.02 | 2 | 0.01 | 0.02 | 2 | 0.00 | 0.02 |
| | 3 | 0.00 | 0.02 | 3 | -0.01 | 0.02 | 3 | 0.04 | 0.02 | 3 | -0.04 | 0.02 |
| | 5 | -0.03 | 0.02 | 5 | -0.02 | 0.02 | 5 | 0.03 | 0.02 | 5 | 0.03 | 0.02 |
| | 10 | 0.00 | 0.02 | 10 | -0.01 | 0.02 | 10 | 0.03 | 0.03 | 10 | -0.04 | 0.02 |
| | 20 | 0.02 | 0.02 | 20 | -0.02 | 0.02 | 20 | 0.00 | 0.02 | 20 | -0.02 | 0.02 |
| | 30 | 0.00 | 0.02 | 30 | 0.00 | 0.02 | 30 | -0.01 | 0.03 | 30 | -0.01 | 0.02 |
| | 50 | 0.00 | 0.02 | 50 | 0.04 | 0.02 | 50 | 0.01 | 0.02 | 50 | 0.02 | 0.02 |
| | 100 | 0.01 | 0.02 | 100 | -0.01 | 0.02 | 100 | 0.00 | 0.04 | 100 | 0.04 | 0.03 |
| | 300 | 0.00 | 0.02 | 300 | 0.01 | 0.02 | 300 | 0.02 | 0.03 | 300 | 0.02 | 0.03 |
| | 500 | 0.02 | 0.02 | 500 | 0.03 | 0.02 | 500 | -0.01 | 0.04 | 500 | 0.10 | 0.03 |
| | 1000 | 0.00 | 0.02 | 1000 | 0.01 | 0.02 | 1000 | 0.06 | 0.06 | 1000 | 0.04 | 0.03 |
| **HB3** | t (ps) | Av Diff (Å) | SEM (Å) | t (ps) | Av Diff (Å) | SEM (Å) | t (ps) | Av Diff (Å) | SEM (Å) | t (ps) | Av Diff (Å) | SEM (Å) |
| | 0.05 | 0.01 | 0.00 | 0.05 | 0.01 | 0.00 | 0.05 | 0.01 | 0.00 | 0.05 | 0.00 | 0.00 |
| | 0.1 | 0.00 | 0.00 | 0.1 | 0.00 | 0.00 | 0.1 | 0.02 | 0.00 | 0.1 | 0.02 | 0.00 |
| | 0.25 | 0.00 | 0.01 | 0.25 | -0.01 | 0.00 | 0.25 | 0.01 | 0.01 | 0.25 | 0.02 | 0.01 |
| | 0.5 | 0.00 | 0.01 | 0.5 | -0.01 | 0.01 | 0.5 | 0.02 | 0.01 | 0.5 | 0.00 | 0.01 |
| | 1 | -0.01 | 0.01 | 1 | -0.02 | 0.01 | 1 | 0.04 | 0.01 | 1 | 0.00 | 0.01 |
| | 2 | 0.00 | 0.01 | 2 | -0.02 | 0.01 | 2 | 0.00 | 0.02 | 2 | 0.01 | 0.01 |
| | 3 | 0.00 | 0.01 | 3 | -0.01 | 0.01 | 3 | 0.02 | 0.02 | 3 | 0.01 | 0.01 |
| | 5 | -0.03 | 0.01 | 5 | -0.03 | 0.02 | 5 | 0.02 | 0.02 | 5 | 0.05 | 0.02 |
| | 10 | 0.01 | 0.01 | 10 | 0.00 | 0.01 | 10 | 0.03 | 0.02 | 10 | 0.00 | 0.01 |
| | 20 | -0.03 | 0.02 | 20 | -0.04 | 0.01 | 20 | -0.01 | 0.02 | 20 | 0.01 | 0.02 |
| | 30 | 0.00 | 0.01 | 30 | -0.01 | 0.01 | 30 | 0.01 | 0.02 | 30 | 0.00 | 0.01 |
| | 50 | 0.02 | 0.01 | 50 | 0.00 | 0.01 | 50 | 0.02 | 0.02 | 50 | 0.02 | 0.02 |
| | 100 | 0.03 | 0.01 | 100 | 0.00 | 0.02 | 100 | 0.01 | 0.02 | 100 | 0.02 | 0.02 |
| | 300 | 0.01 | 0.01 | 300 | 0.01 | 0.02 | 300 | 0.01 | 0.03 | 300 | 0.03 | 0.02 |
| | 500 | 0.00 | 0.01 | 500 | 0.01 | 0.02 | 500 | 0.01 | 0.03 | 500 | 0.06 | 0.02 |
| | 1000 | 0.01 | 0.02 | 1000 | -0.01 | 0.02 | 1000 | 0.06 | 0.04 | 1000 | 0.02 | 0.02 |
| **HB4** | t (ps) | Av Diff (Å) | SEM (Å) | t (ps) | Av Diff (Å) | SEM (Å) | t (ps) | Av Diff (Å) | SEM (Å) | t (ps) | Av Diff (Å) | SEM (Å) |
| | 0.05 | 0.00 | 0.00 | 0.05 | -0.01 | 0.00 | 0.05 | -0.02 | 0.00 | 0.05 | -0.02 | 0.00 |
| | 0.1 | 0.01 | 0.00 | 0.1 | 0.01 | 0.00 | 0.1 | -0.03 | 0.01 | 0.1 | -0.03 | 0.01 |
| | 0.25 | 0.02 | 0.01 | 0.25 | 0.00 | 0.01 | 0.25 | -0.08 | 0.02 | 0.25 | -0.11 | 0.02 |
| | 0.5 | 0.01 | 0.01 | 0.5 | -0.05 | 0.01 | 0.5 | -0.04 | 0.03 | 0.5 | -0.10 | 0.03 |
| | 1 | 0.07 | 0.03 | 1 | 0.00 | 0.02 | 1 | -0.07 | 0.04 | 1 | -0.12 | 0.04 |
| | 2 | 0.08 | 0.03 | 2 | 0.00 | 0.03 | 2 | -0.09 | 0.05 | 2 | -0.15 | 0.06 |
| | 3 | 0.02 | 0.03 | 3 | 0.00 | 0.03 | 3 | -0.06 | 0.05 | 3 | -0.26 | 0.06 |
| | 5 | 0.00 | 0.04 | 5 | -0.02 | 0.04 | 5 | -0.12 | 0.06 | 5 | -0.23 | 0.07 |
| | 10 | -0.01 | 0.04 | 10 | -0.05 | 0.04 | 10 | -0.18 | 0.08 | 10 | -0.36 | 0.07 |
| | 20 | 0.12 | 0.05 | 20 | -0.01 | 0.06 | 20 | -0.12 | 0.09 | 20 | -0.27 | 0.08 |
| | 30 | 0.06 | 0.05 | 30 | 0.01 | 0.05 | 30 | -0.20 | 0.09 | 30 | -0.25 | 0.10 |
| | 50 | 0.04 | 0.08 | 50 | -0.08 | 0.07 | 50 | -0.14 | 0.10 | 50 | -0.27 | 0.10 |
| | 100 | 0.08 | 0.10 | 100 | -0.16 | 0.08 | 100 | -0.37 | 0.12 | 100 | -0.23 | 0.11 |
| | 300 | 0.11 | 0.13 | 300 | -0.12 | 0.10 | 300 | -0.13 | 0.14 | 300 | -0.23 | 0.12 |
| | 500 | -0.07 | 0.15 | 500 | -0.08 | 0.10 | 500 | -0.18 | 0.15 | 500 | -0.10 | 0.12 |
| | 1000 | -0.07 | 0.16 | 1000 | 0.07 | 0.14 | 1000 | -0.33 | 0.16 | 1000 | 0.03 | 0.15 |
| **HB5** | t (ps) | Av Diff (Å) | SEM (Å) | t (ps) | Av Diff (Å) | SEM (Å) | t (ps) | Av Diff (Å) | SEM (Å) | t (ps) | Av Diff (Å) | SEM (Å) |
| | 0.05 | 0.00 | 0.00 | 0.05 | 0.01 | 0.00 | 0.05 | -0.01 | 0.00 | 0.05 | 0.00 | 0.00 |
| | 0.1 | 0.00 | 0.01 | 0.1 | 0.01 | 0.00 | 0.1 | -0.03 | 0.00 | 0.1 | -0.01 | 0.00 |
| | 0.25 | 0.00 | 0.01 | 0.25 | 0.01 | 0.01 | 0.25 | -0.04 | 0.01 | 0.25 | -0.02 | 0.01 |
| | 0.5 | -0.03 | 0.01 | 0.5 | 0.02 | 0.01 | 0.5 | -0.03 | 0.01 | 0.5 | -0.01 | 0.01 |
| | 1 | -0.04 | 0.02 | 1 | -0.03 | 0.01 | 1 | -0.04 | 0.02 | 1 | -0.02 | 0.02 |
| | 2 | -0.05 | 0.03 | 2 | 0.03 | 0.02 | 2 | -0.05 | 0.02 | 2 | -0.03 | 0.03 |
| | 3 | -0.03 | 0.03 | 3 | 0.02 | 0.03 | 3 | -0.01 | 0.03 | 3 | -0.01 | 0.03 |
| | 5 | 0.00 | 0.03 | 5 | 0.03 | 0.03 | 5 | -0.03 | 0.03 | 5 | -0.08 | 0.03 |
| | 10 | 0.03 | 0.03 | 10 | 0.01 | 0.03 | 10 | 0.00 | 0.03 | 10 | -0.07 | 0.03 |
| | 20 | -0.07 | 0.03 | 20 | -0.02 | 0.03 | 20 | -0.01 | 0.03 | 20 | -0.05 | 0.03 |
| | 30 | 0.04 | 0.03 | 30 | 0.02 | 0.03 | 30 | 0.01 | 0.03 | 30 | -0.03 | 0.03 |
| | 50 | -0.01 | 0.03 | 50 | 0.01 | 0.03 | 50 | 0.02 | 0.03 | 50 | -0.03 | 0.03 |
| | 100 | 0.01 | 0.03 | 100 | 0.01 | 0.03 | 100 | -0.03 | 0.03 | 100 | -0.09 | 0.03 |
| | 300 | -0.01 | 0.04 | 300 | 0.07 | 0.03 | 300 | -0.02 | 0.04 | 300 | -0.07 | 0.04 |
| | 500 | 0.00 | 0.04 | 500 | 0.00 | 0.03 | 500 | 0.06 | 0.04 | 500 | -0.01 | 0.04 |
| | 1000 | 0.07 | 0.04 | 1000 | 0.03 | 0.03 | 1000 | 0.09 | 0.04 | 1000 | -0.02 | 0.04 |

**Table S3.3**: The average differences in the donor-acceptor distances of HBs 8-12 following the shown P2 substitutions in s01, s05, s02, and p12. Positive and negative differences with magnitudes > 0.1 Å are in green and red, respectively.

| HB8-12 | s01: P2 L->A | | | s05: P2 L->A | | | s02: P2 F->A | | | p12: P2 W->A | | |
|---|---|---|---|---|---|---|---|---|---|---|---|---|
| HB8 | t (ps) | Av Diff (Å) | SEM (Å) | t (ps) | Av Diff (Å) | SEM (Å) | t (ps) | Av Diff (Å) | SEM (Å) | t (ps) | Av Diff (Å) | SEM (Å) |
| | 0.05 | 0.00 | 0.00 | 0.05 | 0.00 | 0.00 | 0.05 | 0.00 | 0.00 | 0.05 | 0.00 | 0.00 |
| | 0.1 | 0.01 | 0.00 | 0.1 | 0.00 | 0.00 | 0.1 | 0.01 | 0.00 | 0.1 | 0.01 | 0.00 |
| | 0.25 | 0.01 | 0.01 | 0.25 | -0.01 | 0.00 | 0.25 | 0.03 | 0.01 | 0.25 | 0.02 | 0.01 |
| | 0.5 | 0.02 | 0.01 | 0.5 | -0.01 | 0.01 | 0.5 | 0.04 | 0.01 | 0.5 | 0.00 | 0.01 |
| | 1 | 0.06 | 0.02 | 1 | -0.01 | 0.01 | 1 | 0.05 | 0.02 | 1 | 0.03 | 0.02 |
| | 2 | 0.04 | 0.03 | 2 | -0.01 | 0.02 | 2 | 0.08 | 0.03 | 2 | 0.04 | 0.02 |
| | 3 | 0.09 | 0.03 | 3 | -0.03 | 0.02 | 3 | -0.01 | 0.03 | 3 | 0.07 | 0.02 |
| | 5 | 0.10 | 0.03 | 5 | -0.04 | 0.02 | 5 | 0.02 | 0.03 | 5 | 0.06 | 0.02 |
| | 10 | 0.06 | 0.03 | 10 | 0.00 | 0.02 | 10 | 0.04 | 0.03 | 10 | 0.04 | 0.02 |
| | 20 | 0.03 | 0.03 | 20 | 0.01 | 0.03 | 20 | 0.02 | 0.03 | 20 | 0.03 | 0.02 |
| | 30 | 0.05 | 0.03 | 30 | -0.02 | 0.03 | 30 | 0.02 | 0.03 | 30 | 0.03 | 0.03 |
| | 50 | 0.00 | 0.03 | 50 | 0.01 | 0.03 | 50 | 0.01 | 0.03 | 50 | 0.04 | 0.03 |
| | 100 | 0.05 | 0.04 | 100 | -0.06 | 0.03 | 100 | -0.02 | 0.03 | 100 | 0.05 | 0.03 |
| | 300 | 0.06 | 0.03 | 300 | 0.00 | 0.03 | 300 | 0.00 | 0.04 | 300 | 0.01 | 0.03 |
| | 500 | 0.00 | 0.04 | 500 | -0.01 | 0.03 | 500 | 0.08 | 0.04 | 500 | 0.09 | 0.03 |
| | 1000 | 0.04 | 0.03 | 1000 | 0.02 | 0.03 | 1000 | 0.01 | 0.04 | 1000 | 0.06 | 0.03 |
| HB9 | t (ps) | Av Diff (Å) | SEM (Å) | t (ps) | Av Diff (Å) | SEM (Å) | t (ps) | Av Diff (Å) | SEM (Å) | t (ps) | Av Diff (Å) | SEM (Å) |
| | 0.05 | 0.00 | 0.00 | 0.05 | 0.00 | 0.00 | 0.05 | 0.00 | 0.00 | 0.05 | 0.00 | 0.00 |
| | 0.1 | 0.01 | 0.00 | 0.1 | 0.00 | 0.00 | 0.1 | 0.01 | 0.00 | 0.1 | 0.01 | 0.00 |
| | 0.25 | 0.00 | 0.00 | 0.25 | -0.01 | 0.00 | 0.25 | 0.01 | 0.00 | 0.25 | -0.01 | 0.00 |
| | 0.5 | 0.02 | 0.01 | 0.5 | 0.00 | 0.01 | 0.5 | 0.01 | 0.01 | 0.5 | 0.01 | 0.01 |
| | 1 | 0.00 | 0.01 | 1 | 0.00 | 0.01 | 1 | 0.01 | 0.01 | 1 | -0.01 | 0.01 |
| | 2 | 0.02 | 0.02 | 2 | 0.01 | 0.02 | 2 | -0.03 | 0.02 | 2 | -0.02 | 0.02 |
| | 3 | 0.00 | 0.02 | 3 | 0.00 | 0.02 | 3 | -0.01 | 0.02 | 3 | -0.02 | 0.02 |
| | 5 | 0.00 | 0.02 | 5 | 0.00 | 0.02 | 5 | 0.01 | 0.02 | 5 | -0.02 | 0.02 |
| | 10 | 0.01 | 0.02 | 10 | 0.00 | 0.01 | 10 | 0.00 | 0.02 | 10 | -0.05 | 0.02 |
| | 20 | 0.02 | 0.02 | 20 | -0.01 | 0.02 | 20 | 0.00 | 0.02 | 20 | 0.00 | 0.01 |
| | 30 | 0.01 | 0.02 | 30 | -0.01 | 0.02 | 30 | 0.02 | 0.02 | 30 | 0.01 | 0.02 |
| | 50 | -0.01 | 0.02 | 50 | -0.02 | 0.02 | 50 | 0.00 | 0.02 | 50 | -0.02 | 0.02 |
| | 100 | 0.02 | 0.02 | 100 | 0.02 | 0.01 | 100 | 0.03 | 0.02 | 100 | -0.02 | 0.02 |
| | 300 | 0.00 | 0.02 | 300 | -0.02 | 0.02 | 300 | 0.03 | 0.02 | 300 | 0.00 | 0.02 |
| | 500 | 0.00 | 0.02 | 500 | -0.02 | 0.02 | 500 | -0.01 | 0.02 | 500 | -0.03 | 0.02 |
| | 1000 | 0.02 | 0.02 | 1000 | 0.00 | 0.02 | 1000 | 0.00 | 0.02 | 1000 | -0.05 | 0.02 |
| HB10 | t (ps) | Av Diff (Å) | SEM (Å) | t (ps) | Av Diff (Å) | SEM (Å) | t (ps) | Av Diff (Å) | SEM (Å) | t (ps) | Av Diff (Å) | SEM (Å) |
| | 0.05 | 0.00 | 0.00 | 0.05 | 0.00 | 0.00 | 0.05 | 0.00 | 0.00 | 0.05 | 0.00 | 0.00 |
| | 0.1 | 0.00 | 0.00 | 0.1 | 0.00 | 0.00 | 0.1 | 0.00 | 0.00 | 0.1 | 0.00 | 0.00 |
| | 0.25 | 0.00 | 0.00 | 0.25 | 0.00 | 0.00 | 0.25 | 0.02 | 0.00 | 0.25 | 0.02 | 0.00 |
| | 0.5 | 0.00 | 0.00 | 0.5 | 0.00 | 0.00 | 0.5 | 0.01 | 0.00 | 0.5 | -0.01 | 0.00 |
| | 1 | 0.01 | 0.01 | 1 | 0.01 | 0.01 | 1 | 0.02 | 0.01 | 1 | 0.01 | 0.01 |
| | 2 | 0.00 | 0.02 | 2 | -0.01 | 0.02 | 2 | 0.03 | 0.02 | 2 | 0.03 | 0.02 |
| | 3 | 0.00 | 0.02 | 3 | 0.00 | 0.02 | 3 | 0.01 | 0.01 | 3 | 0.01 | 0.02 |
| | 5 | -0.02 | 0.02 | 5 | 0.02 | 0.02 | 5 | 0.04 | 0.02 | 5 | 0.03 | 0.02 |
| | 10 | -0.01 | 0.02 | 10 | 0.00 | 0.02 | 10 | -0.01 | 0.02 | 10 | -0.02 | 0.02 |
| | 20 | -0.03 | 0.02 | 20 | -0.03 | 0.02 | 20 | -0.01 | 0.02 | 20 | -0.01 | 0.02 |
| | 30 | -0.03 | 0.02 | 30 | 0.00 | 0.02 | 30 | 0.01 | 0.02 | 30 | 0.00 | 0.02 |
| | 50 | 0.02 | 0.02 | 50 | 0.00 | 0.02 | 50 | 0.01 | 0.02 | 50 | 0.01 | 0.02 |
| | 100 | -0.01 | 0.02 | 100 | 0.01 | 0.02 | 100 | 0.04 | 0.02 | 100 | 0.02 | 0.02 |
| | 300 | 0.02 | 0.02 | 300 | 0.04 | 0.02 | 300 | 0.00 | 0.02 | 300 | -0.03 | 0.02 |
| | 500 | -0.01 | 0.02 | 500 | -0.02 | 0.02 | 500 | 0.00 | 0.02 | 500 | 0.03 | 0.02 |
| | 1000 | -0.01 | 0.02 | 1000 | 0.03 | 0.02 | 1000 | 0.01 | 0.02 | 1000 | 0.06 | 0.02 |
| HB11 | t (ps) | Av Diff (Å) | SEM (Å) | t (ps) | Av Diff (Å) | SEM (Å) | t (ps) | Av Diff (Å) | SEM (Å) | t (ps) | Av Diff (Å) | SEM (Å) |
| | 0.05 | 0.00 | 0.00 | 0.05 | 0.00 | 0.00 | 0.05 | 0.00 | 0.00 | 0.05 | 0.00 | 0.00 |
| | 0.1 | 0.00 | 0.00 | 0.1 | 0.00 | 0.00 | 0.1 | 0.00 | 0.00 | 0.1 | 0.00 | 0.00 |
| | 0.25 | 0.00 | 0.00 | 0.25 | 0.00 | 0.00 | 0.25 | 0.00 | 0.00 | 0.25 | 0.00 | 0.00 |
| | 0.5 | 0.01 | 0.00 | 0.5 | 0.00 | 0.00 | 0.5 | 0.00 | 0.00 | 0.5 | 0.00 | 0.00 |
| | 1 | -0.01 | 0.01 | 1 | 0.01 | 0.01 | 1 | -0.01 | 0.01 | 1 | -0.01 | 0.01 |
| | 2 | -0.02 | 0.02 | 2 | -0.02 | 0.01 | 2 | -0.01 | 0.02 | 2 | -0.01 | 0.01 |
| | 3 | 0.01 | 0.02 | 3 | 0.00 | 0.02 | 3 | 0.01 | 0.01 | 3 | 0.00 | 0.01 |
| | 5 | -0.02 | 0.02 | 5 | 0.00 | 0.02 | 5 | -0.01 | 0.02 | 5 | 0.01 | 0.01 |
| | 10 | -0.02 | 0.02 | 10 | 0.02 | 0.01 | 10 | -0.01 | 0.01 | 10 | -0.01 | 0.01 |
| | 20 | -0.01 | 0.03 | 20 | 0.03 | 0.02 | 20 | -0.01 | 0.01 | 20 | -0.01 | 0.01 |
| | 30 | 0.00 | 0.02 | 30 | 0.01 | 0.02 | 30 | 0.01 | 0.02 | 30 | -0.03 | 0.01 |
| | 50 | -0.02 | 0.02 | 50 | -0.01 | 0.02 | 50 | 0.01 | 0.02 | 50 | 0.01 | 0.01 |
| | 100 | -0.03 | 0.02 | 100 | 0.01 | 0.02 | 100 | 0.00 | 0.01 | 100 | 0.01 | 0.02 |
| | 300 | -0.01 | 0.02 | 300 | 0.03 | 0.02 | 300 | 0.00 | 0.02 | 300 | 0.02 | 0.01 |
| | 500 | -0.02 | 0.02 | 500 | 0.00 | 0.02 | 500 | -0.03 | 0.02 | 500 | 0.02 | 0.01 |
| | 1000 | 0.03 | 0.02 | 1000 | 0.03 | 0.02 | 1000 | -0.02 | 0.02 | 1000 | 0.04 | 0.01 |
| HB12 | t (ps) | Av Diff (Å) | SEM (Å) | t (ps) | Av Diff (Å) | SEM (Å) | t (ps) | Av Diff (Å) | SEM (Å) | t (ps) | Av Diff (Å) | SEM (Å) |
| | 0.05 | 0.00 | 0.00 | 0.05 | 0.00 | 0.00 | 0.05 | 0.00 | 0.00 | 0.05 | 0.00 | 0.00 |
| | 0.1 | 0.00 | 0.00 | 0.1 | 0.00 | 0.00 | 0.1 | 0.00 | 0.00 | 0.1 | 0.00 | 0.00 |
| | 0.25 | 0.00 | 0.00 | 0.25 | 0.00 | 0.00 | 0.25 | 0.00 | 0.00 | 0.25 | 0.00 | 0.00 |
| | 0.5 | 0.00 | 0.00 | 0.5 | 0.00 | 0.00 | 0.5 | 0.00 | 0.00 | 0.5 | 0.00 | 0.00 |
| | 1 | -0.02 | 0.02 | 1 | 0.00 | 0.01 | 1 | 0.01 | 0.01 | 1 | 0.00 | 0.02 |
| | 2 | -0.01 | 0.03 | 2 | -0.03 | 0.02 | 2 | -0.01 | 0.02 | 2 | 0.00 | 0.02 |
| | 3 | 0.00 | 0.03 | 3 | 0.00 | 0.02 | 3 | -0.01 | 0.02 | 3 | 0.05 | 0.02 |
| | 5 | -0.01 | 0.03 | 5 | -0.03 | 0.02 | 5 | 0.02 | 0.02 | 5 | -0.01 | 0.03 |
| | 10 | -0.05 | 0.03 | 10 | -0.02 | 0.03 | 10 | 0.00 | 0.02 | 10 | -0.01 | 0.03 |
| | 20 | 0.04 | 0.03 | 20 | 0.02 | 0.03 | 20 | 0.00 | 0.02 | 20 | -0.06 | 0.02 |
| | 30 | 0.01 | 0.03 | 30 | 0.01 | 0.03 | 30 | 0.04 | 0.02 | 30 | 0.00 | 0.03 |
| | 50 | -0.09 | 0.04 | 50 | -0.03 | 0.03 | 50 | 0.00 | 0.03 | 50 | -0.02 | 0.02 |
| | 100 | -0.01 | 0.04 | 100 | 0.03 | 0.03 | 100 | -0.04 | 0.02 | 100 | 0.01 | 0.03 |
| | 300 | -0.09 | 0.05 | 300 | 0.00 | 0.03 | 300 | 0.01 | 0.02 | 300 | 0.00 | 0.04 |
| | 500 | 0.02 | 0.05 | 500 | -0.04 | 0.04 | 500 | 0.00 | 0.03 | 500 | -0.03 | 0.04 |
| | 1000 | -0.09 | 0.05 | 1000 | 0.01 | 0.04 | 1000 | -0.01 | 0.03 | 1000 | 0.01 | 0.04 |

**Table S3.4**: The ten most perturbed Cα atoms in the M$^{pro}$-peptide complex in terms of magnitude of the average displacement vector *v*, at selected time points following the P2 substitutions to Ala from Leu, Leu, Phe, and Trp in s01, s05, s02, and p12, respectively. Only displacements ≥ 0.01 Å are shown.

| s01 t(ps) | CA Rank | Atom | Chain | v (Å) | SEM (Å) |
|---|---|---|---|---|---|
| 0.05 | 1 | 5ALA_CA | C | 0.03 | 0.00 |
| | 2 | 165MET_CA | A | 0.01 | 0.00 |
| | 3 | 187ASP_CA | A | 0.01 | 0.00 |
| | 4 | 4VAL_CA | C | 0.01 | 0.00 |
| | 5 | 6GLN_CA | C | 0.01 | 0.00 |
| | 6 | | | | |
| | 7 | | | | |
| | 8 | | | | |
| | 9 | | | | |
| | 10 | | | | |
| 0.5 | 1 | 187ASP_CA | A | 0.10 | 0.01 |
| | 2 | 5ALA_CA | C | 0.10 | 0.02 |
| | 3 | 186VAL_CA | A | 0.08 | 0.01 |
| | 4 | 6GLN_CA | C | 0.08 | 0.01 |
| | 5 | 165MET_CA | A | 0.06 | 0.01 |
| | 6 | 49MET_CA | A | 0.05 | 0.01 |
| | 7 | 4VAL_CA | C | 0.05 | 0.01 |
| | 8 | 41HID_CA | A | 0.04 | 0.00 |
| | 9 | 46SER_CA | A | 0.04 | 0.00 |
| | 10 | 47GLU_CA | A | 0.04 | 0.00 |
| 5 | 1 | 244GLN_CA | B | 0.16 | 0.05 |
| | 2 | 97LYS_CA | B | 0.15 | 0.03 |
| | 3 | 46SER_CA | A | 0.13 | 0.04 |
| | 4 | 301SER_CA | B | 0.13 | 0.03 |
| | 5 | 302GLY_CA | B | 0.13 | 0.04 |
| | 6 | 306GLN_CA | B | 0.13 | 0.03 |
| | 7 | 187ASP_CA | A | 0.12 | 0.03 |
| | 8 | 229ASP_CA | A | 0.12 | 0.04 |
| | 9 | 96PRO_CA | B | 0.12 | 0.04 |
| | 10 | 98THR_CA | B | 0.12 | 0.03 |
| 50 | 1 | 222ARG_CA | B | 0.34 | 0.07 |
| | 2 | 223PHE_CA | B | 0.30 | 0.07 |
| | 3 | 224THR_CA | B | 0.29 | 0.07 |
| | 4 | 225THR_CA | B | 0.25 | 0.06 |
| | 5 | 191ALA_CA | B | 0.24 | 0.07 |
| | 6 | 221ASN_CA | B | 0.22 | 0.05 |
| | 7 | 5ALA_CA | C | 0.22 | 0.03 |
| | 8 | 277ASN_CA | A | 0.20 | 0.09 |
| | 9 | 226THR_CA | B | 0.20 | 0.06 |
| | 10 | 245ASP_CA | B | 0.20 | 0.06 |
| 500 | 1 | 11LYS_CA | C | 0.26 | 0.12 |
| | 2 | 186VAL_CA | A | 0.25 | 0.04 |
| | 3 | 53ASN_CA | A | 0.22 | 0.07 |
| | 4 | 302GLY_CA | B | 0.22 | 0.05 |
| | 5 | 187ASP_CA | A | 0.21 | 0.04 |
| | 6 | 46SER_CA | A | 0.20 | 0.07 |
| | 7 | 93THR_CA | B | 0.20 | 0.08 |
| | 8 | 1THR_CA | C | 0.20 | 0.11 |
| | 9 | 46SER_CA | B | 0.19 | 0.10 |
| | 10 | 2SER_CA | C | 0.19 | 0.07 |
| 1000 | 1 | 11LYS_CA | C | 0.37 | 0.15 |
| | 2 | 47GLU_CA | B | 0.27 | 0.10 |
| | 3 | 306GLN_CA | B | 0.25 | 0.07 |
| | 4 | 45THR_CA | A | 0.23 | 0.07 |
| | 5 | 46SER_CA | B | 0.23 | 0.09 |
| | 6 | 277ASN_CA | B | 0.22 | 0.10 |
| | 7 | 46SER_CA | A | 0.21 | 0.08 |
| | 8 | 186VAL_CA | A | 0.21 | 0.05 |
| | 9 | 193ALA_CA | A | 0.21 | 0.07 |
| | 10 | 248ASP_CA | A | 0.21 | 0.08 |

| s05 t(ps) | CA Rank | Atom | Chain | v (Å) | SEM (Å) |
|---|---|---|---|---|---|
| 0.05 | 1 | 165MET_CA | A | 0.01 | 0.00 |
| | 2 | 4LYS_CA | C | 0.01 | 0.00 |
| | 3 | 5ALA_CA | C | 0.01 | 0.00 |
| | 4 | 6GLN_CA | C | 0.01 | 0.00 |
| | 5 | | | | |
| | 6 | | | | |
| | 7 | | | | |
| | 8 | | | | |
| | 9 | | | | |
| | 10 | | | | |
| 0.5 | 1 | 165MET_CA | A | 0.06 | 0.00 |
| | 2 | 49MET_CA | A | 0.05 | 0.01 |
| | 3 | 4LYS_CA | C | 0.05 | 0.01 |
| | 4 | 44CYS_CA | A | 0.04 | 0.00 |
| | 5 | 47GLU_CA | A | 0.04 | 0.00 |
| | 6 | 48ASP_CA | A | 0.04 | 0.00 |
| | 7 | 143GLY_CA | A | 0.04 | 0.01 |
| | 8 | 173ALA_CA | A | 0.04 | 0.00 |
| | 9 | 6GLN_CA | C | 0.04 | 0.01 |
| | 10 | 41HID_CA | A | 0.03 | 0.00 |
| 5 | 1 | 72ASN_CA | A | 0.16 | 0.05 |
| | 2 | 277ASN_CA | B | 0.16 | 0.05 |
| | 3 | 45THR_CA | A | 0.13 | 0.03 |
| | 4 | 283GLY_CA | B | 0.13 | 0.05 |
| | 5 | 44CYS_CA | A | 0.12 | 0.03 |
| | 6 | 46SER_CA | A | 0.12 | 0.04 |
| | 7 | 73VAL_CA | A | 0.12 | 0.04 |
| | 8 | 194ALA_CA | A | 0.12 | 0.05 |
| | 9 | 227LEU_CA | A | 0.12 | 0.04 |
| | 10 | 228ASN_CA | A | 0.12 | 0.04 |
| 50 | 1 | 47GLU_CA | A | 0.20 | 0.06 |
| | 2 | 53ASN_CA | A | 0.20 | 0.06 |
| | 3 | 46SER_CA | A | 0.19 | 0.06 |
| | 4 | 72ASN_CA | B | 0.19 | 0.09 |
| | 5 | 50LEU_CA | A | 0.18 | 0.06 |
| | 6 | 51ASN_CA | A | 0.18 | 0.06 |
| | 7 | 52PRO_CA | A | 0.18 | 0.06 |
| | 8 | 279ARG_CA | B | 0.18 | 0.06 |
| | 9 | 49MET_CA | A | 0.17 | 0.05 |
| | 10 | 72ASN_CA | A | 0.17 | 0.09 |
| 500 | 1 | 11SER_CA | C | 0.44 | 0.19 |
| | 2 | 277ASN_CA | B | 0.32 | 0.11 |
| | 3 | 278GLY_CA | B | 0.29 | 0.09 |
| | 4 | 10LEU_CA | C | 0.29 | 0.09 |
| | 5 | 277ASN_CA | A | 0.28 | 0.10 |
| | 6 | 53ASN_CA | A | 0.26 | 0.08 |
| | 7 | 72ASN_CA | A | 0.25 | 0.10 |
| | 8 | 190THR_CA | B | 0.25 | 0.08 |
| | 9 | 218TRP_CA | A | 0.23 | 0.06 |
| | 10 | 191ALA_CA | B | 0.23 | 0.10 |
| 1000 | 1 | 11SER_CA | C | 0.36 | 0.17 |
| | 2 | 278GLY_CA | A | 0.21 | 0.12 |
| | 3 | 52PRO_CA | B | 0.21 | 0.10 |
| | 4 | 55GLU_CA | B | 0.21 | 0.08 |
| | 5 | 56ASP_CA | B | 0.21 | 0.11 |
| | 6 | 59ILE_CA | B | 0.20 | 0.10 |
| | 7 | 60ARG_CA | B | 0.20 | 0.09 |
| | 8 | 93THR_CA | B | 0.20 | 0.08 |
| | 9 | 278GLY_CA | B | 0.20 | 0.09 |
| | 10 | 10LEU_CA | C | 0.20 | 0.09 |

| s02 t(ps) | CA Rank | Atom | Chain | v (Å) | SEM (Å) |
|---|---|---|---|---|---|
| 0.05 | 1 | 187ASP_CA | A | 0.02 | 0.00 |
| | 2 | 5ALA_CA | C | 0.02 | 0.00 |
| | 3 | 4THR_CA | C | 0.01 | 0.00 |
| | 4 | | | | |
| | 5 | | | | |
| | 6 | | | | |
| | 7 | | | | |
| | 8 | | | | |
| | 9 | | | | |
| | 10 | | | | |
| 0.5 | 1 | 188ARG_CA | A | 0.13 | 0.01 |
| | 2 | 187ASP_CA | A | 0.12 | 0.01 |
| | 3 | 5ALA_CA | C | 0.11 | 0.01 |
| | 4 | 7SER_CA | C | 0.09 | 0.01 |
| | 5 | 186VAL_CA | A | 0.08 | 0.01 |
| | 6 | 6GLN_CA | C | 0.08 | 0.01 |
| | 7 | 189GLN_CA | A | 0.07 | 0.01 |
| | 8 | 4THR_CA | C | 0.07 | 0.01 |
| | 9 | 49MET_CA | A | 0.06 | 0.01 |
| | 10 | 165MET_CA | A | 0.06 | 0.01 |
| 5 | 1 | 5ALA_CA | C | 0.19 | 0.03 |
| | 2 | 277ASN_CA | B | 0.17 | 0.06 |
| | 3 | 23GLY_CA | A | 0.16 | 0.05 |
| | 4 | 256GLN_CA | B | 0.16 | 0.05 |
| | 5 | 4THR_CA | C | 0.16 | 0.03 |
| | 6 | 47GLU_CA | A | 0.15 | 0.05 |
| | 7 | 277ASN_CA | A | 0.15 | 0.06 |
| | 8 | 278GLY_CA | A | 0.15 | 0.06 |
| | 9 | 46SER_CA | A | 0.14 | 0.04 |

| p12 t(ps) | CA Rank | Atom | Chain | v (Å) | SEM (Å) |
|---|---|---|---|---|---|
| 0.05 | 1 | 5ALA_CA | C | 0.03 | 0.00 |
| | 2 | 187ASP_CA | A | 0.02 | 0.00 |
| | 3 | 49MET_CA | A | 0.01 | 0.00 |
| | 4 | 164HIE_CA | A | 0.01 | 0.00 |
| | 5 | 186VAL_CA | A | 0.01 | 0.00 |
| | 6 | 188ARG_CA | A | 0.01 | 0.00 |
| | 7 | 189GLN_CA | A | 0.01 | 0.00 |
| | 8 | 4PHE_CA | C | 0.01 | 0.00 |
| | 9 | 6GLN_CA | C | 0.01 | 0.00 |
| | 10 | | | | |
| 0.5 | 1 | 186VAL_CA | A | 0.23 | 0.01 |
| | 2 | 187ASP_CA | A | 0.23 | 0.02 |
| | 3 | 5ALA_CA | C | 0.16 | 0.02 |
| | 4 | 164HIE_CA | A | 0.12 | 0.01 |
| | 5 | 165MET_CA | A | 0.12 | 0.01 |
| | 6 | 4PHE_CA | C | 0.12 | 0.01 |
| | 7 | 49MET_CA | A | 0.10 | 0.01 |
| | 8 | 166GLU_CA | A | 0.10 | 0.01 |
| | 9 | 173ALA_CA | A | 0.10 | 0.01 |
| | 10 | 50LEU_CA | A | 0.09 | 0.01 |
| 5 | 1 | 186VAL_CA | A | 0.49 | 0.04 |
| | 2 | 187ASP_CA | A | 0.46 | 0.03 |
| | 3 | 185PHE_CA | A | 0.34 | 0.03 |
| | 4 | 188ARG_CA | A | 0.29 | 0.04 |
| | 5 | 189GLN_CA | A | 0.23 | 0.03 |
| | 6 | 50LEU_CA | A | 0.21 | 0.04 |
| | 7 | 51ASN_CA | A | 0.21 | 0.04 |
| | 8 | 194ALA_CA | A | 0.21 | 0.04 |
| | 9 | 190THR_CA | A | 0.20 | 0.03 |

| | | | | | |
|---|---|---|---|---|---|
| | 10 | 50LEU_CA | A | 0.14 | 0.04 |
| **50** | 1 | 50LEU_CA | A | 0.36 | 0.05 |
| | 2 | 51ASN_CA | A | 0.35 | 0.06 |
| | 3 | 56ASP_CA | A | 0.33 | 0.07 |
| | 4 | 188ARG_CA | A | 0.33 | 0.04 |
| | 5 | 189GLN_CA | A | 0.33 | 0.04 |
| | 6 | 53ASN_CA | A | 0.32 | 0.06 |
| | 7 | 47GLU_CA | A | 0.31 | 0.07 |
| | 8 | 55GLU_CA | A | 0.31 | 0.06 |
| | 9 | 52PRO_CA | A | 0.28 | 0.06 |
| | 10 | 46SER_CA | A | 0.27 | 0.07 |
| **500** | 1 | 1SER_CA | C | 0.57 | 0.19 |
| | 2 | 189GLN_CA | A | 0.36 | 0.05 |
| | 3 | 188ARG_CA | A | 0.35 | 0.05 |
| | 4 | 47GLU_CA | A | 0.29 | 0.09 |
| | 5 | 190THR_CA | A | 0.29 | 0.06 |
| | 6 | 24THR_CA | A | 0.28 | 0.07 |
| | 7 | 50LEU_CA | A | 0.28 | 0.07 |
| | 8 | 187ASP_CA | A | 0.28 | 0.04 |
| | 9 | 48ASP_CA | A | 0.27 | 0.06 |
| | 10 | 51ASN_CA | A | 0.26 | 0.07 |
| **1000** | 1 | 1SER_CA | C | 0.67 | 0.19 |
| | 2 | 47GLU_CA | A | 0.42 | 0.11 |
| | 3 | 189GLN_CA | A | 0.37 | 0.06 |
| | 4 | 50LEU_CA | A | 0.36 | 0.07 |
| | 5 | 188ARG_CA | A | 0.36 | 0.05 |
| | 6 | 46SER_CA | A | 0.35 | 0.10 |
| | 7 | 51ASN_CA | A | 0.35 | 0.07 |
| | 8 | 190THR_CA | A | 0.35 | 0.06 |
| | 9 | 48ASP_CA | A | 0.32 | 0.07 |
| | 10 | 187ASP_CA | A | 0.31 | 0.04 |

| | | | | | |
|---|---|---|---|---|---|
| | 10 | 5ALA_CA | C | 0.19 | 0.03 |
| **50** | 1 | 187ASP_CA | A | 0.44 | 0.04 |
| | 2 | 186VAL_CA | A | 0.41 | 0.04 |
| | 3 | 55GLU_CA | A | 0.35 | 0.06 |
| | 4 | 185PHE_CA | A | 0.33 | 0.04 |
| | 5 | 277ASN_CA | A | 0.33 | 0.09 |
| | 6 | 278GLY_CA | A | 0.31 | 0.09 |
| | 7 | 54TYR_CA | A | 0.30 | 0.04 |
| | 8 | 188ARG_CA | A | 0.30 | 0.05 |
| | 9 | 56ASP_CA | A | 0.29 | 0.06 |
| | 10 | 274ASN_CA | A | 0.29 | 0.08 |
| **500** | 1 | 186VAL_CA | A | 0.51 | 0.05 |
| | 2 | 187ASP_CA | A | 0.47 | 0.04 |
| | 3 | 185PHE_CA | A | 0.39 | 0.04 |
| | 4 | 274ASN_CA | A | 0.38 | 0.09 |
| | 5 | 278GLY_CA | B | 0.36 | 0.10 |
| | 6 | 4PHE_CA | C | 0.36 | 0.05 |
| | 7 | 55GLU_CA | A | 0.35 | 0.07 |
| | 8 | 56ASP_CA | A | 0.32 | 0.08 |
| | 9 | 277ASN_CA | B | 0.31 | 0.11 |
| | 10 | 5ALA_CA | C | 0.29 | 0.04 |
| **1000** | 1 | 186VAL_CA | A | 0.53 | 0.05 |
| | 2 | 187ASP_CA | A | 0.51 | 0.05 |
| | 3 | 4PHE_CA | C | 0.39 | 0.05 |
| | 4 | 185PHE_CA | A | 0.37 | 0.05 |
| | 5 | 47GLU_CA | A | 0.33 | 0.08 |
| | 6 | 59ILE_CA | A | 0.31 | 0.10 |
| | 7 | 5ALA_CA | C | 0.31 | 0.04 |
| | 8 | 46SER_CA | A | 0.26 | 0.08 |
| | 9 | 60ARG_CA | A | 0.26 | 0.11 |
| | 10 | 194ALA_CA | A | 0.25 | 0.07 |

**Table S3.5**: The ten most perturbed non-hydrogen atoms in the M$^{pro}$-peptide complex in terms of magnitude of the average displacement vector *v*, at selected time points following the P2 substitutions to Ala from Leu, Leu, Phe, and Trp in s01, s05, s02, and p12, respectively.

| *s01 t(ps)* | *Non-H Rank* | *Atom* | *Chain* | *v (Å)* | *SEM (Å)* |
|---|---|---|---|---|---|
| **0.05** | 1 | 5ALA_N | C | 0.05 | 0.00 |
| | 2 | 5ALA_CB | C | 0.04 | 0.01 |
| | 3 | 165MET_CB | A | 0.03 | 0.00 |
| | 4 | 5ALA_CA | C | 0.03 | 0.00 |
| | 5 | 4VAL_O | C | 0.03 | 0.00 |
| | 6 | 4VAL_C | C | 0.02 | 0.00 |
| | 7 | 165MET_CG | A | 0.02 | 0.00 |
| | 8 | 41HID_ND1 | A | 0.02 | 0.00 |
| | 9 | 5ALA_C | C | 0.02 | 0.00 |
| | 10 | 49MET_CB | A | 0.02 | 0.00 |
| **0.5** | 1 | 165MET_CE | A | 0.20 | 0.02 |
| | 2 | 49MET_SD | A | 0.20 | 0.03 |
| | 3 | 187ASP_O | A | 0.19 | 0.02 |
| | 4 | 165MET_SD | A | 0.19 | 0.02 |
| | 5 | 5ALA_CB | C | 0.17 | 0.03 |
| | 6 | 165MET_CG | A | 0.15 | 0.02 |
| | 7 | 165MET_CB | A | 0.12 | 0.01 |
| | 8 | 187ASP_C | A | 0.12 | 0.01 |
| | 9 | 5ALA_N | C | 0.12 | 0.01 |
| | 10 | 49MET_CE | A | 0.11 | 0.02 |
| **5** | 1 | 47GLU_OE2 | A | 0.38 | 0.10 |
| | 2 | 49MET_SD | A | 0.34 | 0.06 |
| | 3 | 47GLU_OE2 | B | 0.26 | 0.09 |
| | 4 | 97LYS_CE | B | 0.26 | 0.06 |
| | 5 | 236LYS_NZ | A | 0.26 | 0.10 |
| | 6 | 107GLN_OE1 | A | 0.25 | 0.08 |
| | 7 | 244GLN_OE1 | B | 0.23 | 0.08 |
| | 8 | 245ASP_OD2 | B | 0.23 | 0.08 |
| | 9 | 97LYS_CD | B | 0.23 | 0.05 |
| | 10 | 47GLU_CD | A | 0.23 | 0.08 |
| **50** | 1 | 222ARG_NH1 | B | 0.49 | 0.18 |
| | 2 | 11LYS_NZ | C | 0.49 | 0.17 |
| | 3 | 222ARG_NH1 | A | 0.48 | 0.18 |
| | 4 | 49MET_SD | A | 0.48 | 0.07 |
| | 5 | 244GLN_NE2 | B | 0.45 | 0.15 |
| | 6 | 235MET_CE | B | 0.44 | 0.14 |
| | 7 | 11LYS_CE | C | 0.44 | 0.15 |
| | 8 | 222ARG_CG | B | 0.43 | 0.11 |
| | 9 | 222ARG_NH2 | B | 0.43 | 0.17 |
| | 10 | 222ARG_CZ | B | 0.42 | 0.16 |
| **500** | 1 | 165MET_CE | A | 1.26 | 0.13 |
| | 2 | 165MET_SD | A | 0.96 | 0.10 |
| | 3 | 49MET_CE | A | 0.72 | 0.13 |
| | 4 | 306GLN_NE2 | A | 0.64 | 0.23 |
| | 5 | 165MET_CG | A | 0.63 | 0.07 |
| | 6 | 72ASN_OD1 | B | 0.61 | 0.20 |
| | 7 | 222ARG_NH2 | A | 0.54 | 0.25 |
| | 8 | 49MET_SD | A | 0.53 | 0.09 |
| | 9 | 279ARG_NH1 | B | 0.47 | 0.19 |
| | 10 | 248ASP_OD1 | A | 0.46 | 0.12 |
| **1000** | 1 | 165MET_CE | A | 1.29 | 0.14 |
| | 2 | 49MET_CE | A | 1.18 | 0.14 |
| | 3 | 165MET_SD | A | 1.09 | 0.10 |
| | 4 | 279ARG_NH1 | B | 0.94 | 0.22 |
| | 5 | 11LYS_NZ | C | 0.86 | 0.32 |
| | 6 | 165MET_CG | A | 0.77 | 0.07 |
| | 7 | 11LYS_CE | C | 0.76 | 0.28 |
| | 8 | 279ARG_CZ | B | 0.72 | 0.17 |
| | 9 | 279ARG_NH2 | B | 0.61 | 0.18 |
| | 10 | 279ARG_NE | B | 0.61 | 0.13 |

| *s05 t(ps)* | *Non-H Rank* | *Atom* | *Chain* | *v (Å)* | *SEM (Å)* |
|---|---|---|---|---|---|
| **0.05** | 1 | 5ALA_CB | C | 0.04 | 0.01 |
| | 2 | 5ALA_N | C | 0.04 | 0.00 |
| | 3 | 4LYS_O | C | 0.02 | 0.00 |
| | 4 | 165MET_CB | A | 0.02 | 0.00 |
| | 5 | 5ALA_O | C | 0.02 | 0.00 |
| | 6 | 5ALA_C | C | 0.02 | 0.00 |
| | 7 | 165MET_CG | A | 0.02 | 0.00 |
| | 8 | 41HID_ND1 | A | 0.02 | 0.00 |
| | 9 | 4LYS_CB | C | 0.02 | 0.00 |
| | 10 | 41HID_CG | A | 0.02 | 0.00 |
| **0.5** | 1 | 49MET_SD | A | 0.19 | 0.03 |
| | 2 | 165MET_SD | A | 0.17 | 0.02 |
| | 3 | 165MET_CG | A | 0.13 | 0.02 |
| | 4 | 165MET_CE | A | 0.13 | 0.03 |
| | 5 | 49MET_CE | A | 0.13 | 0.02 |
| | 6 | 165MET_CB | A | 0.10 | 0.01 |
| | 7 | 49MET_CG | A | 0.09 | 0.01 |
| | 8 | 49MET_CB | A | 0.08 | 0.01 |
| | 9 | 165MET_O | A | 0.07 | 0.01 |
| | 10 | 189GLN_CD | A | 0.07 | 0.01 |
| **5** | 1 | 49MET_SD | A | 0.31 | 0.05 |
| | 2 | 244GLN_NE2 | B | 0.27 | 0.08 |
| | 3 | 72ASN_ND2 | A | 0.26 | 0.10 |
| | 4 | 49MET_CE | A | 0.25 | 0.05 |
| | 5 | 47GLU_OE1 | B | 0.25 | 0.11 |
| | 6 | 51ASN_ND2 | A | 0.23 | 0.08 |
| | 7 | 256GLN_NE2 | A | 0.22 | 0.06 |
| | 8 | 235MET_CE | A | 0.21 | 0.10 |
| | 9 | 72ASN_CB | A | 0.21 | 0.07 |
| | 10 | 228ASN_ND2 | A | 0.21 | 0.07 |
| **50** | 1 | 222ARG_NH1 | A | 0.46 | 0.16 |
| | 2 | 49MET_SD | A | 0.46 | 0.06 |
| | 3 | 222ARG_NH2 | A | 0.46 | 0.15 |
| | 4 | 49MET_CE | A | 0.42 | 0.08 |
| | 5 | 222ARG_CZ | A | 0.40 | 0.13 |
| | 6 | 189GLN_NE2 | A | 0.39 | 0.08 |
| | 7 | 49MET_CG | A | 0.37 | 0.06 |
| | 8 | 244GLN_NE2 | A | 0.36 | 0.15 |
| | 9 | 72ASN_ND2 | A | 0.35 | 0.17 |
| | 10 | 244GLN_NE2 | B | 0.34 | 0.15 |
| **500** | 1 | 49MET_CE | A | 0.88 | 0.12 |
| | 2 | 165MET_SD | A | 0.74 | 0.09 |
| | 3 | 49MET_SD | A | 0.72 | 0.08 |
| | 4 | 256GLN_NE2 | A | 0.66 | 0.15 |
| | 5 | 165MET_CG | A | 0.58 | 0.06 |
| | 6 | 11SER_OC2 | C | 0.56 | 0.23 |
| | 7 | 277ASN_OD1 | A | 0.55 | 0.19 |
| | 8 | 165MET_CE | A | 0.53 | 0.10 |
| | 9 | 235MET_CE | B | 0.51 | 0.20 |
| | 10 | 223PHE_CE2 | B | 0.49 | 0.16 |
| **1000** | 1 | 49MET_CE | A | 1.13 | 0.13 |
| | 2 | 235MET_CE | A | 0.87 | 0.22 |
| | 3 | 222ARG_NH2 | B | 0.83 | 0.32 |
| | 4 | 222ARG_NH1 | B | 0.82 | 0.29 |
| | 5 | 222ARG_CZ | B | 0.76 | 0.27 |
| | 6 | 165MET_SD | A | 0.71 | 0.09 |
| | 7 | 49MET_SD | A | 0.67 | 0.08 |
| | 8 | 236LYS_NZ | A | 0.62 | 0.21 |
| | 9 | 235MET_SD | A | 0.61 | 0.17 |
| | 10 | 222ARG_NE | B | 0.61 | 0.24 |

| **s02 t(ps)** | **Non-H** Rank | Atom | Chain | v (Å) | SEM (Å) |
|---|---|---|---|---|---|
| **0.05** | 1 | 5ALA_CB | C | 0.04 | 0.01 |
| | 2 | 5ALA_N | C | 0.03 | 0.00 |
| | 3 | 49MET_CE | A | 0.03 | 0.00 |
| | 4 | 5ALA_CA | C | 0.02 | 0.00 |
| | 5 | 187ASP_CA | A | 0.02 | 0.00 |
| | 6 | 5ALA_C | C | 0.01 | 0.00 |
| | 7 | 49MET_SD | A | 0.01 | 0.00 |
| | 8 | 188ARG_O | A | 0.01 | 0.00 |
| | 9 | 4THR_C | C | 0.01 | 0.00 |
| | 10 | 165MET_SD | A | 0.01 | 0.00 |
| **0.5** | 1 | 165MET_SD | A | 0.32 | 0.03 |
| | 2 | 165MET_CE | A | 0.31 | 0.03 |
| | 3 | 49MET_CE | A | 0.22 | 0.03 |
| | 4 | 165MET_CG | A | 0.19 | 0.02 |
| | 5 | 188ARG_O | A | 0.16 | 0.02 |
| | 6 | 49MET_SD | A | 0.16 | 0.02 |
| | 7 | 5ALA_N | C | 0.14 | 0.02 |
| | 8 | 188ARG_N | A | 0.13 | 0.01 |
| | 9 | 188ARG_CB | A | 0.13 | 0.01 |
| | 10 | 187ASP_O | A | 0.13 | 0.01 |
| **5** | 1 | 165MET_CE | A | 0.38 | 0.07 |
| | 2 | 165MET_SD | A | 0.29 | 0.05 |
| | 3 | 4THR_CG2 | C | 0.25 | 0.05 |
| | 4 | 72ASN_ND2 | B | 0.23 | 0.09 |
| | 5 | 134PHE_CZ | A | 0.23 | 0.07 |
| | 6 | 5ALA_N | C | 0.22 | 0.03 |
| | 7 | 80HID_NE2 | A | 0.22 | 0.07 |
| | 8 | 238ASN_OD1 | B | 0.22 | 0.06 |
| | 9 | 277ASN_OD1 | B | 0.22 | 0.09 |
| | 10 | 107GLN_NE2 | A | 0.22 | 0.09 |
| **50** | 1 | 222ARG_NH2 | B | 0.55 | 0.20 |
| | 2 | 222ARG_NH1 | B | 0.53 | 0.21 |
| | 3 | 165MET_CE | A | 0.52 | 0.09 |
| | 4 | 50LEU_CD2 | A | 0.51 | 0.09 |
| | 5 | 56ASP_OD1 | A | 0.48 | 0.09 |
| | 6 | 222ARG_CZ | B | 0.45 | 0.18 |
| | 7 | 72ASN_OD1 | A | 0.44 | 0.17 |
| | 8 | 67LEU_CD2 | B | 0.43 | 0.10 |
| | 9 | 51ASN_OD1 | A | 0.42 | 0.10 |
| | 10 | 50LEU_CG | A | 0.41 | 0.07 |
| **500** | 1 | 1SER_OG | C | 0.75 | 0.28 |
| | 2 | 1SER_N | C | 0.70 | 0.23 |
| | 3 | 165MET_CE | A | 0.68 | 0.10 |
| | 4 | 1SER_CB | C | 0.58 | 0.23 |
| | 5 | 1SER_CA | C | 0.57 | 0.19 |
| | 6 | 11ARG_NH2 | C | 0.55 | 0.38 |
| | 7 | 235MET_CE | A | 0.54 | 0.22 |
| | 8 | 165MET_SD | A | 0.53 | 0.09 |
| | 9 | 235MET_SD | A | 0.52 | 0.17 |
| | 10 | 60ARG_NH1 | A | 0.51 | 0.23 |
| **1000** | 1 | 1SER_OG | C | 1.04 | 0.27 |
| | 2 | 1SER_CB | C | 0.88 | 0.23 |
| | 3 | 294PHE_CE1 | B | 0.71 | 0.16 |
| | 4 | 294PHE_CZ | B | 0.69 | 0.17 |
| | 5 | 1SER_N | C | 0.68 | 0.24 |
| | 6 | 1SER_CA | C | 0.67 | 0.19 |
| | 7 | 55GLU_OE2 | A | 0.67 | 0.19 |
| | 8 | 165MET_CE | A | 0.63 | 0.12 |
| | 9 | 46SER_OG | A | 0.58 | 0.15 |
| | 10 | 72ASN_ND2 | B | 0.54 | 0.21 |

| **p12 t(ps)** | **Non-H** Rank | Atom | Chain | v (Å) | SEM (Å) |
|---|---|---|---|---|---|
| **0.05** | 1 | 5ALA_N | C | 0.06 | 0.01 |
| | 2 | 5ALA_CB | C | 0.04 | 0.01 |
| | 3 | 187ASP_O | A | 0.04 | 0.00 |
| | 4 | 5ALA_O | C | 0.03 | 0.00 |
| | 5 | 5ALA_CA | C | 0.03 | 0.00 |
| | 6 | 5ALA_C | C | 0.03 | 0.00 |
| | 7 | 187ASP_CA | A | 0.02 | 0.00 |
| | 8 | 186VAL_O | A | 0.02 | 0.00 |
| | 9 | 181PHE_CZ | A | 0.02 | 0.00 |
| | 10 | 4PHE_C | C | 0.02 | 0.00 |
| **0.5** | 1 | 165MET_SD | A | 0.46 | 0.04 |
| | 2 | 165MET_CE | A | 0.37 | 0.03 |
| | 3 | 187ASP_O | A | 0.33 | 0.02 |
| | 4 | 186VAL_O | A | 0.30 | 0.02 |
| | 5 | 165MET_CG | A | 0.27 | 0.02 |
| | 6 | 186VAL_CG2 | A | 0.26 | 0.02 |
| | 7 | 186VAL_C | A | 0.26 | 0.02 |
| | 8 | 186VAL_CB | A | 0.25 | 0.01 |
| | 9 | 187ASP_N | A | 0.24 | 0.02 |
| | 10 | 187ASP_CA | A | 0.24 | 0.02 |
| **5** | 1 | 165MET_SD | A | 0.64 | 0.07 |
| | 2 | 187ASP_O | A | 0.51 | 0.04 |
| | 3 | 186VAL_O | A | 0.50 | 0.04 |
| | 4 | 186VAL_C | A | 0.50 | 0.04 |
| | 5 | 165MET_CE | A | 0.50 | 0.07 |
| | 6 | 186VAL_CA | A | 0.49 | 0.04 |
| | 7 | 187ASP_N | A | 0.49 | 0.04 |
| | 8 | 186VAL_CG2 | A | 0.48 | 0.04 |
| | 9 | 186VAL_CB | A | 0.47 | 0.04 |
| | 10 | 187ASP_CA | A | 0.46 | 0.03 |
| **50** | 1 | 165MET_SD | A | 1.00 | 0.09 |
| | 2 | 165MET_CE | A | 0.77 | 0.08 |
| | 3 | 4PHE_CZ | C | 0.60 | 0.10 |
| | 4 | 4PHE_CE2 | C | 0.57 | 0.10 |
| | 5 | 4PHE_CE1 | C | 0.55 | 0.10 |
| | 6 | 49MET_CE | A | 0.52 | 0.10 |
| | 7 | 277ASN_OD1 | A | 0.52 | 0.15 |
| | 8 | 187ASP_O | A | 0.51 | 0.05 |
| | 9 | 277ASN_ND2 | A | 0.50 | 0.14 |
| | 10 | 49MET_SD | A | 0.49 | 0.09 |
| **500** | 1 | 165MET_SD | A | 1.00 | 0.09 |
| | 2 | 4PHE_CE1 | C | 0.91 | 0.17 |
| | 3 | 165MET_CE | A | 0.89 | 0.09 |
| | 4 | 4PHE_CZ | C | 0.89 | 0.19 |
| | 5 | 4PHE_CD1 | C | 0.78 | 0.13 |
| | 6 | 4PHE_CE2 | C | 0.75 | 0.18 |
| | 7 | 11TYR_OH | C | 0.74 | 0.26 |
| | 8 | 55GLU_OE1 | A | 0.63 | 0.18 |
| | 9 | 49MET_CE | A | 0.62 | 0.13 |
| | 10 | 11TYR_CZ | C | 0.62 | 0.22 |
| **1000** | 1 | 4PHE_CZ | C | 1.11 | 0.18 |
| | 2 | 165MET_SD | A | 1.04 | 0.09 |
| | 3 | 4PHE_CE2 | C | 1.01 | 0.18 |
| | 4 | 4PHE_CE1 | C | 1.00 | 0.17 |
| | 5 | 1LYS_NZ | C | 0.89 | 0.24 |
| | 6 | 165MET_CE | A | 0.88 | 0.10 |
| | 7 | 4PHE_CD1 | C | 0.80 | 0.13 |
| | 8 | 4PHE_CD2 | C | 0.79 | 0.14 |
| | 9 | 1LYS_CE | C | 0.74 | 0.21 |
| | 10 | 49MET_CE | A | 0.71 | 0.15 |

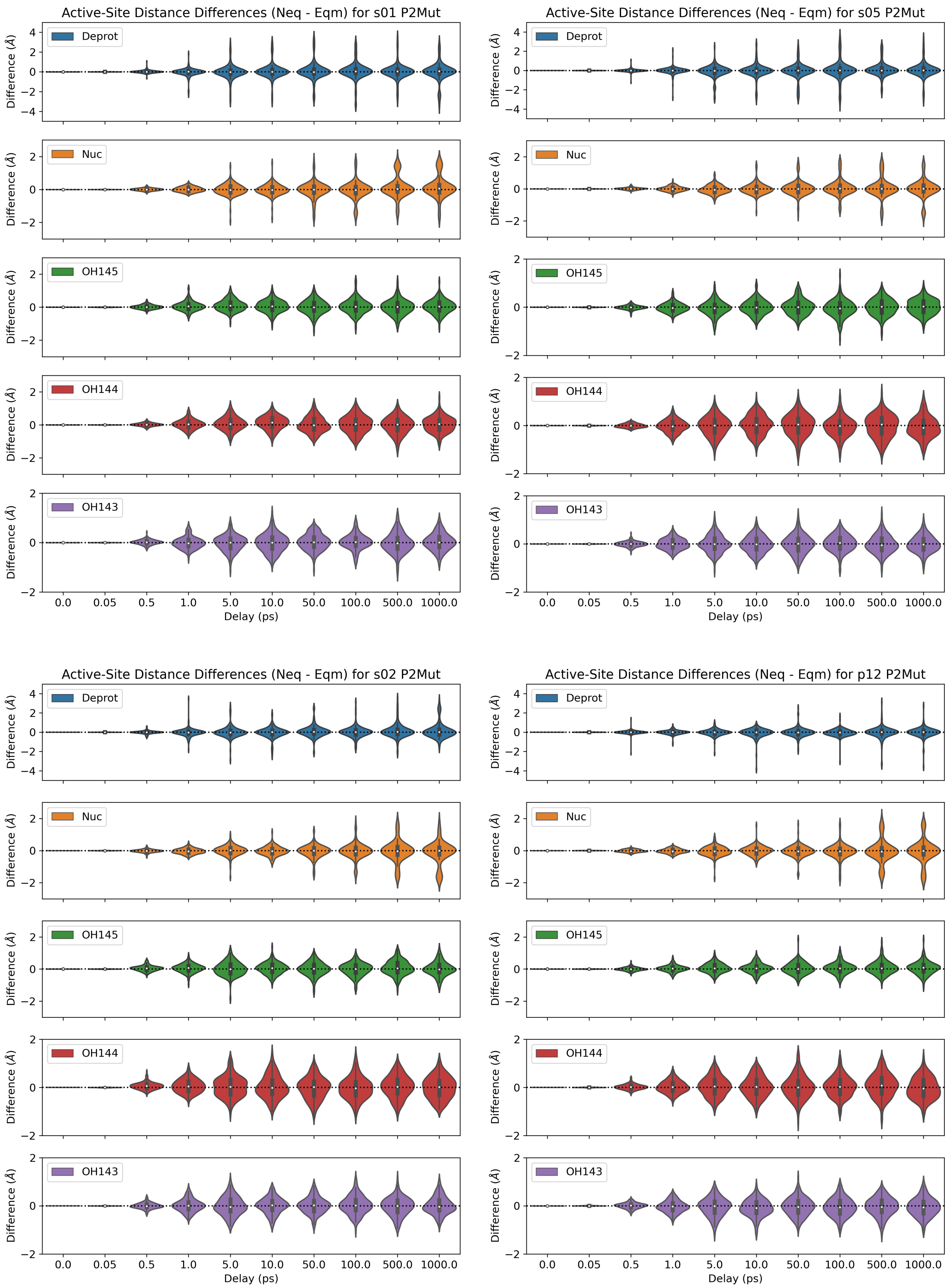

**Figure S3.1**: Distributions of the distance differences in the five active-site distances ($\Delta d = d_{\text{neq}} - d_{\text{eqm}}$) following the P2 substitutions to Ala from Leu, Leu, Phe, and Trp in s01, s05, s02, and p12, respectively.

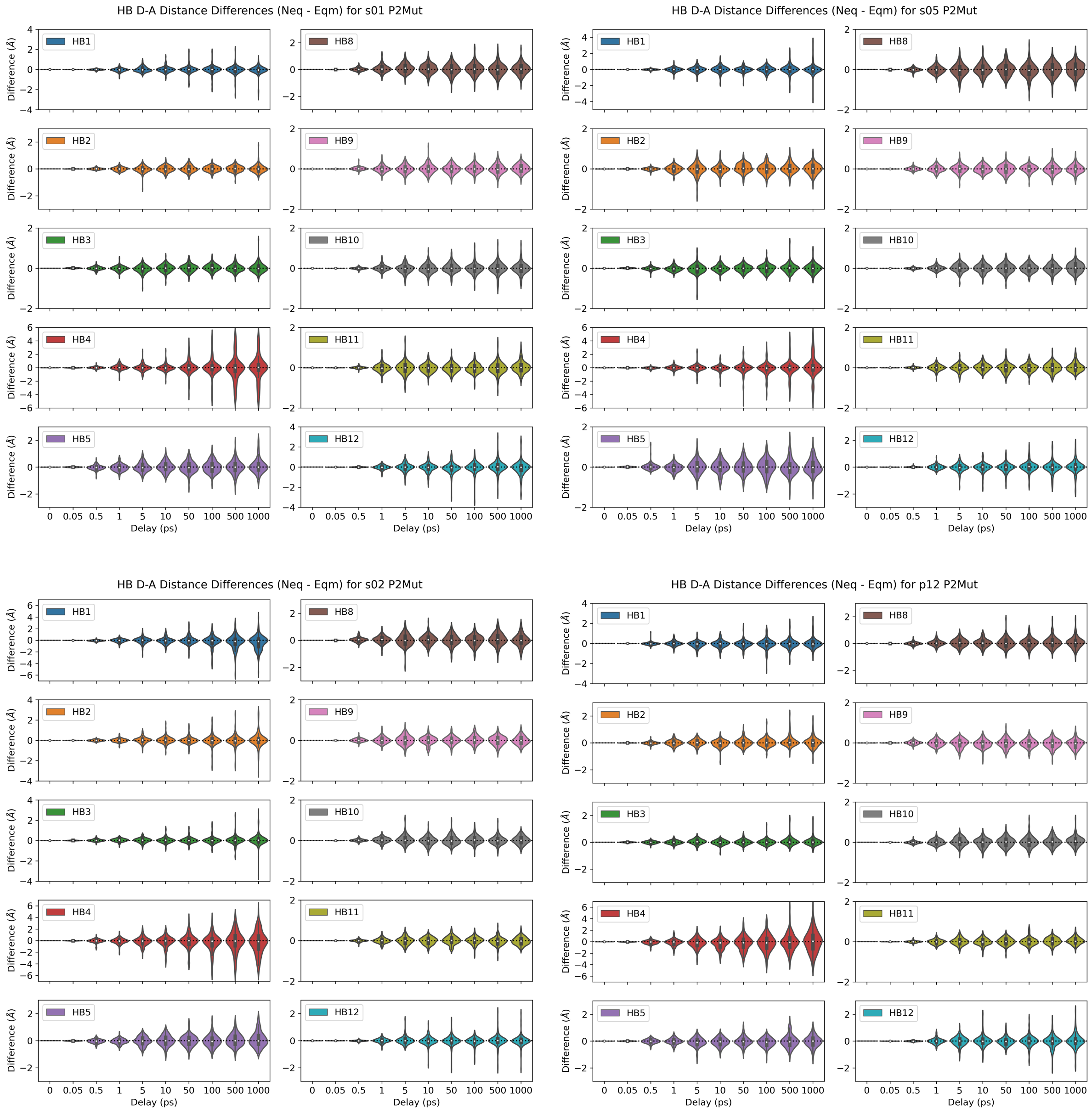

**Figure S3.2**: Distributions of the distance differences in the ten donor-acceptor distances of HBs 1-5 and 8-12 following the P2 substitutions to Ala from Leu, Leu, Phe, and Trp in s01, s05, s02, and p12, respectively.

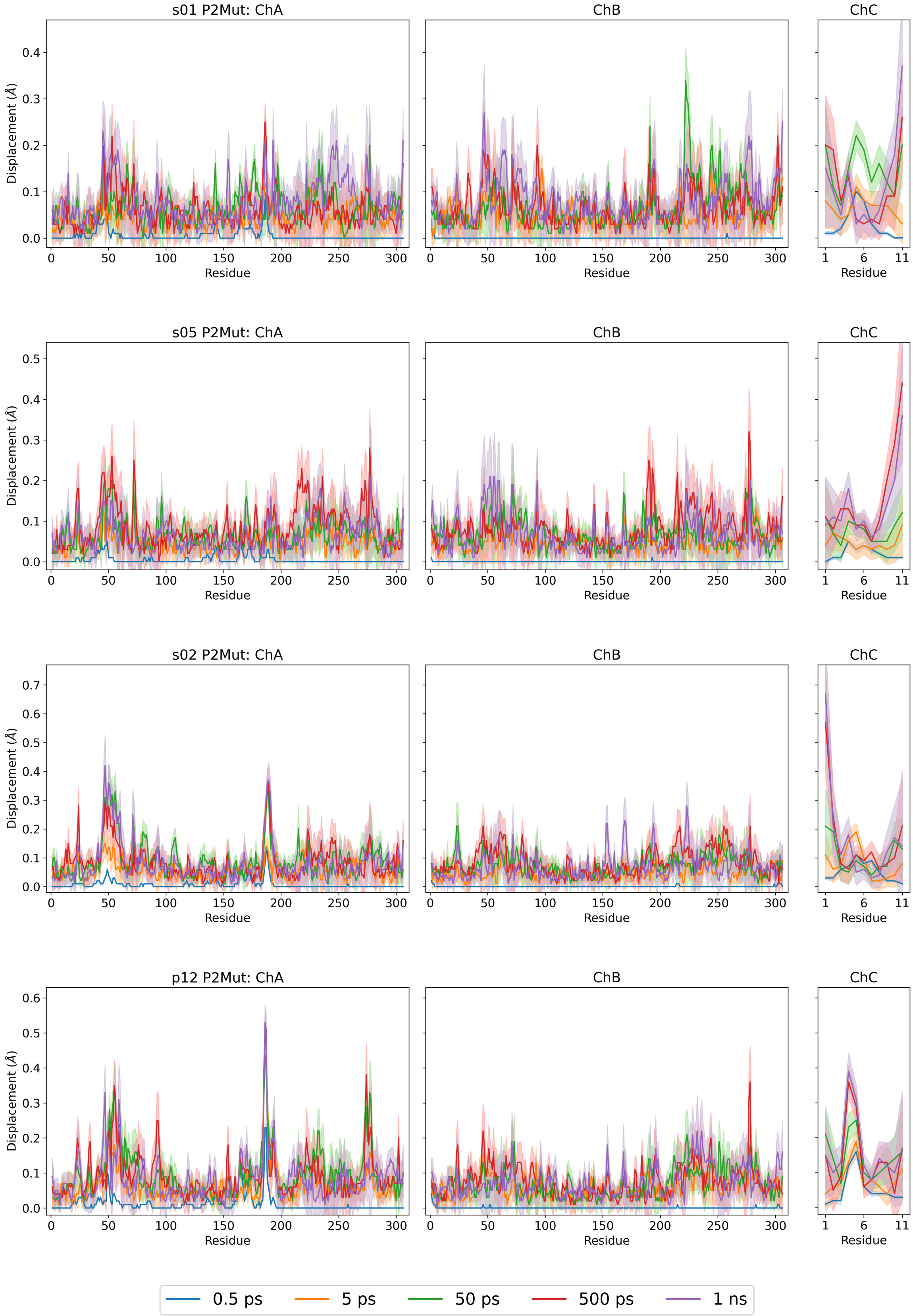

**Figure S3.3**: Magnitudes of the average Cα displacement vectors, with ± SEM shown as shaded regions, following the P2 substitutions to Ala from Leu, Leu, Phe, and Trp in s01, s05, s02, and p12, respectively.

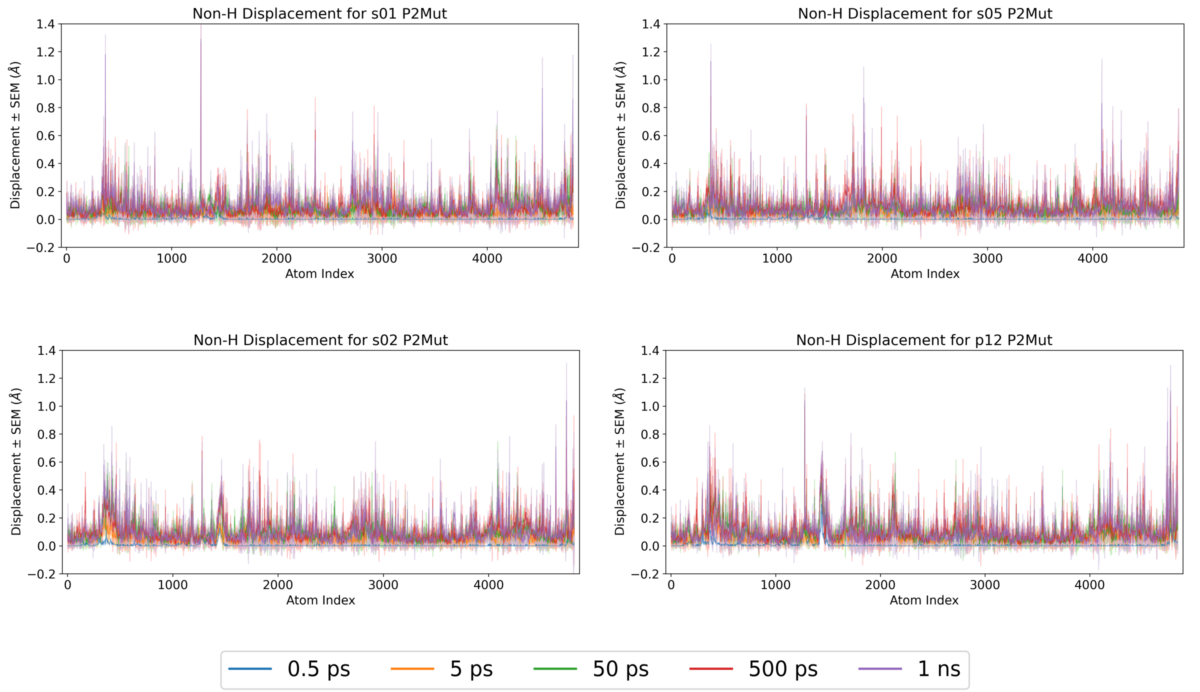

**Figure S3.4**: Magnitudes of the average non-hydrogen atom displacement vectors in the M$^{pro}$-peptide complex, with ± SEM shown as shaded regions, following the P2 substitutions to Ala from Leu, Leu, Phe, and Trp in s01, s05, s02, and p12, respectively.

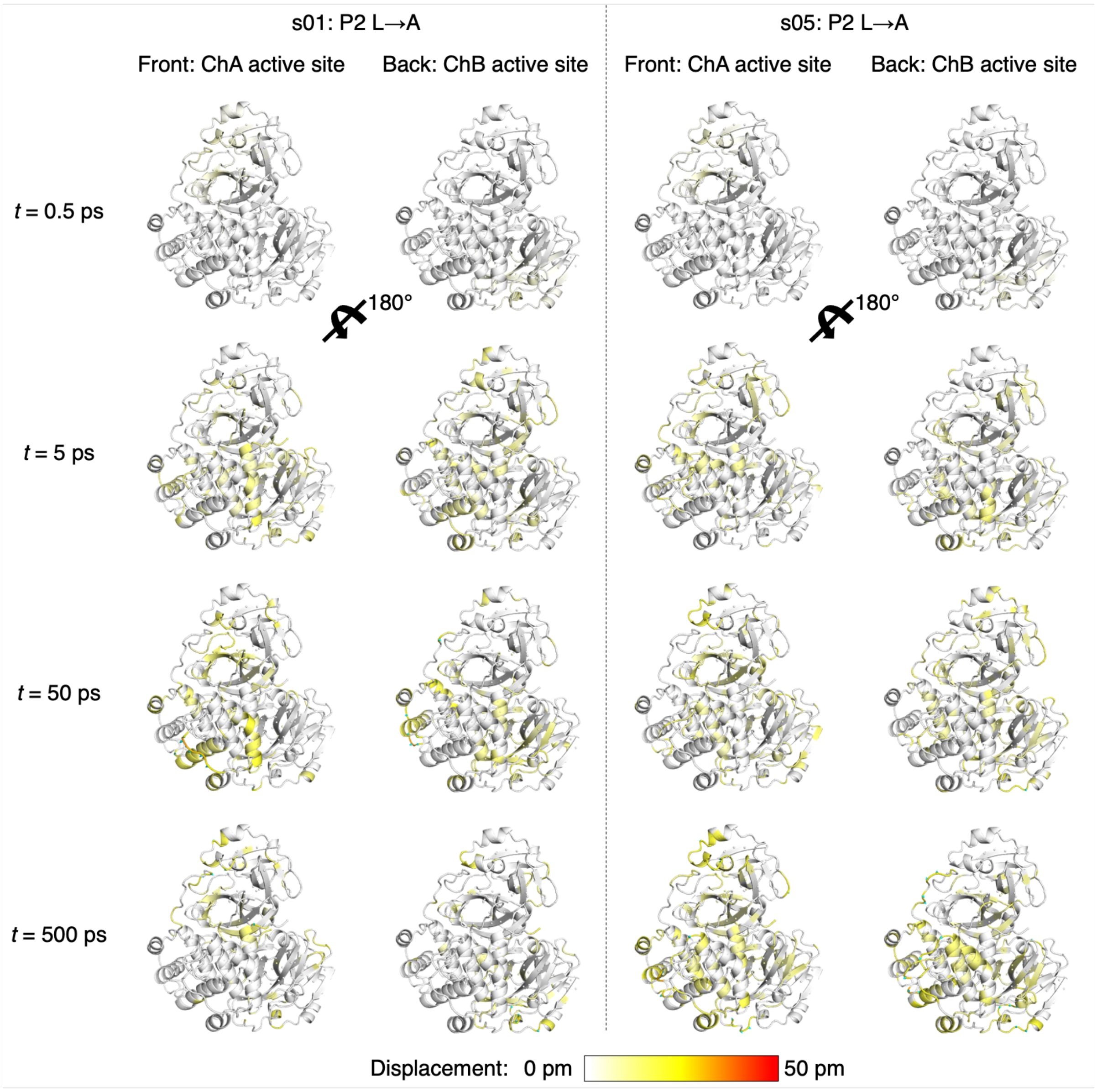
s01: P2 L→A
s05: P2 L→A
Front: ChA active site
Back: ChB active site
Front: ChA active site
Back: ChB active site
t = 0.5 ps
t = 5 ps
t = 50 ps
t = 500 ps
180°
180°
Displacement: 0 pm
50 pm

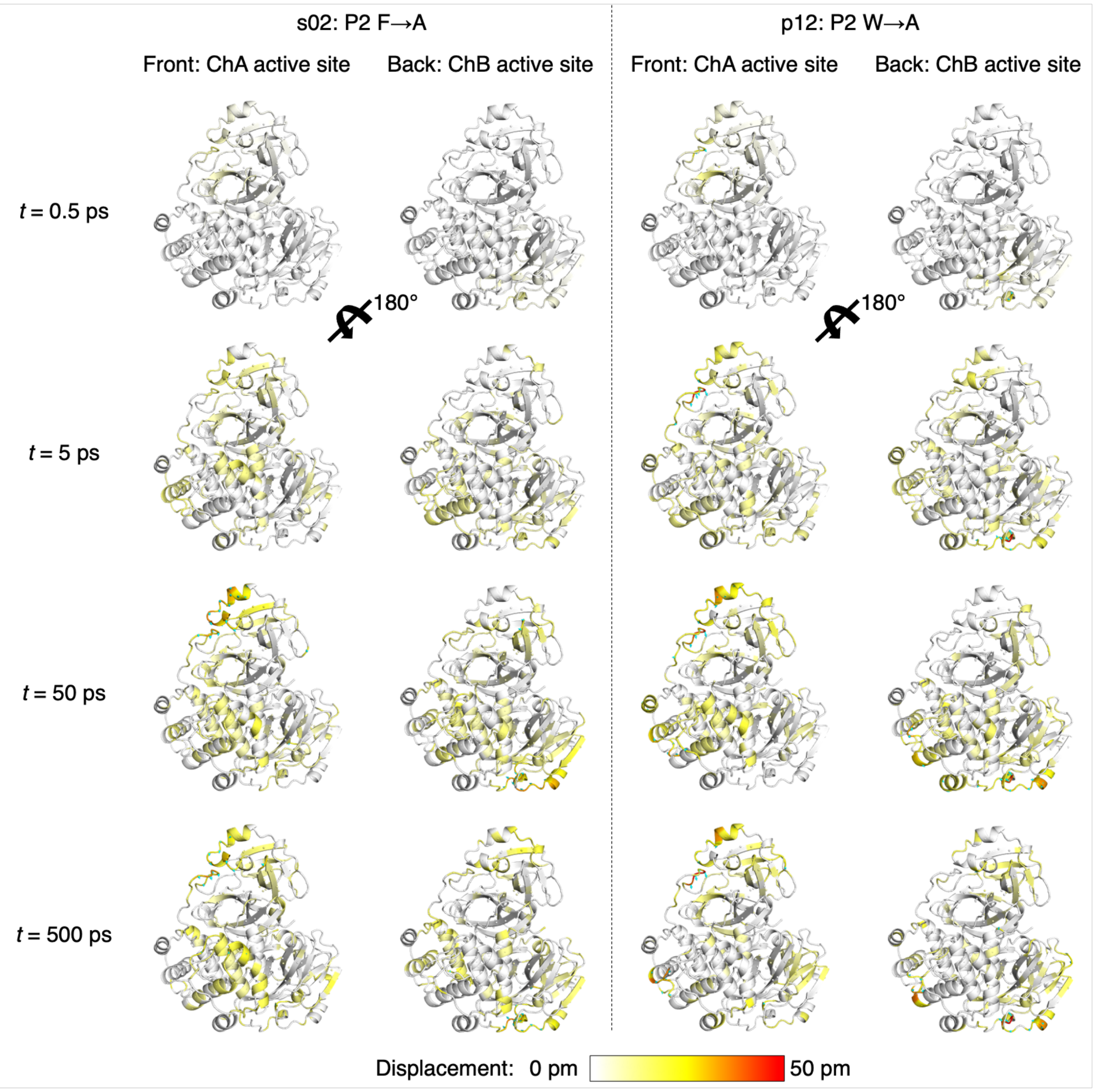

**Figure S3.5**: Views of the M$^{pro}$ response to the shown P2 substitutions in s01, s05, s02, and p12 from averaging Cα displacement vectors, shown using the structure prior to MD simulations. Displacement magnitudes are represented on a white-yellow-red scale.[7] Significant vectors with length ≥20 pm are displayed as cyan arrows with a scale-up factor of 5.[8]

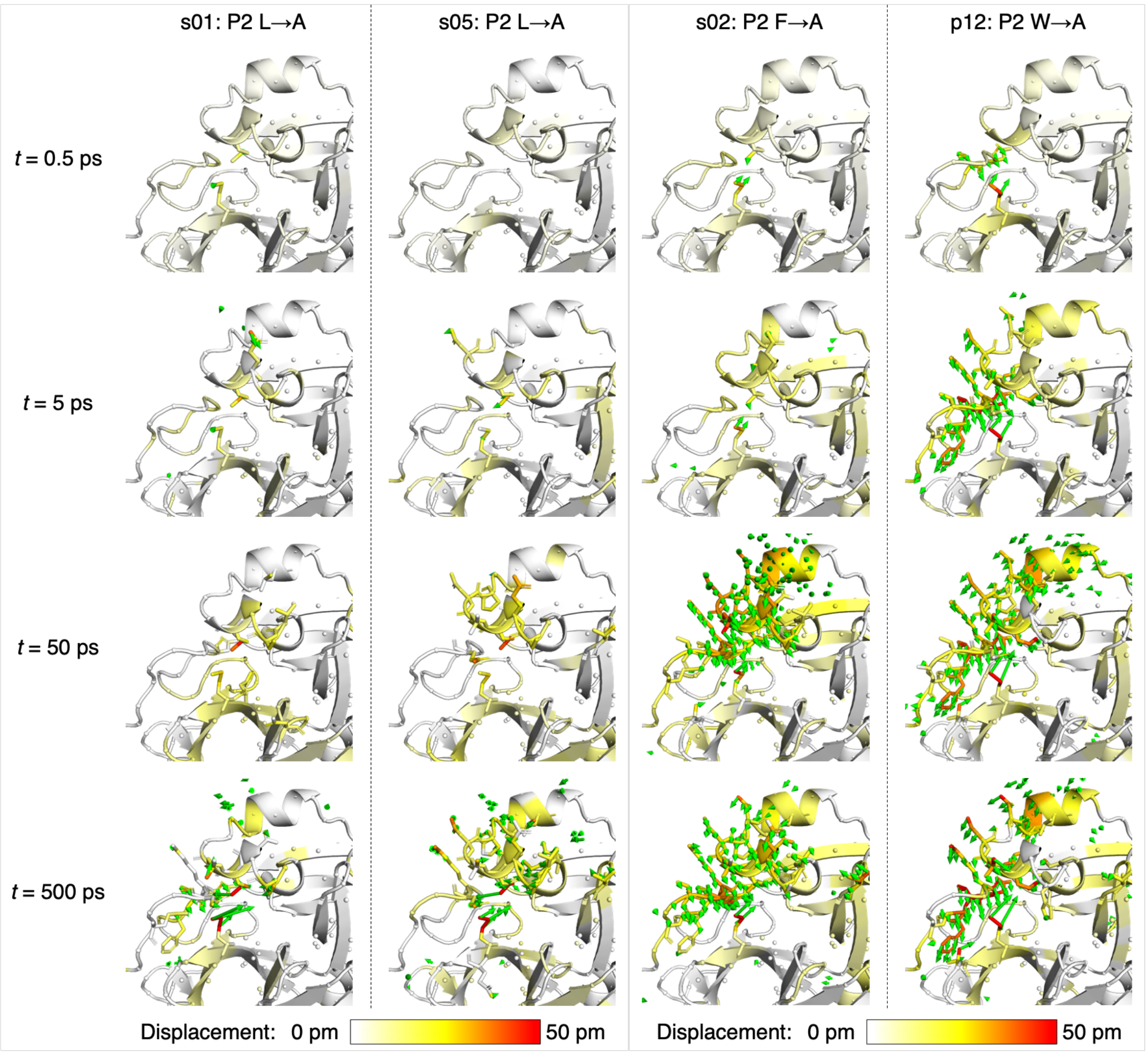

**Figure S3.6**: Views of the M$^{pro}$ response to the shown P2 substitutions in s01, s05, s02, and p12 from averaging non-hydrogen atom displacement vectors, shown using the structure prior to MD simulations, with a focus on the S2 subsite. Displacement magnitudes are represented on a white-yellow-red scale.[7] Significant vectors with length ≥20 pm are displayed as green arrows with a scale-up factor of 5.[8] Residues that show such significant displacements and that are within 10 Å of the substrate P2 residue are shown as sticks.

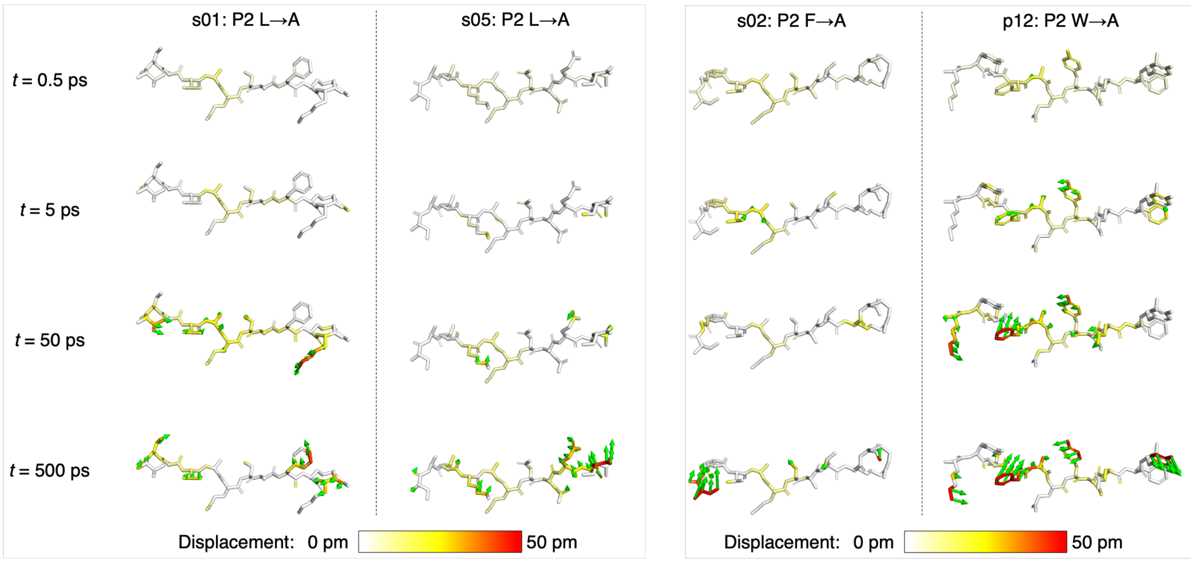

**Figure S3.7**: Views of the peptide response to the shown P2 substitutions in s01, s05, s02, and p12 from averaging non-hydrogen atom displacement vectors, shown using the structure prior to MD simulations. Displacement magnitudes are represented on a white-yellow-red scale.[7] Significant vectors with length ≥20 pm are displayed as green arrows with a scale-up factor of 5.[8]

### Section S4A: Substitution of P3 residue

**Table S4A.1**: The average differences in the five active-site distances ($\Delta d = d_{neq} - d_{eqm}$) following the shown P3 substitutions in s01 and s05. Positive and negative differences with magnitudes > 0.1 Å are in green and red, respectively.

| Active Site | s01: P3 V->A | | | | s05: P3 K->A | | |
|---|---|---|---|---|---|---|---|
| **Deprot** | t (ps) | Av Diff (Å) | SEM (Å) | | t (ps) | Av Diff (Å) | SEM (Å) |
| | 0.05 | 0.00 | 0.00 | | 0.05 | 0.02 | 0.00 |
| | 0.1 | 0.00 | 0.00 | | 0.1 | 0.00 | 0.00 |
| | 0.25 | 0.00 | 0.00 | | 0.25 | 0.01 | 0.01 |
| | 0.5 | 0.00 | 0.00 | | 0.5 | 0.02 | 0.02 |
| | 1 | -0.03 | 0.03 | | 1 | -0.02 | 0.04 |
| | 2 | -0.07 | 0.04 | | 2 | -0.01 | 0.06 |
| | 3 | 0.06 | 0.05 | | 3 | 0.03 | 0.07 |
| | 5 | -0.04 | 0.05 | | 5 | 0.05 | 0.07 |
| | 10 | 0.07 | 0.06 | | 10 | 0.07 | 0.07 |
| | 20 | 0.06 | 0.06 | | 20 | -0.04 | 0.07 |
| | 30 | 0.03 | 0.07 | | 30 | 0.04 | 0.08 |
| | 50 | 0.01 | 0.07 | | 50 | -0.06 | 0.07 |
| | 100 | 0.01 | 0.06 | | 100 | 0.08 | 0.08 |
| | 300 | -0.09 | 0.07 | | 300 | 0.12 | 0.08 |
| | 500 | 0.01 | 0.07 | | 500 | 0.11 | 0.07 |
| | 1000 | 0.15 | 0.09 | | 1000 | 0.04 | 0.07 |
| **Nuc** | t (ps) | Av Diff (Å) | SEM (Å) | | t (ps) | Av Diff (Å) | SEM (Å) |
| | 0.05 | 0.00 | 0.00 | | 0.05 | 0.01 | 0.00 |
| | 0.1 | 0.00 | 0.00 | | 0.1 | 0.01 | 0.00 |
| | 0.25 | 0.00 | 0.00 | | 0.25 | 0.02 | 0.00 |
| | 0.5 | 0.00 | 0.00 | | 0.5 | 0.02 | 0.01 |
| | 1 | -0.01 | 0.01 | | 1 | 0.01 | 0.02 |
| | 2 | -0.01 | 0.02 | | 2 | -0.02 | 0.02 |
| | 3 | -0.05 | 0.02 | | 3 | 0.02 | 0.02 |
| | 5 | -0.01 | 0.02 | | 5 | -0.02 | 0.02 |
| | 10 | -0.02 | 0.02 | | 10 | 0.01 | 0.02 |
| | 20 | 0.04 | 0.03 | | 20 | -0.03 | 0.03 |
| | 30 | -0.04 | 0.03 | | 30 | 0.03 | 0.03 |
| | 50 | -0.01 | 0.04 | | 50 | -0.02 | 0.04 |
| | 100 | -0.03 | 0.04 | | 100 | -0.03 | 0.04 |
| | 300 | -0.09 | 0.05 | | 300 | -0.04 | 0.05 |
| | 500 | 0.10 | 0.04 | | 500 | 0.03 | 0.05 |
| | 1000 | 0.00 | 0.05 | | 1000 | 0.07 | 0.05 |
| **OH145** | t (ps) | Av Diff (Å) | SEM (Å) | | t (ps) | Av Diff (Å) | SEM (Å) |
| | 0.05 | 0.00 | 0.00 | | 0.05 | -0.02 | 0.00 |
| | 0.1 | 0.00 | 0.00 | | 0.1 | -0.05 | 0.00 |
| | 0.25 | -0.01 | 0.00 | | 0.25 | -0.04 | 0.01 |
| | 0.5 | -0.01 | 0.01 | | 0.5 | -0.02 | 0.01 |
| | 1 | -0.01 | 0.02 | | 1 | -0.01 | 0.02 |
| | 2 | 0.02 | 0.03 | | 2 | 0.00 | 0.03 |
| | 3 | 0.01 | 0.03 | | 3 | -0.03 | 0.03 |
| | 5 | 0.04 | 0.03 | | 5 | -0.04 | 0.02 |
| | 10 | 0.00 | 0.03 | | 10 | -0.02 | 0.03 |
| | 20 | -0.04 | 0.03 | | 20 | 0.00 | 0.02 |
| | 30 | 0.01 | 0.03 | | 30 | 0.00 | 0.03 |
| | 50 | -0.05 | 0.03 | | 50 | -0.01 | 0.02 |
| | 100 | -0.05 | 0.04 | | 100 | -0.04 | 0.03 |
| | 300 | -0.04 | 0.03 | | 300 | 0.03 | 0.03 |
| | 500 | -0.07 | 0.03 | | 500 | -0.02 | 0.03 |
| | 1000 | 0.01 | 0.03 | | 1000 | 0.01 | 0.03 |
| **OH144** | t (ps) | Av Diff (Å) | SEM (Å) | | t (ps) | Av Diff (Å) | SEM (Å) |
| | 0.05 | 0.00 | 0.00 | | 0.05 | -0.01 | 0.00 |
| | 0.1 | 0.00 | 0.00 | | 0.1 | -0.02 | 0.00 |
| | 0.25 | -0.01 | 0.00 | | 0.25 | -0.02 | 0.01 |
| | 0.5 | -0.01 | 0.01 | | 0.5 | 0.00 | 0.01 |
| | 1 | 0.01 | 0.02 | | 1 | -0.02 | 0.02 |
| | 2 | 0.02 | 0.03 | | 2 | -0.01 | 0.03 |
| | 3 | 0.02 | 0.04 | | 3 | 0.00 | 0.03 |
| | 5 | 0.05 | 0.04 | | 5 | 0.03 | 0.04 |
| | 10 | 0.08 | 0.03 | | 10 | 0.04 | 0.03 |
| | 20 | -0.07 | 0.04 | | 20 | 0.03 | 0.03 |
| | 30 | 0.04 | 0.04 | | 30 | 0.03 | 0.03 |
| | 50 | -0.03 | 0.04 | | 50 | 0.00 | 0.04 |
| | 100 | -0.04 | 0.04 | | 100 | 0.02 | 0.03 |
| | 300 | -0.03 | 0.04 | | 300 | -0.03 | 0.04 |
| | 500 | -0.04 | 0.04 | | 500 | 0.03 | 0.04 |
| | 1000 | -0.01 | 0.04 | | 1000 | 0.02 | 0.04 |
| **OH143** | t (ps) | Av Diff (Å) | SEM (Å) | | t (ps) | Av Diff (Å) | SEM (Å) |
| | 0.05 | 0.00 | 0.00 | | 0.05 | 0.04 | 0.00 |
| | 0.1 | 0.00 | 0.00 | | 0.1 | 0.07 | 0.00 |
| | 0.25 | 0.00 | 0.00 | | 0.25 | 0.02 | 0.01 |
| | 0.5 | -0.01 | 0.01 | | 0.5 | 0.03 | 0.02 |
| | 1 | -0.03 | 0.02 | | 1 | 0.00 | 0.02 |
| | 2 | -0.01 | 0.03 | | 2 | 0.02 | 0.03 |
| | 3 | 0.00 | 0.03 | | 3 | -0.01 | 0.03 |
| | 5 | -0.04 | 0.03 | | 5 | -0.02 | 0.03 |
| | 10 | 0.00 | 0.03 | | 10 | 0.01 | 0.03 |
| | 20 | 0.05 | 0.03 | | 20 | 0.02 | 0.03 |
| | 30 | 0.01 | 0.03 | | 30 | 0.05 | 0.03 |
| | 50 | 0.00 | 0.03 | | 50 | 0.01 | 0.03 |
| | 100 | 0.08 | 0.03 | | 100 | 0.06 | 0.03 |
| | 300 | -0.03 | 0.03 | | 300 | 0.00 | 0.03 |
| | 500 | -0.02 | 0.03 | | 500 | 0.01 | 0.03 |
| | 1000 | 0.03 | 0.03 | | 1000 | 0.03 | 0.03 |

**Table S4A.2**: The average differences in the donor-acceptor distances of HBs 1-5 following the shown P3 substitutions in s01 and s05. Positive and negative differences with magnitudes > 0.1 Å are in green and red, respectively.

| HB1-5 | s01: P3 V->A | | | | s05: P3 K->A | | |
|---|---|---|---|---|---|---|---|
| **HB1** | t (ps) | Av Diff (Å) | SEM (Å) | | t (ps) | Av Diff (Å) | SEM (Å) |
| | 0.05 | 0.01 | 0.00 | | 0.05 | 0.01 | 0.00 |
| | 0.1 | 0.01 | 0.00 | | 0.1 | 0.02 | 0.00 |
| | 0.25 | 0.00 | 0.00 | | 0.25 | 0.02 | 0.01 |
| | 0.5 | -0.01 | 0.01 | | 0.5 | 0.01 | 0.01 |
| | 1 | -0.02 | 0.01 | | 1 | 0.01 | 0.02 |
| | 2 | -0.01 | 0.02 | | 2 | -0.01 | 0.02 |
| | 3 | 0.00 | 0.02 | | 3 | 0.02 | 0.03 |
| | 5 | -0.01 | 0.02 | | 5 | 0.02 | 0.03 |
| | 10 | 0.03 | 0.02 | | 10 | -0.03 | 0.03 |
| | 20 | 0.00 | 0.02 | | 20 | -0.03 | 0.03 |
| | 30 | 0.03 | 0.03 | | 30 | 0.01 | 0.03 |
| | 50 | 0.04 | 0.03 | | 50 | 0.00 | 0.03 |
| | 100 | 0.04 | 0.03 | | 100 | 0.02 | 0.04 |
| | 300 | 0.08 | 0.04 | | 300 | 0.00 | 0.04 |
| | 500 | 0.02 | 0.03 | | 500 | -0.02 | 0.04 |
| | 1000 | 0.01 | 0.04 | | 1000 | 0.03 | 0.06 |
| **HB2** | t (ps) | Av Diff (Å) | SEM (Å) | | t (ps) | Av Diff (Å) | SEM (Å) |
| | 0.05 | 0.00 | 0.00 | | 0.05 | 0.02 | 0.00 |
| | 0.1 | 0.02 | 0.00 | | 0.1 | 0.01 | 0.00 |
| | 0.25 | -0.01 | 0.01 | | 0.25 | 0.05 | 0.01 |
| | 0.5 | -0.03 | 0.01 | | 0.5 | 0.02 | 0.01 |
| | 1 | -0.02 | 0.01 | | 1 | 0.03 | 0.02 |
| | 2 | -0.06 | 0.02 | | 2 | 0.01 | 0.02 |
| | 3 | -0.03 | 0.02 | | 3 | -0.01 | 0.02 |
| | 5 | -0.05 | 0.02 | | 5 | -0.02 | 0.02 |
| | 10 | -0.01 | 0.02 | | 10 | 0.01 | 0.02 |
| | 20 | -0.02 | 0.02 | | 20 | -0.02 | 0.02 |
| | 30 | -0.05 | 0.02 | | 30 | 0.00 | 0.02 |
| | 50 | -0.05 | 0.02 | | 50 | 0.02 | 0.02 |
| | 100 | 0.00 | 0.02 | | 100 | 0.00 | 0.02 |
| | 300 | -0.03 | 0.02 | | 300 | 0.00 | 0.02 |
| | 500 | -0.02 | 0.02 | | 500 | -0.01 | 0.02 |
| | 1000 | -0.06 | 0.02 | | 1000 | 0.00 | 0.02 |
| **HB3** | t (ps) | Av Diff (Å) | SEM (Å) | | t (ps) | Av Diff (Å) | SEM (Å) |
| | 0.05 | 0.01 | 0.00 | | 0.05 | -0.01 | 0.00 |
| | 0.1 | 0.00 | 0.00 | | 0.1 | 0.00 | 0.00 |
| | 0.25 | -0.01 | 0.01 | | 0.25 | 0.02 | 0.01 |
| | 0.5 | -0.02 | 0.01 | | 0.5 | 0.00 | 0.01 |
| | 1 | -0.01 | 0.01 | | 1 | 0.00 | 0.01 |
| | 2 | -0.04 | 0.01 | | 2 | -0.02 | 0.02 |
| | 3 | 0.00 | 0.01 | | 3 | -0.01 | 0.01 |
| | 5 | -0.05 | 0.02 | | 5 | -0.01 | 0.02 |
| | 10 | 0.01 | 0.01 | | 10 | 0.00 | 0.01 |
| | 20 | -0.03 | 0.01 | | 20 | -0.02 | 0.01 |
| | 30 | 0.00 | 0.01 | | 30 | 0.03 | 0.01 |
| | 50 | -0.02 | 0.01 | | 50 | 0.00 | 0.02 |
| | 100 | 0.01 | 0.01 | | 100 | 0.00 | 0.02 |
| | 300 | -0.03 | 0.01 | | 300 | 0.02 | 0.02 |
| | 500 | -0.02 | 0.01 | | 500 | 0.00 | 0.01 |
| | 1000 | -0.02 | 0.01 | | 1000 | 0.01 | 0.02 |
| **HB4** | t (ps) | Av Diff (Å) | SEM (Å) | | t (ps) | Av Diff (Å) | SEM (Å) |
| | 0.05 | 0.00 | 0.00 | | 0.05 | -0.01 | 0.00 |
| | 0.1 | -0.01 | 0.00 | | 0.1 | -0.01 | 0.01 |
| | 0.25 | -0.01 | 0.01 | | 0.25 | 0.02 | 0.02 |
| | 0.5 | -0.02 | 0.01 | | 0.5 | 0.00 | 0.02 |
| | 1 | -0.02 | 0.02 | | 1 | 0.02 | 0.02 |
| | 2 | 0.02 | 0.03 | | 2 | -0.02 | 0.03 |
| | 3 | 0.03 | 0.04 | | 3 | 0.08 | 0.04 |
| | 5 | -0.03 | 0.04 | | 5 | 0.06 | 0.05 |
| | 10 | 0.01 | 0.04 | | 10 | -0.01 | 0.04 |
| | 20 | 0.08 | 0.05 | | 20 | 0.01 | 0.06 |
| | 30 | 0.02 | 0.06 | | 30 | 0.06 | 0.06 |
| | 50 | -0.03 | 0.07 | | 50 | 0.17 | 0.08 |
| | 100 | 0.03 | 0.09 | | 100 | 0.20 | 0.11 |
| | 300 | -0.05 | 0.09 | | 300 | 0.51 | 0.14 |
| | 500 | -0.01 | 0.13 | | 500 | 0.74 | 0.13 |
| | 1000 | -0.07 | 0.13 | | 1000 | 0.63 | 0.15 |
| **HB5** | t (ps) | Av Diff (Å) | SEM (Å) | | t (ps) | Av Diff (Å) | SEM (Å) |
| | 0.05 | 0.00 | 0.00 | | 0.05 | 0.00 | 0.00 |
| | 0.1 | 0.00 | 0.00 | | 0.1 | 0.01 | 0.00 |
| | 0.25 | 0.02 | 0.00 | | 0.25 | 0.02 | 0.01 |
| | 0.5 | 0.02 | 0.01 | | 0.5 | 0.03 | 0.01 |
| | 1 | 0.00 | 0.01 | | 1 | 0.01 | 0.02 |
| | 2 | -0.02 | 0.03 | | 2 | 0.08 | 0.02 |
| | 3 | 0.02 | 0.03 | | 3 | 0.03 | 0.03 |
| | 5 | 0.01 | 0.03 | | 5 | 0.04 | 0.02 |
| | 10 | 0.02 | 0.03 | | 10 | 0.03 | 0.03 |
| | 20 | 0.02 | 0.03 | | 20 | -0.03 | 0.03 |
| | 30 | 0.03 | 0.03 | | 30 | 0.00 | 0.02 |
| | 50 | -0.01 | 0.03 | | 50 | 0.07 | 0.03 |
| | 100 | 0.08 | 0.03 | | 100 | 0.00 | 0.03 |
| | 300 | 0.01 | 0.03 | | 300 | 0.06 | 0.03 |
| | 500 | -0.01 | 0.03 | | 500 | 0.01 | 0.03 |
| | 1000 | 0.09 | 0.04 | | 1000 | 0.02 | 0.03 |

**Table S4A.3**: The average differences in the donor-acceptor distances of HBs 8-12 following the shown P3 substitutions in s01 and s05. Positive and negative differences with magnitudes > 0.1 Å are in green and red, respectively.

| HB8-12 | s01: P3 V->A | | | | s05: P3 K->A | | |
|---|---|---|---|---|---|---|---|
| **HB8** | t (ps) | Av Diff (Å) | SEM (Å) | | t (ps) | Av Diff (Å) | SEM (Å) |
| | 0.05 | 0.00 | 0.00 | | 0.05 | -0.02 | 0.00 |
| | 0.1 | 0.00 | 0.00 | | 0.1 | -0.05 | 0.00 |
| | 0.25 | -0.01 | 0.00 | | 0.25 | -0.03 | 0.01 |
| | 0.5 | -0.01 | 0.01 | | 0.5 | -0.02 | 0.01 |
| | 1 | -0.01 | 0.02 | | 1 | -0.01 | 0.02 |
| | 2 | 0.02 | 0.03 | | 2 | 0.01 | 0.03 |
| | 3 | 0.01 | 0.03 | | 3 | -0.03 | 0.03 |
| | 5 | 0.04 | 0.03 | | 5 | -0.04 | 0.02 |
| | 10 | 0.00 | 0.03 | | 10 | -0.03 | 0.03 |
| | 20 | -0.04 | 0.03 | | 20 | 0.00 | 0.02 |
| | 30 | 0.00 | 0.03 | | 30 | 0.00 | 0.03 |
| | 50 | -0.05 | 0.03 | | 50 | -0.01 | 0.02 |
| | 100 | -0.05 | 0.04 | | 100 | -0.04 | 0.03 |
| | 300 | -0.05 | 0.03 | | 300 | 0.04 | 0.03 |
| | 500 | -0.08 | 0.03 | | 500 | -0.03 | 0.03 |
| | 1000 | 0.01 | 0.03 | | 1000 | 0.01 | 0.03 |
| **HB9** | t (ps) | Av Diff (Å) | SEM (Å) | | t (ps) | Av Diff (Å) | SEM (Å) |
| | 0.05 | 0.00 | 0.00 | | 0.05 | 0.02 | 0.00 |
| | 0.1 | 0.00 | 0.00 | | 0.1 | 0.04 | 0.00 |
| | 0.25 | 0.00 | 0.00 | | 0.25 | 0.01 | 0.01 |
| | 0.5 | 0.00 | 0.01 | | 0.5 | 0.03 | 0.01 |
| | 1 | -0.02 | 0.01 | | 1 | 0.00 | 0.01 |
| | 2 | 0.01 | 0.02 | | 2 | 0.02 | 0.02 |
| | 3 | 0.00 | 0.02 | | 3 | 0.01 | 0.02 |
| | 5 | -0.02 | 0.01 | | 5 | -0.01 | 0.02 |
| | 10 | 0.01 | 0.02 | | 10 | 0.01 | 0.02 |
| | 20 | 0.01 | 0.02 | | 20 | 0.01 | 0.02 |
| | 30 | 0.01 | 0.02 | | 30 | 0.02 | 0.02 |
| | 50 | 0.00 | 0.02 | | 50 | 0.01 | 0.02 |
| | 100 | 0.04 | 0.02 | | 100 | 0.04 | 0.02 |
| | 300 | -0.01 | 0.02 | | 300 | 0.01 | 0.02 |
| | 500 | -0.02 | 0.02 | | 500 | 0.00 | 0.02 |
| | 1000 | 0.01 | 0.02 | | 1000 | 0.01 | 0.02 |
| **HB10** | t (ps) | Av Diff (Å) | SEM (Å) | | t (ps) | Av Diff (Å) | SEM (Å) |
| | 0.05 | 0.00 | 0.00 | | 0.05 | 0.00 | 0.00 |
| | 0.1 | 0.00 | 0.00 | | 0.1 | 0.00 | 0.00 |
| | 0.25 | 0.00 | 0.00 | | 0.25 | 0.00 | 0.00 |
| | 0.5 | 0.01 | 0.00 | | 0.5 | 0.02 | 0.01 |
| | 1 | 0.00 | 0.01 | | 1 | 0.02 | 0.01 |
| | 2 | 0.02 | 0.02 | | 2 | 0.01 | 0.02 |
| | 3 | 0.01 | 0.02 | | 3 | 0.03 | 0.02 |
| | 5 | -0.01 | 0.02 | | 5 | 0.01 | 0.02 |
| | 10 | -0.01 | 0.02 | | 10 | 0.01 | 0.02 |
| | 20 | -0.03 | 0.02 | | 20 | -0.01 | 0.02 |
| | 30 | 0.00 | 0.02 | | 30 | 0.02 | 0.02 |
| | 50 | 0.00 | 0.02 | | 50 | 0.00 | 0.02 |
| | 100 | -0.01 | 0.02 | | 100 | -0.01 | 0.02 |
| | 300 | 0.00 | 0.02 | | 300 | 0.05 | 0.02 |
| | 500 | -0.03 | 0.02 | | 500 | -0.02 | 0.02 |
| | 1000 | -0.01 | 0.02 | | 1000 | 0.02 | 0.02 |
| **HB11** | t (ps) | Av Diff (Å) | SEM (Å) | | t (ps) | Av Diff (Å) | SEM (Å) |
| | 0.05 | 0.00 | 0.00 | | 0.05 | -0.02 | 0.00 |
| | 0.1 | 0.00 | 0.00 | | 0.1 | -0.03 | 0.00 |
| | 0.25 | 0.00 | 0.00 | | 0.25 | -0.01 | 0.00 |
| | 0.5 | 0.00 | 0.00 | | 0.5 | -0.01 | 0.01 |
| | 1 | 0.01 | 0.01 | | 1 | -0.01 | 0.01 |
| | 2 | 0.00 | 0.02 | | 2 | -0.03 | 0.01 |
| | 3 | 0.01 | 0.02 | | 3 | 0.02 | 0.02 |
| | 5 | -0.03 | 0.02 | | 5 | 0.02 | 0.02 |
| | 10 | -0.02 | 0.02 | | 10 | 0.01 | 0.02 |
| | 20 | 0.03 | 0.03 | | 20 | 0.00 | 0.02 |
| | 30 | -0.02 | 0.02 | | 30 | -0.01 | 0.02 |
| | 50 | -0.03 | 0.02 | | 50 | -0.01 | 0.01 |
| | 100 | 0.00 | 0.02 | | 100 | 0.01 | 0.01 |
| | 300 | -0.01 | 0.02 | | 300 | -0.01 | 0.02 |
| | 500 | -0.01 | 0.02 | | 500 | -0.03 | 0.02 |
| | 1000 | 0.01 | 0.02 | | 1000 | 0.01 | 0.01 |
| **HB12** | t (ps) | Av Diff (Å) | SEM (Å) | | t (ps) | Av Diff (Å) | SEM (Å) |
| | 0.05 | 0.00 | 0.00 | | 0.05 | 0.00 | 0.00 |
| | 0.1 | 0.00 | 0.00 | | 0.1 | 0.00 | 0.00 |
| | 0.25 | 0.00 | 0.00 | | 0.25 | 0.01 | 0.00 |
| | 0.5 | 0.00 | 0.00 | | 0.5 | 0.00 | 0.01 |
| | 1 | 0.00 | 0.02 | | 1 | -0.01 | 0.02 |
| | 2 | 0.01 | 0.03 | | 2 | -0.05 | 0.02 |
| | 3 | 0.03 | 0.03 | | 3 | 0.00 | 0.02 |
| | 5 | -0.04 | 0.03 | | 5 | 0.01 | 0.02 |
| | 10 | -0.04 | 0.03 | | 10 | -0.04 | 0.03 |
| | 20 | -0.03 | 0.03 | | 20 | 0.00 | 0.02 |
| | 30 | -0.02 | 0.03 | | 30 | 0.00 | 0.03 |
| | 50 | -0.05 | 0.03 | | 50 | 0.03 | 0.03 |
| | 100 | 0.02 | 0.04 | | 100 | -0.02 | 0.03 |
| | 300 | -0.03 | 0.05 | | 300 | 0.00 | 0.03 |
| | 500 | 0.00 | 0.04 | | 500 | -0.03 | 0.03 |
| | 1000 | -0.05 | 0.05 | | 1000 | 0.03 | 0.04 |

**Table S4A.4**: The ten most perturbed Cα atoms in the M^pro-peptide complex in terms of magnitude of the average displacement vector $v$, at selected time points following the P3 substitutions to Ala from Val and Lys in s01 and s05, respectively. Only displacements ≥ 0.01 Å are shown.

| s01 t(ps) | CA Rank | Atom | Chain | v (Å) | SEM (Å) |
|---|---|---|---|---|---|
| 0.05 | 1 | 4ALA_CA | C | 0.03 | 0.00 |
| | 2 | 3ALA_CA | C | 0.02 | 0.00 |
| | 3 | 2SER_CA | C | 0.01 | 0.00 |
| | 4 | 5LEU_CA | C | 0.01 | 0.00 |
| | 5 | | | | |
| | 6 | | | | |
| | 7 | | | | |
| | 8 | | | | |
| | 9 | | | | |
| | 10 | | | | |
| 0.5 | 1 | 4ALA_CA | C | 0.08 | 0.01 |
| | 2 | 5LEU_CA | C | 0.05 | 0.01 |
| | 3 | 3ALA_CA | C | 0.03 | 0.01 |
| | 4 | 45THR_CA | A | 0.02 | 0.00 |
| | 5 | 46SER_CA | A | 0.02 | 0.00 |
| | 6 | 138GLY_CA | A | 0.02 | 0.00 |
| | 7 | 167LEU_CA | A | 0.02 | 0.01 |
| | 8 | 168PRO_CA | A | 0.02 | 0.00 |
| | 9 | 169THR_CA | A | 0.02 | 0.00 |
| | 10 | 170GLY_CA | A | 0.02 | 0.00 |
| 5 | 1 | 94ALA_CA | A | 0.15 | 0.04 |
| | 2 | 93THR_CA | A | 0.14 | 0.04 |
| | 3 | 301SER_CA | B | 0.12 | 0.03 |
| | 4 | 4ALA_CA | C | 0.12 | 0.03 |
| | 5 | 24THR_CA | B | 0.11 | 0.05 |
| | 6 | 283GLY_CA | B | 0.11 | 0.04 |
| | 7 | 293PRO_CA | B | 0.11 | 0.03 |
| | 8 | 300CYS_CA | B | 0.11 | 0.03 |
| | 9 | 195GLY_CA | A | 0.10 | 0.04 |
| | 10 | 283GLY_CA | A | 0.10 | 0.04 |
| 50 | 1 | 72ASN_CA | B | 0.26 | 0.09 |
| | 2 | 255ALA_CA | B | 0.22 | 0.06 |
| | 3 | 1THR_CA | C | 0.22 | 0.09 |
| | 4 | 191ALA_CA | B | 0.21 | 0.08 |
| | 5 | 258GLY_CA | B | 0.20 | 0.06 |
| | 6 | 194ALA_CA | B | 0.19 | 0.07 |
| | 7 | 195GLY_CA | B | 0.19 | 0.06 |
| | 8 | 196THR_CA | B | 0.19 | 0.06 |
| | 9 | 246HIE_CA | B | 0.19 | 0.05 |
| | 10 | 248ASP_CA | B | 0.19 | 0.06 |
| 500 | 1 | 72ASN_CA | A | 0.23 | 0.10 |
| | 2 | 47GLU_CA | A | 0.20 | 0.08 |
| | 3 | 303VAL_CA | B | 0.20 | 0.06 |
| | 4 | 72ASN_CA | B | 0.19 | 0.10 |
| | 5 | 73VAL_CA | A | 0.18 | 0.07 |
| | 6 | 92ASP_CA | A | 0.17 | 0.08 |
| | 7 | 260ALA_CA | A | 0.17 | 0.07 |
| | 8 | 251GLY_CA | B | 0.17 | 0.06 |
| | 9 | 274ASN_CA | B | 0.17 | 0.08 |
| | 10 | 275GLY_CA | B | 0.17 | 0.07 |
| 1000 | 1 | 277ASN_CA | A | 0.27 | 0.12 |
| | 2 | 277ASN_CA | B | 0.25 | 0.09 |
| | 3 | 256GLN_CA | B | 0.23 | 0.07 |
| | 4 | 255ALA_CA | B | 0.22 | 0.07 |
| | 5 | 51ASN_CA | A | 0.20 | 0.08 |
| | 6 | 52PRO_CA | A | 0.20 | 0.08 |
| | 7 | 55GLU_CA | A | 0.19 | 0.07 |
| | 8 | 283GLY_CA | A | 0.19 | 0.06 |
| | 9 | 50LEU_CA | B | 0.19 | 0.09 |
| | 10 | 224THR_CA | B | 0.19 | 0.09 |

| s05 t(ps) | CA Rank | Atom | Chain | v (Å) | SEM (Å) |
|---|---|---|---|---|---|
| 0.05 | 1 | 4ALA_CA | C | 0.03 | 0.00 |
| | 2 | 46SER_CA | A | 0.01 | 0.00 |
| | 3 | 141LEU_CA | A | 0.01 | 0.00 |
| | 4 | 143GLY_CA | A | 0.01 | 0.00 |
| | 5 | 170GLY_CA | A | 0.01 | 0.00 |
| | 6 | 188ARG_CA | A | 0.01 | 0.00 |
| | 7 | 189GLN_CA | A | 0.01 | 0.00 |
| | 8 | 190THR_CA | A | 0.01 | 0.00 |
| | 9 | 1SER_CA | B | 0.01 | 0.00 |
| | 10 | 2GLY_CA | B | 0.01 | 0.00 |
| 0.5 | 1 | 7ASN_CA | C | 0.06 | 0.01 |
| | 2 | 8ASN_CA | C | 0.05 | 0.01 |
| | 3 | 9GLU_CA | C | 0.05 | 0.01 |
| | 4 | 10LEU_CA | C | 0.05 | 0.01 |
| | 5 | 45THR_CA | A | 0.04 | 0.01 |
| | 6 | 170GLY_CA | A | 0.04 | 0.01 |
| | 7 | 5LEU_CA | C | 0.04 | 0.01 |
| | 8 | 23GLY_CA | A | 0.03 | 0.01 |
| | 9 | 24THR_CA | A | 0.03 | 0.01 |
| | 10 | 44CYS_CA | A | 0.03 | 0.01 |
| 5 | 1 | 50LEU_CA | B | 0.18 | 0.04 |
| | 2 | 46SER_CA | B | 0.16 | 0.05 |
| | 3 | 51ASN_CA | B | 0.16 | 0.04 |
| | 4 | 47GLU_CA | B | 0.15 | 0.05 |
| | 5 | 24THR_CA | B | 0.14 | 0.05 |
| | 6 | 189GLN_CA | B | 0.14 | 0.04 |
| | 7 | 71GLY_CA | A | 0.13 | 0.06 |
| | 8 | 45THR_CA | B | 0.13 | 0.04 |
| | 9 | 49MET_CA | B | 0.13 | 0.04 |
| | 10 | 64HIE_CA | B | 0.13 | 0.05 |
| 50 | 1 | 11SER_CA | C | 0.26 | 0.11 |
| | 2 | 251GLY_CA | A | 0.25 | 0.06 |
| | 3 | 252PRO_CA | A | 0.25 | 0.06 |
| | 4 | 253LEU_CA | A | 0.22 | 0.05 |
| | 5 | 24THR_CA | B | 0.22 | 0.06 |
| | 6 | 236LYS_CA | B | 0.20 | 0.07 |
| | 7 | 232LEU_CA | A | 0.19 | 0.06 |
| | 8 | 23GLY_CA | B | 0.19 | 0.05 |
| | 9 | 275GLY_CA | B | 0.19 | 0.07 |
| | 10 | 276MET_CA | B | 0.19 | 0.07 |
| 500 | 1 | 72ASN_CA | A | 0.36 | 0.11 |
| | 2 | 11SER_CA | C | 0.30 | 0.16 |
| | 3 | 55GLU_CA | A | 0.29 | 0.07 |
| | 4 | 93THR_CA | B | 0.26 | 0.09 |
| | 5 | 277ASN_CA | B | 0.25 | 0.11 |
| | 6 | 73VAL_CA | A | 0.23 | 0.09 |
| | 7 | 278GLY_CA | A | 0.23 | 0.09 |
| | 8 | 193ALA_CA | B | 0.22 | 0.07 |
| | 9 | 194ALA_CA | B | 0.22 | 0.07 |
| | 10 | 195GLY_CA | B | 0.22 | 0.08 |
| 1000 | 1 | 11SER_CA | C | 0.84 | 0.18 |
| | 2 | 10LEU_CA | C | 0.48 | 0.11 |
| | 3 | 278GLY_CA | A | 0.33 | 0.12 |
| | 4 | 9GLU_CA | C | 0.31 | 0.07 |
| | 5 | 55GLU_CA | B | 0.29 | 0.09 |
| | 6 | 56ASP_CA | B | 0.29 | 0.11 |
| | 7 | 306GLN_CA | A | 0.28 | 0.10 |
| | 8 | 46SER_CA | A | 0.23 | 0.07 |
| | 9 | 59ILE_CA | B | 0.23 | 0.10 |
| | 10 | 277ASN_CA | A | 0.22 | 0.11 |

**Table S4A.5**: The ten most perturbed non-hydrogen atoms in the M$^{pro}$-peptide complex in terms of magnitude of the average displacement vector *v*, at selected time points following the P3 substitutions to Ala from Val and Lys in s01 and s05, respectively.

| *s01 t(ps)* | *Non-H Rank* | *Atom* | *Chain* | *v (Å)* | *SEM (Å)* |
|---|---|---|---|---|---|
| **0.05** | 1 | 4ALA_CB | C | 0.09 | 0.01 |
| | 2 | 4ALA_N | C | 0.04 | 0.00 |
| | 3 | 3ALA_CB | C | 0.03 | 0.00 |
| | 4 | 4ALA_O | C | 0.03 | 0.00 |
| | 5 | 3ALA_C | C | 0.03 | 0.00 |
| | 6 | 4ALA_CA | C | 0.03 | 0.00 |
| | 7 | 3ALA_CA | C | 0.02 | 0.00 |
| | 8 | 166GLU_CB | A | 0.02 | 0.00 |
| | 9 | 5LEU_CA | C | 0.01 | 0.00 |
| | 10 | 3ALA_O | C | 0.01 | 0.00 |
| **0.5** | 1 | 4ALA_CB | C | 0.18 | 0.02 |
| | 2 | 189GLN_NE2 | A | 0.09 | 0.02 |
| | 3 | 4ALA_CA | C | 0.08 | 0.01 |
| | 4 | 5LEU_O | C | 0.07 | 0.02 |
| | 5 | 3ALA_O | C | 0.07 | 0.02 |
| | 6 | 166GLU_OE2 | A | 0.07 | 0.01 |
| | 7 | 5LEU_CB | C | 0.07 | 0.01 |
| | 8 | 5LEU_N | C | 0.06 | 0.01 |
| | 9 | 189GLN_CD | A | 0.06 | 0.01 |
| | 10 | 189GLN_OE1 | A | 0.06 | 0.02 |
| **5** | 1 | 232LEU_CD2 | A | 0.27 | 0.08 |
| | 2 | 232LEU_CD1 | A | 0.26 | 0.07 |
| | 3 | 10ARG_NH2 | C | 0.23 | 0.07 |
| | 4 | 76ARG_NH2 | A | 0.23 | 0.08 |
| | 5 | 222ARG_NH1 | B | 0.22 | 0.10 |
| | 6 | 53ASN_OD1 | A | 0.21 | 0.08 |
| | 7 | 236LYS_NZ | A | 0.21 | 0.11 |
| | 8 | 92ASP_OD2 | A | 0.21 | 0.06 |
| | 9 | 4ALA_CB | C | 0.21 | 0.04 |
| | 10 | 232LEU_CG | A | 0.20 | 0.06 |
| **50** | 1 | 72ASN_ND2 | B | 0.47 | 0.14 |
| | 2 | 11LYS_NZ | C | 0.40 | 0.18 |
| | 3 | 222ARG_NH2 | A | 0.39 | 0.19 |
| | 4 | 55GLU_OE1 | B | 0.38 | 0.14 |
| | 5 | 60ARG_NH1 | A | 0.37 | 0.12 |
| | 6 | 1THR_OG1 | C | 0.36 | 0.14 |
| | 7 | 248ASP_OD1 | B | 0.36 | 0.09 |
| | 8 | 72ASN_OD1 | A | 0.35 | 0.14 |
| | 9 | 222ARG_NH2 | B | 0.35 | 0.16 |
| | 10 | 72ASN_CG | B | 0.35 | 0.12 |
| **500** | 1 | 244GLN_OE1 | B | 0.60 | 0.16 |
| | 2 | 64HIE_NE2 | B | 0.52 | 0.15 |
| | 3 | 222ARG_NH1 | B | 0.49 | 0.25 |
| | 4 | 97LYS_NZ | B | 0.48 | 0.14 |
| | 5 | 72ASN_OD1 | B | 0.47 | 0.21 |
| | 6 | 244GLN_NE2 | A | 0.47 | 0.21 |
| | 7 | 9PHE_CZ | C | 0.47 | 0.16 |
| | 8 | 72ASN_ND2 | A | 0.46 | 0.20 |
| | 9 | 47GLU_OE2 | A | 0.45 | 0.21 |
| | 10 | 64HIE_CE1 | B | 0.44 | 0.15 |
| **1000** | 1 | 277ASN_OD1 | A | 0.62 | 0.20 |
| | 2 | 222ARG_NH2 | B | 0.58 | 0.25 |
| | 3 | 244GLN_NE2 | B | 0.56 | 0.23 |
| | 4 | 142ASN_ND2 | B | 0.53 | 0.14 |
| | 5 | 279ARG_NH1 | B | 0.52 | 0.20 |
| | 6 | 55GLU_OE1 | B | 0.52 | 0.16 |
| | 7 | 222ARG_NH1 | B | 0.51 | 0.22 |
| | 8 | 222ARG_CZ | B | 0.50 | 0.21 |
| | 9 | 306GLN_NE2 | A | 0.49 | 0.22 |
| | 10 | 55GLU_OE2 | B | 0.48 | 0.16 |

| *s05 t(ps)* | *Non-H Rank* | *Atom* | *Chain* | *v (Å)* | *SEM (Å)* |
|---|---|---|---|---|---|
| **0.05** | 1 | 7ASN_O | C | 0.05 | 0.00 |
| | 2 | 142ASN_OD1 | A | 0.05 | 0.00 |
| | 3 | 5LEU_O | C | 0.05 | 0.00 |
| | 4 | 7ASN_OD1 | C | 0.05 | 0.00 |
| | 5 | 4ALA_O | C | 0.04 | 0.00 |
| | 6 | 189GLN_OE1 | A | 0.03 | 0.00 |
| | 7 | 4ALA_N | C | 0.03 | 0.01 |
| | 8 | 4ALA_CB | C | 0.03 | 0.01 |
| | 9 | 166GLU_OE1 | A | 0.03 | 0.00 |
| | 10 | 9GLU_OE1 | C | 0.03 | 0.00 |
| **0.5** | 1 | 142ASN_ND2 | A | 0.21 | 0.04 |
| | 2 | 9GLU_OE1 | C | 0.18 | 0.03 |
| | 3 | 9GLU_OE2 | C | 0.17 | 0.03 |
| | 4 | 142ASN_OD1 | A | 0.17 | 0.03 |
| | 5 | 9GLU_CD | C | 0.15 | 0.02 |
| | 6 | 10LEU_O | C | 0.10 | 0.02 |
| | 7 | 47GLU_OE2 | A | 0.10 | 0.02 |
| | 8 | 47GLU_OE1 | A | 0.09 | 0.02 |
| | 9 | 9GLU_CG | C | 0.09 | 0.02 |
| | 10 | 142ASN_CG | A | 0.09 | 0.02 |
| **5** | 1 | 142ASN_ND2 | A | 0.33 | 0.07 |
| | 2 | 279ARG_NH1 | B | 0.30 | 0.07 |
| | 3 | 107GLN_NE2 | B | 0.28 | 0.08 |
| | 4 | 189GLN_NE2 | B | 0.27 | 0.07 |
| | 5 | 279ARG_CZ | B | 0.26 | 0.06 |
| | 6 | 107GLN_NE2 | A | 0.26 | 0.08 |
| | 7 | 279ARG_NH2 | B | 0.26 | 0.07 |
| | 8 | 24THR_CG2 | B | 0.25 | 0.09 |
| | 9 | 47GLU_OE1 | B | 0.25 | 0.10 |
| | 10 | 244GLN_NE2 | B | 0.24 | 0.09 |
| **50** | 1 | 142ASN_OD1 | A | 0.57 | 0.08 |
| | 2 | 236LYS_NZ | B | 0.48 | 0.19 |
| | 3 | 142ASN_ND2 | A | 0.44 | 0.08 |
| | 4 | 236LYS_CE | B | 0.44 | 0.16 |
| | 5 | 244GLN_NE2 | B | 0.43 | 0.14 |
| | 6 | 11SER_OC2 | C | 0.43 | 0.16 |
| | 7 | 279ARG_NH2 | B | 0.41 | 0.12 |
| | 8 | 279ARG_NH1 | A | 0.41 | 0.12 |
| | 9 | 306GLN_OE1 | A | 0.39 | 0.12 |
| | 10 | 236LYS_CD | B | 0.35 | 0.13 |
| **500** | 1 | 189GLN_OE1 | A | 0.75 | 0.11 |
| | 2 | 256GLN_NE2 | A | 0.68 | 0.14 |
| | 3 | 72ASN_ND2 | A | 0.68 | 0.24 |
| | 4 | 55GLU_OE2 | A | 0.59 | 0.17 |
| | 5 | 76ARG_NH1 | B | 0.56 | 0.17 |
| | 6 | 11SER_OC1 | C | 0.55 | 0.24 |
| | 7 | 46SER_OG | A | 0.55 | 0.12 |
| | 8 | 72ASN_CG | A | 0.54 | 0.20 |
| | 9 | 9GLU_OE1 | C | 0.53 | 0.14 |
| | 10 | 47GLU_OE2 | A | 0.52 | 0.19 |
| **1000** | 1 | 11SER_OG | C | 1.14 | 0.24 |
| | 2 | 11SER_CB | C | 1.08 | 0.23 |
| | 3 | 11SER_OC2 | C | 0.95 | 0.25 |
| | 4 | 11SER_CA | C | 0.84 | 0.19 |
| | 5 | 9GLU_OE2 | C | 0.81 | 0.14 |
| | 6 | 11SER_C | C | 0.81 | 0.22 |
| | 7 | 11SER_N | C | 0.71 | 0.15 |
| | 8 | 47GLU_OE1 | A | 0.68 | 0.19 |
| | 9 | 189GLN_OE1 | A | 0.68 | 0.12 |
| | 10 | 11SER_OC1 | C | 0.66 | 0.25 |

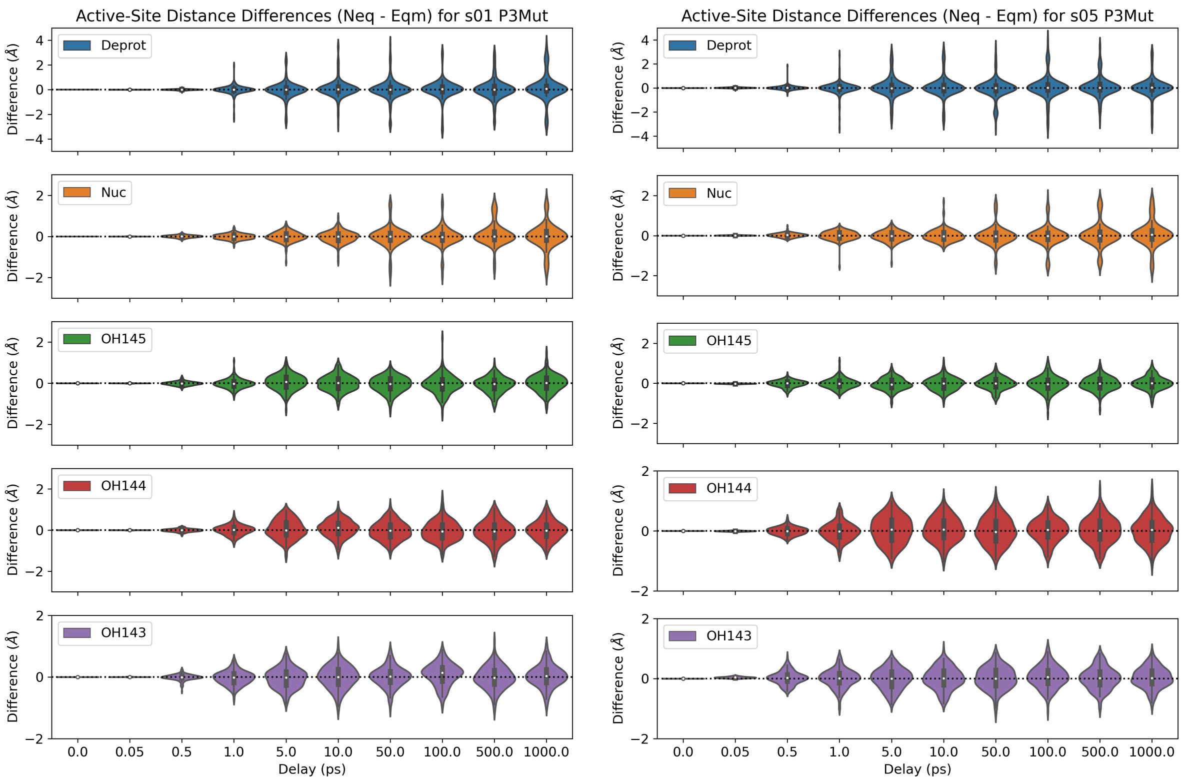

**Figure S4A.1**: Distributions of the distance differences in the five active-site distances ($\Delta d = d_{\mathrm{neq}} - d_{\mathrm{eqm}}$) following the P3 substitutions to Ala from Val and Lys in s01 and s05, respectively.

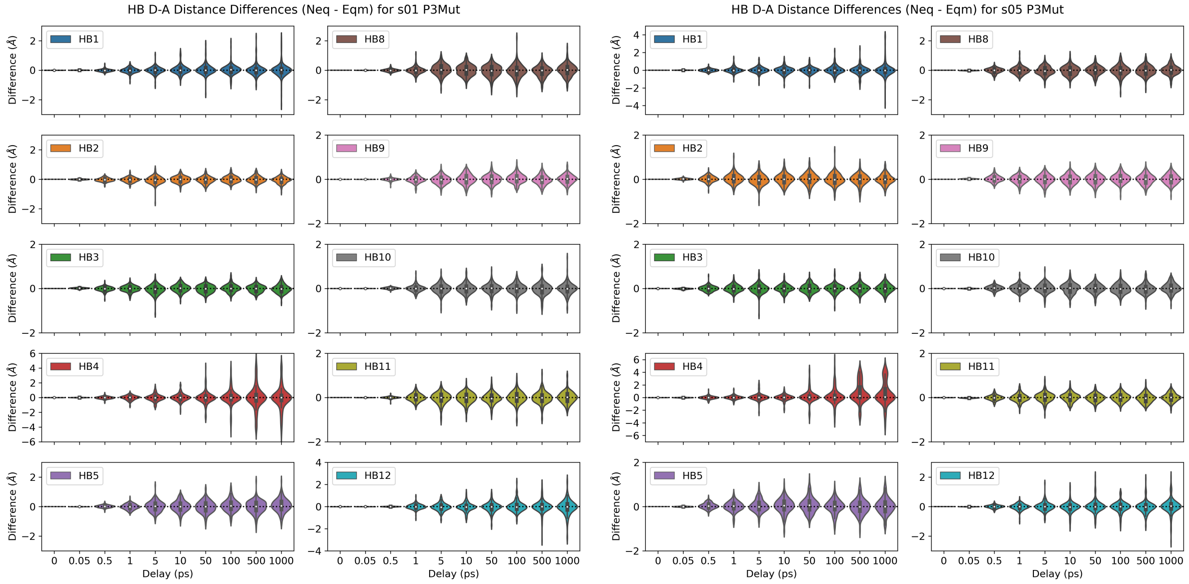

**Figure S4A.2**: Distributions of the distance differences in the ten donor-acceptor distances of HBs 1-5 and 8-12 following the P3 substitutions to Ala from Val and Lys in s01 and s05, respectively.

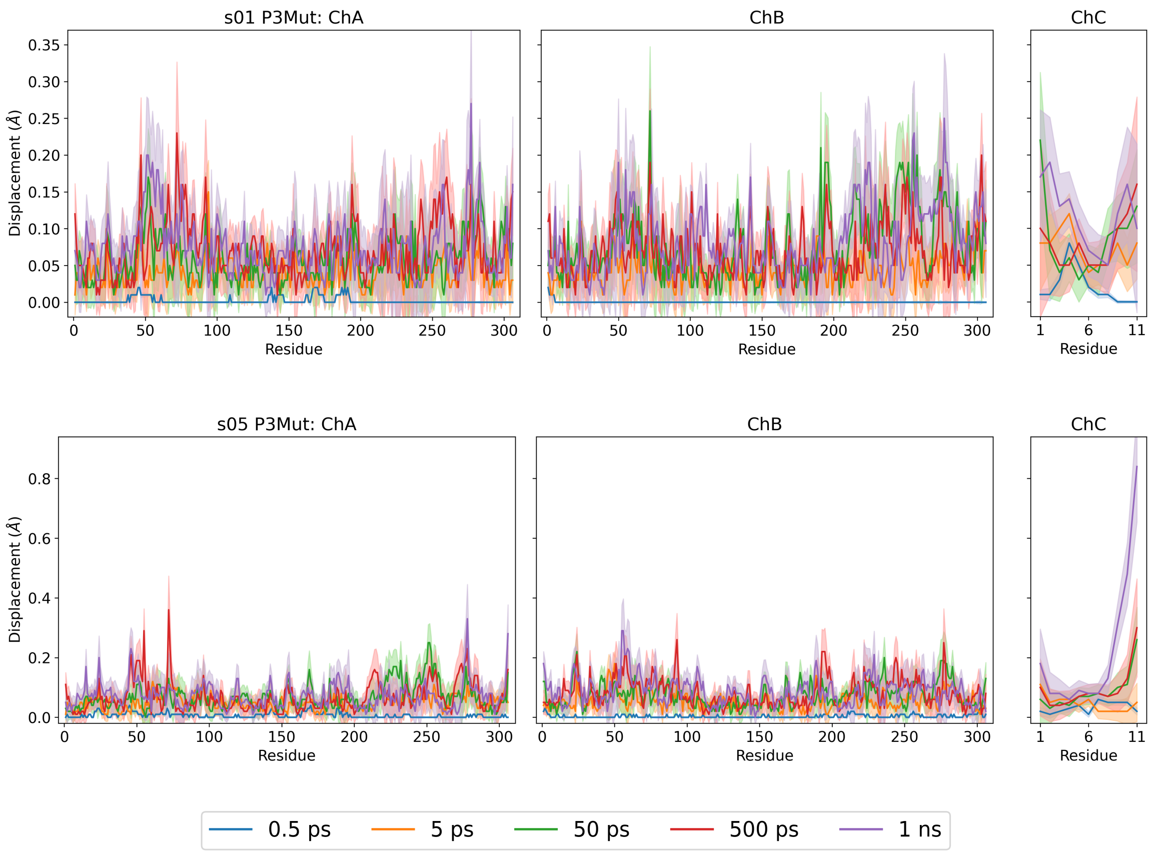

**Figure S4A.3**: Magnitudes of the average Cα displacement vectors, with ± SEM shown as shaded regions, following the P3 substitutions to Ala from Val and Lys in s01 and s05, respectively.

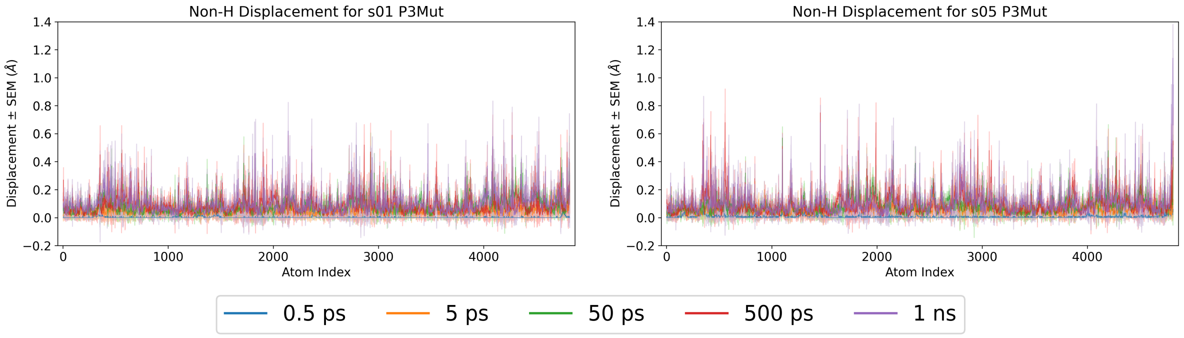

**Figure S4A.4**: Magnitudes of the average non-hydrogen atom displacement vectors in the M$^{pro}$-peptide complex, with ± SEM shown as shaded regions, following the P3 substitutions to Ala from Val and Lys in s01 and s05, respectively.

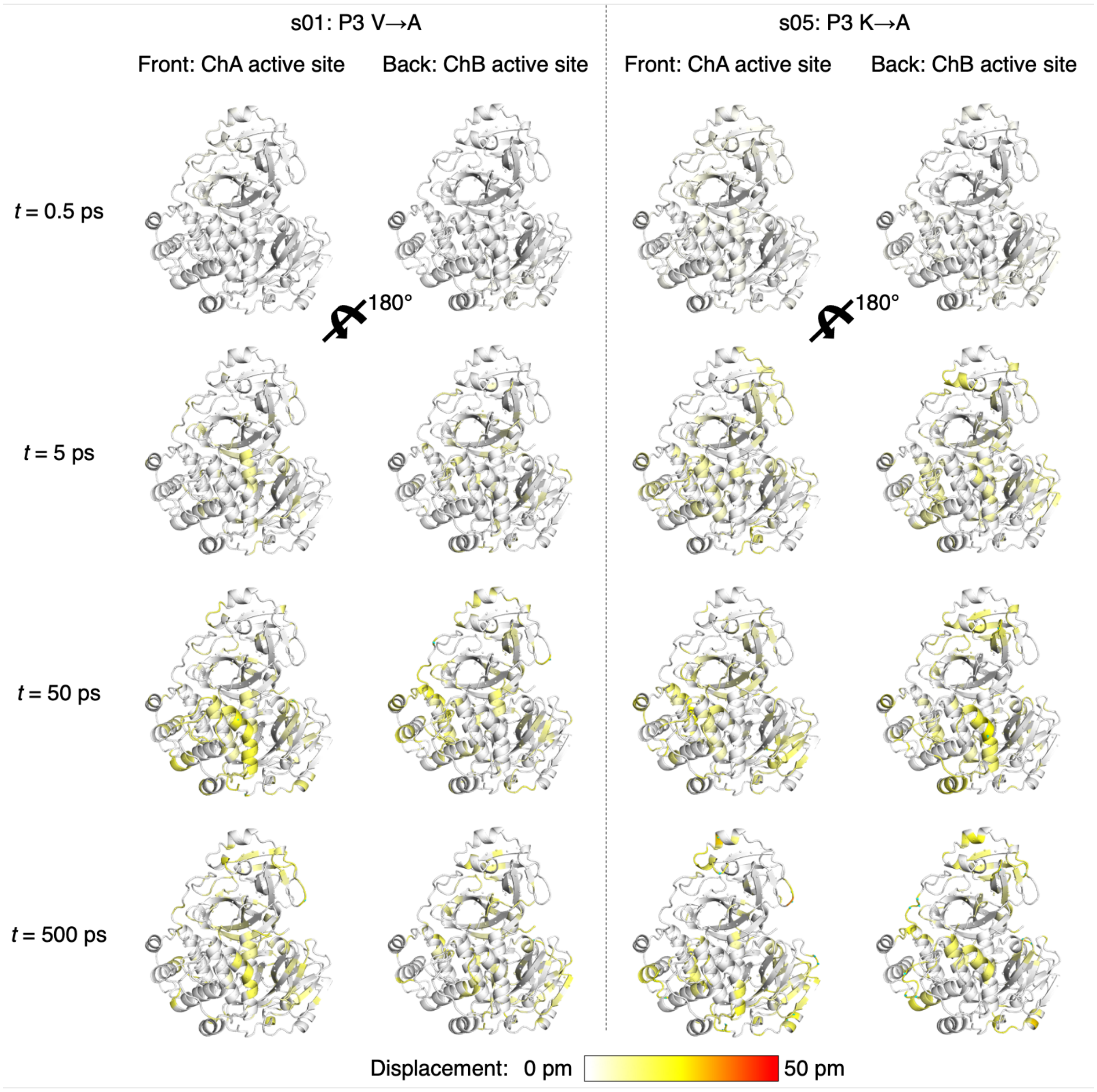

**Figure S4A.5**: Views of the M^pro^ response to the shown P3 substitutions in s01 and s05 from averaging Cα displacement vectors, shown using the structure prior to MD simulations. Displacement magnitudes are represented on a white-yellow-red scale.[7] Significant vectors with length ≥20 pm are displayed as cyan arrows with a scale-up factor of 5.[8]

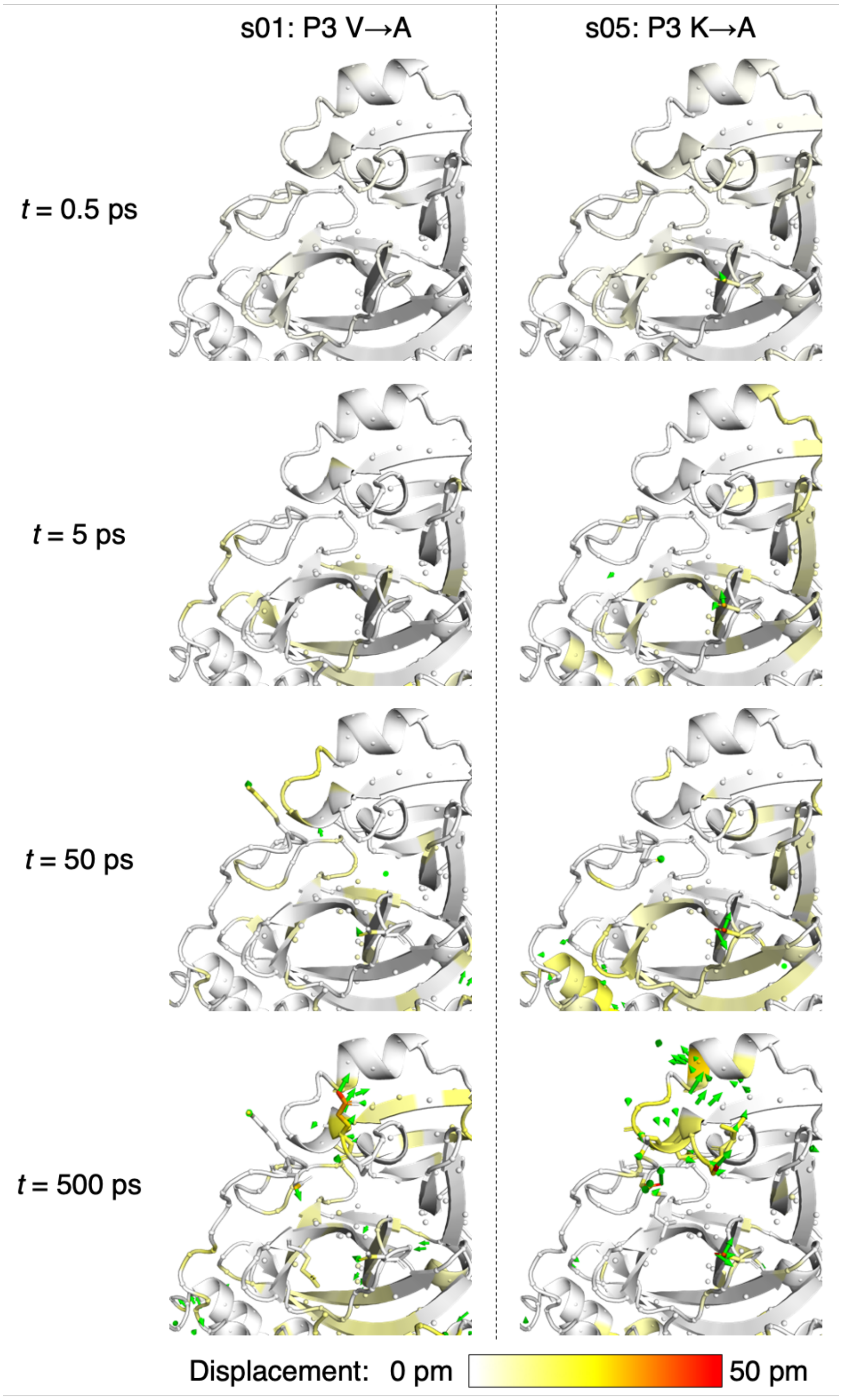

**Figure S4A.6**: Views of the M<sup>pro</sup> response to the shown P3 substitutions in s01 and s05 from averaging non-hydrogen atom displacement vectors, shown using the structure prior to MD simulations, with a focus on the S3 subsite. Displacement magnitudes are represented on a white-yellow-red scale.[7] Significant vectors with length ≥20 pm are displayed as green arrows with a scale-up factor of 5.[8] Residues that show such significant displacements and that are within 10 Å of the substrate P3 residue are shown as sticks.

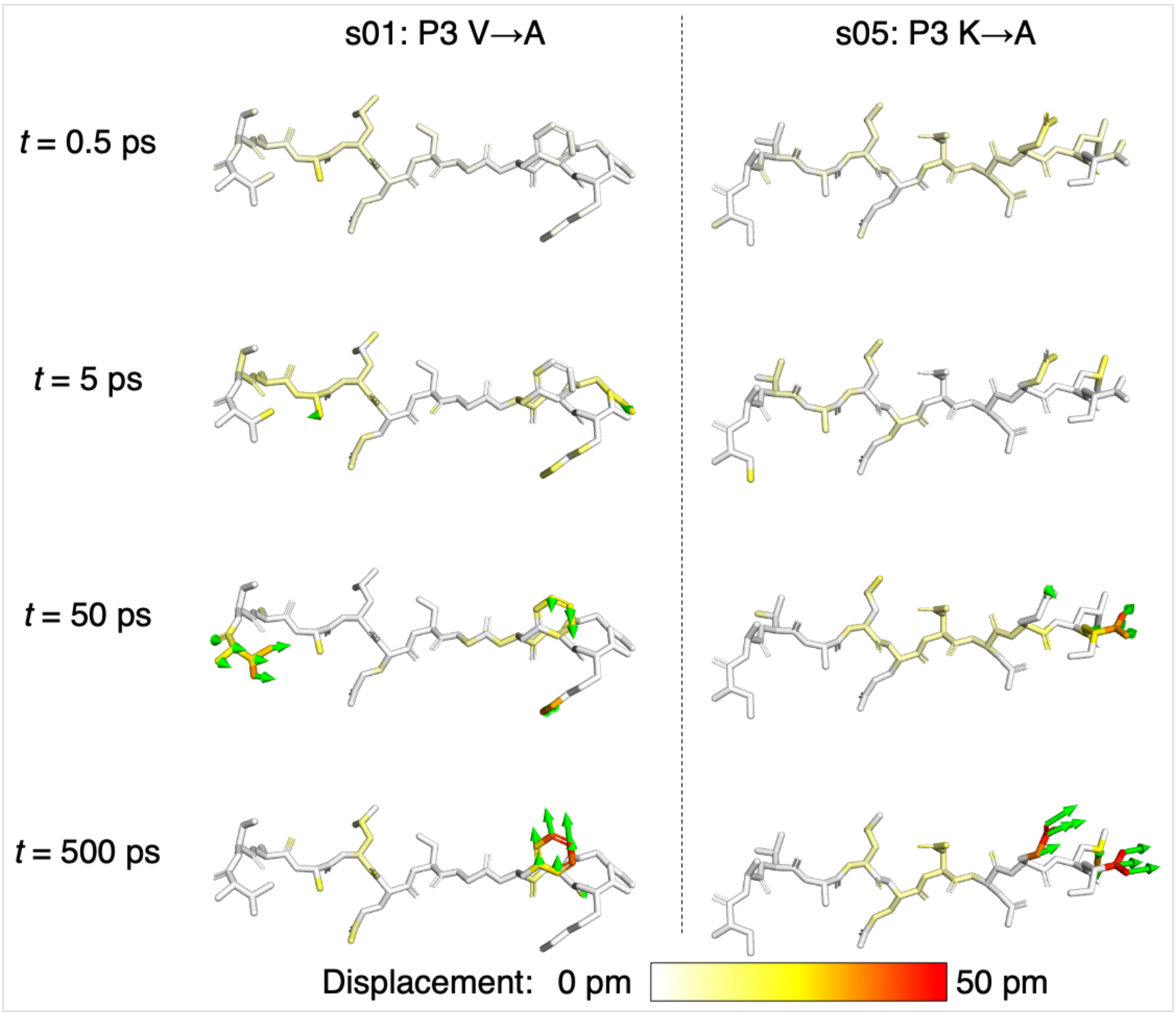

**Figure S4A.7**: Views of the peptide response to the shown P3 substitutions in s01 and s05 from averaging non-hydrogen atom displacement vectors, shown using the structure prior to MD simulations. Displacement magnitudes are represented on a white-yellow-red scale.[7] Significant vectors with length ≥20 pm are displayed as green arrows with a scale-up factor of 5.[8]

### Section S4B: Substitution of P4 residue

**Table S4B.1**: The average differences in the five active-site distances ($\Delta d = d_{neq} - d_{eqm}$) following the P4 Val to Ala substitution in s05. Positive and negative differences with magnitudes > 0.1 Å are in green and red, respectively.

| Active Site | s05: P4 V->A | | |
|---|---|---|---|
| **Deprot** | t (ps) | Av Diff (Å) | SEM (Å) |
| | 0.05 | 0.00 | 0.00 |
| | 0.1 | 0.00 | 0.00 |
| | 0.25 | 0.00 | 0.00 |
| | 0.5 | 0.01 | 0.01 |
| | 1 | 0.02 | 0.03 |
| | 2 | 0.00 | 0.05 |
| | 3 | 0.05 | 0.07 |
| | 5 | 0.01 | 0.07 |
| | 10 | -0.01 | 0.06 |
| | 20 | -0.05 | 0.07 |
| | 30 | 0.03 | 0.06 |
| | 50 | 0.00 | 0.07 |
| | 100 | -0.03 | 0.07 |
| | 300 | 0.06 | 0.07 |
| | 500 | 0.02 | 0.07 |
| | 1000 | 0.12 | 0.08 |
| **Nuc** | t (ps) | Av Diff (Å) | SEM (Å) |
| | 0.05 | 0.00 | 0.00 |
| | 0.1 | 0.00 | 0.00 |
| | 0.25 | 0.01 | 0.00 |
| | 0.5 | 0.00 | 0.00 |
| | 1 | 0.00 | 0.01 |
| | 2 | 0.01 | 0.02 |
| | 3 | 0.03 | 0.02 |
| | 5 | -0.02 | 0.02 |
| | 10 | -0.05 | 0.03 |
| | 20 | -0.04 | 0.03 |
| | 30 | 0.01 | 0.03 |
| | 50 | -0.02 | 0.04 |
| | 100 | 0.01 | 0.04 |
| | 300 | -0.05 | 0.05 |
| | 500 | -0.03 | 0.05 |
| | 1000 | -0.01 | 0.05 |
| **OH145** | t (ps) | Av Diff (Å) | SEM (Å) |
| | 0.05 | 0.00 | 0.00 |
| | 0.1 | 0.00 | 0.00 |
| | 0.25 | 0.01 | 0.00 |
| | 0.5 | 0.01 | 0.00 |
| | 1 | 0.00 | 0.01 |
| | 2 | -0.01 | 0.02 |
| | 3 | -0.01 | 0.03 |
| | 5 | 0.00 | 0.03 |
| | 10 | -0.02 | 0.03 |
| | 20 | 0.01 | 0.02 |
| | 30 | 0.01 | 0.03 |
| | 50 | -0.01 | 0.02 |
| | 100 | -0.05 | 0.03 |
| | 300 | 0.01 | 0.03 |
| | 500 | 0.04 | 0.03 |
| | 1000 | 0.02 | 0.03 |
| **OH144** | t (ps) | Av Diff (Å) | SEM (Å) |
| | 0.05 | 0.00 | 0.00 |
| | 0.1 | 0.00 | 0.00 |
| | 0.25 | 0.00 | 0.00 |
| | 0.5 | 0.00 | 0.00 |
| | 1 | -0.01 | 0.02 |
| | 2 | -0.05 | 0.03 |
| | 3 | -0.01 | 0.03 |
| | 5 | 0.03 | 0.03 |
| | 10 | 0.02 | 0.03 |
| | 20 | -0.02 | 0.03 |
| | 30 | 0.03 | 0.03 |
| | 50 | -0.02 | 0.04 |
| | 100 | -0.01 | 0.04 |
| | 300 | -0.02 | 0.04 |
| | 500 | -0.02 | 0.04 |
| | 1000 | -0.05 | 0.04 |
| **OH143** | t (ps) | Av Diff (Å) | SEM (Å) |
| | 0.05 | 0.00 | 0.00 |
| | 0.1 | 0.00 | 0.00 |
| | 0.25 | 0.00 | 0.00 |
| | 0.5 | -0.01 | 0.00 |
| | 1 | 0.01 | 0.02 |
| | 2 | 0.03 | 0.03 |
| | 3 | -0.02 | 0.03 |
| | 5 | 0.00 | 0.03 |
| | 10 | 0.02 | 0.03 |
| | 20 | -0.04 | 0.03 |
| | 30 | 0.00 | 0.03 |
| | 50 | -0.01 | 0.03 |
| | 100 | 0.02 | 0.03 |
| | 300 | 0.00 | 0.03 |
| | 500 | -0.02 | 0.03 |
| | 1000 | -0.03 | 0.03 |

**Table S4B.2**: The average differences in the donor-acceptor distances of HBs 1-5 following the P4 Val to Ala substitution in s05. Positive and negative differences with magnitudes > 0.1 Å are in green and red, respectively.

| HB1-5 | s05: P4 V->A | | |
|---|---|---|---|
| HB1 | t (ps) | Av Diff (Å) | SEM (Å) |
| | 0.05 | -0.01 | 0.00 |
| | 0.1 | -0.01 | 0.01 |
| | 0.25 | -0.06 | 0.01 |
| | 0.5 | -0.11 | 0.02 |
| | 1 | -0.13 | 0.03 |
| | 2 | -0.10 | 0.04 |
| | 3 | -0.09 | 0.03 |
| | 5 | -0.11 | 0.04 |
| | 10 | -0.12 | 0.03 |
| | 20 | -0.15 | 0.04 |
| | 30 | -0.14 | 0.04 |
| | 50 | -0.14 | 0.04 |
| | 100 | -0.12 | 0.04 |
| | 300 | -0.14 | 0.06 |
| | 500 | -0.19 | 0.06 |
| | 1000 | -0.10 | 0.07 |
| HB2 | t (ps) | Av Diff (Å) | SEM (Å) |
| | 0.05 | 0.00 | 0.00 |
| | 0.1 | -0.02 | 0.00 |
| | 0.25 | -0.05 | 0.01 |
| | 0.5 | -0.06 | 0.01 |
| | 1 | -0.03 | 0.01 |
| | 2 | -0.06 | 0.02 |
| | 3 | -0.04 | 0.02 |
| | 5 | -0.08 | 0.02 |
| | 10 | -0.06 | 0.02 |
| | 20 | -0.05 | 0.02 |
| | 30 | -0.06 | 0.02 |
| | 50 | -0.02 | 0.02 |
| | 100 | -0.04 | 0.02 |
| | 300 | -0.05 | 0.02 |
| | 500 | -0.04 | 0.02 |
| | 1000 | -0.06 | 0.02 |
| HB3 | t (ps) | Av Diff (Å) | SEM (Å) |
| | 0.05 | 0.00 | 0.00 |
| | 0.1 | 0.00 | 0.00 |
| | 0.25 | 0.00 | 0.00 |
| | 0.5 | -0.02 | 0.01 |
| | 1 | -0.01 | 0.01 |
| | 2 | -0.01 | 0.01 |
| | 3 | 0.00 | 0.01 |
| | 5 | -0.04 | 0.02 |
| | 10 | -0.02 | 0.01 |
| | 20 | -0.02 | 0.01 |
| | 30 | 0.01 | 0.01 |
| | 50 | 0.00 | 0.01 |
| | 100 | -0.02 | 0.01 |
| | 300 | 0.00 | 0.01 |
| | 500 | -0.01 | 0.01 |
| | 1000 | -0.01 | 0.02 |
| HB4 | t (ps) | Av Diff (Å) | SEM (Å) |
| | 0.05 | 0.00 | 0.00 |
| | 0.1 | 0.00 | 0.00 |
| | 0.25 | 0.01 | 0.00 |
| | 0.5 | 0.01 | 0.01 |
| | 1 | -0.01 | 0.02 |
| | 2 | -0.06 | 0.03 |
| | 3 | 0.00 | 0.03 |
| | 5 | 0.01 | 0.04 |
| | 10 | -0.07 | 0.04 |
| | 20 | -0.08 | 0.05 |
| | 30 | -0.08 | 0.06 |
| | 50 | -0.15 | 0.06 |
| | 100 | -0.09 | 0.08 |
| | 300 | -0.13 | 0.09 |
| | 500 | 0.00 | 0.10 |
| | 1000 | -0.09 | 0.14 |
| HB5 | t (ps) | Av Diff (Å) | SEM (Å) |
| | 0.05 | 0.00 | 0.00 |
| | 0.1 | 0.00 | 0.00 |
| | 0.25 | 0.01 | 0.00 |
| | 0.5 | 0.02 | 0.01 |
| | 1 | -0.02 | 0.01 |
| | 2 | 0.04 | 0.02 |
| | 3 | 0.03 | 0.02 |
| | 5 | 0.07 | 0.02 |
| | 10 | 0.00 | 0.02 |
| | 20 | 0.02 | 0.03 |
| | 30 | 0.01 | 0.03 |
| | 50 | 0.02 | 0.03 |
| | 100 | -0.01 | 0.03 |
| | 300 | 0.08 | 0.03 |
| | 500 | 0.03 | 0.03 |
| | 1000 | 0.03 | 0.03 |

**Table S4B.3**: The average differences in the donor-acceptor distances of HBs 8-12 following the P4 Val to Ala substitution in s05. Positive and negative differences with magnitudes > 0.1 Å are in green and red, respectively.

| HB8-12 | s05: P4 V->A | | |
|---|---|---|---|
| HB8 | t (ps) | Av Diff (Å) | SEM (Å) |
| | 0.05 | 0.00 | 0.00 |
| | 0.1 | 0.00 | 0.00 |
| | 0.25 | 0.01 | 0.00 |
| | 0.5 | 0.01 | 0.00 |
| | 1 | 0.00 | 0.01 |
| | 2 | -0.01 | 0.02 |
| | 3 | -0.01 | 0.02 |
| | 5 | 0.00 | 0.03 |
| | 10 | -0.02 | 0.03 |
| | 20 | 0.02 | 0.02 |
| | 30 | 0.00 | 0.03 |
| | 50 | -0.01 | 0.02 |
| | 100 | -0.06 | 0.03 |
| | 300 | 0.02 | 0.03 |
| | 500 | 0.03 | 0.03 |
| | 1000 | 0.02 | 0.03 |
| HB9 | t (ps) | Av Diff (Å) | SEM (Å) |
| | 0.05 | 0.00 | 0.00 |
| | 0.1 | 0.00 | 0.00 |
| | 0.25 | 0.00 | 0.00 |
| | 0.5 | 0.00 | 0.00 |
| | 1 | 0.01 | 0.01 |
| | 2 | 0.00 | 0.02 |
| | 3 | -0.02 | 0.02 |
| | 5 | 0.00 | 0.02 |
| | 10 | 0.01 | 0.02 |
| | 20 | -0.01 | 0.01 |
| | 30 | 0.00 | 0.02 |
| | 50 | 0.00 | 0.02 |
| | 100 | 0.01 | 0.02 |
| | 300 | -0.01 | 0.02 |
| | 500 | -0.02 | 0.02 |
| | 1000 | -0.02 | 0.02 |
| HB10 | t (ps) | Av Diff (Å) | SEM (Å) |
| | 0.05 | 0.00 | 0.00 |
| | 0.1 | 0.00 | 0.00 |
| | 0.25 | 0.00 | 0.00 |
| | 0.5 | 0.00 | 0.00 |
| | 1 | 0.02 | 0.01 |
| | 2 | 0.00 | 0.02 |
| | 3 | 0.03 | 0.02 |
| | 5 | 0.02 | 0.02 |
| | 10 | -0.01 | 0.02 |
| | 20 | 0.00 | 0.02 |
| | 30 | 0.00 | 0.02 |
| | 50 | 0.01 | 0.02 |
| | 100 | -0.01 | 0.02 |
| | 300 | 0.01 | 0.02 |
| | 500 | -0.03 | 0.02 |
| | 1000 | 0.05 | 0.02 |
| HB11 | t (ps) | Av Diff (Å) | SEM (Å) |
| | 0.05 | 0.00 | 0.00 |
| | 0.1 | 0.00 | 0.00 |
| | 0.25 | 0.00 | 0.00 |
| | 0.5 | 0.00 | 0.00 |
| | 1 | 0.01 | 0.01 |
| | 2 | -0.02 | 0.02 |
| | 3 | 0.01 | 0.02 |
| | 5 | -0.01 | 0.02 |
| | 10 | 0.01 | 0.01 |
| | 20 | 0.03 | 0.02 |
| | 30 | 0.00 | 0.02 |
| | 50 | 0.01 | 0.02 |
| | 100 | -0.02 | 0.02 |
| | 300 | 0.01 | 0.02 |
| | 500 | 0.01 | 0.02 |
| | 1000 | 0.03 | 0.02 |
| HB12 | t (ps) | Av Diff (Å) | SEM (Å) |
| | 0.05 | 0.00 | 0.00 |
| | 0.1 | 0.00 | 0.00 |
| | 0.25 | 0.00 | 0.00 |
| | 0.5 | 0.00 | 0.00 |
| | 1 | 0.00 | 0.01 |
| | 2 | -0.04 | 0.02 |
| | 3 | -0.01 | 0.02 |
| | 5 | -0.05 | 0.02 |
| | 10 | 0.00 | 0.02 |
| | 20 | 0.02 | 0.03 |
| | 30 | 0.01 | 0.03 |
| | 50 | 0.01 | 0.03 |
| | 100 | -0.01 | 0.03 |
| | 300 | 0.01 | 0.03 |
| | 500 | -0.06 | 0.04 |
| | 1000 | 0.02 | 0.04 |

**Table S4B.4**: The ten most perturbed Cα atoms in the $M^{pro}$-peptide complex in terms of magnitude of the average displacement vector *v*, at selected time points following the P4 Val to Ala substitution in s05. Only displacements ≥ 0.01 Å are shown.

| s05 t(ps) | CA Rank | Atom | Chain | v (Å) | SEM (Å) |
|---|---|---|---|---|---|
| **0.05** | 1 | 3ALA_CA | C | 0.03 | 0.00 |
| | 2 | 2ALA_CA | C | 0.02 | 0.00 |
| | 3 | 167LEU_CA | A | 0.01 | 0.00 |
| | 4 | 188ARG_CA | A | 0.01 | 0.00 |
| | 5 | 189GLN_CA | A | 0.01 | 0.00 |
| | 6 | 190THR_CA | A | 0.01 | 0.00 |
| | 7 | 192GLN_CA | A | 0.01 | 0.00 |
| | 8 | 1SER_CA | C | 0.01 | 0.00 |
| | 9 | 4LYS_CA | C | 0.01 | 0.00 |
| | 10 | | | | |
| **0.5** | 1 | 4LYS_CA | C | 0.16 | 0.01 |
| | 2 | 192GLN_CA | A | 0.15 | 0.01 |
| | 3 | 3ALA_CA | C | 0.15 | 0.01 |
| | 4 | 188ARG_CA | A | 0.10 | 0.01 |
| | 5 | 167LEU_CA | A | 0.08 | 0.01 |
| | 6 | 171VAL_CA | A | 0.08 | 0.01 |
| | 7 | 189GLN_CA | A | 0.08 | 0.01 |
| | 8 | 190THR_CA | A | 0.08 | 0.01 |
| | 9 | 193ALA_CA | A | 0.07 | 0.01 |
| | 10 | 186VAL_CA | A | 0.06 | 0.01 |
| **5** | 1 | 192GLN_CA | A | 0.28 | 0.04 |
| | 2 | 193ALA_CA | A | 0.26 | 0.06 |
| | 3 | 188ARG_CA | A | 0.18 | 0.03 |
| | 4 | 190THR_CA | A | 0.18 | 0.04 |
| | 5 | 191ALA_CA | A | 0.18 | 0.04 |
| | 6 | 189GLN_CA | A | 0.17 | 0.03 |
| | 7 | 1SER_CA | C | 0.17 | 0.07 |
| | 8 | 186VAL_CA | A | 0.16 | 0.03 |
| | 9 | 2ALA_CA | C | 0.16 | 0.04 |
| | 10 | 3ALA_CA | C | 0.16 | 0.03 |
| **50** | 1 | 193ALA_CA | A | 0.69 | 0.07 |
| | 2 | 194ALA_CA | A | 0.58 | 0.07 |
| | 3 | 192GLN_CA | A | 0.50 | 0.05 |
| | 4 | 195GLY_CA | A | 0.48 | 0.07 |
| | 5 | 1SER_CA | C | 0.42 | 0.11 |
| | 6 | 191ALA_CA | A | 0.30 | 0.05 |
| | 7 | 196THR_CA | A | 0.28 | 0.07 |
| | 8 | 2ALA_CA | C | 0.28 | 0.06 |
| | 9 | 11SER_CA | C | 0.27 | 0.13 |
| | 10 | 72ASN_CA | A | 0.26 | 0.09 |
| **500** | 1 | 194ALA_CA | A | 0.88 | 0.09 |
| | 2 | 193ALA_CA | A | 0.87 | 0.09 |
| | 3 | 195GLY_CA | A | 0.63 | 0.09 |
| | 4 | 192GLN_CA | A | 0.42 | 0.05 |
| | 5 | 168PRO_CA | A | 0.30 | 0.06 |
| | 6 | 1SER_CA | C | 0.29 | 0.10 |
| | 7 | 11SER_CA | C | 0.28 | 0.19 |
| | 8 | 189GLN_CA | A | 0.27 | 0.04 |
| | 9 | 2ALA_CA | C | 0.27 | 0.08 |
| | 10 | 191ALA_CA | B | 0.26 | 0.11 |
| **1000** | 1 | 193ALA_CA | A | 0.91 | 0.08 |
| | 2 | 194ALA_CA | A | 0.79 | 0.08 |
| | 3 | 195GLY_CA | A | 0.62 | 0.09 |
| | 4 | 11SER_CA | C | 0.53 | 0.20 |
| | 5 | 192GLN_CA | A | 0.51 | 0.06 |
| | 6 | 1SER_CA | C | 0.44 | 0.15 |
| | 7 | 196THR_CA | A | 0.39 | 0.08 |
| | 8 | 2ALA_CA | C | 0.37 | 0.09 |
| | 9 | 168PRO_CA | A | 0.32 | 0.06 |
| | 10 | 191ALA_CA | A | 0.31 | 0.06 |

**Table S4B.5**: The ten most perturbed non-hydrogen atoms in the M$^{pro}$-peptide complex in terms of magnitude of the average displacement vector *v*, at selected time points following the P4 Val to Ala substitution in s05.

| ***s05 t(ps)*** | ***Non-H Rank*** | *Atom* | *Chain* | *v (Å)* | *SEM (Å)* |
|---|---|---|---|---|---|
| **0.05** | 1 | 3ALA_CB | C | 0.11 | 0.01 |
| | 2 | 192GLN_CB | A | 0.05 | 0.00 |
| | 3 | 3ALA_N | C | 0.04 | 0.00 |
| | 4 | 3ALA_CA | C | 0.03 | 0.00 |
| | 5 | 4LYS_N | C | 0.02 | 0.00 |
| | 6 | 2ALA_O | C | 0.02 | 0.00 |
| | 7 | 167LEU_CD2 | A | 0.02 | 0.00 |
| | 8 | 190THR_O | A | 0.02 | 0.00 |
| | 9 | 188ARG_O | A | 0.02 | 0.00 |
| | 10 | 165MET_CE | A | 0.02 | 0.00 |
| **0.5** | 1 | 167LEU_CD1 | A | 0.33 | 0.04 |
| | 2 | 167LEU_CD2 | A | 0.32 | 0.03 |
| | 3 | 165MET_CE | A | 0.29 | 0.04 |
| | 4 | 167LEU_CG | A | 0.28 | 0.02 |
| | 5 | 165MET_SD | A | 0.27 | 0.03 |
| | 6 | 3ALA_CB | C | 0.23 | 0.02 |
| | 7 | 3ALA_O | C | 0.22 | 0.02 |
| | 8 | 192GLN_CB | A | 0.21 | 0.01 |
| | 9 | 4LYS_N | C | 0.20 | 0.01 |
| | 10 | 192GLN_NE2 | A | 0.20 | 0.02 |
| **5** | 1 | 165MET_CE | A | 0.54 | 0.08 |
| | 2 | 193ALA_CB | A | 0.33 | 0.07 |
| | 3 | 167LEU_CD2 | A | 0.32 | 0.05 |
| | 4 | 192GLN_CB | A | 0.31 | 0.04 |
| | 5 | 3ALA_CB | C | 0.29 | 0.04 |
| | 6 | 192GLN_CA | A | 0.28 | 0.04 |
| | 7 | 192GLN_C | A | 0.28 | 0.05 |
| | 8 | 193ALA_N | A | 0.28 | 0.05 |
| | 9 | 192GLN_O | A | 0.27 | 0.05 |
| | 10 | 193ALA_CA | A | 0.26 | 0.06 |
| **50** | 1 | 167LEU_CD2 | A | 0.87 | 0.08 |
| | 2 | 193ALA_CB | A | 0.83 | 0.10 |
| | 3 | 193ALA_CA | A | 0.69 | 0.07 |
| | 4 | 194ALA_N | A | 0.64 | 0.07 |
| | 5 | 193ALA_N | A | 0.63 | 0.06 |
| | 6 | 193ALA_C | A | 0.62 | 0.07 |
| | 7 | 192GLN_O | A | 0.60 | 0.07 |
| | 8 | 165MET_CE | A | 0.59 | 0.09 |
| | 9 | 191ALA_O | A | 0.59 | 0.08 |
| | 10 | 194ALA_CA | A | 0.58 | 0.07 |
| **500** | 1 | 193ALA_CB | A | 1.13 | 0.11 |
| | 2 | 167LEU_CD2 | A | 1.11 | 0.08 |
| | 3 | 194ALA_N | A | 0.90 | 0.09 |
| | 4 | 194ALA_CB | A | 0.89 | 0.10 |
| | 5 | 194ALA_CA | A | 0.88 | 0.09 |
| | 6 | 193ALA_CA | A | 0.87 | 0.09 |
| | 7 | 193ALA_C | A | 0.82 | 0.08 |
| | 8 | 194ALA_O | A | 0.80 | 0.09 |
| | 9 | 194ALA_C | A | 0.79 | 0.08 |
| | 10 | 193ALA_O | A | 0.76 | 0.08 |
| **1000** | 1 | 193ALA_CB | A | 1.13 | 0.11 |
| | 2 | 167LEU_CD2 | A | 1.11 | 0.09 |
| | 3 | 11SER_OG | C | 0.92 | 0.26 |
| | 4 | 193ALA_CA | A | 0.91 | 0.08 |
| | 5 | 194ALA_N | A | 0.86 | 0.08 |
| | 6 | 222ARG_NH2 | B | 0.85 | 0.33 |
| | 7 | 196THR_OG1 | A | 0.81 | 0.14 |
| | 8 | 194ALA_O | A | 0.81 | 0.09 |
| | 9 | 193ALA_C | A | 0.80 | 0.08 |
| | 10 | 194ALA_CA | A | 0.79 | 0.08 |

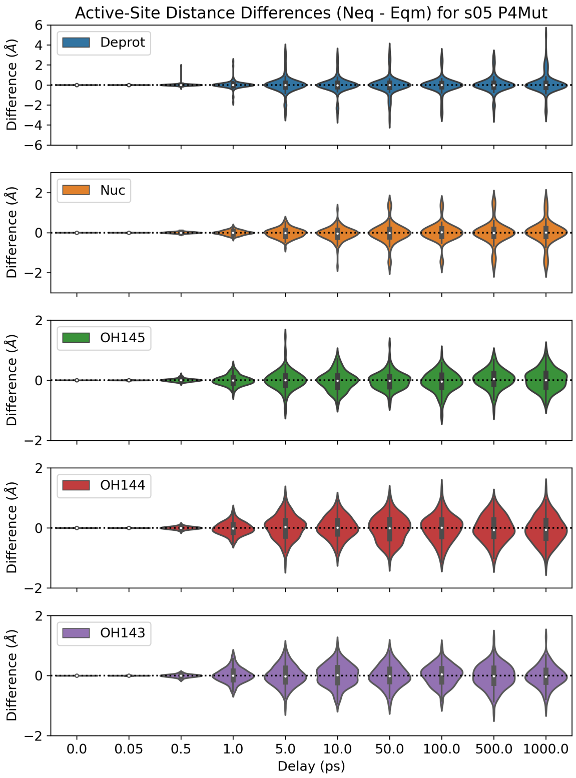

**Figure S4B.1**: Distributions of the distance differences in the five active-site distances ($\Delta d = d_{neq} - d_{eqm}$) following the P4 Val to Ala substitution in s05.

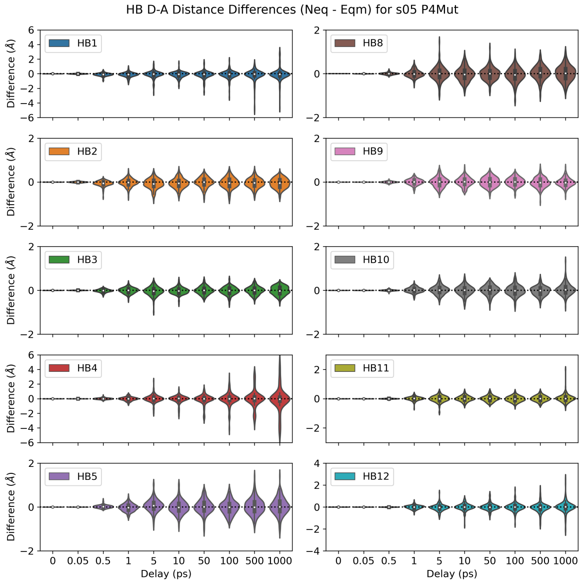

**Figure S4B.2**: Distributions of the distance differences in the ten donor-acceptor distances of HBs 1-5 and 8-12 following the P4 Val to Ala substitution in s05.

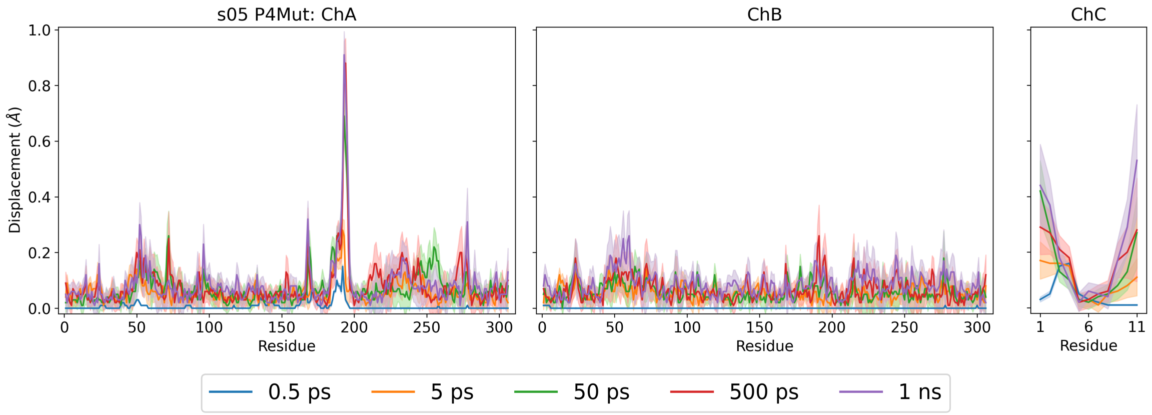

**Figure S4B.3**: Magnitudes of the average Cα displacement vectors, with ± SEM shown as shaded regions, following the P4 Val to Ala substitution in s05.

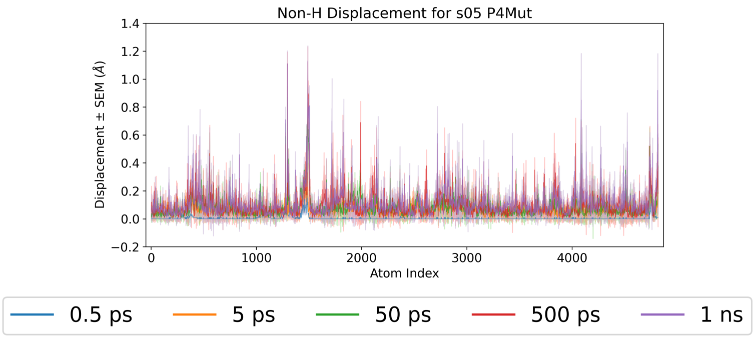

**Figure S4B.4**: Magnitudes of the average non-hydrogen atom displacement vectors in the $M^{pro}$-peptide complex, with ± SEM shown as shaded regions, following the P4 Val to Ala substitution in s05.

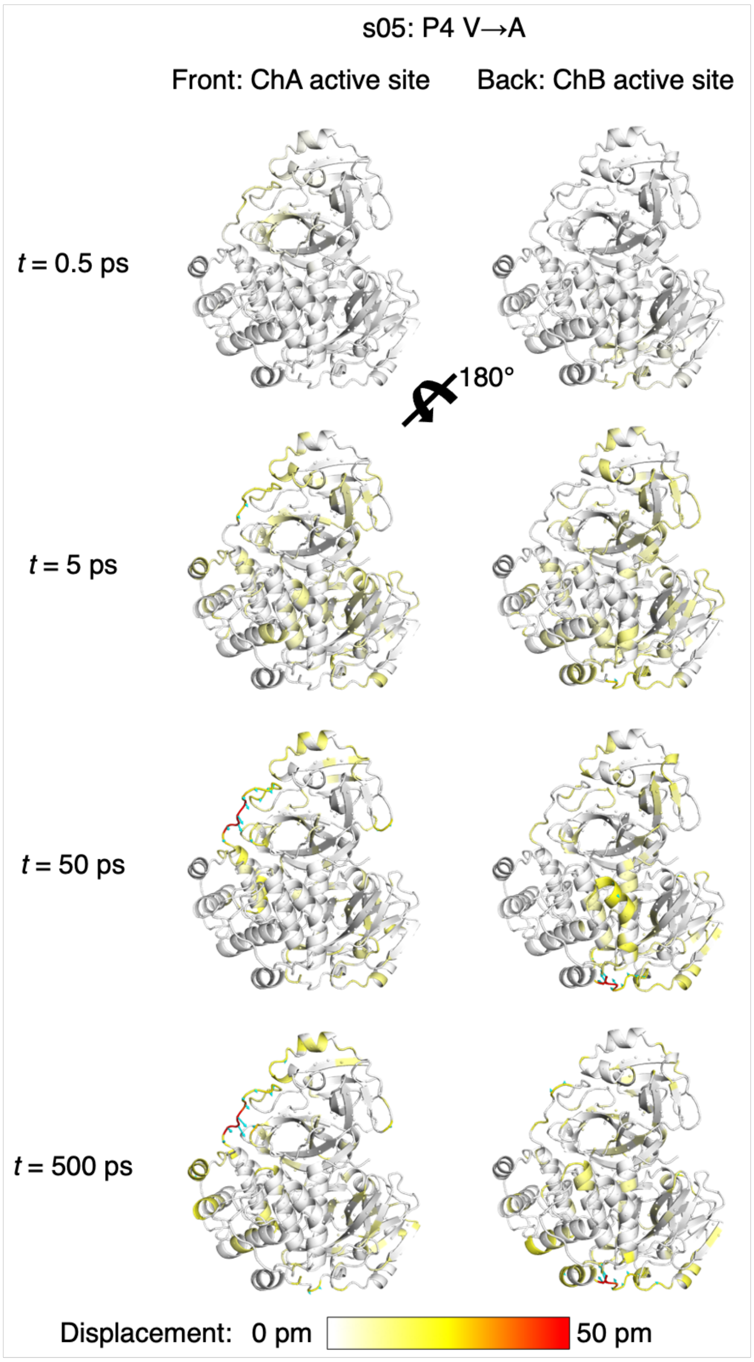

**Figure S4B.5**: Views of the M$^{pro}$ response to the P4 Val to Ala substitution in s05 from averaging Cα displacement vectors, shown using the structure prior to MD simulations. Displacement magnitudes are represented on a white-yellow-red scale.[7] Significant vectors with length ≥20 pm are displayed as cyan arrows with a scale-up factor of 5.[8]

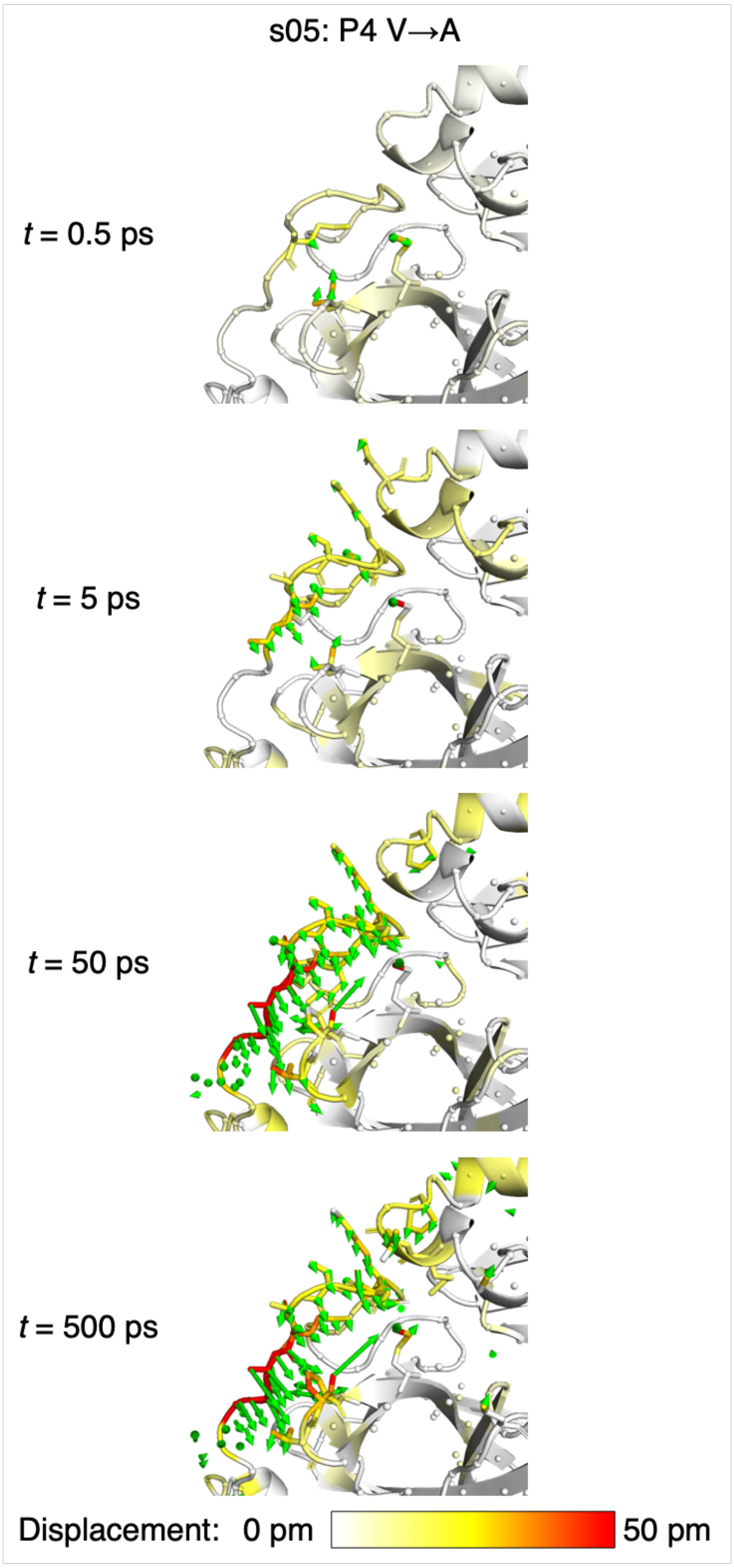

**Figure S4B.6**: Views of the M$^{pro}$ response to the P4 Val to Ala substitution in s05 from averaging non-hydrogen atom displacement vectors, shown using the structure prior to MD simulations, with a focus on the S4 subsite. Displacement magnitudes are represented on a white-yellow-red scale.[7] Significant vectors with length ≥20 pm are displayed as green arrows with a scale-up factor of 5.[8] Residues that show such significant displacements and that are within 10 Å of the substrate P4 residue are shown as sticks.

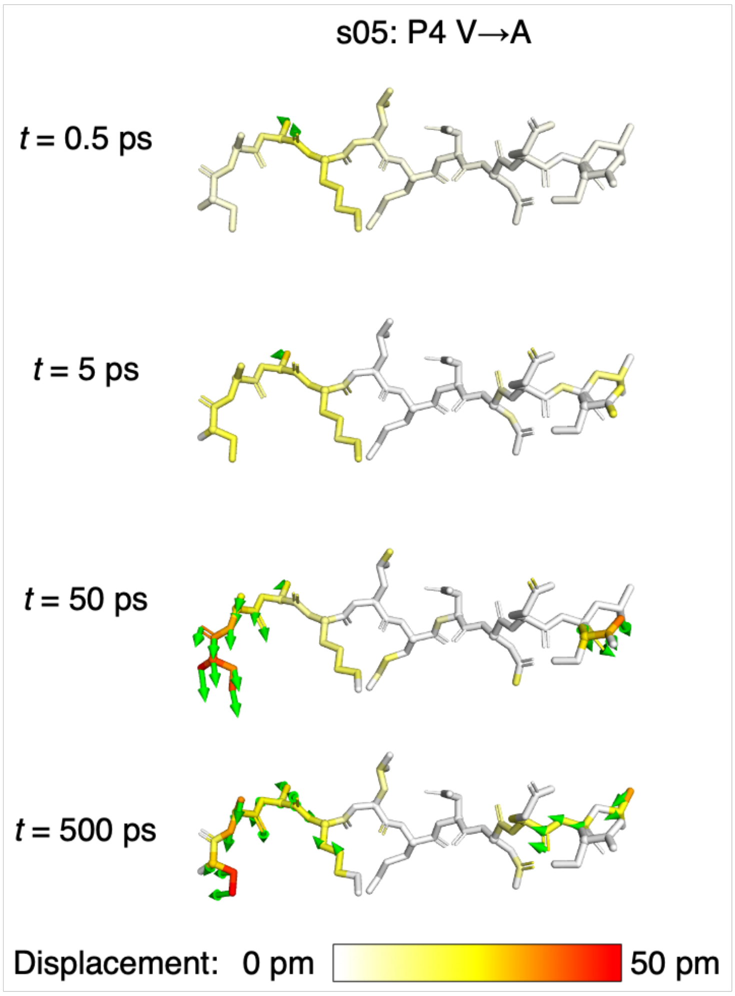

**Figure S4B.7**: Views of the peptide response to the P4 Val to Ala substitution in s05 from averaging non-hydrogen atom displacement vectors, shown using the structure prior to MD simulations. Displacement magnitudes are represented on a white-yellow-red scale.[7] Significant vectors with length ≥20 pm are displayed as green arrows with a scale-up factor of 5.[8]

### Section S5A: Substitution of P1′ residue

**Table S5A.1**: The average differences in the five active-site distances ($\Delta d = d_{neq} - d_{eqm}$) following the shown P1′ substitutions in s01 and s05. Positive and negative differences with magnitudes > 0.1 Å are in green and red, respectively.

| **Active Site** | s01: P1' S->A | | | | s05: P1' N->A | | |
|---|---|---|---|---|---|---|---|
| **Deprot** | t (ps) | Av Diff (Å) | SEM (Å) | | t (ps) | Av Diff (Å) | SEM (Å) |
| | 0.05 | 0.00 | 0.00 | | 0.05 | -0.01 | 0.00 |
| | 0.1 | -0.01 | 0.00 | | 0.1 | 0.00 | 0.01 |
| | 0.25 | -0.02 | 0.01 | | 0.25 | -0.06 | 0.02 |
| | 0.5 | 0.00 | 0.02 | | 0.5 | -0.04 | 0.02 |
| | 1 | -0.06 | 0.04 | | 1 | -0.07 | 0.04 |
| | 2 | -0.12 | 0.04 | | 2 | -0.05 | 0.06 |
| | 3 | -0.02 | 0.04 | | 3 | -0.07 | 0.07 |
| | 5 | -0.10 | 0.06 | | 5 | -0.22 | 0.06 |
| | 10 | 0.02 | 0.06 | | 10 | -0.10 | 0.06 |
| | 20 | 0.00 | 0.04 | | 20 | -0.17 | 0.07 |
| | 30 | -0.10 | 0.06 | | 30 | -0.11 | 0.07 |
| | 50 | -0.04 | 0.06 | | 50 | -0.17 | 0.07 |
| | 100 | -0.05 | 0.07 | | 100 | -0.25 | 0.06 |
| | 300 | -0.23 | 0.06 | | 300 | -0.12 | 0.06 |
| | 500 | -0.14 | 0.05 | | 500 | -0.10 | 0.05 |
| | 1000 | -0.12 | 0.06 | | 1000 | -0.06 | 0.06 |
| **Nuc** | t (ps) | Av Diff (Å) | SEM (Å) | | t (ps) | Av Diff (Å) | SEM (Å) |
| | 0.05 | 0.00 | 0.00 | | 0.05 | 0.00 | 0.00 |
| | 0.1 | 0.00 | 0.00 | | 0.1 | 0.01 | 0.00 |
| | 0.25 | -0.01 | 0.01 | | 0.25 | 0.01 | 0.01 |
| | 0.5 | 0.01 | 0.01 | | 0.5 | -0.01 | 0.01 |
| | 1 | -0.01 | 0.01 | | 1 | 0.00 | 0.01 |
| | 2 | 0.00 | 0.02 | | 2 | 0.02 | 0.02 |
| | 3 | -0.04 | 0.02 | | 3 | 0.02 | 0.02 |
| | 5 | -0.03 | 0.02 | | 5 | -0.04 | 0.02 |
| | 10 | 0.00 | 0.02 | | 10 | 0.01 | 0.02 |
| | 20 | 0.06 | 0.03 | | 20 | -0.02 | 0.03 |
| | 30 | -0.02 | 0.03 | | 30 | 0.00 | 0.03 |
| | 50 | -0.04 | 0.04 | | 50 | 0.03 | 0.04 |
| | 100 | -0.01 | 0.04 | | 100 | 0.01 | 0.04 |
| | 300 | -0.04 | 0.05 | | 300 | 0.01 | 0.05 |
| | 500 | 0.07 | 0.05 | | 500 | -0.08 | 0.05 |
| | 1000 | 0.04 | 0.05 | | 1000 | -0.07 | 0.06 |
| **OH145** | t (ps) | Av Diff (Å) | SEM (Å) | | t (ps) | Av Diff (Å) | SEM (Å) |
| | 0.05 | 0.01 | 0.00 | | 0.05 | 0.01 | 0.00 |
| | 0.1 | 0.00 | 0.00 | | 0.1 | 0.02 | 0.00 |
| | 0.25 | 0.00 | 0.01 | | 0.25 | 0.05 | 0.01 |
| | 0.5 | 0.03 | 0.01 | | 0.5 | 0.09 | 0.02 |
| | 1 | 0.01 | 0.02 | | 1 | 0.06 | 0.02 |
| | 2 | 0.00 | 0.03 | | 2 | 0.09 | 0.03 |
| | 3 | 0.04 | 0.03 | | 3 | 0.08 | 0.03 |
| | 5 | 0.04 | 0.03 | | 5 | 0.10 | 0.03 |
| | 10 | 0.06 | 0.03 | | 10 | 0.06 | 0.03 |
| | 20 | -0.03 | 0.03 | | 20 | 0.07 | 0.03 |
| | 30 | 0.04 | 0.03 | | 30 | 0.08 | 0.03 |
| | 50 | -0.02 | 0.03 | | 50 | 0.09 | 0.03 |
| | 100 | -0.04 | 0.03 | | 100 | 0.07 | 0.03 |
| | 300 | -0.02 | 0.03 | | 300 | 0.09 | 0.03 |
| | 500 | -0.05 | 0.03 | | 500 | 0.09 | 0.03 |
| | 1000 | 0.02 | 0.03 | | 1000 | 0.04 | 0.03 |
| **OH144** | t (ps) | Av Diff (Å) | SEM (Å) | | t (ps) | Av Diff (Å) | SEM (Å) |
| | 0.05 | 0.00 | 0.00 | | 0.05 | 0.00 | 0.00 |
| | 0.1 | -0.02 | 0.01 | | 0.1 | -0.02 | 0.01 |
| | 0.25 | -0.04 | 0.01 | | 0.25 | 0.01 | 0.01 |
| | 0.5 | 0.00 | 0.01 | | 0.5 | 0.04 | 0.01 |
| | 1 | 0.00 | 0.02 | | 1 | 0.00 | 0.02 |
| | 2 | 0.01 | 0.03 | | 2 | 0.00 | 0.04 |
| | 3 | 0.00 | 0.04 | | 3 | 0.04 | 0.03 |
| | 5 | 0.07 | 0.04 | | 5 | 0.05 | 0.04 |
| | 10 | 0.06 | 0.03 | | 10 | 0.07 | 0.03 |
| | 20 | -0.09 | 0.04 | | 20 | 0.02 | 0.03 |
| | 30 | 0.00 | 0.04 | | 30 | 0.07 | 0.03 |
| | 50 | 0.01 | 0.03 | | 50 | 0.04 | 0.04 |
| | 100 | 0.00 | 0.04 | | 100 | 0.01 | 0.03 |
| | 300 | 0.04 | 0.04 | | 300 | 0.07 | 0.04 |
| | 500 | -0.03 | 0.04 | | 500 | 0.00 | 0.04 |
| | 1000 | 0.00 | 0.04 | | 1000 | 0.01 | 0.04 |
| **OH143** | t (ps) | Av Diff (Å) | SEM (Å) | | t (ps) | Av Diff (Å) | SEM (Å) |
| | 0.05 | -0.01 | 0.00 | | 0.05 | -0.02 | 0.00 |
| | 0.1 | -0.01 | 0.00 | | 0.1 | -0.03 | 0.01 |
| | 0.25 | -0.01 | 0.01 | | 0.25 | -0.05 | 0.01 |
| | 0.5 | -0.03 | 0.01 | | 0.5 | -0.05 | 0.02 |
| | 1 | -0.04 | 0.02 | | 1 | -0.06 | 0.02 |
| | 2 | -0.01 | 0.03 | | 2 | -0.06 | 0.03 |
| | 3 | -0.01 | 0.03 | | 3 | -0.09 | 0.03 |
| | 5 | -0.03 | 0.03 | | 5 | -0.09 | 0.03 |
| | 10 | -0.04 | 0.03 | | 10 | -0.03 | 0.03 |
| | 20 | 0.00 | 0.03 | | 20 | -0.05 | 0.03 |
| | 30 | -0.04 | 0.03 | | 30 | -0.07 | 0.03 |
| | 50 | 0.04 | 0.03 | | 50 | -0.09 | 0.03 |
| | 100 | 0.02 | 0.03 | | 100 | -0.04 | 0.03 |
| | 300 | -0.04 | 0.03 | | 300 | -0.06 | 0.03 |
| | 500 | 0.01 | 0.03 | | 500 | -0.04 | 0.03 |
| | 1000 | -0.02 | 0.03 | | 1000 | 0.01 | 0.03 |

**Table S5A.2**: The average differences in the donor-acceptor distances of HBs 1-5 following the shown P1′ substitutions in s01 and s05. Positive and negative differences with magnitudes > 0.1 Å are in green and red, respectively.

| HB1-5 | s01: P1' S->A | | | | s05: P1' N->A | | |
|---|---|---|---|---|---|---|---|
| **HB1** | t (ps) | Av Diff (Å) | SEM (Å) | | t (ps) | Av Diff (Å) | SEM (Å) |
| | 0.05 | 0.00 | 0.00 | | 0.05 | 0.00 | 0.00 |
| | 0.1 | 0.00 | 0.00 | | 0.1 | 0.00 | 0.00 |
| | 0.25 | 0.00 | 0.00 | | 0.25 | 0.01 | 0.00 |
| | 0.5 | 0.00 | 0.00 | | 0.5 | 0.00 | 0.01 |
| | 1 | 0.00 | 0.01 | | 1 | -0.02 | 0.02 |
| | 2 | 0.00 | 0.02 | | 2 | -0.01 | 0.03 |
| | 3 | 0.02 | 0.02 | | 3 | -0.01 | 0.03 |
| | 5 | -0.02 | 0.02 | | 5 | 0.02 | 0.03 |
| | 10 | 0.06 | 0.03 | | 10 | 0.02 | 0.03 |
| | 20 | 0.07 | 0.03 | | 20 | -0.01 | 0.03 |
| | 30 | 0.04 | 0.04 | | 30 | 0.02 | 0.03 |
| | 50 | 0.01 | 0.02 | | 50 | 0.04 | 0.03 |
| | 100 | 0.01 | 0.03 | | 100 | 0.01 | 0.04 |
| | 300 | 0.05 | 0.03 | | 300 | 0.00 | 0.04 |
| | 500 | 0.00 | 0.04 | | 500 | -0.01 | 0.04 |
| | 1000 | -0.09 | 0.04 | | 1000 | 0.08 | 0.06 |
| **HB2** | t (ps) | Av Diff (Å) | SEM (Å) | | t (ps) | Av Diff (Å) | SEM (Å) |
| | 0.05 | 0.00 | 0.00 | | 0.05 | 0.00 | 0.00 |
| | 0.1 | 0.00 | 0.00 | | 0.1 | 0.00 | 0.00 |
| | 0.25 | 0.01 | 0.00 | | 0.25 | 0.01 | 0.00 |
| | 0.5 | 0.00 | 0.00 | | 0.5 | 0.01 | 0.00 |
| | 1 | 0.00 | 0.01 | | 1 | 0.01 | 0.01 |
| | 2 | -0.01 | 0.02 | | 2 | 0.00 | 0.02 |
| | 3 | 0.01 | 0.02 | | 3 | -0.02 | 0.02 |
| | 5 | -0.01 | 0.02 | | 5 | -0.05 | 0.02 |
| | 10 | 0.00 | 0.02 | | 10 | 0.03 | 0.02 |
| | 20 | -0.01 | 0.02 | | 20 | 0.00 | 0.02 |
| | 30 | -0.03 | 0.02 | | 30 | -0.01 | 0.02 |
| | 50 | 0.00 | 0.02 | | 50 | 0.01 | 0.02 |
| | 100 | 0.01 | 0.02 | | 100 | 0.01 | 0.02 |
| | 300 | 0.00 | 0.02 | | 300 | -0.01 | 0.02 |
| | 500 | 0.02 | 0.02 | | 500 | 0.00 | 0.02 |
| | 1000 | -0.02 | 0.02 | | 1000 | -0.02 | 0.02 |
| **HB3** | t (ps) | Av Diff (Å) | SEM (Å) | | t (ps) | Av Diff (Å) | SEM (Å) |
| | 0.05 | 0.00 | 0.00 | | 0.05 | 0.00 | 0.00 |
| | 0.1 | 0.00 | 0.00 | | 0.1 | 0.01 | 0.00 |
| | 0.25 | 0.00 | 0.00 | | 0.25 | 0.02 | 0.00 |
| | 0.5 | 0.00 | 0.00 | | 0.5 | 0.02 | 0.01 |
| | 1 | 0.01 | 0.01 | | 1 | 0.01 | 0.01 |
| | 2 | -0.02 | 0.01 | | 2 | 0.00 | 0.02 |
| | 3 | 0.00 | 0.01 | | 3 | 0.01 | 0.01 |
| | 5 | -0.02 | 0.01 | | 5 | -0.02 | 0.02 |
| | 10 | 0.01 | 0.01 | | 10 | 0.03 | 0.01 |
| | 20 | -0.04 | 0.01 | | 20 | -0.02 | 0.01 |
| | 30 | 0.02 | 0.01 | | 30 | 0.02 | 0.01 |
| | 50 | -0.01 | 0.01 | | 50 | 0.02 | 0.01 |
| | 100 | 0.02 | 0.01 | | 100 | 0.02 | 0.01 |
| | 300 | 0.01 | 0.01 | | 300 | 0.03 | 0.01 |
| | 500 | 0.02 | 0.01 | | 500 | 0.03 | 0.01 |
| | 1000 | 0.00 | 0.01 | | 1000 | 0.01 | 0.02 |
| **HB4** | t (ps) | Av Diff (Å) | SEM (Å) | | t (ps) | Av Diff (Å) | SEM (Å) |
| | 0.05 | 0.00 | 0.00 | | 0.05 | 0.00 | 0.00 |
| | 0.1 | 0.00 | 0.00 | | 0.1 | 0.01 | 0.00 |
| | 0.25 | 0.01 | 0.00 | | 0.25 | 0.01 | 0.01 |
| | 0.5 | -0.01 | 0.01 | | 0.5 | 0.00 | 0.01 |
| | 1 | 0.01 | 0.03 | | 1 | 0.01 | 0.02 |
| | 2 | 0.06 | 0.03 | | 2 | -0.05 | 0.03 |
| | 3 | 0.05 | 0.03 | | 3 | 0.01 | 0.03 |
| | 5 | -0.01 | 0.03 | | 5 | 0.01 | 0.03 |
| | 10 | -0.04 | 0.04 | | 10 | -0.07 | 0.04 |
| | 20 | -0.03 | 0.04 | | 20 | -0.10 | 0.06 |
| | 30 | -0.05 | 0.06 | | 30 | -0.09 | 0.06 |
| | 50 | -0.09 | 0.07 | | 50 | -0.11 | 0.06 |
| | 100 | -0.06 | 0.09 | | 100 | -0.14 | 0.08 |
| | 300 | -0.16 | 0.11 | | 300 | 0.19 | 0.12 |
| | 500 | -0.33 | 0.13 | | 500 | 0.33 | 0.12 |
| | 1000 | -0.35 | 0.15 | | 1000 | -0.05 | 0.14 |
| **HB5** | t (ps) | Av Diff (Å) | SEM (Å) | | t (ps) | Av Diff (Å) | SEM (Å) |
| | 0.05 | 0.00 | 0.00 | | 0.05 | 0.00 | 0.00 |
| | 0.1 | 0.00 | 0.00 | | 0.1 | 0.01 | 0.00 |
| | 0.25 | 0.00 | 0.00 | | 0.25 | 0.01 | 0.00 |
| | 0.5 | 0.00 | 0.01 | | 0.5 | 0.02 | 0.01 |
| | 1 | -0.03 | 0.02 | | 1 | -0.02 | 0.02 |
| | 2 | -0.03 | 0.03 | | 2 | 0.01 | 0.02 |
| | 3 | -0.03 | 0.03 | | 3 | 0.03 | 0.03 |
| | 5 | -0.01 | 0.03 | | 5 | 0.05 | 0.03 |
| | 10 | 0.03 | 0.03 | | 10 | 0.00 | 0.02 |
| | 20 | 0.00 | 0.03 | | 20 | 0.00 | 0.03 |
| | 30 | 0.03 | 0.03 | | 30 | 0.02 | 0.02 |
| | 50 | 0.03 | 0.03 | | 50 | 0.02 | 0.03 |
| | 100 | 0.02 | 0.03 | | 100 | -0.01 | 0.03 |
| | 300 | 0.00 | 0.03 | | 300 | 0.11 | 0.03 |
| | 500 | -0.01 | 0.04 | | 500 | 0.05 | 0.03 |
| | 1000 | 0.04 | 0.04 | | 1000 | -0.03 | 0.03 |

**Table S5A.3**: The average differences in the donor-acceptor distances of HBs 8-12 following the shown P1′ substitutions in s01 and s05. Positive and negative differences with magnitudes > 0.1 Å are in green and red, respectively.

| HB8-12 | s01: P1' S->A | | | | s05: P1' N->A | | |
|---|---|---|---|---|---|---|---|
| **HB8** | t (ps) | Av Diff (Å) | SEM (Å) | | t (ps) | Av Diff (Å) | SEM (Å) |
| | 0.05 | 0.01 | 0.00 | | 0.05 | 0.01 | 0.00 |
| | 0.1 | 0.00 | 0.00 | | 0.1 | 0.02 | 0.00 |
| | 0.25 | 0.00 | 0.01 | | 0.25 | 0.05 | 0.01 |
| | 0.5 | 0.03 | 0.01 | | 0.5 | 0.09 | 0.02 |
| | 1 | 0.01 | 0.02 | | 1 | 0.06 | 0.02 |
| | 2 | -0.01 | 0.03 | | 2 | 0.09 | 0.03 |
| | 3 | 0.04 | 0.03 | | 3 | 0.08 | 0.03 |
| | 5 | 0.04 | 0.03 | | 5 | 0.09 | 0.03 |
| | 10 | 0.06 | 0.03 | | 10 | 0.06 | 0.03 |
| | 20 | -0.03 | 0.03 | | 20 | 0.07 | 0.03 |
| | 30 | 0.03 | 0.03 | | 30 | 0.08 | 0.03 |
| | 50 | -0.02 | 0.03 | | 50 | 0.09 | 0.03 |
| | 100 | -0.04 | 0.03 | | 100 | 0.06 | 0.03 |
| | 300 | -0.02 | 0.03 | | 300 | 0.09 | 0.03 |
| | 500 | -0.06 | 0.03 | | 500 | 0.09 | 0.03 |
| | 1000 | 0.02 | 0.03 | | 1000 | 0.04 | 0.03 |
| **HB9** | t (ps) | Av Diff (Å) | SEM (Å) | | t (ps) | Av Diff (Å) | SEM (Å) |
| | 0.05 | -0.01 | 0.00 | | 0.05 | -0.01 | 0.00 |
| | 0.1 | -0.01 | 0.00 | | 0.1 | -0.02 | 0.00 |
| | 0.25 | -0.01 | 0.00 | | 0.25 | -0.03 | 0.01 |
| | 0.5 | -0.01 | 0.01 | | 0.5 | -0.02 | 0.01 |
| | 1 | -0.01 | 0.01 | | 1 | -0.04 | 0.01 |
| | 2 | 0.00 | 0.02 | | 2 | -0.03 | 0.02 |
| | 3 | -0.01 | 0.02 | | 3 | -0.05 | 0.02 |
| | 5 | -0.01 | 0.02 | | 5 | -0.05 | 0.02 |
| | 10 | -0.02 | 0.02 | | 10 | -0.01 | 0.02 |
| | 20 | 0.00 | 0.02 | | 20 | -0.02 | 0.02 |
| | 30 | -0.02 | 0.02 | | 30 | -0.03 | 0.02 |
| | 50 | 0.02 | 0.02 | | 50 | -0.04 | 0.02 |
| | 100 | 0.01 | 0.02 | | 100 | -0.02 | 0.02 |
| | 300 | -0.02 | 0.02 | | 300 | -0.03 | 0.02 |
| | 500 | 0.00 | 0.02 | | 500 | -0.02 | 0.02 |
| | 1000 | -0.01 | 0.02 | | 1000 | 0.01 | 0.02 |
| **HB10** | t (ps) | Av Diff (Å) | SEM (Å) | | t (ps) | Av Diff (Å) | SEM (Å) |
| | 0.05 | 0.01 | 0.00 | | 0.05 | 0.00 | 0.00 |
| | 0.1 | 0.00 | 0.00 | | 0.1 | 0.00 | 0.00 |
| | 0.25 | 0.01 | 0.00 | | 0.25 | 0.02 | 0.01 |
| | 0.5 | 0.01 | 0.01 | | 0.5 | 0.02 | 0.01 |
| | 1 | 0.00 | 0.01 | | 1 | 0.00 | 0.01 |
| | 2 | 0.03 | 0.02 | | 2 | 0.01 | 0.02 |
| | 3 | 0.02 | 0.02 | | 3 | 0.03 | 0.02 |
| | 5 | 0.00 | 0.02 | | 5 | 0.03 | 0.02 |
| | 10 | 0.00 | 0.02 | | 10 | 0.02 | 0.02 |
| | 20 | -0.02 | 0.02 | | 20 | 0.01 | 0.02 |
| | 30 | -0.01 | 0.02 | | 30 | 0.04 | 0.02 |
| | 50 | 0.02 | 0.02 | | 50 | 0.04 | 0.02 |
| | 100 | 0.00 | 0.02 | | 100 | 0.02 | 0.02 |
| | 300 | -0.01 | 0.02 | | 300 | 0.02 | 0.02 |
| | 500 | -0.03 | 0.02 | | 500 | -0.02 | 0.02 |
| | 1000 | -0.01 | 0.02 | | 1000 | 0.05 | 0.02 |
| **HB11** | t (ps) | Av Diff (Å) | SEM (Å) | | t (ps) | Av Diff (Å) | SEM (Å) |
| | 0.05 | 0.00 | 0.00 | | 0.05 | -0.02 | 0.00 |
| | 0.1 | 0.01 | 0.00 | | 0.1 | -0.06 | 0.01 |
| | 0.25 | 0.00 | 0.01 | | 0.25 | -0.05 | 0.01 |
| | 0.5 | 0.01 | 0.01 | | 0.5 | -0.06 | 0.01 |
| | 1 | -0.02 | 0.02 | | 1 | -0.07 | 0.02 |
| | 2 | -0.02 | 0.02 | | 2 | -0.07 | 0.01 |
| | 3 | 0.00 | 0.02 | | 3 | -0.04 | 0.02 |
| | 5 | -0.01 | 0.02 | | 5 | -0.04 | 0.02 |
| | 10 | 0.01 | 0.02 | | 10 | -0.03 | 0.02 |
| | 20 | 0.01 | 0.03 | | 20 | -0.06 | 0.02 |
| | 30 | -0.02 | 0.02 | | 30 | -0.05 | 0.02 |
| | 50 | 0.02 | 0.03 | | 50 | -0.01 | 0.02 |
| | 100 | 0.00 | 0.02 | | 100 | -0.04 | 0.02 |
| | 300 | 0.01 | 0.02 | | 300 | -0.03 | 0.02 |
| | 500 | -0.03 | 0.02 | | 500 | -0.01 | 0.02 |
| | 1000 | 0.02 | 0.02 | | 1000 | 0.00 | 0.02 |
| **HB12** | t (ps) | Av Diff (Å) | SEM (Å) | | t (ps) | Av Diff (Å) | SEM (Å) |
| | 0.05 | 0.00 | 0.00 | | 0.05 | 0.00 | 0.00 |
| | 0.1 | 0.00 | 0.00 | | 0.1 | 0.00 | 0.00 |
| | 0.25 | -0.01 | 0.00 | | 0.25 | 0.01 | 0.00 |
| | 0.5 | -0.02 | 0.01 | | 0.5 | 0.01 | 0.01 |
| | 1 | 0.00 | 0.02 | | 1 | 0.01 | 0.02 |
| | 2 | -0.01 | 0.02 | | 2 | -0.02 | 0.02 |
| | 3 | 0.06 | 0.03 | | 3 | 0.03 | 0.02 |
| | 5 | 0.00 | 0.03 | | 5 | -0.04 | 0.03 |
| | 10 | -0.05 | 0.03 | | 10 | 0.05 | 0.03 |
| | 20 | 0.03 | 0.03 | | 20 | 0.05 | 0.03 |
| | 30 | -0.02 | 0.03 | | 30 | 0.04 | 0.04 |
| | 50 | -0.08 | 0.03 | | 50 | 0.03 | 0.03 |
| | 100 | 0.02 | 0.04 | | 100 | 0.00 | 0.03 |
| | 300 | -0.05 | 0.05 | | 300 | 0.03 | 0.03 |
| | 500 | 0.02 | 0.04 | | 500 | 0.01 | 0.03 |
| | 1000 | -0.06 | 0.05 | | 1000 | 0.04 | 0.04 |

**Table S5A.4**: The ten most perturbed Cα atoms in the M$^{pro}$-peptide complex in terms of magnitude of the average displacement vector $v$, at selected time points following the P1′ substitutions to Ala from Ser and Asn in s01 and s05, respectively. Only displacements ≥ 0.01 Å are shown.

| s01 t(ps) | CA Rank | Atom | Chain | v (Å) | SEM (Å) |
|---|---|---|---|---|---|
| **0.05** | 1 | 7ALA CA | C | 0.02 | 0.00 |
| | 2 | 8GLY_CA | C | 0.01 | 0.00 |
| | 3 | | | | |
| | 4 | | | | |
| | 5 | | | | |
| | 6 | | | | |
| | 7 | | | | |
| | 8 | | | | |
| | 9 | | | | |
| | 10 | | | | |
| **0.5** | 1 | 27LEU_CA | A | 0.02 | 0.00 |
| | 2 | 42VAL_CA | A | 0.02 | 0.00 |
| | 3 | 46SER_CA | A | 0.02 | 0.01 |
| | 4 | 187ASP_CA | A | 0.02 | 0.00 |
| | 5 | 5LEU_CA | C | 0.02 | 0.01 |
| | 6 | 7ALA_CA | C | 0.02 | 0.01 |
| | 7 | 8GLY_CA | C | 0.02 | 0.01 |
| | 8 | 19GLN_CA | A | 0.01 | 0.00 |
| | 9 | 20VAL_CA | A | 0.01 | 0.00 |
| | 10 | 22CYS_CA | A | 0.01 | 0.00 |
| **5** | 1 | 72ASN_CA | B | 0.17 | 0.05 |
| | 2 | 277ASN_CA | B | 0.15 | 0.05 |
| | 3 | 71GLY_CA | B | 0.12 | 0.06 |
| | 4 | 196THR_CA | B | 0.12 | 0.05 |
| | 5 | 218TRP_CA | B | 0.12 | 0.04 |
| | 6 | 278GLY_CA | B | 0.12 | 0.05 |
| | 7 | 219PHE_CA | B | 0.11 | 0.03 |
| | 8 | 275GLY_CA | B | 0.11 | 0.05 |
| | 9 | 279ARG_CA | B | 0.11 | 0.04 |
| | 10 | 283GLY_CA | B | 0.11 | 0.04 |
| **50** | 1 | 72ASN CA | B | 0.25 | 0.08 |
| | 2 | 191ALA_CA | B | 0.23 | 0.08 |
| | 3 | 232LEU_CA | A | 0.19 | 0.06 |
| | 4 | 1THR_CA | C | 0.19 | 0.09 |
| | 5 | 73VAL_CA | B | 0.18 | 0.07 |
| | 6 | 142ASN_CA | B | 0.14 | 0.04 |
| | 7 | 221ASN_CA | B | 0.14 | 0.05 |
| | 8 | 225THR_CA | B | 0.14 | 0.06 |
| | 9 | 226THR_CA | B | 0.14 | 0.06 |
| | 10 | 83GLN_CA | A | 0.13 | 0.04 |
| **500** | 1 | 274ASN_CA | A | 0.24 | 0.09 |
| | 2 | 72ASN_CA | B | 0.24 | 0.10 |
| | 3 | 73VAL_CA | B | 0.24 | 0.08 |
| | 4 | 277ASN_CA | B | 0.24 | 0.10 |
| | 5 | 278GLY_CA | B | 0.22 | 0.11 |
| | 6 | 191ALA_CA | B | 0.21 | 0.09 |
| | 7 | 59ILE_CA | A | 0.20 | 0.06 |
| | 8 | 275GLY_CA | A | 0.20 | 0.08 |
| | 9 | 74GLN_CA | B | 0.20 | 0.06 |
| | 10 | 169THR_CA | B | 0.20 | 0.07 |
| **1000** | 1 | 244GLN_CA | A | 0.24 | 0.08 |
| | 2 | 236LYS_CA | A | 0.23 | 0.08 |
| | 3 | 11LYS_CA | C | 0.23 | 0.14 |
| | 4 | 235MET_CA | A | 0.22 | 0.08 |
| | 5 | 245ASP_CA | A | 0.22 | 0.07 |
| | 6 | 51ASN_CA | A | 0.21 | 0.07 |
| | 7 | 232LEU_CA | A | 0.21 | 0.07 |
| | 8 | 283GLY_CA | A | 0.21 | 0.07 |
| | 9 | 243THR_CA | A | 0.20 | 0.07 |
| | 10 | 92ASP_CA | A | 0.19 | 0.07 |

| s05 t(ps) | CA Rank | Atom | Chain | v (Å) | SEM (Å) |
|---|---|---|---|---|---|
| **0.05** | 1 | 25THR CA | A | 0.01 | 0.00 |
| | 2 | 6GLN_CA | C | 0.01 | 0.00 |
| | 3 | 7ALA_CA | C | 0.01 | 0.00 |
| | 4 | 8ASN_CA | C | 0.01 | 0.00 |
| | 5 | | | | |
| | 6 | | | | |
| | 7 | | | | |
| | 8 | | | | |
| | 9 | | | | |
| | 10 | | | | |
| **0.5** | 1 | 7ALA_CA | C | 0.11 | 0.02 |
| | 2 | 44CYS_CA | A | 0.07 | 0.01 |
| | 3 | 6GLN_CA | C | 0.07 | 0.01 |
| | 4 | 41HID_CA | A | 0.06 | 0.01 |
| | 5 | 43ILE_CA | A | 0.06 | 0.01 |
| | 6 | 5LEU_CA | C | 0.06 | 0.01 |
| | 7 | 8ASN_CA | C | 0.06 | 0.01 |
| | 8 | 9GLU_CA | C | 0.06 | 0.01 |
| | 9 | 42VAL_CA | A | 0.05 | 0.01 |
| | 10 | 49MET_CA | A | 0.05 | 0.01 |
| **5** | 1 | 285ALA_CA | B | 0.16 | 0.03 |
| | 2 | 51ASN_CA | B | 0.15 | 0.04 |
| | 3 | 286LEU_CA | B | 0.15 | 0.03 |
| | 4 | 49MET_CA | A | 0.13 | 0.04 |
| | 5 | 72ASN_CA | A | 0.13 | 0.05 |
| | 6 | 284SER_CA | B | 0.13 | 0.03 |
| | 7 | 44CYS_CA | A | 0.12 | 0.03 |
| | 8 | 50LEU_CA | A | 0.12 | 0.04 |
| | 9 | 64HIE_CA | A | 0.12 | 0.04 |
| | 10 | 235MET_CA | A | 0.12 | 0.04 |
| **50** | 1 | 72ASN CA | A | 0.29 | 0.08 |
| | 2 | 275GLY_CA | B | 0.22 | 0.06 |
| | 3 | 274ASN_CA | B | 0.21 | 0.07 |
| | 4 | 277ASN_CA | B | 0.21 | 0.09 |
| | 5 | 71GLY_CA | A | 0.20 | 0.08 |
| | 6 | 236LYS_CA | A | 0.20 | 0.06 |
| | 7 | 24THR_CA | B | 0.20 | 0.07 |
| | 8 | 72ASN_CA | B | 0.18 | 0.08 |
| | 9 | 64HIE_CA | A | 0.17 | 0.06 |
| | 10 | 227LEU_CA | A | 0.17 | 0.05 |
| **500** | 1 | 278GLY_CA | A | 0.33 | 0.10 |
| | 2 | 10LEU_CA | C | 0.32 | 0.09 |
| | 3 | 277ASN_CA | A | 0.30 | 0.11 |
| | 4 | 72ASN_CA | A | 0.28 | 0.10 |
| | 5 | 244GLN_CA | B | 0.28 | 0.07 |
| | 6 | 9GLU_CA | C | 0.27 | 0.07 |
| | 7 | 245ASP_CA | B | 0.25 | 0.07 |
| | 8 | 274ASN_CA | A | 0.24 | 0.09 |
| | 9 | 243THR_CA | B | 0.24 | 0.07 |
| | 10 | 11SER_CA | C | 0.24 | 0.16 |
| **1000** | 1 | 11SER_CA | C | 0.39 | 0.18 |
| | 2 | 59ILE_CA | B | 0.21 | 0.09 |
| | 3 | 60ARG_CA | B | 0.21 | 0.09 |
| | 4 | 10LEU_CA | C | 0.21 | 0.09 |
| | 5 | 1SER_CA | C | 0.20 | 0.12 |
| | 6 | 15GLY_CA | A | 0.19 | 0.05 |
| | 7 | 222ARG_CA | B | 0.19 | 0.10 |
| | 8 | 234ALA_CA | A | 0.18 | 0.06 |
| | 9 | 215GLY_CA | B | 0.18 | 0.10 |
| | 10 | 9GLU_CA | C | 0.17 | 0.06 |

**Table S5A.5**: The ten most perturbed non-hydrogen atoms in the M$^{pro}$-peptide complex in terms of magnitude of the average displacement vector *v*, at selected time points following the P1′ substitutions to Ala from Ser and Asn in s01 and s05, respectively.

| *s01 t(ps)* | *Non-H Rank* | *Atom* | *Chain* | *v (Å)* | *SEM (Å)* |
|---|---|---|---|---|---|
| **0.05** | 1 | 7ALA_CB | C | 0.06 | 0.01 |
| | 2 | 5LEU_O | C | 0.02 | 0.00 |
| | 3 | 26THR_O | A | 0.02 | 0.00 |
| | 4 | 7ALA_CA | C | 0.02 | 0.00 |
| | 5 | 6GLN_O | C | 0.01 | 0.00 |
| | 6 | 8GLY_N | C | 0.01 | 0.00 |
| | 7 | 7ALA_C | C | 0.01 | 0.00 |
| | 8 | 7ALA_N | C | 0.01 | 0.00 |
| | 9 | 41HID_NE2 | A | 0.01 | 0.00 |
| | 10 | 7ALA_O | C | 0.01 | 0.00 |
| **0.5** | 1 | 7ALA_CB | C | 0.08 | 0.02 |
| | 2 | 41HID_NE2 | A | 0.08 | 0.02 |
| | 3 | 41HID_CE1 | A | 0.08 | 0.02 |
| | 4 | 49MET_CE | A | 0.07 | 0.02 |
| | 5 | 5LEU_O | C | 0.07 | 0.02 |
| | 6 | 41HID_ND1 | A | 0.06 | 0.01 |
| | 7 | 41HID_CD2 | A | 0.06 | 0.01 |
| | 8 | 7ALA_O | C | 0.06 | 0.02 |
| | 9 | 26THR_O | A | 0.06 | 0.01 |
| | 10 | 25THR_CG2 | A | 0.05 | 0.02 |
| **5** | 1 | 72ASN_ND2 | B | 0.30 | 0.09 |
| | 2 | 72ASN_CG | B | 0.26 | 0.08 |
| | 3 | 72ASN_CB | B | 0.24 | 0.07 |
| | 4 | 10ARG_NH2 | C | 0.23 | 0.07 |
| | 5 | 306GLN_OE1 | B | 0.23 | 0.07 |
| | 6 | 72ASN_OD1 | B | 0.22 | 0.09 |
| | 7 | 294PHE_CZ | A | 0.21 | 0.07 |
| | 8 | 277ASN_O | B | 0.21 | 0.06 |
| | 9 | 53ASN_OD1 | B | 0.20 | 0.07 |
| | 10 | 47GLU_OE2 | B | 0.20 | 0.09 |
| **50** | 1 | 222ARG_NH2 | B | 0.42 | 0.19 |
| | 2 | 235MET_CE | A | 0.39 | 0.15 |
| | 3 | 55GLU_OE1 | B | 0.39 | 0.13 |
| | 4 | 222ARG_NH1 | A | 0.38 | 0.19 |
| | 5 | 72ASN_ND2 | B | 0.37 | 0.15 |
| | 6 | 222ARG_NH1 | B | 0.36 | 0.19 |
| | 7 | 11LYS_NZ | C | 0.36 | 0.19 |
| | 8 | 142ASN_OD1 | B | 0.33 | 0.10 |
| | 9 | 217ARG_NH1 | B | 0.32 | 0.12 |
| | 10 | 191ALA_CB | B | 0.32 | 0.10 |
| **500** | 1 | 222ARG_NH2 | B | 0.77 | 0.22 |
| | 2 | 64HIE_NE2 | B | 0.69 | 0.15 |
| | 3 | 222ARG_NH1 | B | 0.69 | 0.22 |
| | 4 | 189GLN_NE2 | B | 0.61 | 0.14 |
| | 5 | 222ARG_CZ | B | 0.59 | 0.18 |
| | 6 | 222ARG_NH2 | A | 0.57 | 0.23 |
| | 7 | 64HIE_CD2 | B | 0.56 | 0.12 |
| | 8 | 64HIE_CE1 | B | 0.55 | 0.15 |
| | 9 | 107GLN_OE1 | B | 0.49 | 0.17 |
| | 10 | 189GLN_OE1 | B | 0.49 | 0.15 |
| **1000** | 1 | 236LYS_NZ | A | 0.84 | 0.24 |
| | 2 | 236LYS_CE | A | 0.77 | 0.21 |
| | 3 | 279ARG_NH1 | B | 0.70 | 0.22 |
| | 4 | 60ARG_NH2 | A | 0.63 | 0.19 |
| | 5 | 47GLU_OE2 | B | 0.60 | 0.22 |
| | 6 | 222ARG_NH1 | B | 0.60 | 0.27 |
| | 7 | 236LYS_CD | A | 0.57 | 0.16 |
| | 8 | 244GLN_NE2 | A | 0.56 | 0.24 |
| | 9 | 235MET_CE | B | 0.54 | 0.27 |
| | 10 | 235MET_SD | B | 0.53 | 0.19 |

| *s05 t(ps)* | *Non-H Rank* | *Atom* | *Chain* | *v (Å)* | *SEM (Å)* |
|---|---|---|---|---|---|
| **0.05** | 1 | 5LEU_O | C | 0.06 | 0.00 |
| | 2 | 8ASN_O | C | 0.04 | 0.00 |
| | 3 | 7ALA_CB | C | 0.04 | 0.01 |
| | 4 | 7ALA_O | C | 0.04 | 0.00 |
| | 5 | 4LYS_NZ | C | 0.02 | 0.00 |
| | 6 | 8ASN_N | C | 0.02 | 0.00 |
| | 7 | 25THR_CG2 | A | 0.02 | 0.00 |
| | 8 | 26THR_O | A | 0.02 | 0.00 |
| | 9 | 5LEU_C | C | 0.02 | 0.00 |
| | 10 | 49MET_CE | A | 0.02 | 0.00 |
| **0.5** | 1 | 7ALA_O | C | 0.28 | 0.03 |
| | 2 | 7ALA_C | C | 0.18 | 0.02 |
| | 3 | 49MET_CE | A | 0.17 | 0.02 |
| | 4 | 5LEU_O | C | 0.16 | 0.01 |
| | 5 | 49MET_SD | A | 0.13 | 0.02 |
| | 6 | 8ASN_O | C | 0.13 | 0.02 |
| | 7 | 41HID_NE2 | A | 0.12 | 0.01 |
| | 8 | 7ALA_CB | C | 0.12 | 0.02 |
| | 9 | 25THR_CG2 | A | 0.12 | 0.02 |
| | 10 | 7ALA_CA | C | 0.12 | 0.02 |
| **5** | 1 | 7ALA_O | C | 0.34 | 0.05 |
| | 2 | 49MET_CE | A | 0.26 | 0.04 |
| | 3 | 228ASN_OD1 | A | 0.24 | 0.08 |
| | 4 | 217ARG_NH1 | A | 0.23 | 0.08 |
| | 5 | 107GLN_NE2 | A | 0.23 | 0.08 |
| | 6 | 107GLN_NE2 | B | 0.23 | 0.09 |
| | 7 | 189GLN_NE2 | B | 0.23 | 0.07 |
| | 8 | 235MET_SD | A | 0.23 | 0.08 |
| | 9 | 41HID_NE2 | A | 0.22 | 0.03 |
| | 10 | 49MET_SD | A | 0.22 | 0.04 |
| **50** | 1 | 72ASN_OD1 | A | 0.51 | 0.14 |
| | 2 | 279ARG_NH2 | B | 0.50 | 0.11 |
| | 3 | 277ASN_ND2 | B | 0.41 | 0.13 |
| | 4 | 72ASN_CG | A | 0.40 | 0.13 |
| | 5 | 223PHE_CZ | A | 0.39 | 0.12 |
| | 6 | 222ARG_NH2 | A | 0.39 | 0.16 |
| | 7 | 7ALA_O | C | 0.38 | 0.05 |
| | 8 | 279ARG_CZ | B | 0.37 | 0.10 |
| | 9 | 72ASN_ND2 | A | 0.36 | 0.16 |
| | 10 | 72ASN_CB | A | 0.35 | 0.11 |
| **500** | 1 | 256GLN_NE2 | A | 0.69 | 0.15 |
| | 2 | 222ARG_NH2 | B | 0.62 | 0.28 |
| | 3 | 235MET_CE | B | 0.56 | 0.20 |
| | 4 | 10LEU_CD1 | C | 0.55 | 0.14 |
| | 5 | 189GLN_NE2 | B | 0.51 | 0.13 |
| | 6 | 107GLN_NE2 | B | 0.49 | 0.14 |
| | 7 | 60ARG_NH2 | A | 0.49 | 0.17 |
| | 8 | 245ASP_OD1 | B | 0.49 | 0.15 |
| | 9 | 60ARG_NH1 | A | 0.48 | 0.18 |
| | 10 | 189GLN_OE1 | B | 0.47 | 0.14 |
| **1000** | 1 | 222ARG_NH2 | B | 0.89 | 0.30 |
| | 2 | 235MET_CE | A | 0.80 | 0.24 |
| | 3 | 222ARG_CZ | B | 0.72 | 0.27 |
| | 4 | 235MET_SD | A | 0.70 | 0.19 |
| | 5 | 222ARG_NH1 | B | 0.68 | 0.31 |
| | 6 | 76ARG_NH1 | B | 0.65 | 0.22 |
| | 7 | 76ARG_NH2 | B | 0.63 | 0.23 |
| | 8 | 222ARG_NE | B | 0.62 | 0.23 |
| | 9 | 76ARG_CZ | B | 0.52 | 0.19 |
| | 10 | 11SER_OG | C | 0.51 | 0.27 |

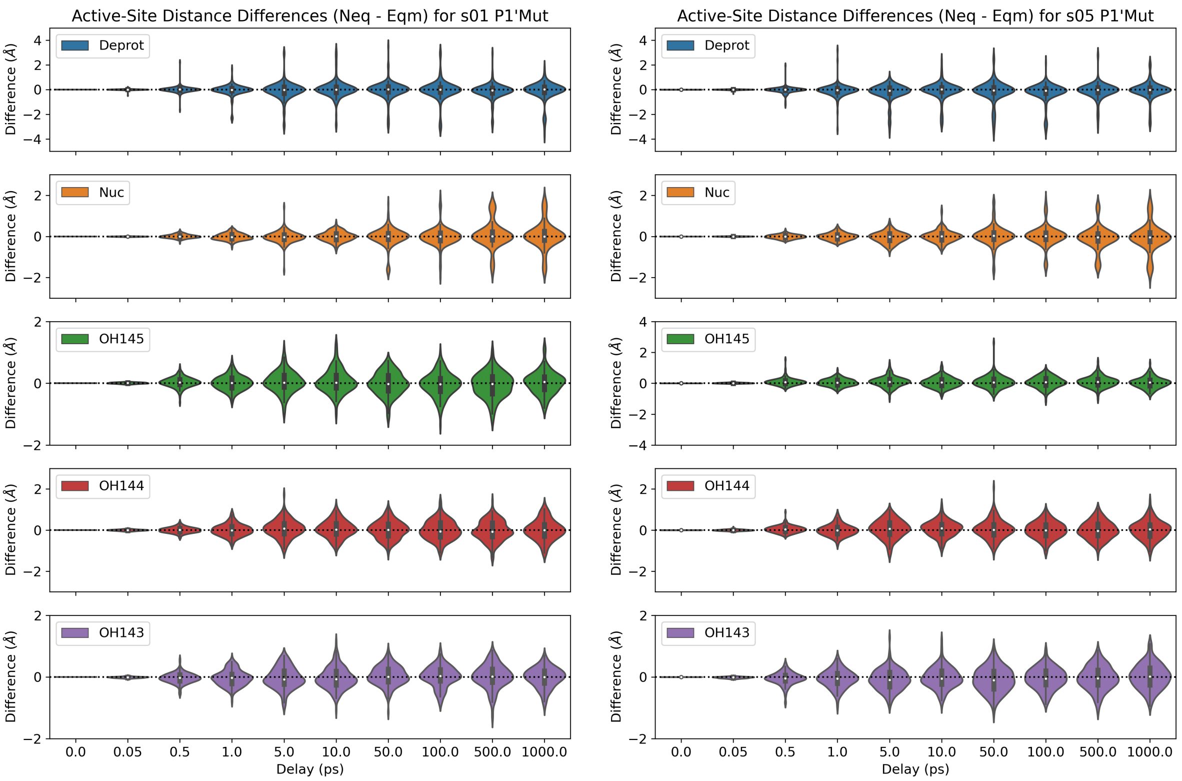

**Figure S5A.1**: Distributions of the distance differences in the five active-site distances ($\Delta d = d_{\text{neq}} - d_{\text{eqm}}$) following the P1′ substitutions to Ala from Ser and Asn in s01 and s05, respectively.

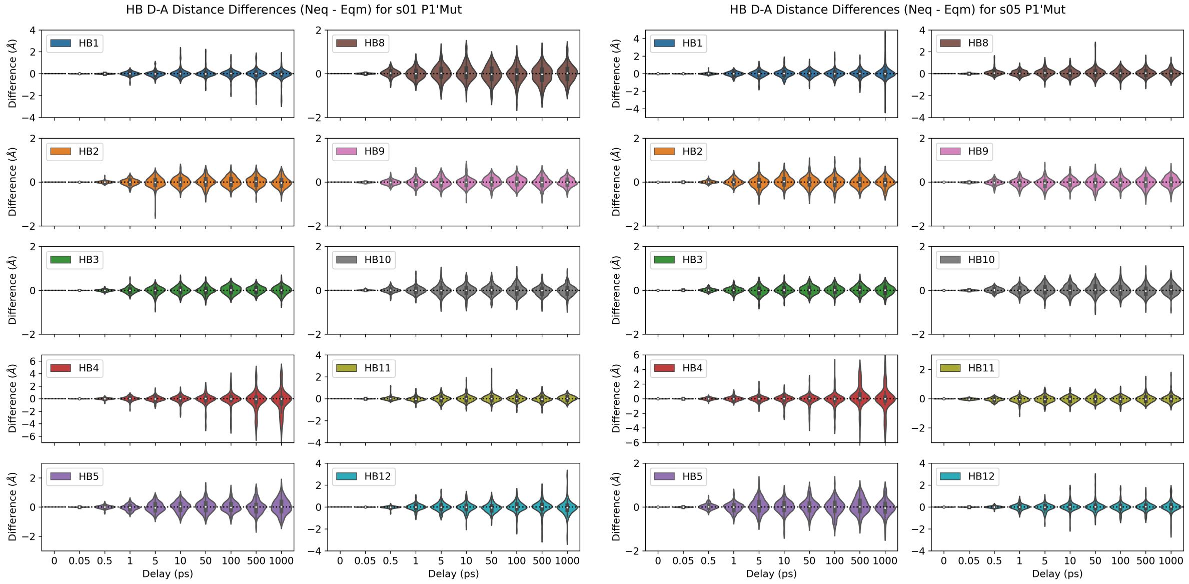

**Figure S5A.2**: Distributions of the distance differences in the ten donor-acceptor distances of HBs 1-5 and 8-12 following the P1′ substitutions to Ala from Ser and Asn in s01 and s05, respectively.

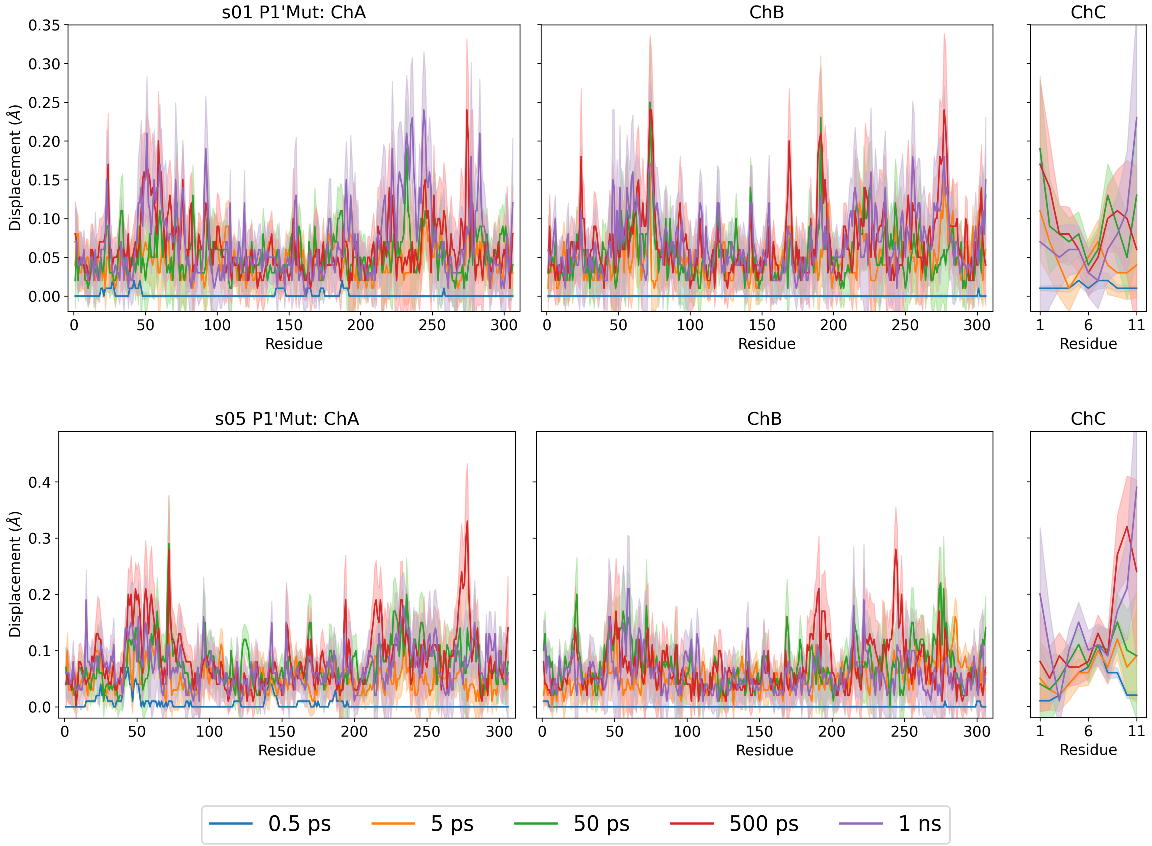

**Figure S5A.3**: Magnitudes of the average Cα displacement vectors, with ± SEM shown as shaded regions, following the P1′ substitutions to Ala from Ser and Asn in s01 and s05, respectively.

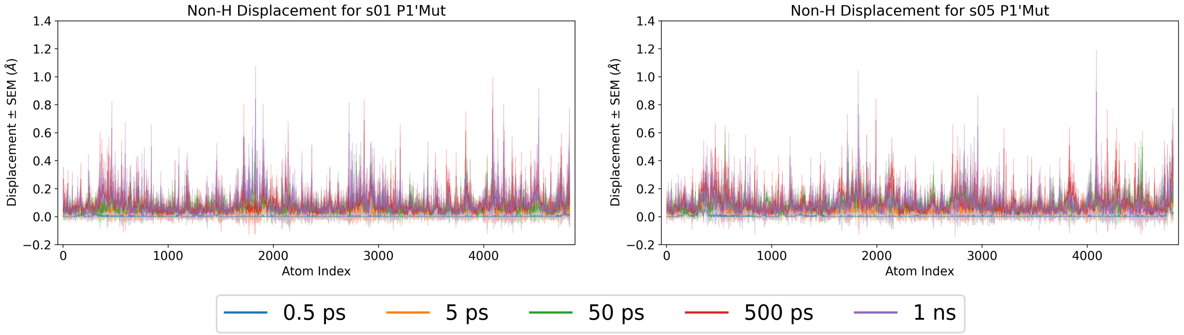

**Figure S5A.4**: Magnitudes of the average non-hydrogen atom displacement vectors in the M$^{pro}$-peptide complex, with ± SEM shown as shaded regions, following the P1′ substitutions to Ala from Ser and Asn in s01 and s05, respectively.

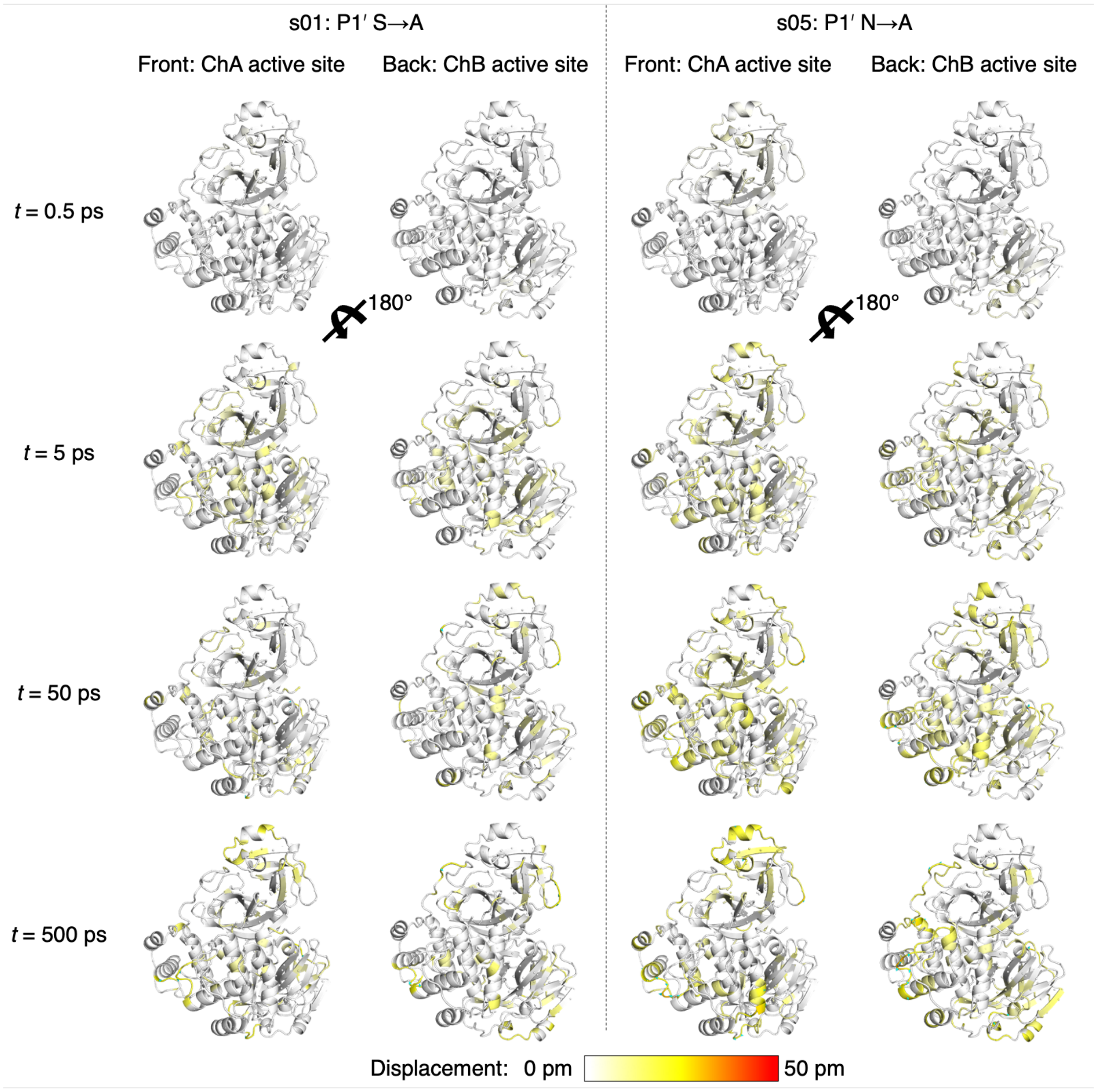

**Figure S5A.5**: Views of the M^pro response to the shown P1′ substitutions in s01 and s05 from averaging Cα displacement vectors, shown using the structure prior to MD simulations. Displacement magnitudes are represented on a white-yellow-red scale.[7] Significant vectors with length ≥20 pm are displayed as cyan arrows with a scale-up factor of 5.[8]

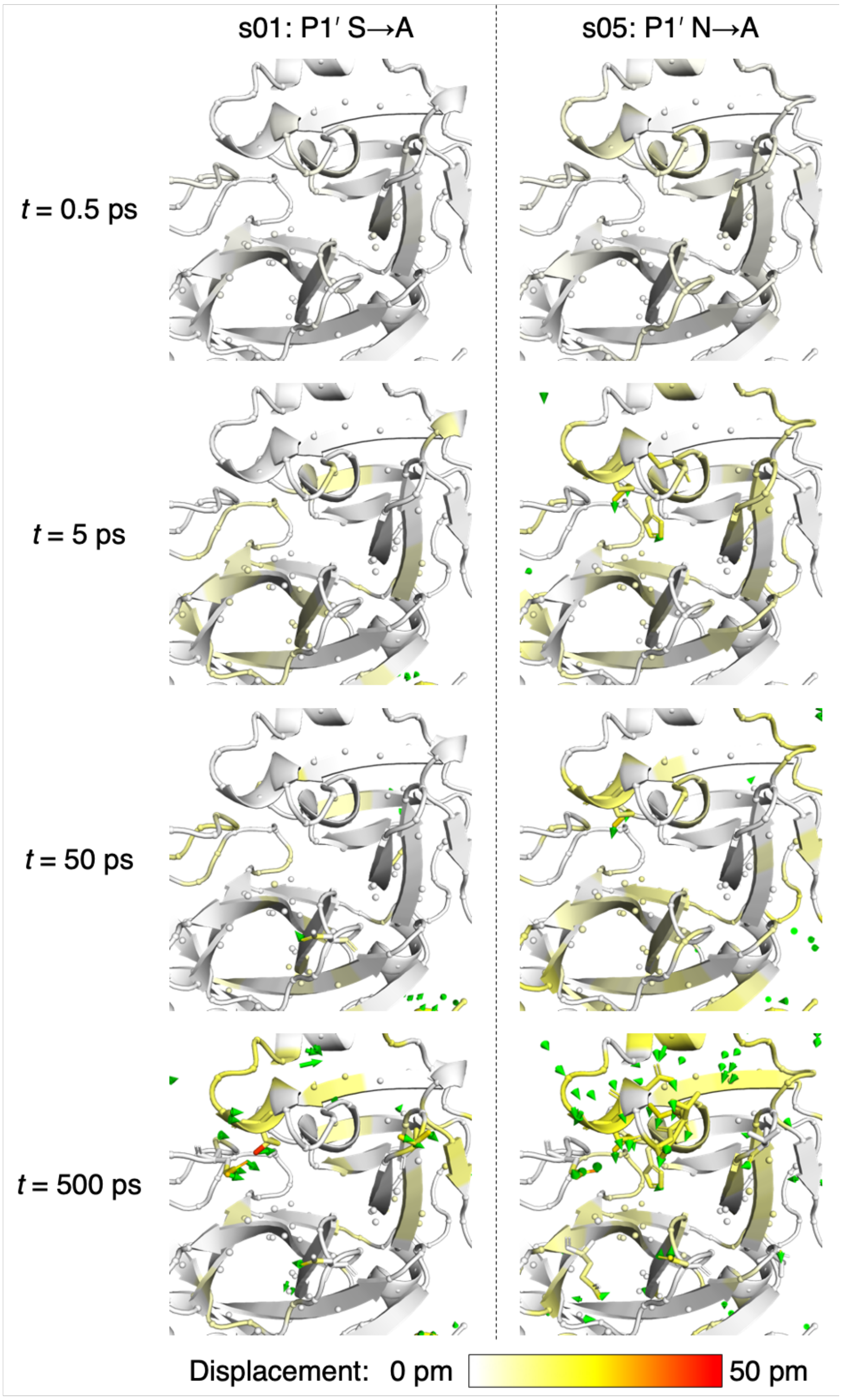

**Figure S5A.6**: Views of the M^pro response to the shown P1′ substitutions in s01 and s05 from averaging non-hydrogen atom displacement vectors, shown using the structure prior to MD simulations, with a focus on the S1′ subsite. Displacement magnitudes are represented on a white-yellow-red scale.[7] Significant vectors with length ≥20 pm are displayed as green arrows with a scale-up factor of 5.[8] Residues that show such significant displacements and that are within 10 Å of the substrate P1′ residue are shown as sticks.

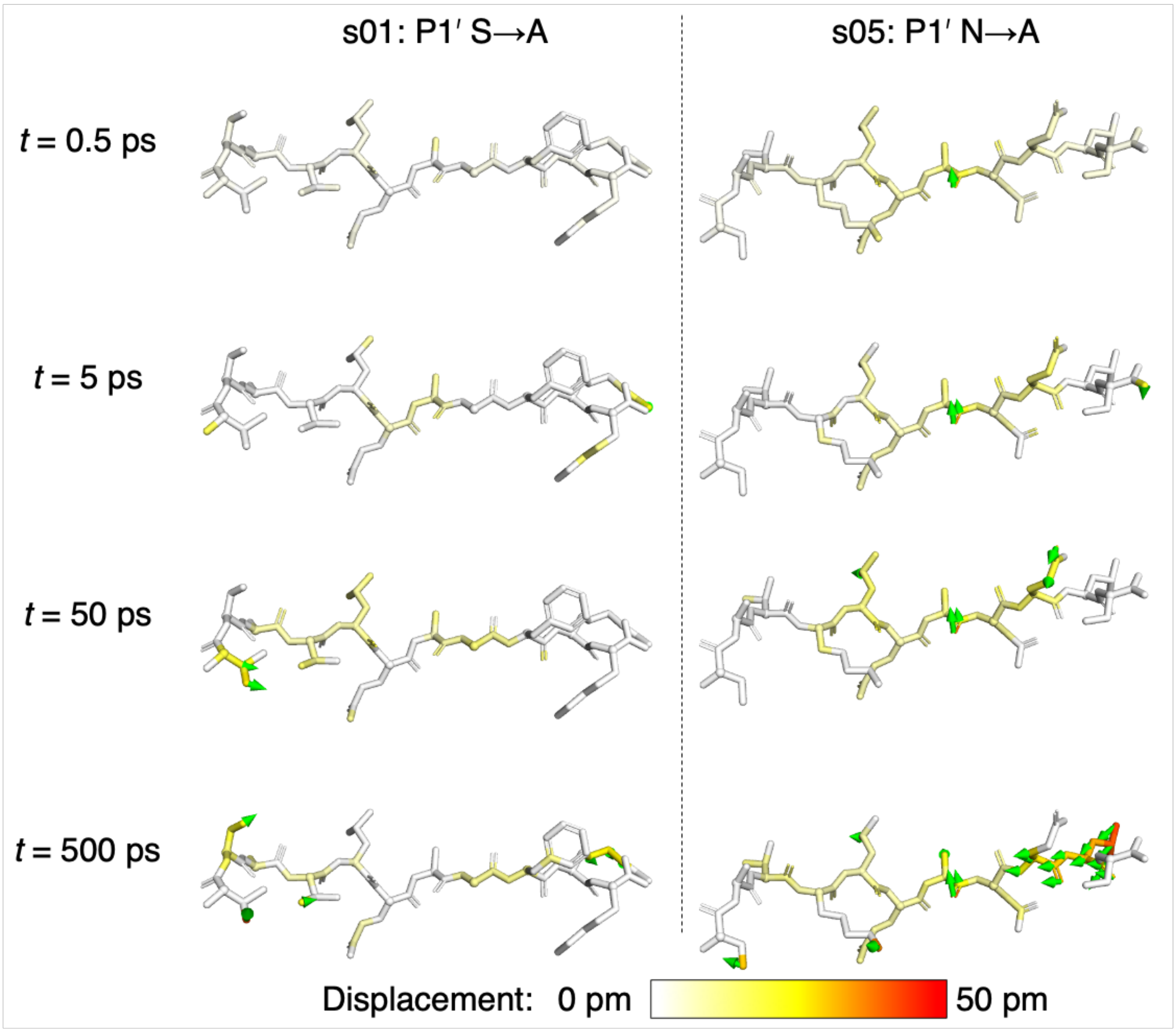

**Figure S5A.7**: Views of the peptide response to the shown P1′ substitutions in s01 and s05 from averaging non-hydrogen atom displacement vectors, shown using the structure prior to MD simulations. Displacement magnitudes are represented on a white-yellow-red scale.[7] Significant vectors with length ≥20 pm are displayed as green arrows with a scale-up factor of 5.[8]

### Section S5B: Substitution of P2′ residue

**Table S5B.1**: The average differences in the five active-site distances ($\Delta d = d_{neq} - d_{eqm}$) following the P2′ Asn to Ala substitution in s05. Positive and negative differences with magnitudes > 0.1 Å are in green and red, respectively.

| Active Site | s05: P2' N->A | | |
|---|---|---|---|
| **Deprot** | t (ps) | Av Diff (Å) | SEM (Å) |
| | 0.05 | 0.00 | 0.00 |
| | 0.1 | 0.00 | 0.00 |
| | 0.25 | 0.00 | 0.00 |
| | 0.5 | 0.00 | 0.01 |
| | 1 | -0.02 | 0.03 |
| | 2 | 0.00 | 0.05 |
| | 3 | -0.07 | 0.06 |
| | 5 | -0.05 | 0.07 |
| | 10 | -0.04 | 0.05 |
| | 20 | 0.03 | 0.06 |
| | 30 | 0.01 | 0.08 |
| | 50 | 0.01 | 0.07 |
| | 100 | -0.01 | 0.08 |
| | 300 | -0.01 | 0.07 |
| | 500 | -0.02 | 0.07 |
| | 1000 | 0.12 | 0.07 |
| **Nuc** | t (ps) | Av Diff (Å) | SEM (Å) |
| | 0.05 | 0.00 | 0.00 |
| | 0.1 | 0.00 | 0.00 |
| | 0.25 | 0.00 | 0.00 |
| | 0.5 | 0.00 | 0.01 |
| | 1 | 0.01 | 0.01 |
| | 2 | -0.02 | 0.02 |
| | 3 | 0.00 | 0.02 |
| | 5 | -0.01 | 0.02 |
| | 10 | -0.02 | 0.03 |
| | 20 | 0.00 | 0.04 |
| | 30 | 0.00 | 0.04 |
| | 50 | -0.04 | 0.04 |
| | 100 | 0.01 | 0.05 |
| | 300 | 0.04 | 0.05 |
| | 500 | 0.03 | 0.04 |
| | 1000 | -0.04 | 0.05 |
| **OH145** | t (ps) | Av Diff (Å) | SEM (Å) |
| | 0.05 | 0.00 | 0.00 |
| | 0.1 | -0.01 | 0.00 |
| | 0.25 | -0.01 | 0.00 |
| | 0.5 | 0.00 | 0.01 |
| | 1 | -0.03 | 0.02 |
| | 2 | -0.02 | 0.02 |
| | 3 | -0.01 | 0.02 |
| | 5 | 0.00 | 0.03 |
| | 10 | -0.01 | 0.02 |
| | 20 | -0.04 | 0.02 |
| | 30 | 0.00 | 0.03 |
| | 50 | -0.01 | 0.02 |
| | 100 | -0.02 | 0.03 |
| | 300 | -0.02 | 0.03 |
| | 500 | 0.00 | 0.02 |
| | 1000 | -0.01 | 0.03 |
| **OH144** | t (ps) | Av Diff (Å) | SEM (Å) |
| | 0.05 | 0.00 | 0.00 |
| | 0.1 | -0.02 | 0.00 |
| | 0.25 | -0.02 | 0.01 |
| | 0.5 | -0.02 | 0.01 |
| | 1 | -0.01 | 0.02 |
| | 2 | -0.04 | 0.03 |
| | 3 | -0.05 | 0.03 |
| | 5 | 0.02 | 0.04 |
| | 10 | 0.06 | 0.03 |
| | 20 | 0.00 | 0.03 |
| | 30 | 0.00 | 0.04 |
| | 50 | -0.01 | 0.03 |
| | 100 | -0.06 | 0.03 |
| | 300 | 0.00 | 0.04 |
| | 500 | 0.03 | 0.03 |
| | 1000 | -0.06 | 0.04 |
| **OH143** | t (ps) | Av Diff (Å) | SEM (Å) |
| | 0.05 | 0.01 | 0.00 |
| | 0.1 | 0.02 | 0.00 |
| | 0.25 | 0.03 | 0.01 |
| | 0.5 | -0.01 | 0.01 |
| | 1 | 0.01 | 0.02 |
| | 2 | 0.04 | 0.03 |
| | 3 | -0.01 | 0.03 |
| | 5 | -0.03 | 0.03 |
| | 10 | -0.01 | 0.03 |
| | 20 | 0.00 | 0.03 |
| | 30 | 0.03 | 0.03 |
| | 50 | -0.03 | 0.03 |
| | 100 | 0.02 | 0.03 |
| | 300 | 0.00 | 0.03 |
| | 500 | -0.01 | 0.03 |
| | 1000 | -0.02 | 0.03 |

**Table S5B.2**: The average differences in the donor-acceptor distances of HBs 1-5 following the P2′ Asn to Ala substitution in s05. Positive and negative differences with magnitudes > 0.1 Å are in green and red, respectively.

| HB1-5 | s05: P2' N->A | | |
|---|---|---|---|
| HB1 | t (ps) | Av Diff (Å) | SEM (Å) |
| | 0.05 | 0.00 | 0.00 |
| | 0.1 | 0.00 | 0.00 |
| | 0.25 | 0.00 | 0.00 |
| | 0.5 | -0.01 | 0.00 |
| | 1 | -0.01 | 0.02 |
| | 2 | 0.00 | 0.02 |
| | 3 | -0.01 | 0.03 |
| | 5 | 0.02 | 0.03 |
| | 10 | -0.03 | 0.03 |
| | 20 | -0.04 | 0.03 |
| | 30 | 0.01 | 0.03 |
| | 50 | 0.03 | 0.03 |
| | 100 | 0.01 | 0.03 |
| | 300 | -0.01 | 0.04 |
| | 500 | -0.01 | 0.05 |
| | 1000 | -0.03 | 0.05 |
| HB2 | t (ps) | Av Diff (Å) | SEM (Å) |
| | 0.05 | 0.00 | 0.00 |
| | 0.1 | 0.00 | 0.00 |
| | 0.25 | 0.00 | 0.00 |
| | 0.5 | 0.00 | 0.00 |
| | 1 | 0.01 | 0.01 |
| | 2 | -0.03 | 0.02 |
| | 3 | 0.03 | 0.02 |
| | 5 | -0.05 | 0.02 |
| | 10 | 0.02 | 0.02 |
| | 20 | -0.01 | 0.02 |
| | 30 | 0.00 | 0.02 |
| | 50 | 0.02 | 0.02 |
| | 100 | -0.01 | 0.02 |
| | 300 | 0.01 | 0.02 |
| | 500 | 0.03 | 0.02 |
| | 1000 | -0.02 | 0.02 |
| HB3 | t (ps) | Av Diff (Å) | SEM (Å) |
| | 0.05 | 0.00 | 0.00 |
| | 0.1 | 0.00 | 0.00 |
| | 0.25 | 0.00 | 0.00 |
| | 0.5 | 0.00 | 0.00 |
| | 1 | -0.01 | 0.01 |
| | 2 | -0.02 | 0.02 |
| | 3 | 0.02 | 0.01 |
| | 5 | -0.04 | 0.02 |
| | 10 | 0.01 | 0.01 |
| | 20 | -0.02 | 0.02 |
| | 30 | -0.01 | 0.01 |
| | 50 | 0.01 | 0.01 |
| | 100 | -0.01 | 0.01 |
| | 300 | 0.00 | 0.01 |
| | 500 | 0.02 | 0.02 |
| | 1000 | -0.01 | 0.02 |
| HB4 | t (ps) | Av Diff (Å) | SEM (Å) |
| | 0.05 | 0.00 | 0.00 |
| | 0.1 | 0.00 | 0.00 |
| | 0.25 | 0.00 | 0.00 |
| | 0.5 | 0.00 | 0.01 |
| | 1 | 0.01 | 0.02 |
| | 2 | -0.02 | 0.03 |
| | 3 | 0.01 | 0.03 |
| | 5 | 0.05 | 0.04 |
| | 10 | -0.07 | 0.04 |
| | 20 | 0.00 | 0.05 |
| | 30 | 0.01 | 0.05 |
| | 50 | 0.02 | 0.07 |
| | 100 | 0.05 | 0.08 |
| | 300 | 0.08 | 0.11 |
| | 500 | 0.11 | 0.11 |
| | 1000 | -0.24 | 0.13 |
| HB5 | t (ps) | Av Diff (Å) | SEM (Å) |
| | 0.05 | 0.00 | 0.00 |
| | 0.1 | 0.00 | 0.00 |
| | 0.25 | -0.01 | 0.00 |
| | 0.5 | 0.00 | 0.01 |
| | 1 | -0.06 | 0.02 |
| | 2 | 0.03 | 0.02 |
| | 3 | -0.04 | 0.02 |
| | 5 | -0.01 | 0.02 |
| | 10 | -0.03 | 0.03 |
| | 20 | -0.05 | 0.03 |
| | 30 | -0.01 | 0.02 |
| | 50 | 0.02 | 0.03 |
| | 100 | -0.04 | 0.03 |
| | 300 | -0.03 | 0.02 |
| | 500 | -0.04 | 0.03 |
| | 1000 | -0.01 | 0.03 |

**Table S5B.3**: The average differences in the donor-acceptor distances of HBs 8-12 following the P2′ Asn to Ala substitution in s05. Positive and negative differences with magnitudes > 0.1 Å are in green and red, respectively.

| HB8-12 | s05: P2' N->A | | |
|---|---|---|---|
| HB8 | t (ps) | Av Diff (Å) | SEM (Å) |
| | 0.05 | 0.00 | 0.00 |
| | 0.1 | -0.01 | 0.00 |
| | 0.25 | -0.01 | 0.00 |
| | 0.5 | 0.00 | 0.01 |
| | 1 | -0.02 | 0.02 |
| | 2 | -0.02 | 0.02 |
| | 3 | -0.01 | 0.02 |
| | 5 | 0.00 | 0.03 |
| | 10 | -0.02 | 0.02 |
| | 20 | -0.03 | 0.02 |
| | 30 | 0.00 | 0.03 |
| | 50 | -0.01 | 0.02 |
| | 100 | -0.03 | 0.03 |
| | 300 | -0.02 | 0.03 |
| | 500 | 0.00 | 0.02 |
| | 1000 | 0.00 | 0.03 |
| HB9 | t (ps) | Av Diff (Å) | SEM (Å) |
| | 0.05 | 0.01 | 0.00 |
| | 0.1 | 0.01 | 0.00 |
| | 0.25 | 0.01 | 0.00 |
| | 0.5 | 0.00 | 0.01 |
| | 1 | 0.01 | 0.01 |
| | 2 | 0.02 | 0.02 |
| | 3 | 0.00 | 0.02 |
| | 5 | -0.01 | 0.02 |
| | 10 | 0.01 | 0.02 |
| | 20 | 0.01 | 0.02 |
| | 30 | 0.00 | 0.02 |
| | 50 | -0.01 | 0.02 |
| | 100 | 0.02 | 0.02 |
| | 300 | 0.02 | 0.02 |
| | 500 | 0.00 | 0.02 |
| | 1000 | -0.01 | 0.02 |
| HB10 | t (ps) | Av Diff (Å) | SEM (Å) |
| | 0.05 | 0.00 | 0.00 |
| | 0.1 | -0.01 | 0.00 |
| | 0.25 | 0.00 | 0.01 |
| | 0.5 | 0.00 | 0.01 |
| | 1 | 0.01 | 0.01 |
| | 2 | -0.04 | 0.02 |
| | 3 | 0.00 | 0.02 |
| | 5 | 0.02 | 0.02 |
| | 10 | -0.01 | 0.02 |
| | 20 | -0.03 | 0.02 |
| | 30 | 0.00 | 0.02 |
| | 50 | -0.01 | 0.02 |
| | 100 | -0.02 | 0.02 |
| | 300 | 0.00 | 0.02 |
| | 500 | -0.06 | 0.02 |
| | 1000 | 0.02 | 0.02 |
| HB11 | t (ps) | Av Diff (Å) | SEM (Å) |
| | 0.05 | -0.01 | 0.00 |
| | 0.1 | 0.01 | 0.00 |
| | 0.25 | -0.01 | 0.01 |
| | 0.5 | 0.00 | 0.01 |
| | 1 | 0.03 | 0.02 |
| | 2 | -0.01 | 0.02 |
| | 3 | 0.00 | 0.01 |
| | 5 | 0.01 | 0.02 |
| | 10 | 0.00 | 0.02 |
| | 20 | -0.01 | 0.02 |
| | 30 | 0.00 | 0.02 |
| | 50 | 0.02 | 0.02 |
| | 100 | 0.02 | 0.02 |
| | 300 | 0.00 | 0.02 |
| | 500 | 0.01 | 0.02 |
| | 1000 | 0.06 | 0.02 |
| HB12 | t (ps) | Av Diff (Å) | SEM (Å) |
| | 0.05 | 0.00 | 0.00 |
| | 0.1 | 0.00 | 0.00 |
| | 0.25 | 0.00 | 0.01 |
| | 0.5 | 0.01 | 0.01 |
| | 1 | 0.01 | 0.02 |
| | 2 | -0.05 | 0.02 |
| | 3 | 0.02 | 0.02 |
| | 5 | -0.02 | 0.03 |
| | 10 | 0.01 | 0.03 |
| | 20 | 0.03 | 0.03 |
| | 30 | 0.00 | 0.03 |
| | 50 | 0.02 | 0.03 |
| | 100 | 0.03 | 0.04 |
| | 300 | 0.05 | 0.03 |
| | 500 | 0.01 | 0.04 |
| | 1000 | 0.07 | 0.04 |

**Table S5B.4**: The ten most perturbed Cα atoms in the $M^{pro}$-peptide complex in terms of magnitude of the average displacement vector *v*, at selected time points following the P2′ Asn to Ala substitution in s05. Only displacements ≥ 0.01 Å are shown.

| s05 t(ps) | CA Rank | Atom | Chain | v (Å) | SEM (Å) |
|---|---|---|---|---|---|
| **0.05** | 1 | 8ALA_CA | C | 0.02 | 0.00 |
| | 2 | 143GLY_CA | A | 0.01 | 0.00 |
| | 3 | 7ASN_CA | C | 0.01 | 0.00 |
| | 4 | 9GLU_CA | C | 0.01 | 0.00 |
| | 5 | | | | |
| | 6 | | | | |
| | 7 | | | | |
| | 8 | | | | |
| | 9 | | | | |
| | 10 | | | | |
| **0.5** | 1 | 8ALA_CA | C | 0.04 | 0.01 |
| | 2 | 9GLU_CA | C | 0.04 | 0.01 |
| | 3 | 26THR_CA | A | 0.03 | 0.01 |
| | 4 | 142ASN_CA | A | 0.03 | 0.01 |
| | 5 | 143GLY_CA | A | 0.03 | 0.01 |
| | 6 | 144SER_CA | A | 0.03 | 0.01 |
| | 7 | 27LEU_CA | A | 0.02 | 0.00 |
| | 8 | 46SER_CA | A | 0.02 | 0.00 |
| | 9 | 47GLU_CA | A | 0.02 | 0.00 |
| | 10 | 119ASN_CA | A | 0.02 | 0.01 |
| **5** | 1 | 251GLY_CA | B | 0.17 | 0.04 |
| | 2 | 252PRO_CA | B | 0.17 | 0.04 |
| | 3 | 183GLY_CA | B | 0.14 | 0.04 |
| | 4 | 50LEU_CA | B | 0.13 | 0.04 |
| | 5 | 51ASN_CA | B | 0.13 | 0.04 |
| | 6 | 235MET_CA | B | 0.13 | 0.04 |
| | 7 | 11SER_CA | C | 0.13 | 0.07 |
| | 8 | 218TRP_CA | A | 0.12 | 0.04 |
| | 9 | 221ASN_CA | A | 0.12 | 0.04 |
| | 10 | 222ARG_CA | A | 0.12 | 0.05 |
| **50** | 1 | 11SER_CA | C | 0.28 | 0.13 |
| | 2 | 277ASN_CA | B | 0.24 | 0.10 |
| | 3 | 215GLY_CA | B | 0.23 | 0.07 |
| | 4 | 278GLY_CA | B | 0.23 | 0.09 |
| | 5 | 279ARG_CA | B | 0.21 | 0.07 |
| | 6 | 50LEU_CA | A | 0.20 | 0.06 |
| | 7 | 23GLY_CA | B | 0.20 | 0.06 |
| | 8 | 24THR_CA | B | 0.20 | 0.07 |
| | 9 | 275GLY_CA | B | 0.20 | 0.08 |
| | 10 | 276MET_CA | B | 0.19 | 0.07 |
| **500** | 1 | 278GLY_CA | A | 0.26 | 0.11 |
| | 2 | 191ALA_CA | B | 0.26 | 0.10 |
| | 3 | 23GLY_CA | B | 0.25 | 0.07 |
| | 4 | 215GLY_CA | B | 0.24 | 0.09 |
| | 5 | 155ASP_CA | A | 0.23 | 0.06 |
| | 6 | 215GLY_CA | A | 0.23 | 0.08 |
| | 7 | 277ASN_CA | A | 0.23 | 0.11 |
| | 8 | 276MET_CA | A | 0.22 | 0.07 |
| | 9 | 190THR_CA | B | 0.22 | 0.07 |
| | 10 | 252PRO_CA | B | 0.22 | 0.08 |
| **1000** | 1 | 11SER_CA | C | 0.81 | 0.22 |
| | 2 | 10LEU_CA | C | 0.47 | 0.11 |
| | 3 | 9GLU_CA | C | 0.39 | 0.07 |
| | 4 | 276MET_CA | A | 0.24 | 0.10 |
| | 5 | 215GLY_CA | A | 0.23 | 0.09 |
| | 6 | 194ALA_CA | B | 0.21 | 0.07 |
| | 7 | 215GLY_CA | B | 0.21 | 0.11 |
| | 8 | 55GLU_CA | B | 0.20 | 0.09 |
| | 9 | 273GLN_CA | B | 0.20 | 0.10 |
| | 10 | 274ASN_CA | B | 0.20 | 0.11 |

**Table S5B.5**: The ten most perturbed non-hydrogen atoms in the M^pro^-peptide complex in terms of magnitude of the average displacement vector *v*, at selected time points following the P2′ Asn to Ala substitution in s05.

| s05 t(ps) | Non-H Rank | Atom | Chain | v (Å) | SEM (Å) |
|---|---|---|---|---|---|
| **0.05** | 1 | 8ALA_CB | C | 0.05 | 0.01 |
| | 2 | 8ALA_N | C | 0.02 | 0.01 |
| | 3 | 8ALA_CA | C | 0.02 | 0.00 |
| | 4 | 119ASN_OD1 | A | 0.02 | 0.00 |
| | 5 | 8ALA_C | C | 0.02 | 0.00 |
| | 6 | 9GLU_CB | C | 0.01 | 0.00 |
| | 7 | 8ALA_O | C | 0.01 | 0.00 |
| | 8 | 7ASN_O | C | 0.01 | 0.00 |
| | 9 | 7ASN_C | C | 0.01 | 0.00 |
| | 10 | 9GLU_N | C | 0.01 | 0.00 |
| **0.5** | 1 | 119ASN_OD1 | A | 0.11 | 0.03 |
| | 2 | 9GLU_O | C | 0.10 | 0.02 |
| | 3 | 8ALA_CB | C | 0.08 | 0.03 |
| | 4 | 7ASN_O | C | 0.06 | 0.02 |
| | 5 | 119ASN_ND2 | A | 0.06 | 0.03 |
| | 6 | 9GLU_C | C | 0.05 | 0.01 |
| | 7 | 9GLU_CB | C | 0.05 | 0.01 |
| | 8 | 46SER_OG | A | 0.05 | 0.01 |
| | 9 | 9GLU_CA | C | 0.04 | 0.01 |
| | 10 | 19GLN_NE2 | A | 0.04 | 0.02 |
| **5** | 1 | 11SER_OC1 | C | 0.31 | 0.10 |
| | 2 | 47GLU_OE2 | A | 0.31 | 0.09 |
| | 3 | 252PRO_CG | B | 0.27 | 0.06 |
| | 4 | 47GLU_OE1 | B | 0.26 | 0.11 |
| | 5 | 107GLN_NE2 | B | 0.25 | 0.09 |
| | 6 | 256GLN_NE2 | A | 0.25 | 0.07 |
| | 7 | 142ASN_ND2 | B | 0.24 | 0.08 |
| | 8 | 223PHE_CZ | B | 0.23 | 0.08 |
| | 9 | 223PHE_CE2 | B | 0.22 | 0.08 |
| | 10 | 11SER_C | C | 0.22 | 0.09 |
| **50** | 1 | 11SER_OC2 | C | 0.56 | 0.17 |
| | 2 | 11SER_C | C | 0.42 | 0.16 |
| | 3 | 279ARG_NH2 | B | 0.41 | 0.12 |
| | 4 | 11SER_OC1 | C | 0.38 | 0.19 |
| | 5 | 256GLN_NE2 | A | 0.36 | 0.11 |
| | 6 | 279ARG_CZ | B | 0.35 | 0.11 |
| | 7 | 217ARG_NH2 | B | 0.34 | 0.14 |
| | 8 | 279ARG_NH1 | B | 0.34 | 0.12 |
| | 9 | 82MET_CE | A | 0.33 | 0.08 |
| | 10 | 217ARG_NH1 | A | 0.32 | 0.11 |
| **500** | 1 | 235MET_CE | A | 0.54 | 0.23 |
| | 2 | 72ASN_OD1 | B | 0.50 | 0.19 |
| | 3 | 256GLN_NE2 | A | 0.49 | 0.16 |
| | 4 | 306GLN_OE1 | B | 0.48 | 0.17 |
| | 5 | 90LYS_NZ | B | 0.46 | 0.15 |
| | 6 | 277ASN_OD1 | A | 0.44 | 0.18 |
| | 7 | 277ASN_ND2 | A | 0.44 | 0.18 |
| | 8 | 248ASP_OD2 | A | 0.43 | 0.13 |
| | 9 | 279ARG_NH1 | A | 0.43 | 0.19 |
| | 10 | 306GLN_OE1 | A | 0.43 | 0.17 |
| **1000** | 1 | 11SER_OG | C | 1.06 | 0.29 |
| | 2 | 11SER_CB | C | 0.99 | 0.28 |
| | 3 | 11SER_CA | C | 0.81 | 0.22 |
| | 4 | 11SER_C | C | 0.70 | 0.24 |
| | 5 | 9GLU_O | C | 0.69 | 0.10 |
| | 6 | 11SER_OC1 | C | 0.68 | 0.24 |
| | 7 | 11SER_OC2 | C | 0.68 | 0.28 |
| | 8 | 11SER_N | C | 0.66 | 0.18 |
| | 9 | 235MET_CE | A | 0.66 | 0.25 |
| | 10 | 9GLU_OE2 | C | 0.66 | 0.14 |

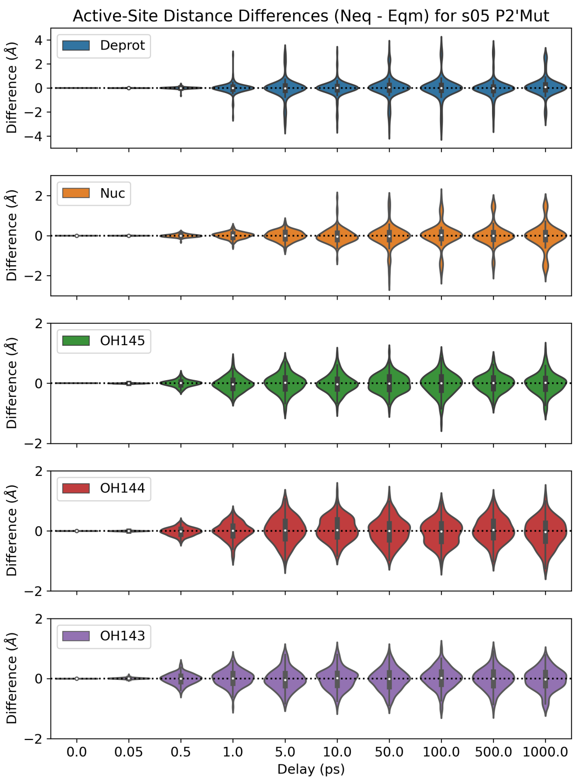

**Figure S5B.1**: Distributions of the distance differences in the five active-site distances ($\Delta d = d_{\text{neq}} - d_{\text{eqm}}$) following the P2′ Asn to Ala substitution in s05.

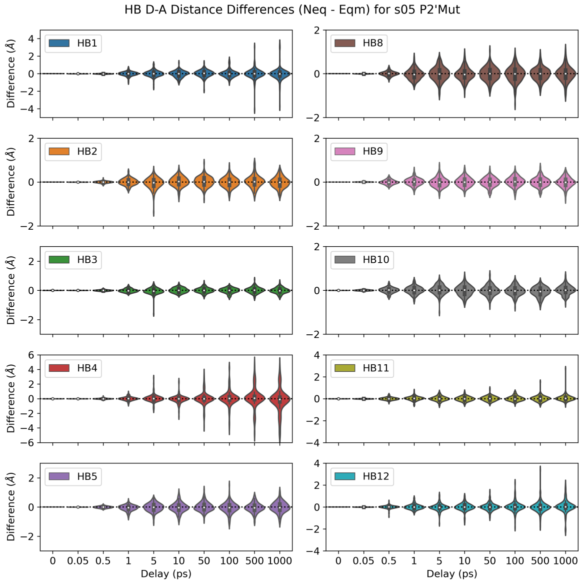

**Figure S5B.2**: Distributions of the distance differences in the ten donor-acceptor distances of HBs 1-5 and 8-12 following the P2′ Asn to Ala substitution in s05.

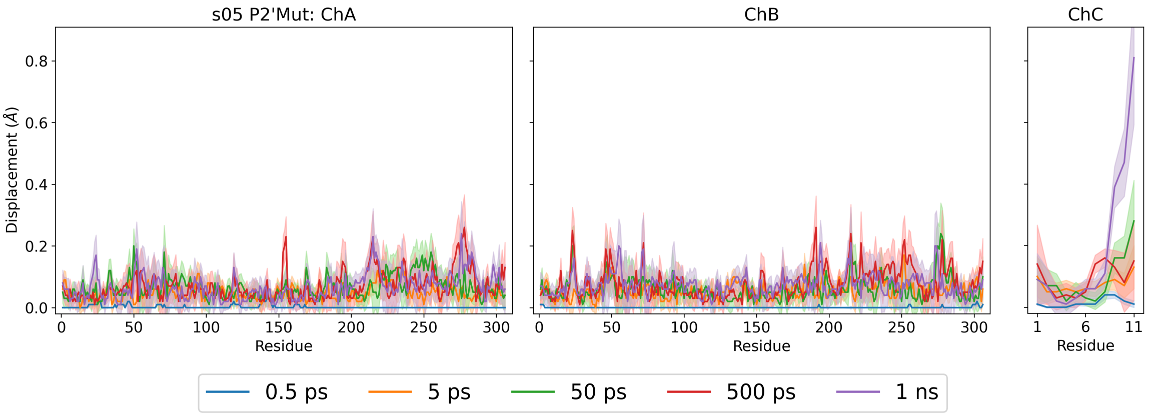

**Figure S5B.3**: Magnitudes of the average Cα displacement vectors, with ± SEM shown as shaded regions, following the P2′ Asn to Ala substitution in s05.

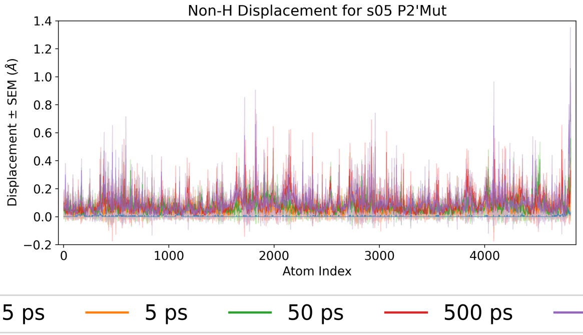

**Figure S5B.4**: Magnitudes of the average non-hydrogen atom displacement vectors in the M$^{pro}$-peptide complex, with ± SEM shown as shaded regions, following the P2′ Asn to Ala substitution in s05.

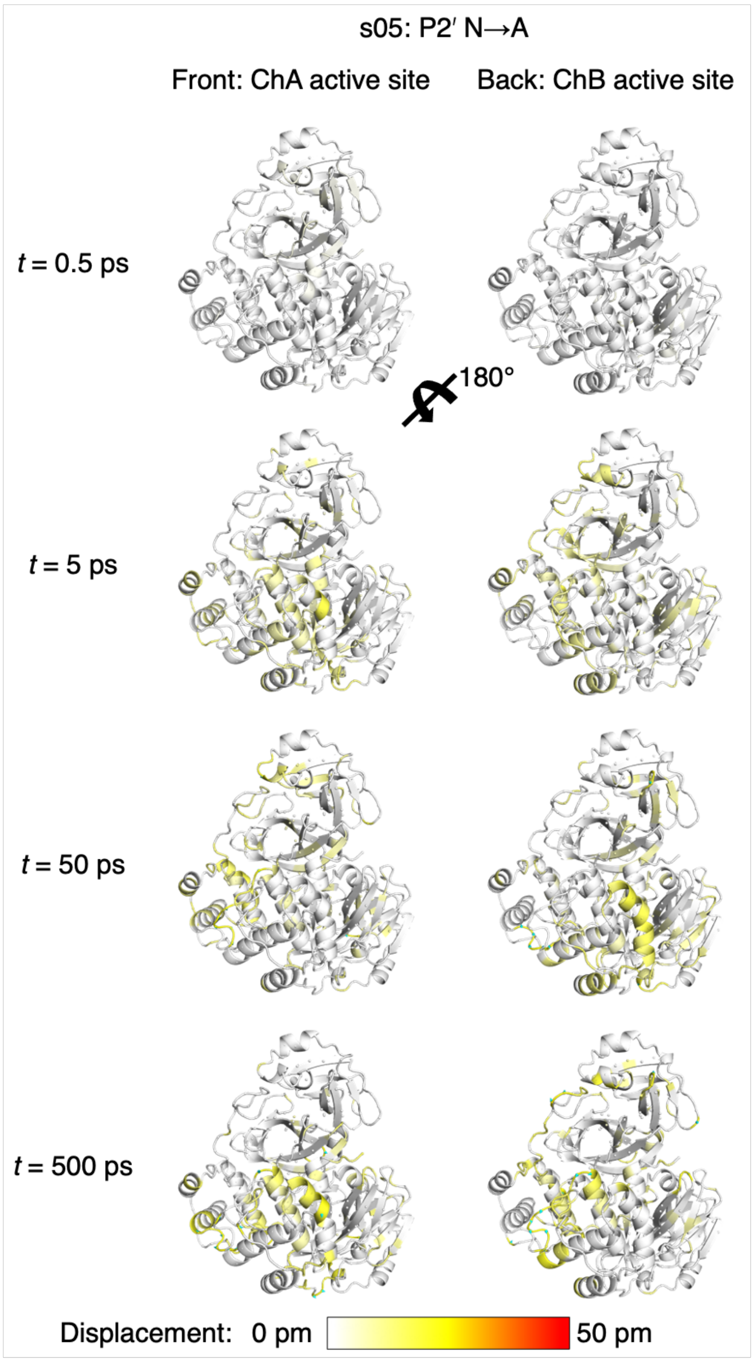

**Figure S5B.5**: Views of the M$^{pro}$ response to the P2′ Asn to Ala substitution in s05 from averaging Cα displacement vectors, shown using the structure prior to MD simulations. Displacement magnitudes are represented on a white-yellow-red scale.[7] Significant vectors with length ≥20 pm are displayed as cyan arrows with a scale-up factor of 5.[8]

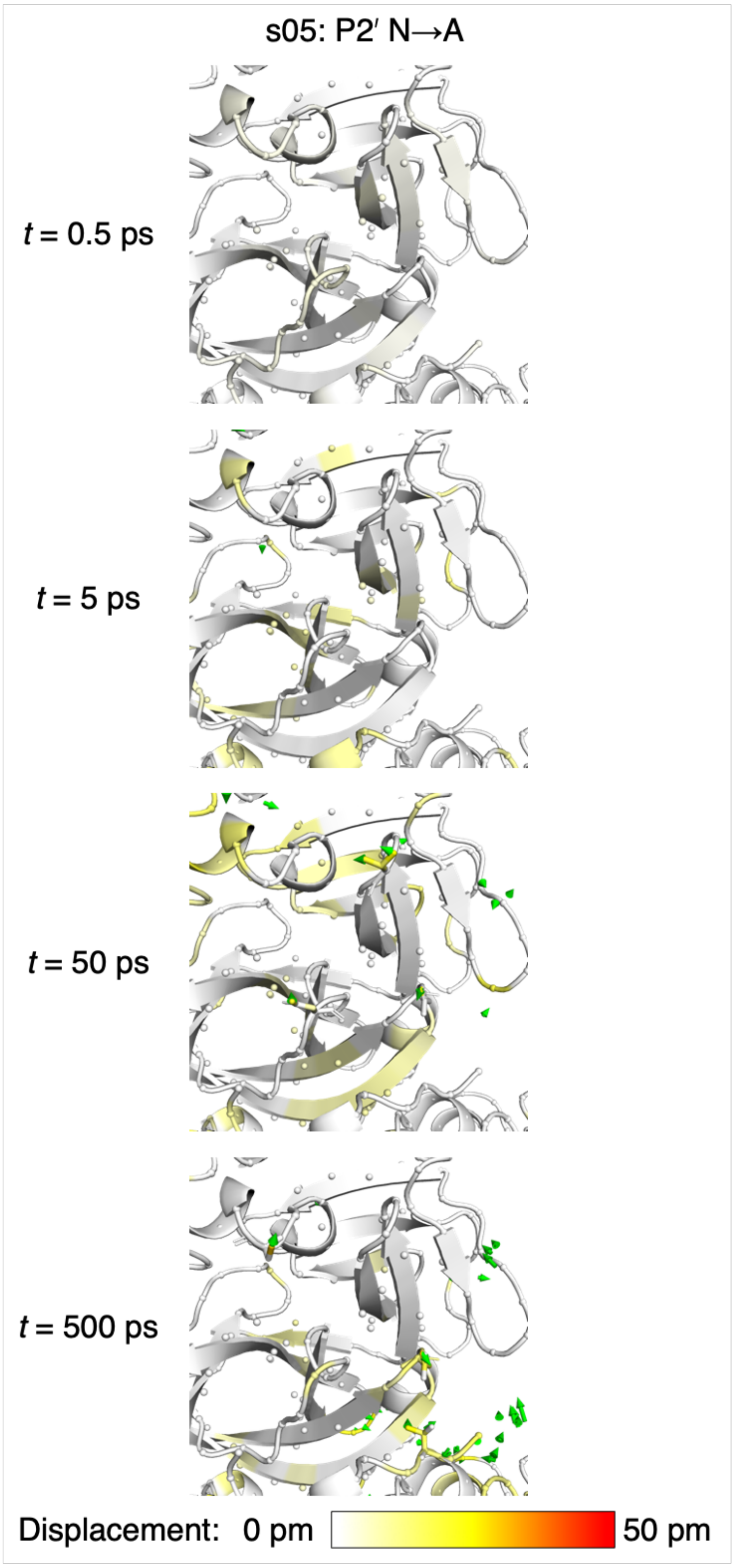

**Figure S5B.6**: Views of the M^pro response to the P2′ Asn to Ala substitution in s05 from averaging non-hydrogen atom displacement vectors, shown using the structure prior to MD simulations, with a focus on the S2′ subsite. Displacement magnitudes are represented on a white-yellow-red scale.[7] Significant vectors with length ≥20 pm are displayed as green arrows with a scale-up factor of 5.[8] Residues that show such significant displacements and that are within 10 Å of the substrate P2′ residue are shown as sticks.

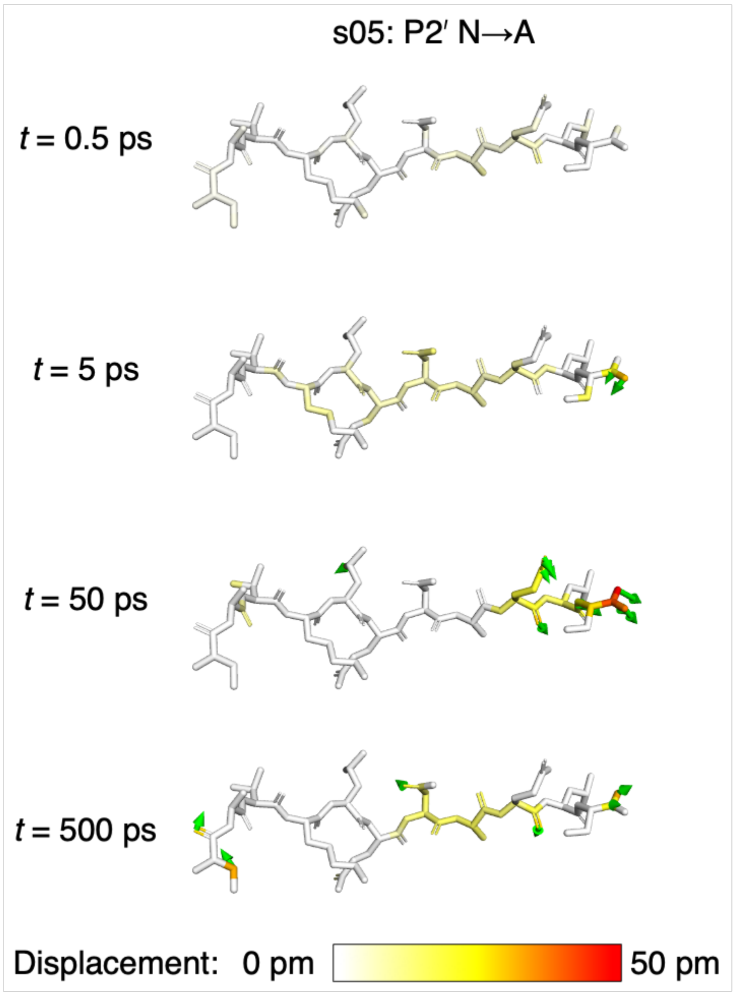

**Figure S5B.7**: Views of the peptide response to the P2′ Asn to Ala substitution in s05 from averaging non-hydrogen atom displacement vectors, shown using the structure prior to MD simulations. Displacement magnitudes are represented on a white-yellow-red scale.[7] Significant vectors with length ≥20 pm are displayed as green arrows with a scale-up factor of 5.[8]

### Section SM: Supplementary Methods

**Table SM.1**: Assignment of ionic and protonation states of titratable residues of SARS-CoV-2 M$^{pro}$ (PDB 6YB7)[3] employed in this study. The setup is identical to that previously reported.[1, 2] Neutral histidine residues that are N$\delta$-protonated and N$\epsilon$-protonated are referred to as "HID" and "HIE" respectively, according to AMBER force field nomenclature.[9] None of the N-terminal (NT) and C-terminal (CT) in M$^{pro}$ and peptides are capped.

| Residue | LYS (+1) | ARG (+1) | ASP (-1) | GLU (-1) | HID (0) | HIE (0) | NT (+1) | CT (-1) | **TOTAL** |
|---|---|---|---|---|---|---|---|---|---|
| ChA/ChB | 5 | 4 | 33 | 14 | 41 | 64 | 1 | 306 | |
| | 12 | 40 | 34 | 47 | 80 | 163 | | | |
| | 61 | 60 | 48 | 55 | | 164 | | | |
| | 88 | 76 | 56 | 166 | | 172 | | | |
| | 90 | 105 | 92 | 178 | | 246 | | | |
| | 97 | 131 | 153 | 240 | | | | | |
| | 100 | 188 | 155 | 270 | | | | | |
| | 102 | 217 | 176 | 288 | | | | | |
| | 137 | 222 | 187 | 290 | | | | | |
| | 236 | 279 | 197 | | | | | | |
| | 269 | 298 | 216 | | | | | | |
| | | | 229 | | | | | | |
| | | | 245 | | | | | | |
| | | | 248 | | | | | | |
| | | | 263 | | | | | | |
| | | | 289 | | | | | | |
| | | | 295 | | | | | | |
| **Charge** | **+11** | **+11** | **-17** | **-9** | **0** | **0** | **+1** | **-1** | **-4** |
| Peptides | | | | | | | | | |
| s01 | 11 | 10 | | | | | 1 | 11 | |
| **Charge** | **+1** | **+1** | | | | | **+1** | **-1** | **+2** |
| s02 | 10 | 11 | | | | | 1 | 11 | |
| **Charge** | **+1** | **+1** | | | | | **+1** | **-1** | **+2** |
| p12 | 1 | | | | | | 1 | 11 | |
| **Charge** | **+1** | | | | | | **+1** | **-1** | **+1** |
| s01-QP1A | 11 | 10 | | | | | 1 | 11 | |
| **Charge** | **+1** | **+1** | | | | | **+1** | **-1** | **+2** |
| s05-QP1A | 4 | | | 9 | | | 1 | 11 | |
| **Charge** | **+1** | | | **-1** | | | **+1** | **-1** | **0** |
